# An Alternative Framework for Fluorescence Correlation Spectroscopy

**DOI:** 10.1101/426114

**Authors:** Sina Jazani, Ioannis Sgouralis, Omer M. Shafraz, Marcia Levitus, Sanjeevi Sivasankar, Steve Pressé

## Abstract

Fluorescence correlation spectroscopy (FCS), is a flexible and widely used tool routinely exploited for *in vivo* and *in vitro* applications. While FCS provides estimates of dynamical quantities, such as diffusion coefficients, it demands high signal to noise ratios and long time traces, typically in the minute range. In principle, the same information can be extracted from *µ*-s long time traces; however, an appropriate analysis method is missing. To overcome these limitations, we adapt novel tools inspired by Bayesian non-parametrics, which starts from the direct analysis of the observed photon counts. With this approach, we are able to analyze time traces, which are too short to be analyzed by existing methods, including FCS. Our new analysis extends the capability of single molecule fluorescence confocal microscopy based approaches, to probe processes several orders of magnitude faster in time and permits a reduction of phototoxic effects on living samples induced by long periods of light exposure.

## INTRODUCTION

Due to its flexibility and limited invasiveness for *in vivo* applications, single molecule fluorescence confocal microscopy^1–4^ is one of the most widely used experimental techniques of modern Biophysics. In this technique, fluorescent molecules are allowed to freely diffuse into a volume illuminated by a tightly focused laser beam of a conventional single-focus confocal setup. Molecular motion inside the illuminated volume generates fluctuations in the emitted fluorescence that is recorded and subsequently temporally autocorrelated^1–4^ or, jointly spatiotemporally autocorrelated^5–7^, to deduce physical quantities of interest. For example, fluorescence correlation spectroscopy (FCS)^1,2^ as well as complementary methods - such as Fluorescence Cross-Correlation Spectroscopy (FCCS)^8^, scanning FCS^9,10^, spot variation Fluorescence Correlation Spectroscopy (svFCS)^11^, Fluorescence Resonance Energy Transfer-Fluorescence Correlation Spectroscopy (FCS-FRET)^12,13^, etc - estimate diffusion coefficients, reaction kinetic, binding affinities and concentrations of labeled molecules^14,15^.

While single molecule fluorescence confocal microscopy data is acquired on the micro- to millisecond timescales (*µ*s-ms), fluorescence correlation methods typically require the analysis of long time traces, several seconds to tens of minutes long, depending on the molecular concentrations and emission properties of the fluorophores employed^16,17^. These traces, capturing multiple molecule traversals of the confocal volume, provide the statistics needed for the post-processing steps used in traditional FCS analysis^16^ (e.g. autocorrelation, and nonlinear fitting to theoretical curves). However, processing steps like these downgrade the quality of the available data and demand either relatively high concentrations or excessively long time traces to yield reliable estimates. The same downgrades are encountered even with less traditional FCS analyses, including Bayesian approaches^18–22^, that also rely on auto-correlations.

In principle, within milliseconds, for the fluorophore concentrations and confocal volumes used in most experiments,^1,2,23^ thousands of data points are already available. Accordingly, if one could, somehow, estimate diffusion coefficients within tens of ms with the same accuracy as FCS, one could hypothetically use tens of minutes worth of data to discriminate between very small differences in diffusion coefficients. Alternatively, one could opt for shorter traces in the first place and, in doing so, reduce the sample’s light exposure to only a few ms, thereby minimizing photo-toxic effects which remain a severe limitation of fluorescence microscopy^24–26^.

Exploiting data on ms timescales would require a method that, simultaneously and self-consistently, estimates the number of fluorescent molecules at any given time within the (inhomogenously) illuminated volume and deduce their dynamical properties based on their photon emissions, which, in turn, depend on their evolving location within the confocal volume. The mathematics to do so in a rigorous and efficient manner have, so far, been unavailable as analyzing ms traces would demand that we consider all possible populations of molecules responsible for the observed traces, their diffusion coefficients, and every possible location (and, thus, photon emission rate) of those molecules at any given time.

Indeed, with current technology, this global optimization is prohibitively computationally expensive. To wit, maximum likelihood approaches^15,27^, popular in a variety of applications, are excluded since they require that the, otherwise unknown, population of molecules in the confocal volume at any given time be specified in advance by other means. These considerations motivate an entirely new framework for FCS.

Here we introduce a novel approach that exploits Bayesian non-parametrics,^15,28,29^ a branch of Statistics first suggested^30^ in 1973 and only broadly popularized in physical applications over the last few years^15,28,29,31–37^. This approach allows us to account for an arbitrary number of molecules responsible for emitting detected photons. With the proposed method, we are able to estimate physical variables, otherwise determined from FCS, with: (i) significantly shorter time traces; and (ii) nearly single molecule resolution. Furthermore, our overall framework is generalizable, and can estimate not only diffusion coefficients and molecular populations but also track molecules through time as well as determine their molecular brightness and the background photon emission rate.

## RESULTS

The method we propose for the analysis of traces from single molecule fluorescence confocal microscopy follows the Bayesian paradigm. Within this paradigm,^15,27,29,38^ our goal is to estimate posterior probability distributions over unknown parameters such as diffusion coefficients as well as molecular populations over time.

In this section, we first demonstrate and validate our method by computing posterior distributions using synthetic (simulated) traces mimicking the properties of real single molecule fluorescence confocal experiments. We subsequently benchmark our estimates with traces from control *in vitro* experiments.

### A. Demonstration and validation with simulated data

To demonstrate the robustness of our method, we simulate fluorescent time traces under a broad range of: (i) numbers of labeled molecules in the effective volume, Fig. 1; (ii) diffusion coefficients, Fig. 2a; (iii) trace lengths, Fig. 2b; and (iv) molecular brightness, Fig. 3. Since, the majority of our time traces are too short to be meaningfully analyzed with traditional FCS, we compare our posteriors directly to the ground truth that we used in the simulations.

**FIG. 1.**
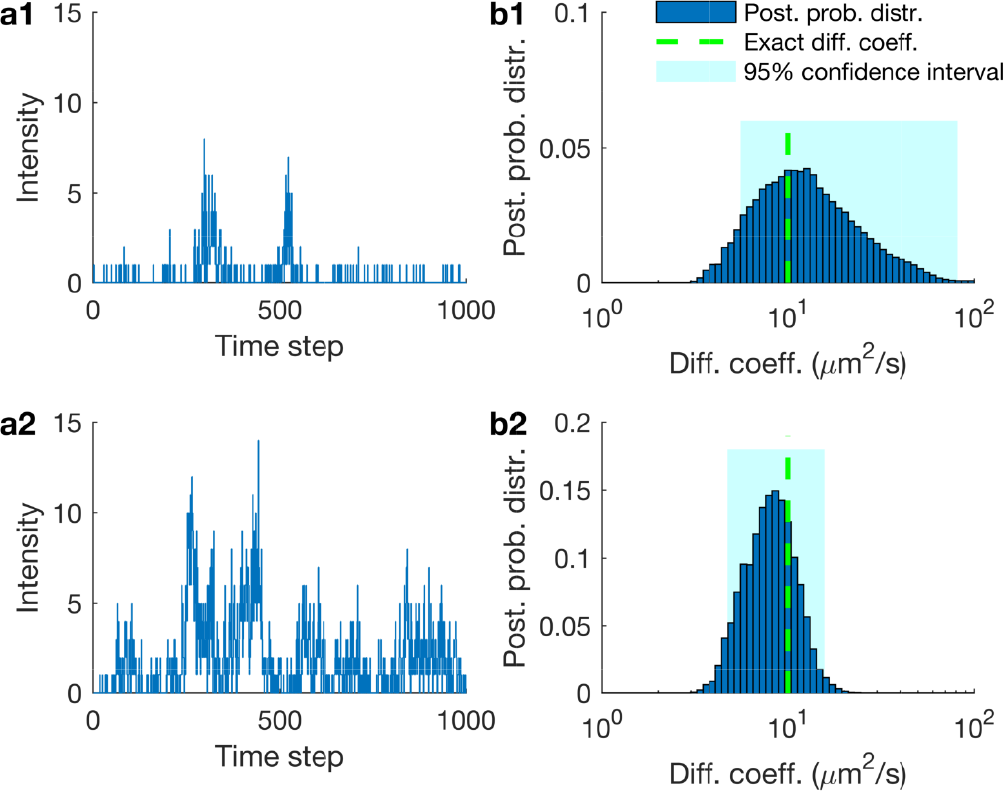
Testing the effects of the number of molecules inside the confocal volume on diffusion coefficient estimates. **(a1)** Synthetic fluorescent intensity trace produced by 1 molecule inside the confocal volume. For this simulation we used a molecular brightness of 5 × 10^4^ photons/s and a background photon emission rate of 10^3^ photons/s. **(b1)** Posterior probability distribution over the diffusion coefficient estimated from the trace in (a1). **(a2)** Synthetic fluorescent intensity trace produced by 5 molecules inside the confocal volume otherwise identical to (a1). **(b2)** Posterior probability distribution over the diffusion coefficient estimated from (a2). Traces shown in (a1) and (a2) are acquired at 100 *µ*s for a total of 100 ms and the highlighted regions in (b1) and (b2) represent the 95% confidence intervals. For clarity, the horizontal axis is shown in logarithmic scale.

**FIG. 2.**
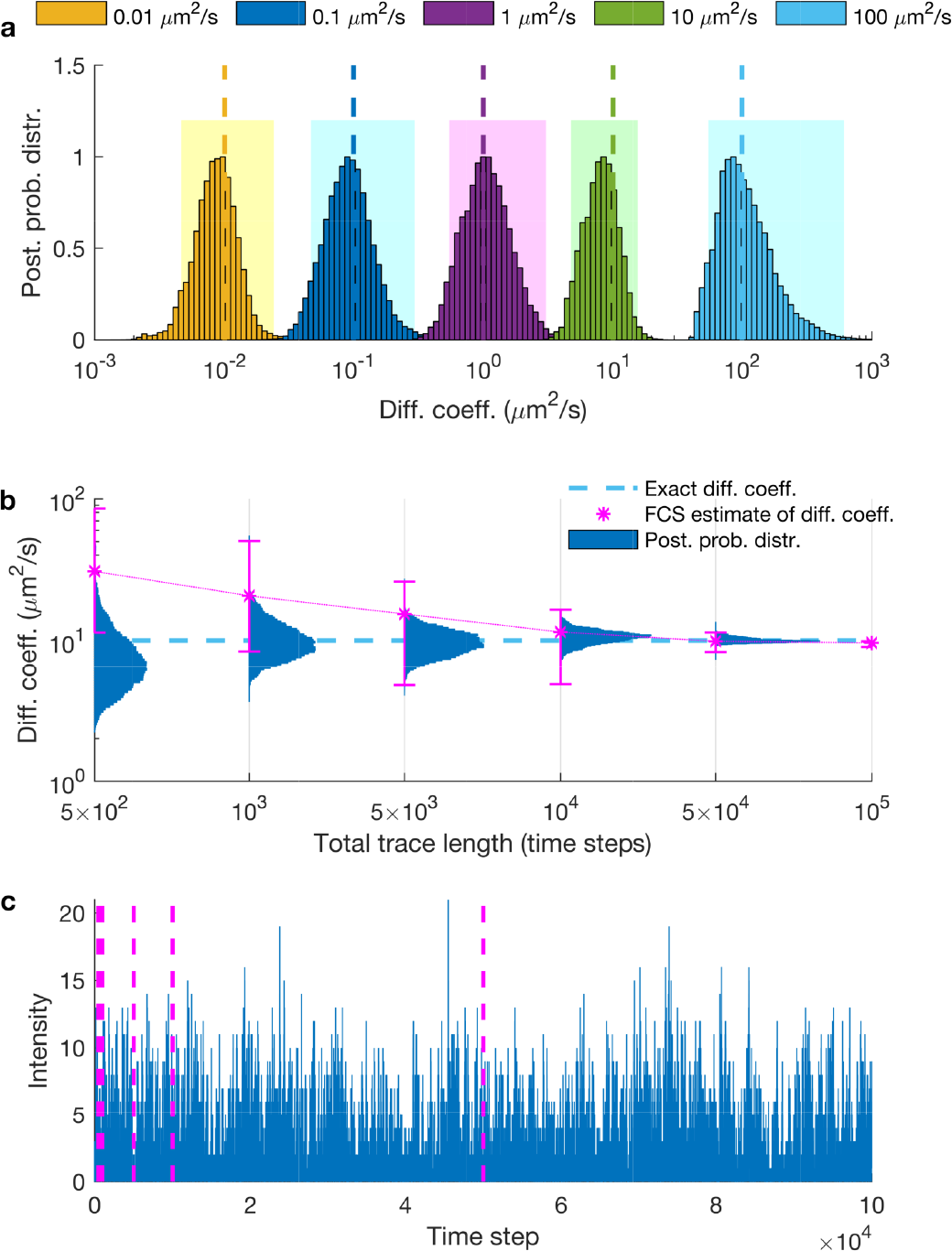
Testing the effects of diffusion coefficient and trace length on diffusion coefficient estimates. **(a)** Posterior probability distributions deduced from traces produced from molecules with diffusion coefficients of 0.01, 0.1, 1, 10 and 100 *µm*^2^/*s*. For clarity, posteriors are normalized to maximum 1 and the horizontal axis is shown in logarithmic scale. Shaded regions illustrate the 95% confidence intervals. **(b)** Posterior probability distributions deduced from traces acquired at 100 *µ*s with total trace lengths of 5 × 10^2^, 1 × 10^3^, 5 × 10^3^, 1 × 10^4^, 5 × 10^4^ and 1 × 10^5^ time steps. For comparison, exact values and FCS estimates are also shown and, for clarity, the vertical axis is shown in logarithmic scale. As can be seen from (b), it is typical for FCS to require ≈1000× more data than our method before estimating diffusion coefficients within 2× of the correct value. **(c)** The entire trace used to deduce diffusion coefficients in (a) and (b). Each segment marked by dashed-lines represents the portion used in (b). The molecular brightness and background photon emission rates used to generate the time traces used are 5 × 10^4^ and 10^3^ photons/s.

**FIG. 3.**
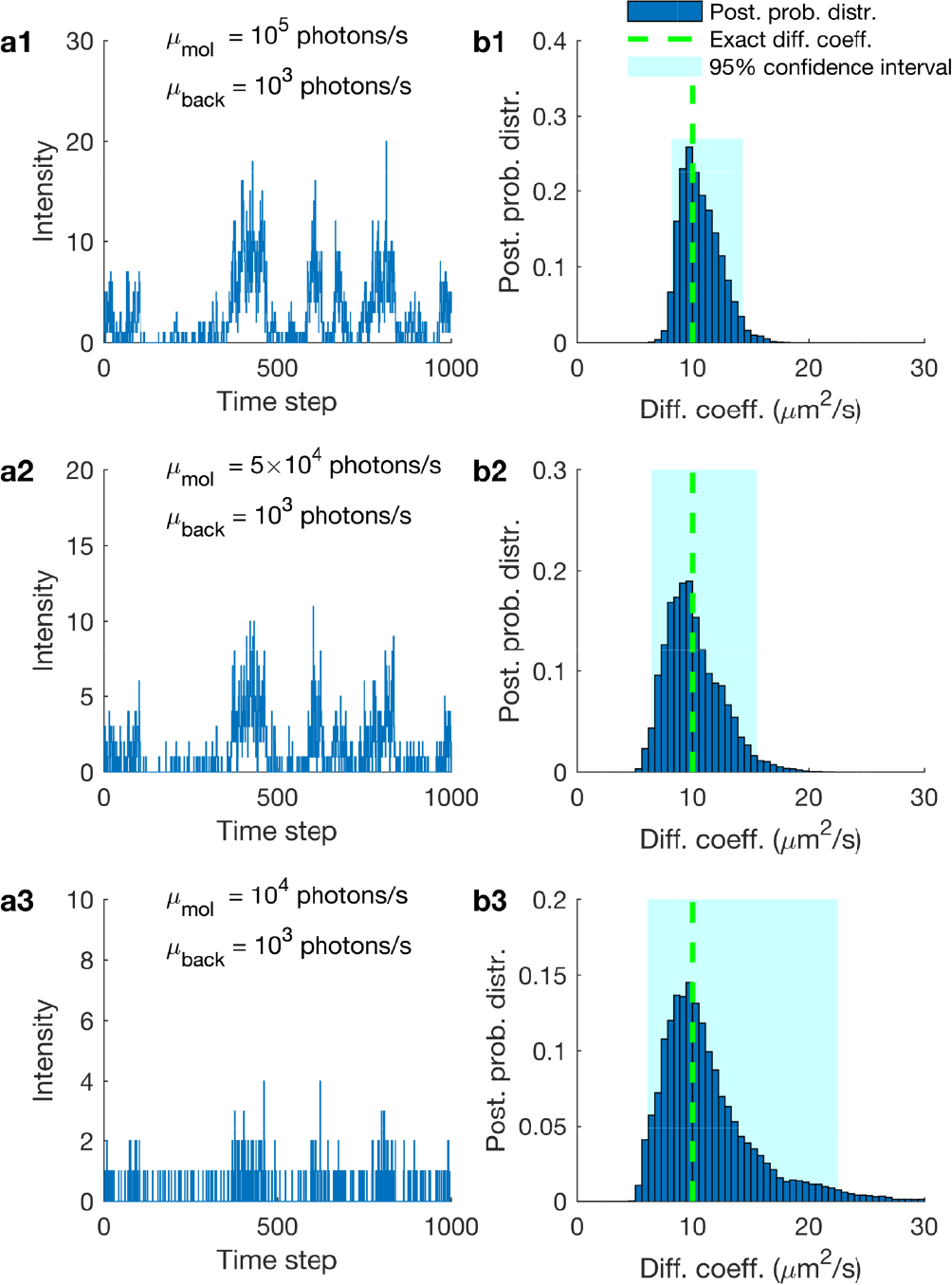
Testing the effects of molecular brightness on diffusion coefficient estimates. **(a)** Intensity traces produced by the same molecular trajectories under a molecular brightness of 10^5^, 5 × 10^4^ and 1 × 10^4^ photons/s and back-ground photon emission rate fixed at 10^3^ photons/s. **(b)** Posterior probability distributions and exact values of diffusion coefficients obtained from the corresponding traces. Shaded regions illustrate the 95% confidence intervals.

The posteriors we obtain, in all figures, are informed from the analysis of a single trace. In those, the breadth of the posterior (i.e. variance), which is a measure of the accuracy of our estimate, therefore indicates the uncertainty introduced by the finiteness of the data and the inherent noise in this single time trace.

To begin, in Fig. 1 we simulate a 3D Gaussian confocal volume of size (*ω*_*xy*_ = 0.3 *µm* and *ω*_*z*_ = 1.5 *µm*) and 1 molecule inside the effective volume (Fig. 1a1) or 5 molecules inside the effective volume (Fig. 1a2) diffusing at 10 *µ*m^2^/s for a total period of 100 ms.

As can be seen in Fig. 1, a low intensity leads to a wide estimate of the diffusion coefficient. However, the higher the intensity, the sharper (i.e. more conclusive) the estimate of the diffusion coefficient becomes (e.g. note a narrower posterior in Fig. 1b2 as compared to Fig. 1b1). Thus, diffusion coefficients are determined more accurately when the number of labeled molecules are higher. Accordingly, the most difficult data to analyze are those where concentrations of molecules are so low that, on average, only one molecule ventures, albeit rarely, into the effective region of the confocal volume where it can be appreciably excited. Put differently, for an equal length time trace, the posterior estimate over the diffusion coefficient is broader (i.e. less conclusive) for lower numbers of molecules inside the effective volume, Fig. 1b1, than it is for larger numbers of molecules, Fig. 1b2.

Following a similar reasoning, the slower a molecule diffuses, the more photons are collected, leading to a sharper posterior estimate of the corresponding diffusion coefficient, Fig. 2a. Likewise, the longer the trace is, Fig. 2b, or the greater the molecular brightness is, Fig. 3, the sharper the diffusion coefficient estimate becomes.

We emphasize that our definition of molecular brightness is based on the the maximum number of detected photons emitted from a single fluorophore when located at the center of the confocal volume and we provide more details in the supplementary materials.

In Fig. 3 we demonstrate the robustness of the diffusion coefficient estimates when varying the molecular brightness. While we keep the background photon emission rate fixed, we simulate gradually dimmer fluorophores such as those encountered in an experiment under lower laser powers, until the molecular signature is virtually lost. As can be seen, such traces lead to broader posterior estimates over diffusion coefficients, as one would expect, since these traces are associated with greater uncertainty. Also, as such traces lead to a weaker (i.e. less constraining) likelihood, the posterior resembles more closely the prior (not shown) and naturally starts to deviate from the exact value.

### B. Benchmarking on experimental data with elongated confocal volume shapes

Here we apply our method on experimental traces captured with an elongated confocal volume that we approximate by a cylinder. To do so, we apply our method on fluorescent beads (with average diameter of 45 nm) diffusing in water. We benchmark our estimated diffusion coefficients against the Stokes-Einstein prediction and results from FCS. In particular, Fig 4 illustrates our method’s performance in the analysis of traces too short to be meaningfully analyzed by FCS. Both the precise FCS formulation used here as well as additional results of our method on traces generated with free Cy3B dyes in glycerol/water mixtures with 70% glycerol can be found in the Supplementary materials.

**FIG. 4.**
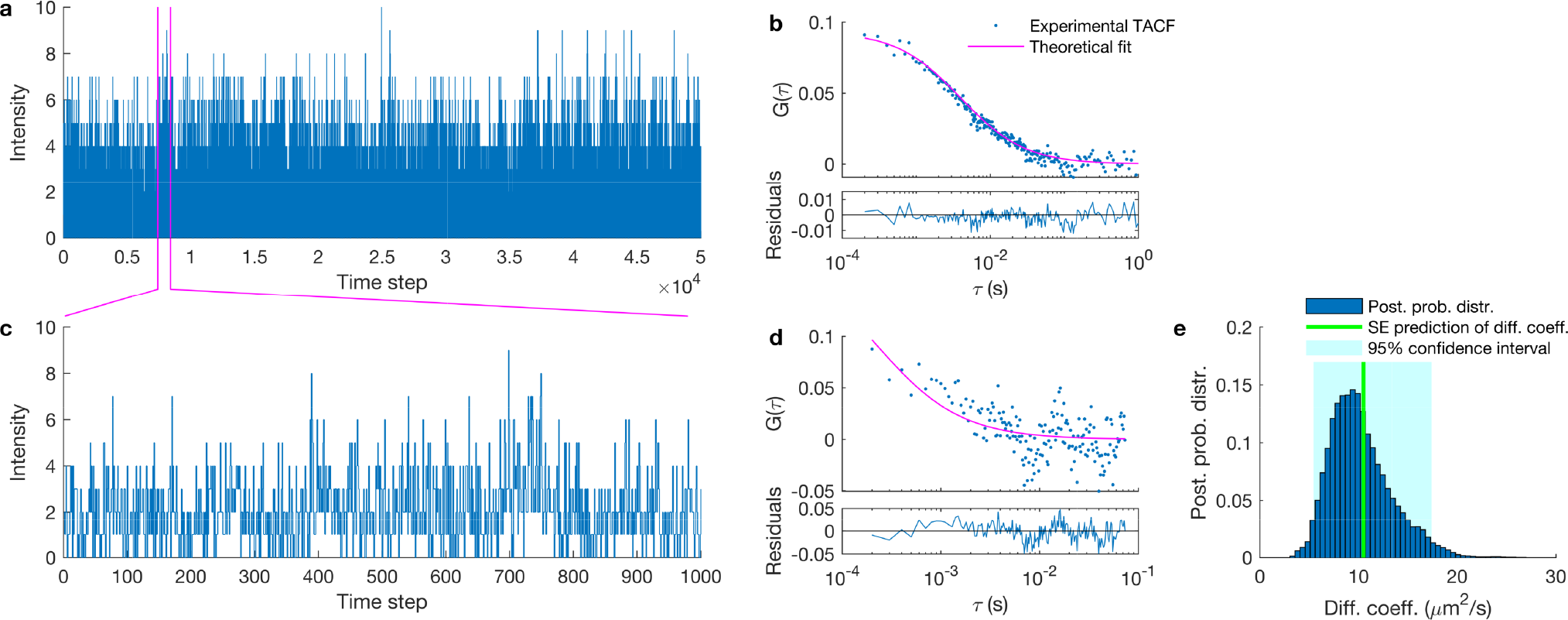
Diffusion coefficient estimates from experimental traces of free fluorescent beads using an elongated confocal volume. **(a)** Experimental fluorescent intensity trace used in FCS. **(b)** Auto-correlation curve of the trace in (a) and best theoretical fit. **(c)** Portion of the trace in (a) to be used as the input to FCS and our method. **(d)** Auto-correlation curve of trace in (c). **(e)** Posterior probability distribution over diffusion coefficient estimated from the trace in (c). Stokes-Einstein prediction are denoted by a green line. Traces shown in (a) and (c) are acquired in bins of 100 *µ*s for a total period of 5 seconds and 0.1 second respectively. The laser power used to generate the trace (a) is 100 *µW* (measured before the beam enters the objective). The estimate of diffusion coefficient resulting by autocorrelation fitting in (a) matches with the Stokes-Einstein prediction (i.e. 10.5 *µm*^2^/*s*); while in (d) is almost ten fold higher (~100 *µm*^2^/*s*).

### C. Benchmarking on experimental data with elliptical confocal volume shapes

#### 1. Fluorescent dyes

Next, we apply our method on experimental time traces derived from single molecule fluorescence confocal microscopy. In our setup, we monitor Cy3 dyes which diffuse freely in a mixture of water and glycerol. We benchmark our estimated diffusion coefficients against two values: those predicted by the Stokes-Einstein formula^40^, which is parametrized by physical quantities such as temperature and viscosity; and those estimated by FCS. To analyze the data using FCS, each time we used the full (6 min) trace available. This is by contrast to our the estimates provided by our method which are obtained on traces ~1000× shorter (100 ms).

In benchmarking, we obtained and analyzed measurements with different: (i) numbers of Cy3 dyes inside the effective volume (tuned by varying Cy3 concentrations); (ii) diffusion coefficients (tuned by adjusting the viscosity of the solution); and (iii) molecular brightness (tuned by adjusting the laser power). For example, in Fig. 5b1-b4, we illustrate the effect of different dye concentrations where a trace with stronger signal, anticipated when concentrations are higher, is expected to lead to better diffusion coefficient estimates (and thus sharper posteriors) on traces of equal length due to the higher number of labeled-molecules inside the confocal volume. Consistent with the synthetic data shown earlier, we obtain a broader posterior over diffusion coefficients when the number of dyes inside the effective volume is low and sharper posteriors for higher numbers of dyes.

**FIG. 5.**
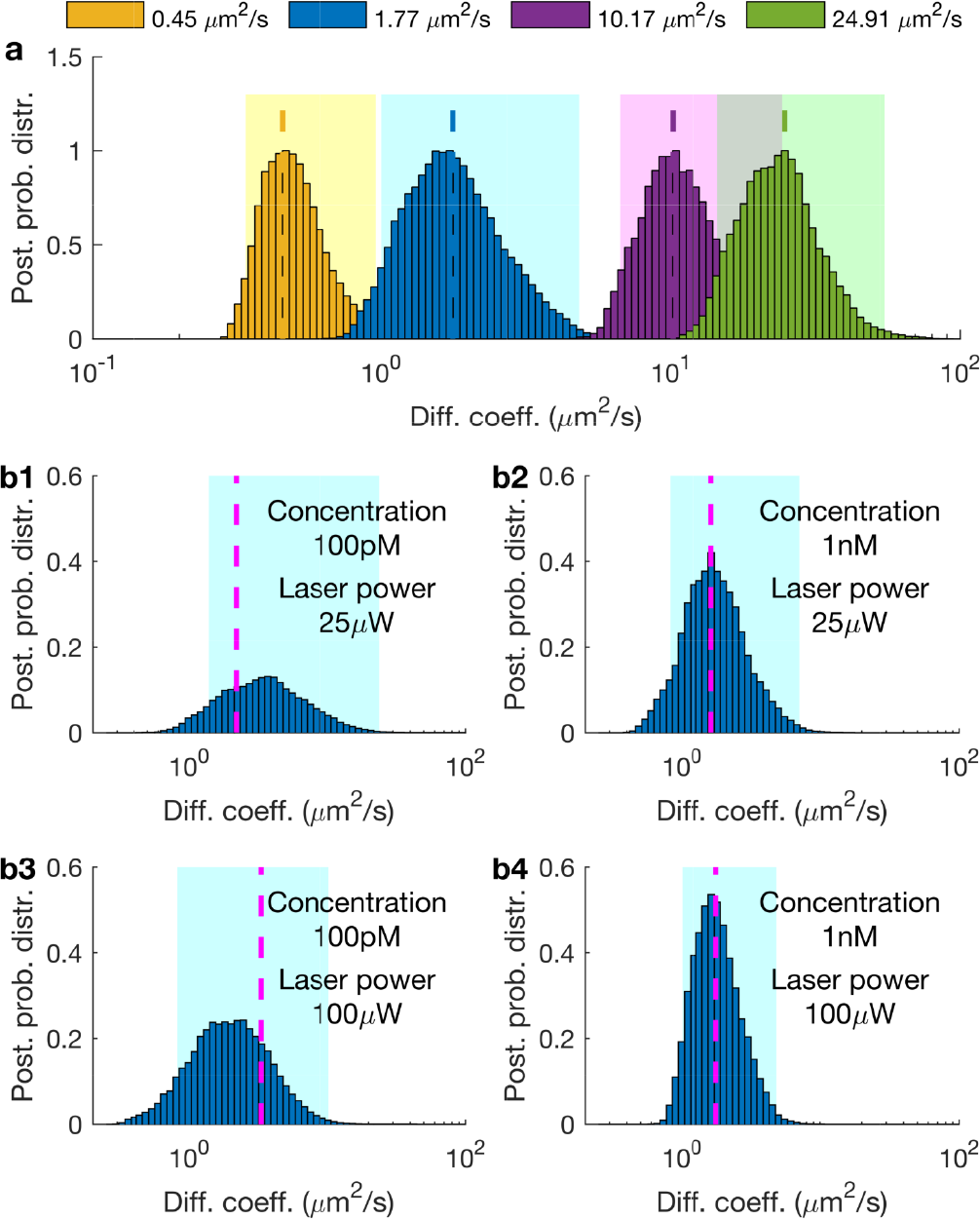
Estimating diffusion coefficients of free Cy3. **(a)** Posterior probability distributions of diffusion coefficients of free Cy3 in different concentrations of glycerol/water mixture. The legend labels the posteriors according to FCS estimates of long time traces. For clarity, posteriors are normalized to maximum 1 and the horizontal axis is shown in logarithmic scale. Also, the 95% confidence intervals are shown by highlighted regions. Posteriors are obtained from the analyses of time traces acquired at 100 *µ*s for total periods of 100 ms. Different diffusion coefficients are obtained by varying the amount of glycerol from 99% to 50% in the glycerol/water mixture. **(b)** Posterior probability distributions over the diffusion coefficients of traces generated by different laser powers (25 and 100 *µW*, respectively) with different concentrations of Cy3 (100 pM and 1 nM, respectively) in a glycerol/water mixture of 94% glycerol. For comparison, FCS estimates shown by dashed lines are obtained by traces, each of 6 min, i.e. ≈1000× longer than the segments used in our method.

Just as before, the slower a molecule diffuses, the more time it spends in the vicinity of the confocal volume, so the more photons are collected, thereby leading to sharper posterior estimates for the diffusion coefficient; see Fig. 5a.

A posterior’s sharpness depends strongly on the number of molecules in a time trace, their respective locations, and thus their photon emission rates. As the molecular population near the center of the confocal volume may exhibit strong fluctuations, the width of the posterior may also fluctuate from trace to trace, especially when the individual traces are short. Thus, the individual posteriors become sharper only *on average* as we move to higher numbers of molecules inside the effective volume or molecular brightness

#### 2. Labeled proteins

To test our method beyond free beads and dyes, we used labeled proteins, namely freely diffusing streptavidin labeled by Cy3. Similar to the previous cases, we tested a range of concentrations, diffusion coefficients, and laser powers. Figure 6 summarizes characteristic results and compares our analyses against the results of FCS applied on longer time traces. As can be seen, our method provides acceptable estimates of the diffusion coefficient even with ≈1000× less datapoints than FCS.

**FIG. 6.**
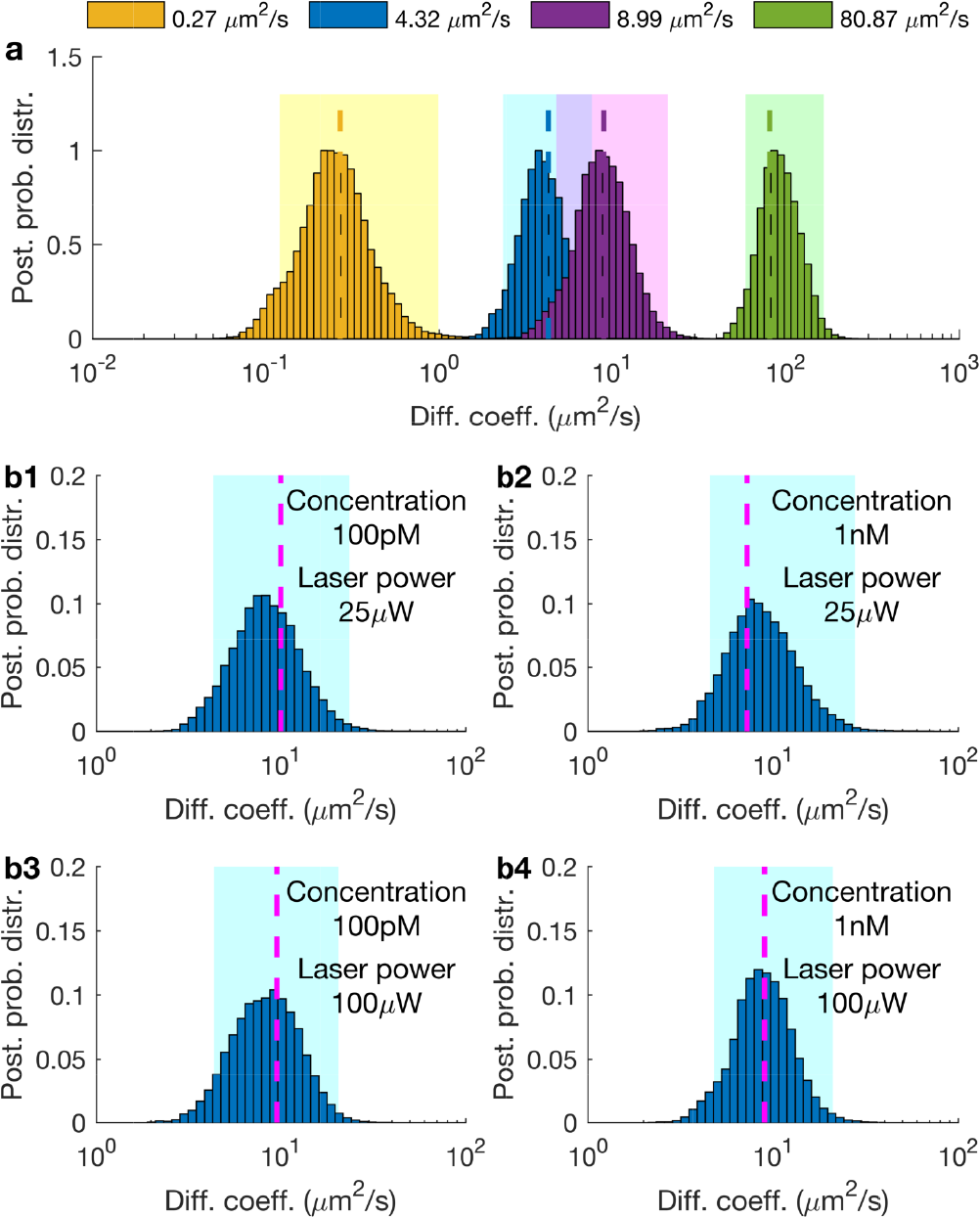
Estimating diffusion coefficients of free streptavidin. **(a)** Posterior probability distributions of diffusion coefficients of free streptavidin labeled by Cy3 in different concentrations of glycerol/water mixture. The legend labels the posteriors according to FCS estimates of long time traces. For clarity, posteriors are normalized to maximum 1 and the horizontal axis is shown in logarithmic scale. Also, the 95% confidence intervals are shown by highlighted regions. Posteriors are obtained from the analyses of time traces acquired at 100 *µ*s for total periods of 100 ms. Different diffusion coefficients are obtained by varying the amount of glycerol from 94% to 0% in the glycerol/water mixture. **(b)** Posterior probability distributions over the diffusion coefficients of traces generated by different laser powers (25 and 100 *µW*, respectively) with different concentrations of Cy3 (100 pM and 1 nM, respectively) in a glycerol/water mixture of 94% glycerol. For comparison, FCS estimates shown by dashed lines are obtained by traces, each of 6 min, i.e. ≈1000× longer than the segments used in our method.

### D. Additional results

For all cases described so far, we estimated more than just diffusion coefficients. For example, we also estimate the population of molecules contributing photons to the traces, their instantaneous photon emission rates and locations relative to the center of the confocal volume, as well as the background photon emission rate. A more detailed report of our estimates, with discussions of full joint posterior distributions, can be found in the Supplementary materials.

In addition to cases involving a single diffusion coefficient that we have considered thus far, our method can be generalized to treat multiple diffusion coefficients as well. To show this, we artificially mixed (summed) and analyzed experimental traces where dyes diffuse in different amounts of glycerol and so they exhibit different diffusion coefficients. On account of the additivity of photon emissions and detections, artificial mixing of traces allows us to obtain realistic traces of different diffusive species that can be analyzed as if they were diffusing simultaneously within the same confocal volume and separately as well. In Fig. 7, we compare the analysis of intensities created by mixing traces containing slow and fast diffusing Cy3 (94% and 75% glycerol/water, respectively). As can be seen, our estimates obtained under simultaneous diffusion compare favorably to the estimates under separate diffusion, indicating that our method can also identify robustly multiple diffusion coefficients at once.

**FIG. 7.**
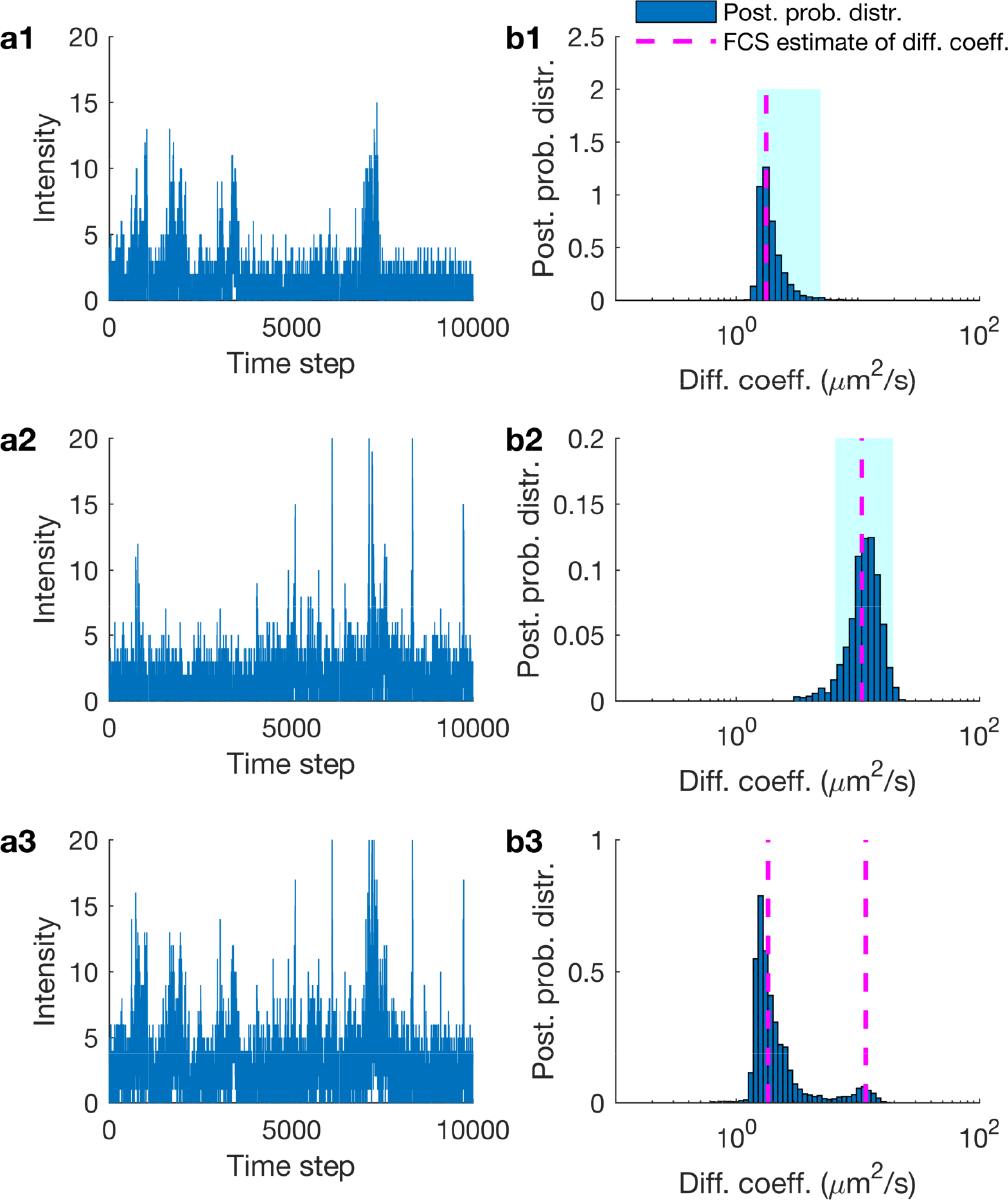
Estimating multiple diffusion coefficients in experimental Cy3 traces. **(a1)–(a2)** Experimental traces of free Cy3 in glycerol/water mixtures with 94% and 75% glycerol, respectively. **(a3)** Trace resulting by mixing the traces in (a1) and (a2). **(b1)–(b2)** Posterior probability distributions resulting from the analysis of the traces in (a1) and (a2). **(b3)** Posterior probability distribution resulting from the analysis of the trace in (a3). For comparison, FCS estimates shown by dashed lines are produced from five different traces, each of 6 min, i.e. ≈100 longer than the segments shown and analyzed in out method. FCS estimates are highlighted by dashed lines. Posteriors are obtained from the analyses of time traces acquired at 100 *µ*s for a total period of 1 s.

## DISCUSSION

Single molecule fluorescence confocal microscopy has the potential to reveal dynamical information at timescales that may be as short as a hundred milliseconds. Here, we have exploited Bayesian non-parametrics to overcome the limitations of specifically fluorescent correlative methods in utilizing short, ≈100 ms, and noisy time traces to deduce molecular properties such as diffusion coefficients. Exploiting novel analysis, to obtain reliable results from such short traces or excessively noisy traces as those obtained under low laser power, is key to minimizing photo-damage inherent to all methods relying on illumination and especially critical to gaining insight on rapid or light-sensitive processes^24,26^. The analysis of similarly short traces may also be required when monitoring non-equilibrium processes that can be resolved only within hundreds of milliseconds. Furthermore, novel analysis with increased sensitivity may reserve longer traces to tease out subtle dynamical features (such as deducing multiple diffusion coefficients at once).

The deep implication of our method is that it places single molecule fluorescence confocal microscopy at a competitive advantage over wide-field techniques used in single particle tracking. Indeed, wide-field techniques provide high, super-resolved, spatial accuracy^15^, but with diminished temporal resolution, since molecule localization requires the collection of sufficient photons obtained only after long frame exposures^15^. Such a requirement is especially problematic for photo-sensitive or rapidly diffusing biomolecules^15^.

By contrast to wide-field microscopy, single molecule fluorescence confocal microscopy yields minimal spatial resolution. However, as our analysis shows, although spatial resolution may be diminished, reduced photo-damage and exceptionally high temporal resolution can be achieved instead.

Since their inception, over half a century ago, correlative methods, such as FCS, have demanded very long traces in order to extract dynamical features from single molecule fluorescence confocal microscopy data^2,11,41–44^. In this study, we have developed a principled framework capable of taking advantage of all spatio-temporal information nested within time traces of photon counts and, together with novel Mathematics, we have reformulated the analysis of single molecule fluorescence confocal microscopy data.

Existing methods, even those that apply Bayesian techniques such as FCS-Bayes^18–22^, still utilize auto-correlation functions. Therefore, they demand equally long time traces as FCS and implicitly assume that the physical system under study is at a stationary or equilibrium phase throughout the entire trace. By contrast, our method only requires short traces and therefore it avoids stationarity or equilibrium requirements on timescales longer than those of the data analyzed. In addition, our method also: (i) provide interpretable estimation of errors (i.e. posterior variance) determined exclusively from the information content of the trace supplied (i.e. length and noise) as opposed to *ad hoc* metric fitting (i.e. chi square); (ii) track instantaneous molecule photon emissions and locations; and (iii) estimate the molecular brightness and background photon emission rates which, if left undetermined, can bias the other estimates.

Since our method is formulated exclusively in the time-domain, it offers a versatile framework for further modifications. For example, it is possible to adapt the present formulation to incorporate scanning-FCS^9,10,45^ which involves moving the confocal volume or incorporate demanding illumination profiles, such as those arising in two photon excitation^43,46^, TIRF microscopy^42^ or even Airy patterns^47^ with or without aberrations^48^ by changing the specified point spread function (see Methods section). Additionally, it is possible to extend our framework to treat multiple diffusion coefficients (see Supplementary materials), confining forces or photon emission kinetics as would be relevant for FCS-FRET^49,50^ and FLIM^51,52^ applications. Also, our method could be extended to handle more complex photophysics^23,53–55^, and, since we explicitly track individual molecules over time, extensions appropriate for fast bimolecular reaction kinetics are also conceivable.

## METHODS

Here we describe the formulation and mathematical foundation of our model. Our overarching goal is to start from an experimental time series of photon counts, 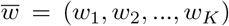 where *w*_*k*_ denotes the photon intensity assessed at time *t*_*k*_ (which includes both background photons as well as photons derived from the labeled molecules of interest), and derive estimates of kinetic quantities such as molecular locations with respect to the center of the confocal volume as well as diffusion coefficients.

To derive estimates for the desired quantities, we need to compute intermediate quantities which include: (i) molecular brightness; (ii) background photon emission rate; and, most importantly, (iii) the unknown population of moving molecules and their relative location with respect to the center of the confocal volume. Below we explain each one of these in detail. Computational details and a working implementation of the entire method are available in the Supplementary materials.

### E. Model description

The starting point of our analysis is the raw data, namely the photon counts. As our current focus is on deducing dynamical information on timescales exceeding ≈1 *µ*s, we ignore triplet state and photon anti-bunching effects which occur on vastly different timescales^16,56,57^.

At the timescale of interest, individual photon detections happen stochastically and independently from each other. Accordingly, the total number of photon counts *w*_*k*_ between successive assessments follows Poisson^15,27^ (shot noise) statistics

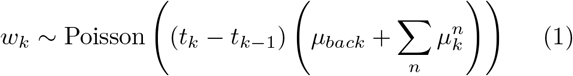

where *µ*_*back*_ is a background photon emission rate and 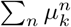 gathers the photon emission rates 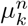 from individual fluorescent molecules that we index with *n* = 1, 2,…. The the total number of molecules involved in the summation above is to be determined. This is the key reason we invoke Bayesian non-parametrics in the model inference section (see below). Since we only collect a small fraction of the total photons emitted by the fluorescent molecules, in our framework 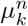 coincides with the emission rate of detected photons, as opposed to the true photon emission rate which might be larger.

Each rate 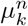 depends on the position 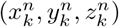 of the corresponding molecule relative to the center of the confocal volume as well as other features such as laser intensity, laser wavelength, quantum yield and camera pinhole size^58^. Similar to other studies^41,59,60^, we combine all these effects into a characteristic point spread function (PSF) that combines excitation and emission PSFs

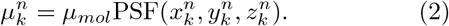

The parameter *µ*_*mol*_ represents the molecular brightness and as we discuss in the supplementary materials it is related to the maximum photon emission rate of a single molecule that is located at the center of the confocal volume. Specific choices of PSF models, such as Gaussian or Gaussian-Lorentzian, are also detailed in the Supplementary materials.

Finally, we associate individual molecular locations across time by adopting a motion model. Here we assume that molecules are purely diffusive and arrive at

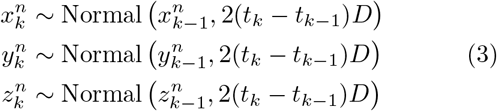

where *D* denotes the diffusion coefficient, which we assume is the same for all molecules. As we explain in the Supplementary materials, these probabilities result directly from the diffusion equation. Additionally, in the Supplementary materials, we illustrate how this motion model can be generalized to capture more than one diffusion coefficients.

A graphical summary of the entire formulation is shown on Fig. 8.

**FIG. 8.**
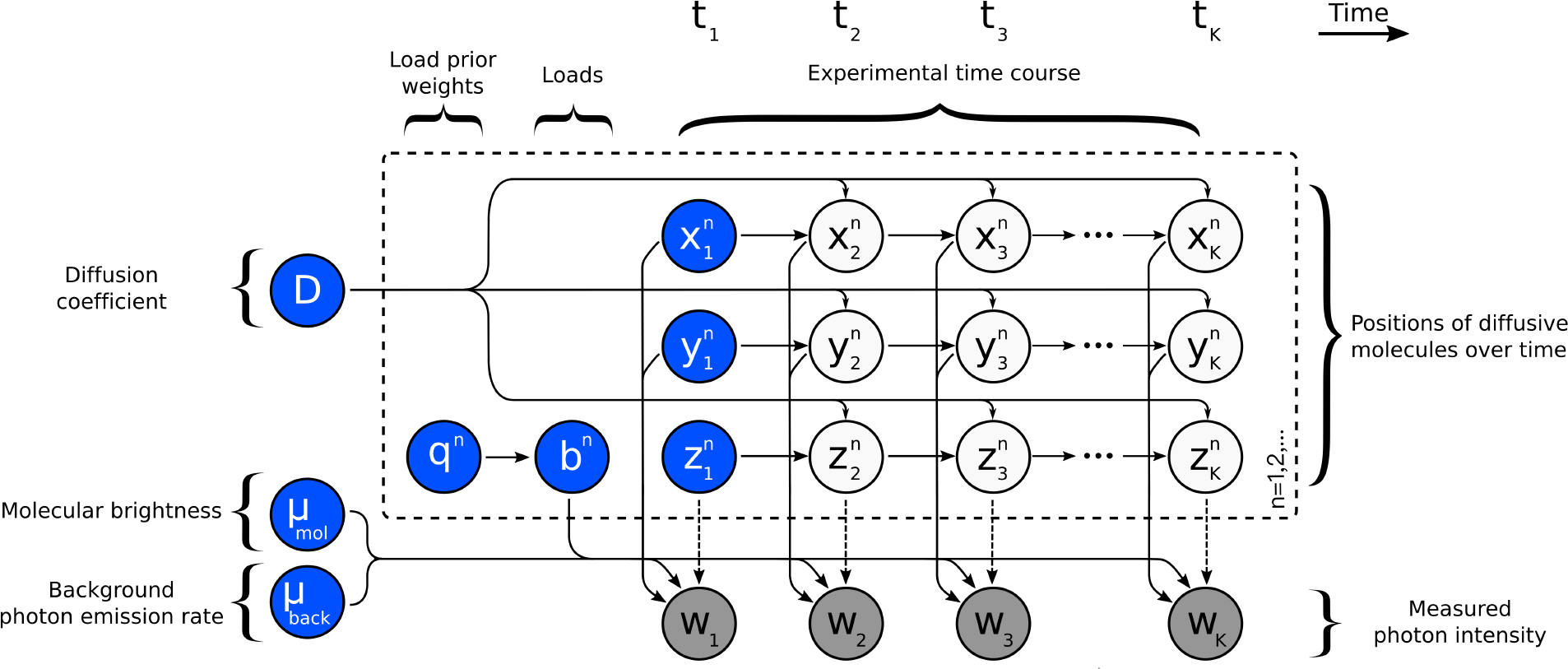
Graphical representation of the formulation used in the analysis of fluorescence time traces. A population of model molecules, labeled by *n* = 1, 2, …, evolves over the course of the experiment which is marked by *k* = 1, 2, …, K. Here, 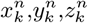 denote the location in Cartesian space of molecule *n* at time *t*_*k*_; *µ*_*mol*_ denotes the brightness of an individual molecule; and *µ*_*back*_ denotes the background photon emission rate. During the experiment, only a single observation *w*_*k*_, combining photon emissions between *t*_*k*−1_ and *t*_*k*_ from every molecule and background is recorded at every time step. The diffusion coefficient *D* determines the evolution of the molecular locations which, in turn, influence the photon emission rates and ultimately the recorded photon intensity *w*_*k*_. Auxiliary variables *b*^*n*^, or “loads”, and corresponding prior weights *q*^*n*^, are introduced in order to estimate the unknown population size. The dashed arrows apply for the 3D-Gaussian and 2D-Gaussian-Lorentzian PSFs; while in the case of the 2D-Gaussian-Cylindrical there is no dependency of the measurements *w*_*k*_ on the 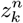 coordinates of the molecules (see the Supplementary materials for the definitions of these PSFs).

### F. Model inference

The quantities which we want to estimate, for example the diffusion coefficient *D*, molecular locations through time 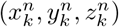, molecular brightness *µ*_*mol*_ and background photon emission rate *µ*_*back*_ or the molecular population are introduced as model variables in the preceding formulation. To estimate values for these variables, we follow the Bayesian paradigm^15,28,38,60^.

Variables such as *D*, *µ*_*mol*_ and *µ*_*back*_ are parameters of the model and, as such, require priors. Choices for these priors are straightforward and, for interpretational and computational convenience, we adopt the distributions described in the Supplementary materials.

Additionally, we must place priors on the initial molecular locations, 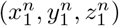, i.e. the locations of the molecules at the onset of the measurement period. Specifying a prior on initial molecular locations also entails specifying a prior on the molecular population.

In particular, to allow the dimensionality or, alternatively, the complexity of our model to fluctuate based on the number of molecules that contribute to the fluorescent trace, we abandon traditional Bayesian parametric priors and turn to the non-parametric formulation described below.

Before we proceed any further, we recast equation (2) as

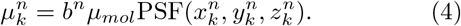

The newly introduced variables *b*^*n*^, one for each model molecule, may take only values 1 or 0. In particular, the possibility that *b*^*n*^ = 0, coinciding with the case where molecules do not contribute to the observation, allows us to introduce an arbitrarily large number of molecules, technically an infinite number. With the introduction of *b*^*n*^, we can estimate the number of molecules that contribute photons (termed “active” to distinguish them from those that do not contribute termed “inactive”) simultaneously with the rest of the parameters simply by treating each *b*^*n*^ as a separate parameter and estimating its value (of 1 for active molecules and 0 for inactive ones).

To estimate *b*^*n*^, we place a prior *b*^*n*^ ~ Bernoulli(*q*^*n*^) and subsequently a hyperprior on *q*^*n*^ in order to learn precisely how many model molecules are active. For the latter, we choose *q*^*n*^ ~ Beta(*A*_*q*_, *B*_*q*_) with hyperparameters *A*_*q*_ and *B*_*q*_. Both steps can be combined by invoking the newly developed Beta-Bernoulli process^36,61^ which is described in more detail in the Supplementary materials.

Once the choices for the priors above are made, we form a joint posterior probability 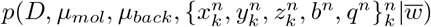 encompassing all unknown variables which we may wish to determine.

The nonlinearities in the PSF, with respect to variables 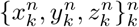, and the non-parametric prior on {*b*^*n*^, *q*^*n*^}^*n*^ exclude analytic forms for our posterior. For this reason, we develop a computational scheme exploiting Markov chain Monte Carlo^38,62^ that can be used to generate pseudo-random samples from this posterior.

The main bottleneck of a naive implementation of our method, as compared to correlative methods, is its higher computational cost. As we explain in the Supplementary materials, to have computations run on an average desktop computer, we adopt mathematical approximations (e.g. photon binning, Anscombe transform^63^ and filter updates^64,65^) that are tested on the synthetic data presented.

A working implementation (source code and GUI) is given in the Supporting Materials.

### G. Data acquisition

#### 1. Synthetic data

We obtain the synthetic data presented in the Results section by standard pseudo-random computer simulations^66–68^ that mimic the common single molecule fluorescence confocal setup. We provide details and complete parameter choices in the Supplementary materials.

#### 2. Experimental data

For the experimental data acquired with elongated confocal volumes, a stock solution of Cy3B (monoreactive NHS ester, GE Healthcare) was prepared by dissolving a small amount of solid in 1 mL of doubly-distilled water, and its concentration was determined from the absorbance of the solution using the extinction coefficient provided by the vendors. A 10 nM solution was then prepared by appropriate dilution of the stock and measured on a silicone perfusion chamber mounted on a glass coverslip. Fluorescent beads were purchased from ThermoFisher (Catalog number: F8792. Lot number: 1604237). The average diameter was 0.046 *µ*m as indicated in the certificate of analysis provided by the vendors. Suspensions for FCS measurements were prepared by adding 3 *µ*L of stock solution (9.4 × 1014 particles/mL) to 1 mL of water and sonicating the mixture for 20 minutes. Measurements were carried out using a home-built instrument. A 532 nm continuous-wave laser (Compass 215M-10, Coherent, Santa Clara, CA) was attenuated to 100 *µ*W and focused onto an PlanApo 100×, 1.4 NA, oil-immersion, objective (Olympus, Center Valley, PA). Emitted fluorescence was collected using the same objective and then passed through a 50 *µ*m pinhole to reject the out-of-focus light. The signal was detected using a silicon avalanche photodiode (SPCM-AQR-14; Perkin-Elmer, Fremont, CA). A band-pass filter (Omega 3RD560-620) in front of the detector was employed to further reduce the background signal and an ALV correlator card (ALV 5000/EPP, ALV-GmbH, Langen, Germany) was used to correlate the detected fluorescence signal. Data for our analysis were acquired with 100 *µ*s resolution using a PCI-6602 acquisition card (National Instruments, Austin, TX).

For the experimental data acquired with elliptical confocal volumes, Cy3 dye and Cy3-labeled streptavidin solutions were prepared by suspending Cy3 or streptavidin in glycerol/buffer (pH 7.5, 10 mM Tris-HCl,100 mM NaCl and 10 mM KCl, 2.5 mM CaCl_2_) at different v/v, to a final concentration of either 100 pM or 1 nM. The solutions were added onto a glass-bottomed fluid-cell, mounted on a custom designed single molecule fluorescence confocal microscope^69,70^ and a 532 nm laser beam was focused to a diffraction-limited spot on the glass coverslip of the fluid-cell using a 60×, 1.42 NA, oil-immersion objective (Olympus). the laser power was measured before the objective and the beam is reflected by a dichroic and focused by the objective on to the sample. The dichroic reflected 95% of the intensity on to the objective. Emitted fluorescence was collected by the same objective and focused onto the detection face of a Single Photon Avalanche Diode (SPAD, Micro Photon Devices) that has a maximum count rate of 11.8 Mc/s. A bandpass filter was placed in front of the detector to transmit only the fluorescence from Cy3 and to block the back-scattered excitation light. TTL pulses, triggered by the arrival of individual photons on the SPAD, were timestamped and recorded at 80 MHz by a field programmable gated array (FPGA, NI Instruments) using custom LabVIEW software and initially binned at 100 *µs*^70^.

## ACKNOWLEDGMENTS

SP acknowledges support from NSF CAREER grant MCB-1719537. SS acknowledge support from the National Institute of Health grant R01GM121885. ML thanks Anirban Purohit for assistance with the experiments.

## AUTHOR CONTRIBUTIONS

SJ developed analysis software and analyzed data; SJ, IS developed computational tools; OS, SS, ML contributed experimental data; SJ, IS, SP conceived research; SP oversaw all aspects of the projects. SJ and IS contributed equally to this work.

## CODE AVAILABILITY

Source code and GUI versions of the methods developed herewith are available through the Supporting Materials.

## Supplementary Material

Here we provide supplementary materials and technical details that complement the main text. These include: (i) Additional analysis results that demonstrate the estimation of molecular brightness and background photon emission rates, joint posterior probability distributions, molecule locations, and additional results for multiple diffusive species. These results are repeated for simulated and experimental data. (ii) Additional details of the methods used including descriptions of the motion model, the Stokes-Einstein model, point spread functions (PSFs), and time trace preparation. (iii) A complete description of the inference framework developed that includes choices for the prior probability distributions and a computational implementation. (iv) A description of the modifications necessary for the model with multiple diffusive species. (v) Summary of notation and other conventions used throughout this study as well as detailed parameter choices for the simulations and analyses.

## S1. Additional results

### S1.1. Analysis of additional simulated data

**FIG. S1.**
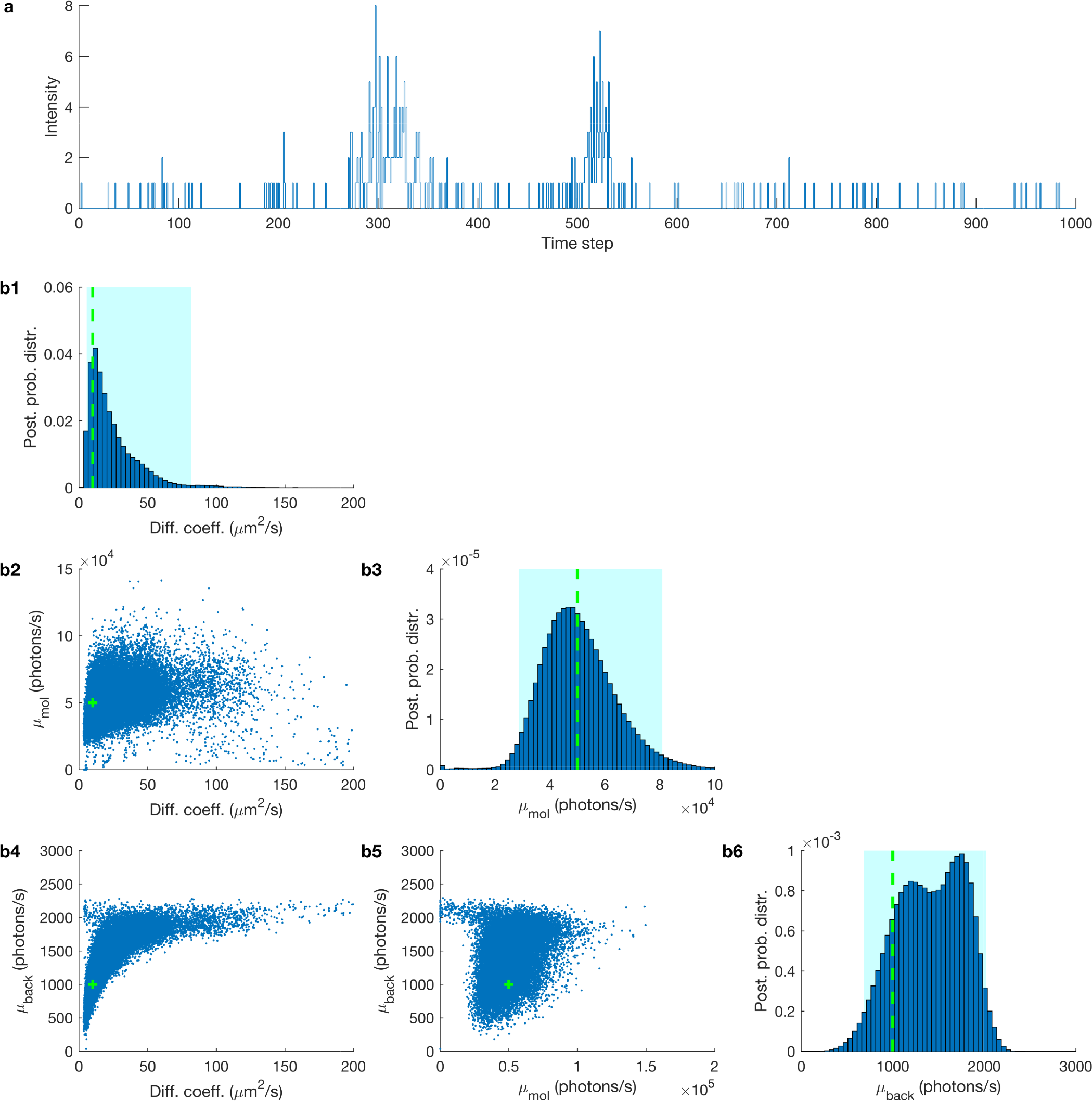
Estimated joint posterior probability distribution. **(a)** Synthetic fluorescent intensity trace used in Fig. 1a1 with a length of 1000 data points and time step 100 *µ*s. The true values of the diffusion coefficient, molecular brightness and background emission rates are, 10 *µm*^2^/*s*, 5 × 10^4^ photons/s and 10^3^ photons/s (shown by green dashed lines). **(b1)** The posterior of the diffusion coefficient. **(b2)** The joint probability distribution of the diffusion coefficient and molecular brightness. **(b3)** The posterior probability distribution of the molecular brightness. **(b4)** The joint probability distribution of the diffusion coefficient and molecular brightness. **(b5)** The joint probability distribution of the molecular brightness and background photon emission rates. **(b6)** The posterior probability distribution of the background photon emission rate. The 95% confidence intervals are shown with pink highlighted regions.

**FIG. S2.**
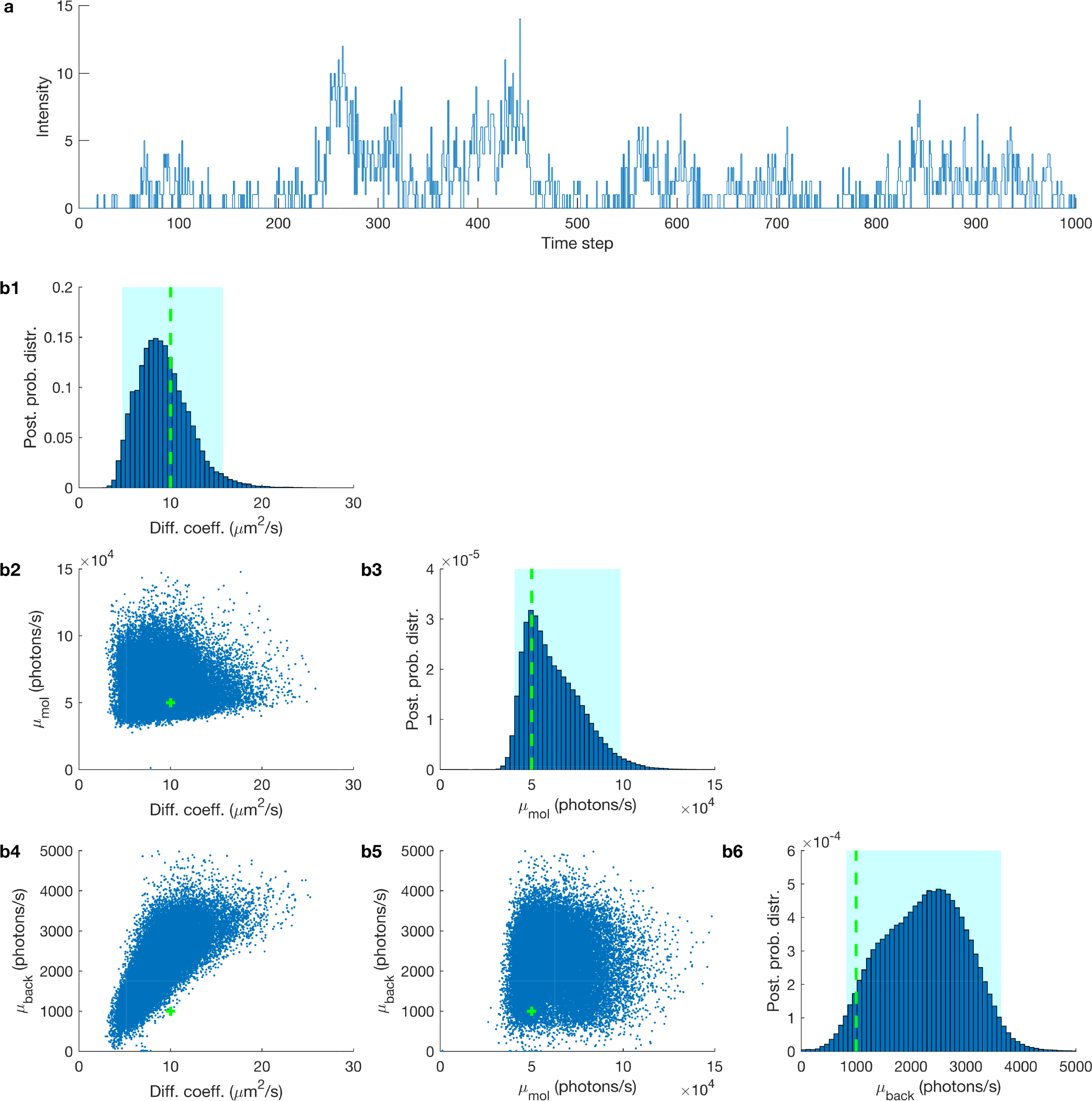
Estimated joint posterior probability distribution. **(a)** Synthetic fluorescent intensity trace used in Fig. 1a2 with a length of 1000 data points and time step 100 *µs*. The true values of the diffusion coefficient, molecular brightness and background emission rates are, 10 *µm*^2^/*s*, 5 × 10^4^ photons/s and 10^3^ photons/s (shown by green dashed lines). **(b1)** The posterior of the diffusion coefficient. **(b2)** The joint probability distribution of diffusion coefficient and molecular brightness. **(b3)** The posterior probability distribution of the molecular brightness. **(b4)** The joint probability distribution of the diffusion coefficient and molecular brightness. **(b5)** The joint probability distribution of the molecular brightness and background photon emission rates. **(b6)** The posterior probability distribution of the background photon emission rate. The 95% confidence intervals of the posterior over the number of molecules is highlighted in cyan.

**FIG. S3.**
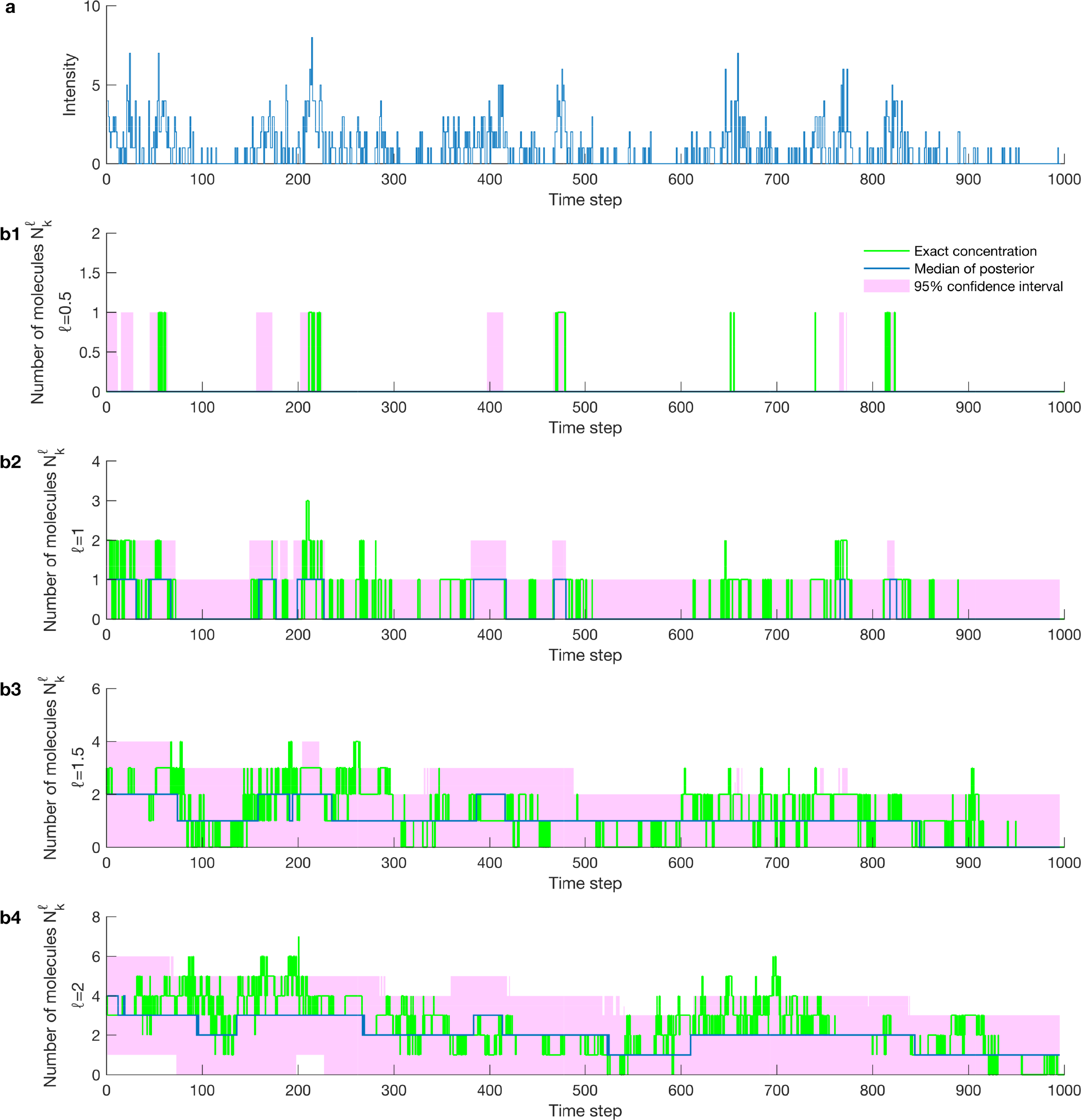
Estimated number of molecules/concentrations. **(a)** Synthetic fluorescent intensity trace produced with a molecular brightness of 5 × 10^4^ photons/s and a background photon emission rate of 10^3^ photons/s, diffusion coefficient of 10 *µm*^2^/s and 50 molecules. **(b1)–(b4)** Number of molecules estimated from the trace in (a) corresponding to normalized distances from the confocal center of *ℓ* = 0.5, 1, 1.5, 2. The exact number of molecules is shown by the green lines, the median of the posterior over the number of the molecules is shown by the blue lines, and the 95% confidence intervals of the posteriors over the number of the molecules are highlighted in pink. For details of the definition of number of molecules 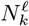 and the normalized distance ℓ, see Eq. (S23), below.

**FIG. S4.**
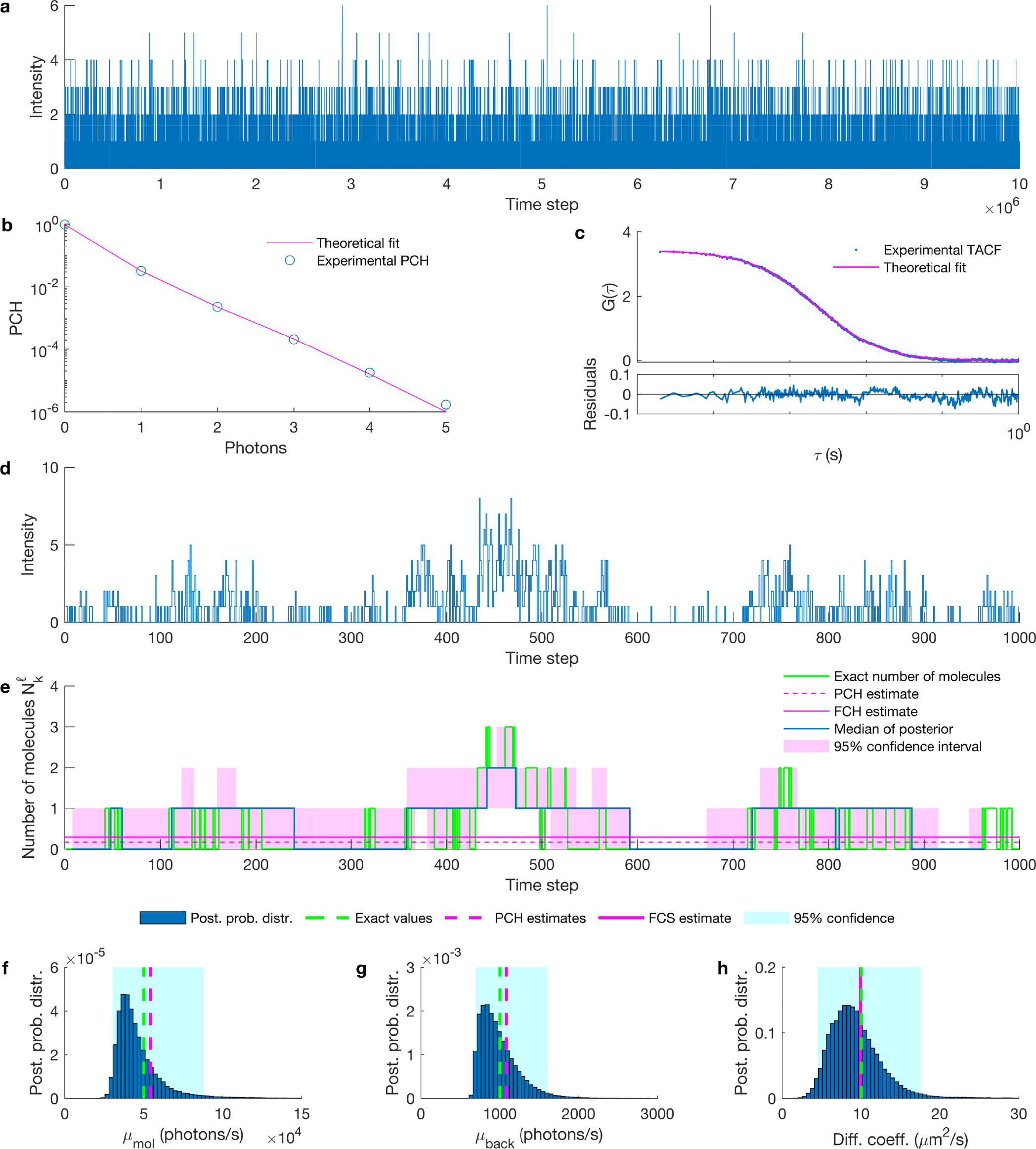
Comparison of estimated photon emission rates and concentration with FCS and PCH. **(a)** Targeted synthetic fluorescent intensity trace. The time step is 10 *µ*s and the total duration of the trace is 100 s. **(b)** PCH curve and the theoretical fit. **(c)** FCS curve and best theoretical fit. **(d)** The portion of the trace analyzed by our method rebinned at 100 *µs*. **(e)** The number of molecules in the effective volume with ℓ =1, arising from the trace in (d). Exact value of the number of molecules is shown by the green line and the PCH and FCS estimates are shown by the dashed and solid pink lines. **(g)** On the posterior probability distribution of the molecular brightness we superpose the PCH estimate of the molecular brightness (pink dashed line) and the true value (green dashed line). **(h)** On the posterior probability distribution of the background photon emission rate we superpose the PCH estimate of the background photon emission rate (pink dashed line) and the true value (green dashed line). **(f)** The posterior probability distribution of the diffusion coefficient obtained by analyzing the trace in (d). The FCS estimate of the diffusion coefficient obtained by analyzing the total trace, shown in (a), illustrated by a pink dashed line with the exact value (green dashed line). The targeted synthetic trace is generated by freely diffusive molecules with diffusion coefficient, molecular brightness and background photon emission rates of of 10 *µm*^2^/s, 5 × 10^4^ photons/s and 10^3^ photons/s, respectively.

**FIG. S5.**
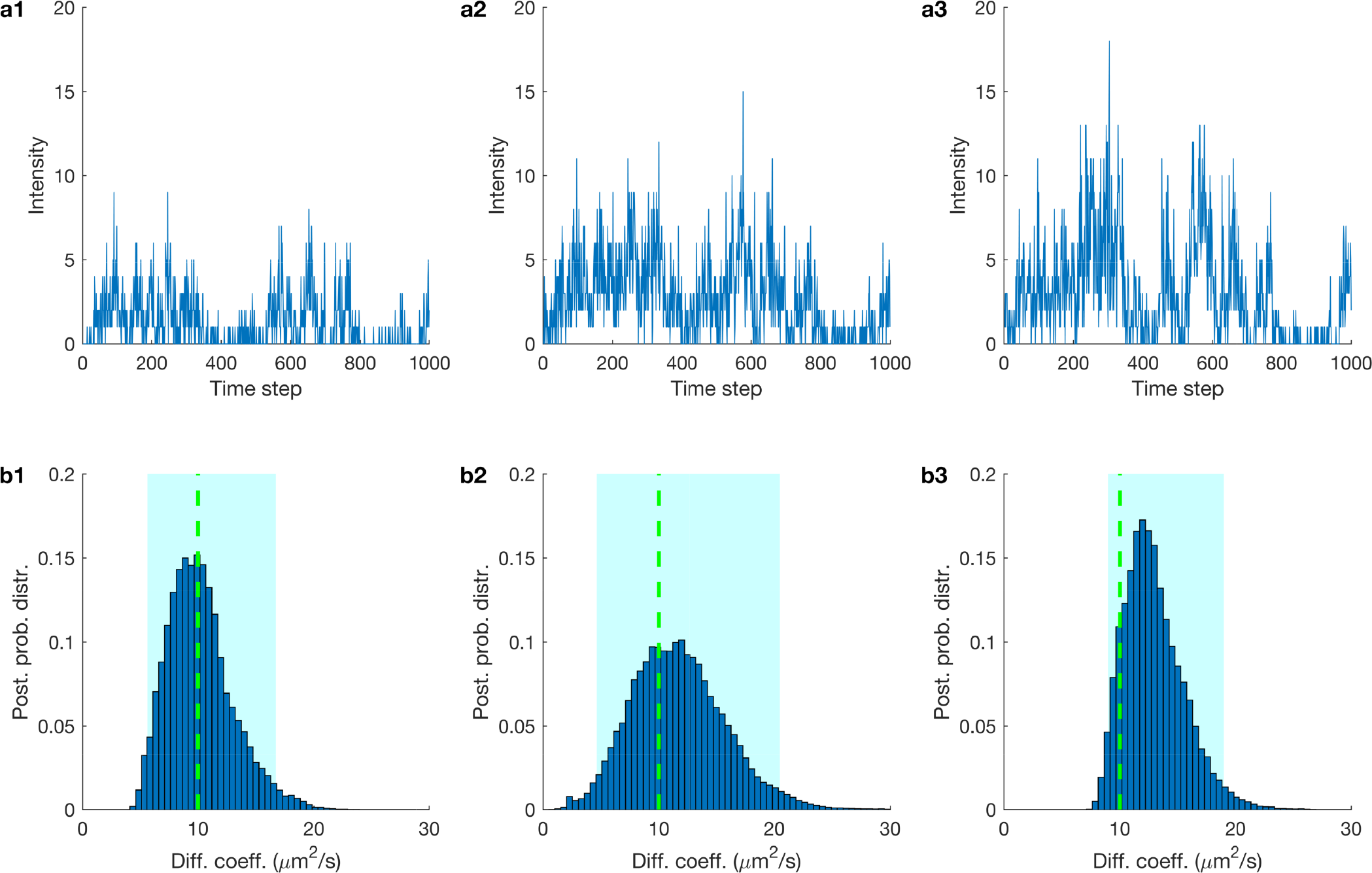
Comparison of the posterior on the diffusion coefficient obtained from synthetic fluorescent intensity traces under different PSF models. **(a1)** Synthetic fluorescent intensity trace produced with a 3DG PSF, Eq. (S16). **(a2)** Synthetic fluorescent intensity trace produced with a 2DGL PSF, Eq. (S18). **(a3)** Synthetic fluorescent intensity trace produced with a 2DGC PSF, Eq. (S17). **(b1)** Posterior of the diffusion coefficient using a 3DG PSF model on the trace in (a1). **(b2)** Posterior of the diffusion coefficient using a 2DGL PSF model on the trace in (a2). **(b3)** Posterior of the diffusion coefficient using a 2DGC PSF model on the trace in (a3). To facilitate the comparison both traces analyzed are generated using the same underlying molecule trajectories with molecular brightness and background photon emission rates of 5 × 10^4^ photons/s and 10^3^ photons/s, diffusion coefficient of 1 *µm*^2^/s (shown by green dashed lines).

### S1.2. Analysis of additional experimental data

**FIG. S6.**
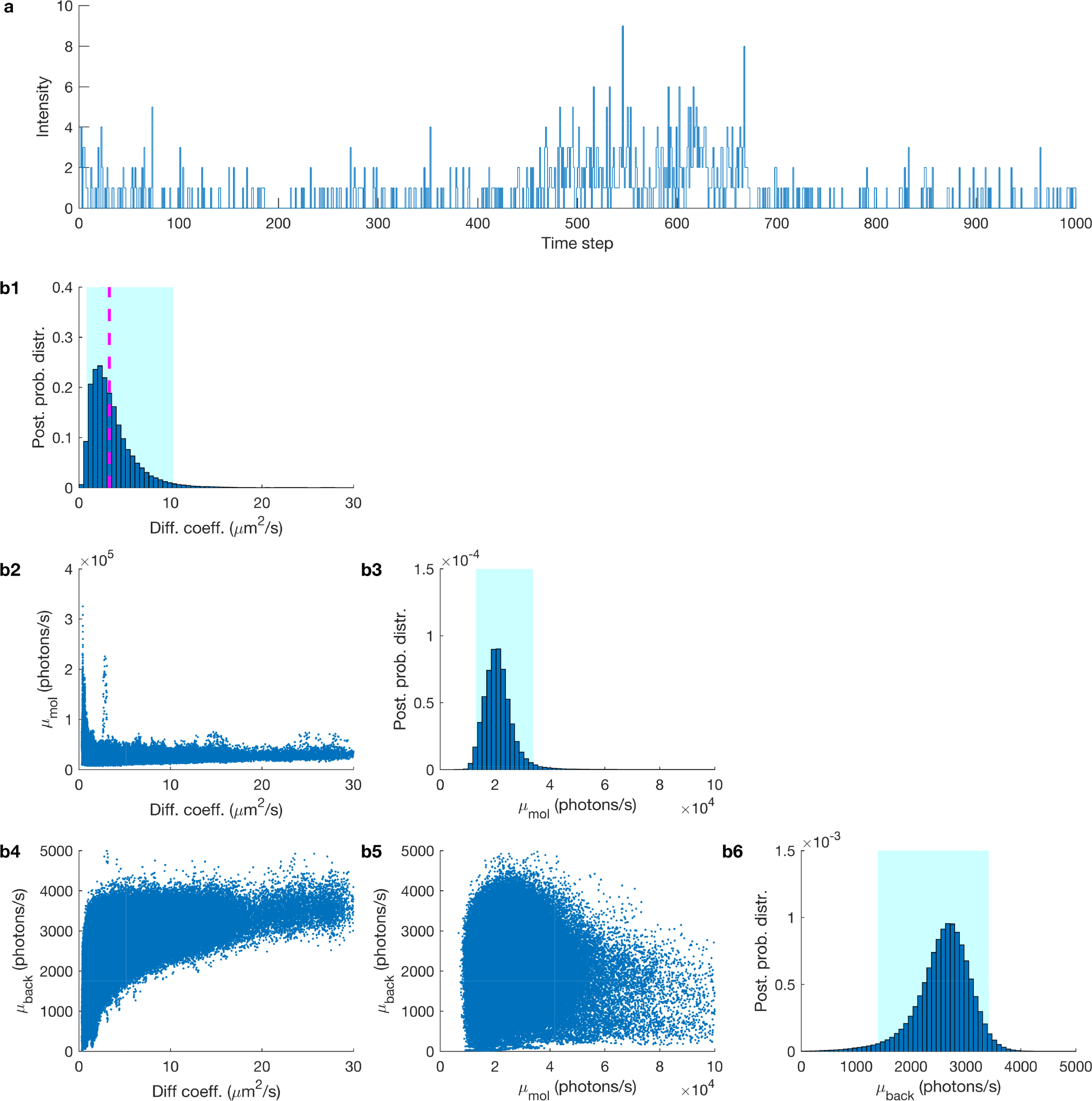
Estimated joint posterior probability distribution. **(a)** Experimental fluorescent intensity trace used in Fig. 5b3 with a length of 1000 data points and time step 100 *µs*. **(b1)** The posterior of the diffusion coefficient. The FCS estimate is shown by a magenta dashed line.**(b2)** The joint probability distribution of the diffusion coefficient and the molecular brightness. **(b3)** The posterior probability distribution of the molecular brightness. **(b4)** The joint probability distribution of the diffusion coefficient and molecular brightness. **(b5)** The joint probability distribution of the molecular brightness and background photon emission rates. **(b5)** The posterior probability distribution of the background photon emission rate and the 95% confidence intervals of the posteriors are highlighted in cyan. The experimental fluorescent intensity trace was produced with a concentration of 100 pM of Cy3 in a 94% glycerol/water mixture.

**FIG. S7.**
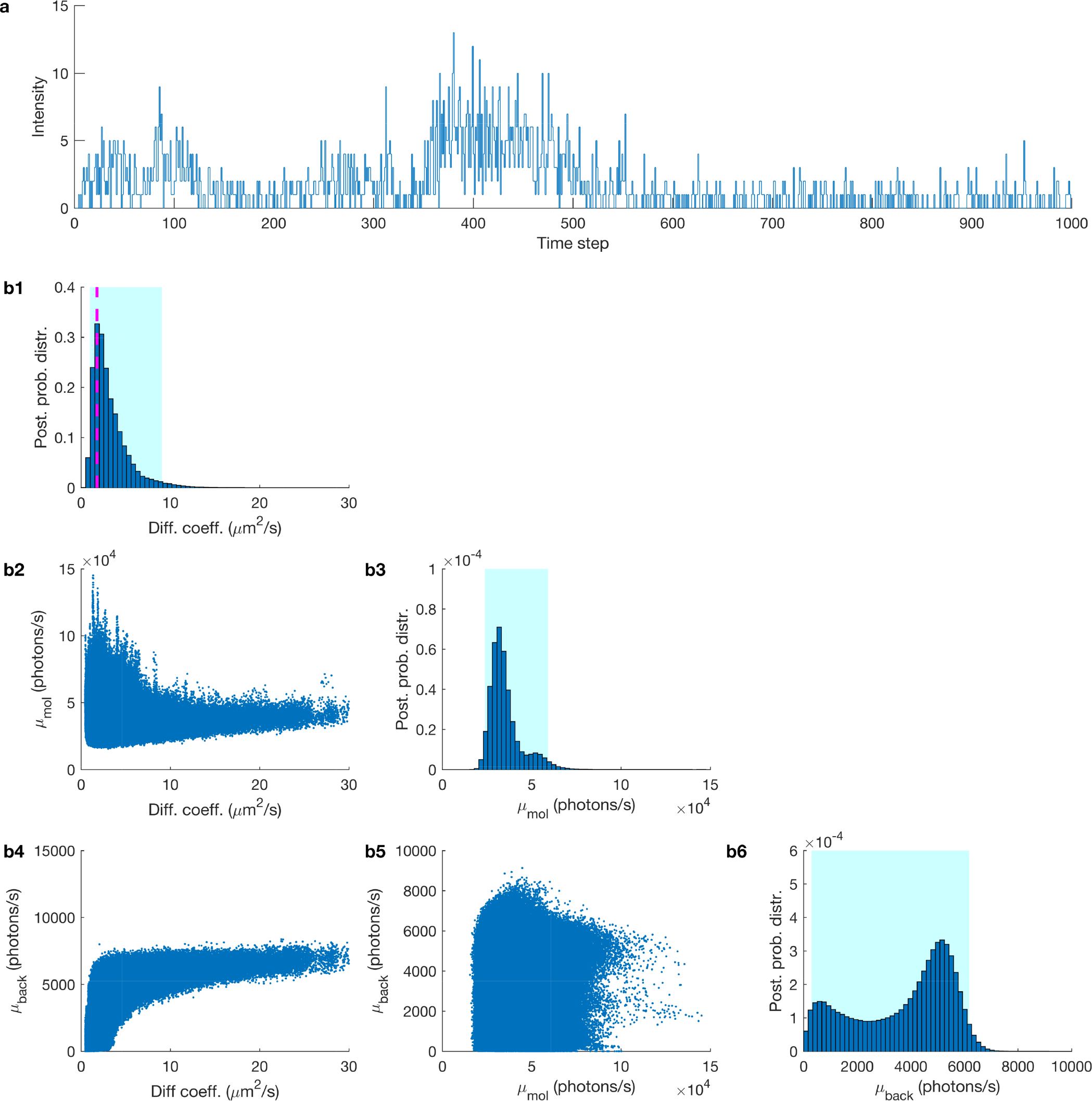
Estimated joint posterior probability distribution. **(a)** Experimental fluorescent intensity trace used in Fig. 5b4 with a length of 1000 data points and time step 100 *µs*. **(b1)** The posterior of the diffusion coefficient. The FCS estimate is shown by a magenta dashed line. **(b2)** The joint probability distribution of the diffusion coefficient and the molecular brightness. **(b3)** The posterior probability distribution of the molecular brightness. **(b4)** The joint probability distribution of the diffusion coefficient and the molecular brightness. **(b5)** The joint probability distribution of the molecular brightness and background photon emission rates. **(b5)** The posterior probability distribution of the background photon emission rate and the 95% confidence intervals of the posteriors are highlighted in cyan. The experimental fluorescent intensity trace produced is with a concentration of 1 nM of Cy3 in a 94% glycerol/water mixture.

**FIG. S8.**
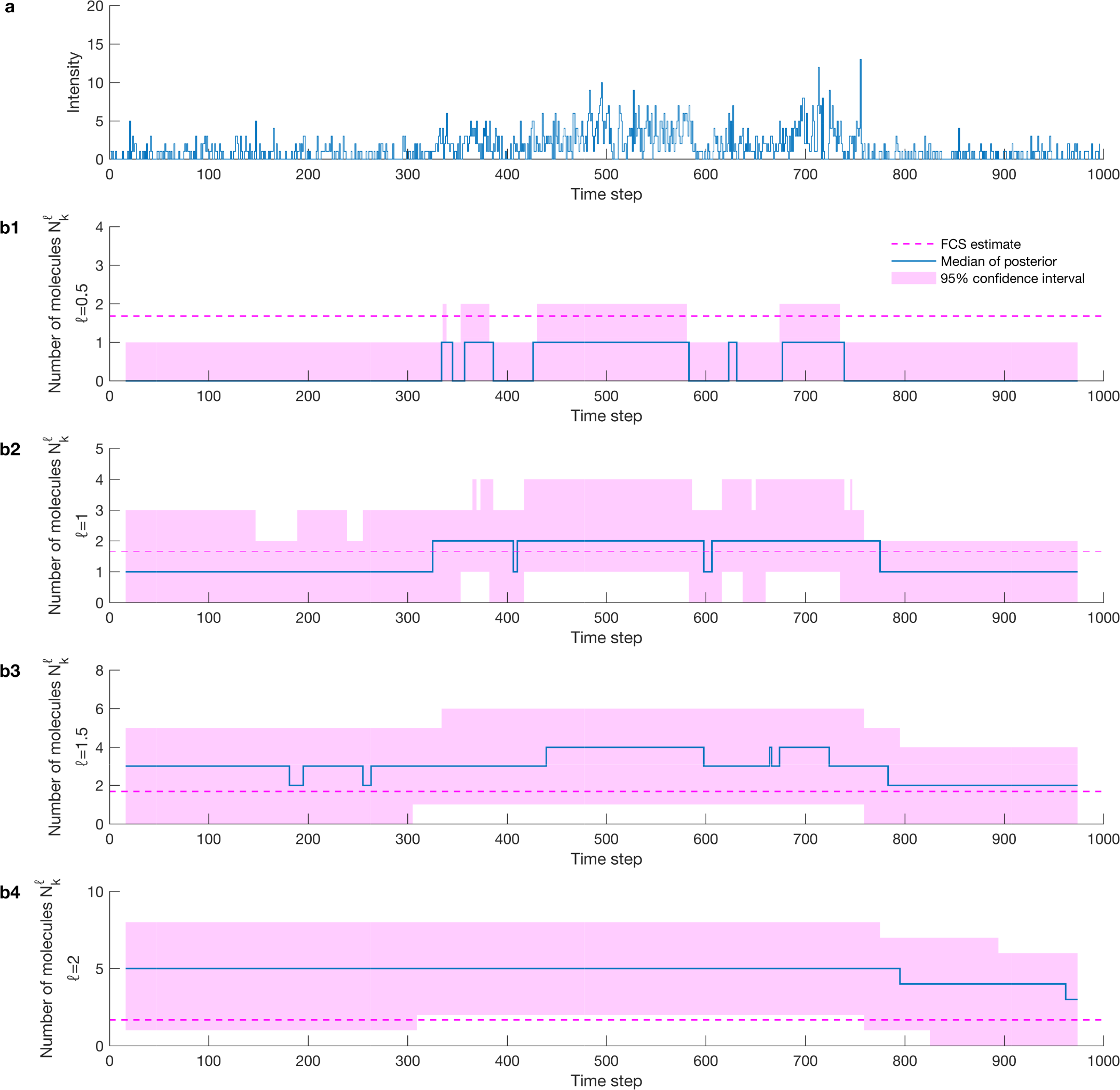
Estimated number of molecules/concentrations. **(a)** Experimental fluorescent intensity trace produced with a concentration of 1 nM of Cy3 in a 94% glycerol/water mixture. **(b1)–(b4)** Number of molecules estimated from the trace in (a) with ℓ = 0.5, 1, 1.5, 2, respectively. The FCS estimate of the average number of molecules in the effective volume (~1.68 *molecules*) by analyzing a 3 minutes long time trace, is shown by the magenta dashed lines and the median of the posterior over the number of the molecules is shown by a blue line. The 95% confidence interval of the posterior over the number of the molecules is highlighted in pink. For details of the definition of number of the molecules 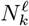 and the normalized distance ℓ, see Eq. (S23), below.

**FIG. S9.**
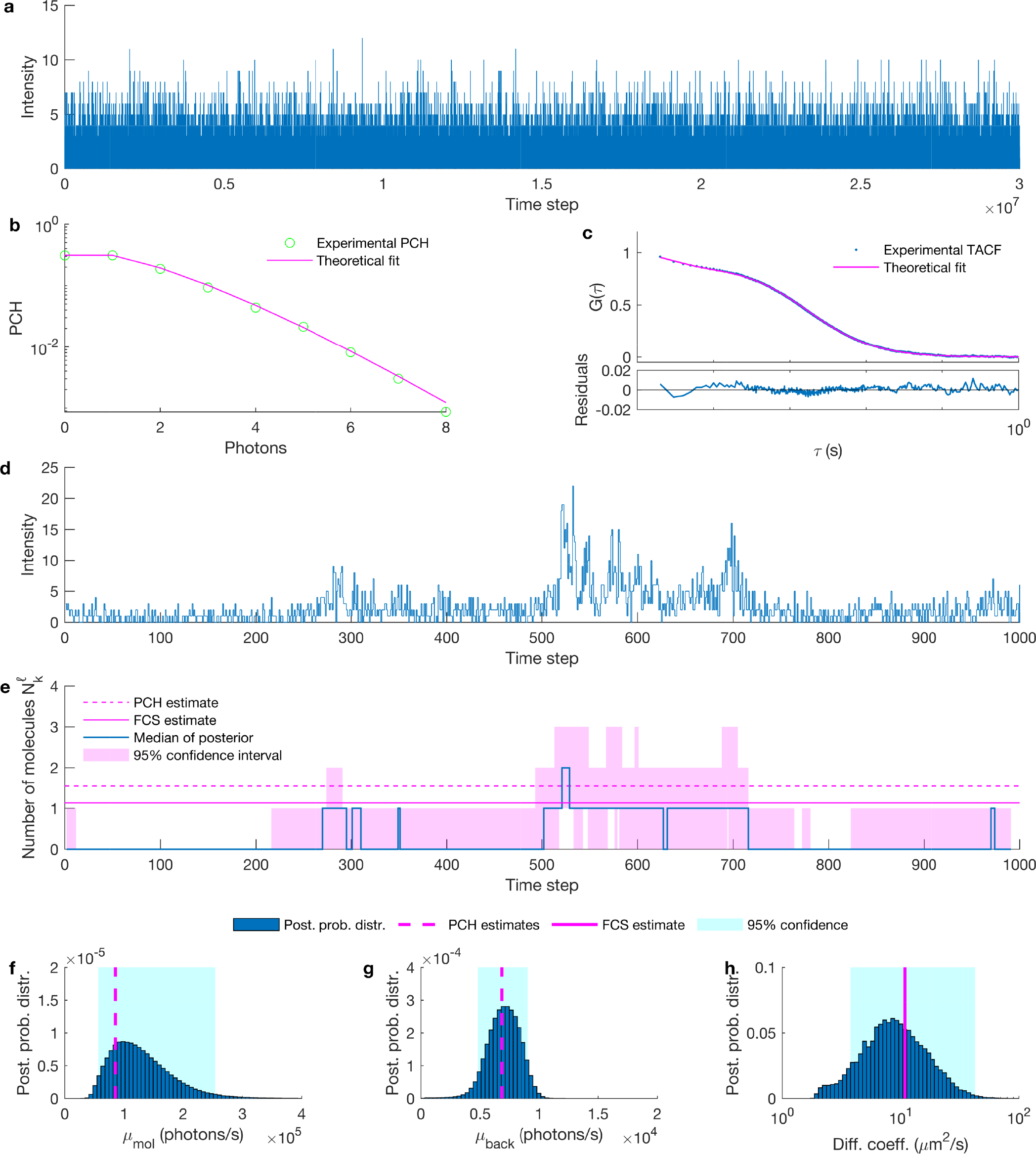
Comparison of estimated photon emission rates and concentration with FCS and PCH. **(a)** Targeted experimental fluorescent intensity trace. The time step is 10 *µ*s with a total time of 5 min. **(b)** PCH curve and the theoretical fit. **(c)** FCS curve and the best theoretical fit. **(d)** The portion of the trace analyzed by our method rebinned at 100 *µs*. **(e)** The concentration of Cy3 in the effective volume with ℓ =1, arising from the trace in (d). The experimental concentration is shown by the green line and the PCH estimated is shown by the pink line. **(g)** The posterior probability distribution of the molecular brightness with the PCH estimated of the molecular brightness shown by a solid green line. **(h)** The posterior probability distribution of the background photon emission rate with the PCH estimate of the background photon emission rate shown by a solid green line. **(f)** The posterior probability distribution of the diffusion coefficient obtained by analyzing the trace in (d). The FCS estimate of the diffusion coefficient obtained by analyzing the total time trace, shown in (a), is denoted by a pink solid line. The targeted experimental trace is generated by free diffusive Cy3 in a mixture of water and glycerol with 75% glycerol, a laser power of 100 *µ*W and a concentration of Cy3 at 1 nM, excitation wavelength, NA and refractive index used are 532 nm, 1.42 and 1.4, respectively.

**FIG. S10.**
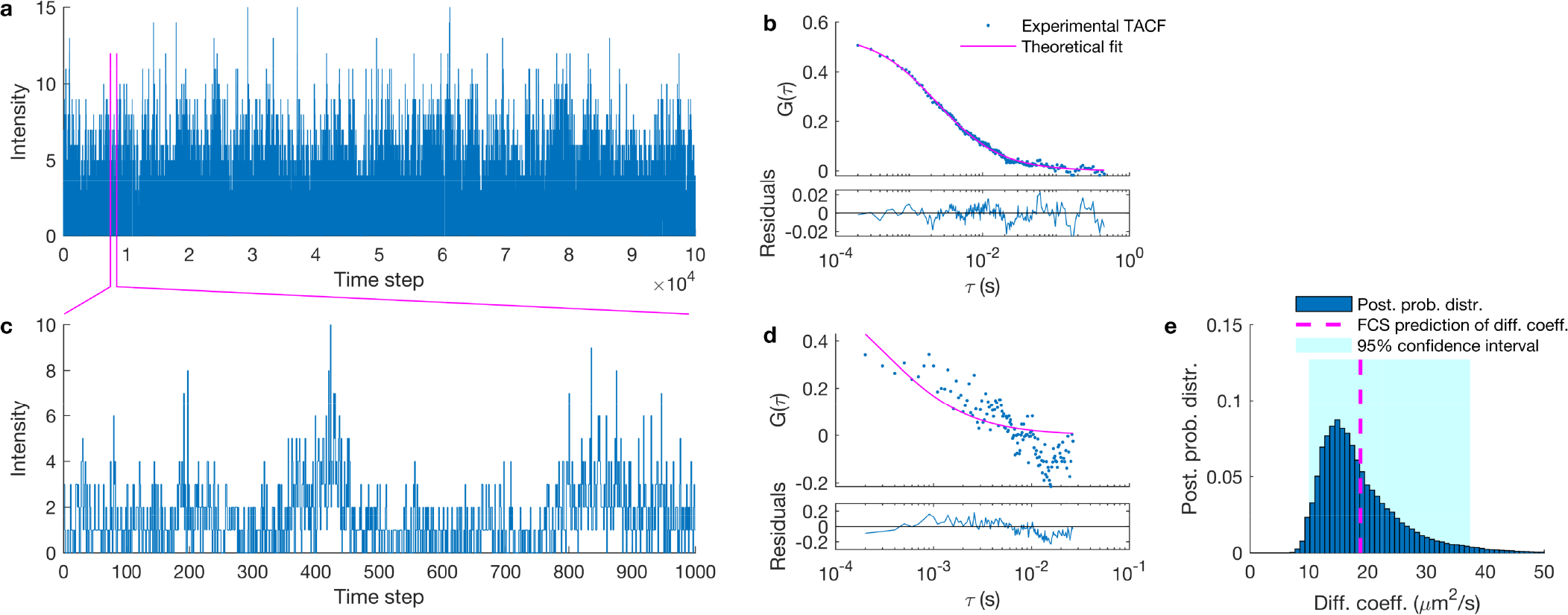
Testing diffusion coefficient estimates in experimental traces of free Cy3B dyes using an elongated confocal volume. **(a)** Experimental fluorescent intensity trace used in FCS. The time trace is generated by 2.5 nM Cy3B dyes in glycerol/water mixture with 70% glycerol and laser power of 100 *µW*. **(b)** Auto-correlation curve of the trace in (a) and best theoretical fit. **(c)** Portion of the trace in (a) to be used as the input to FCS and our method. **(d)** Auto-correlation curve of trace in (c). **(e)** Posterior probability distribution over diffusion coefficient estimated from the trace in (c). Traces shown in (a) and (c) are acquired at 100 *µ*s for a total of 10 second and 0.1 second respectively. The laser power use to generate the signal (a) is 100 *µW* measured before the beam enters the objective. The estimation of the diffusion coefficient as the results of autocorrelation fitting in (a) matched with Stokes-Einstein prediction, equal to 18.79 *µm*^2^/*s* and in (d) is 145.75 *µm*^2^/*s*.

### S1.3. Analysis of additional data for multiple diffusive species

**FIG. S11.**
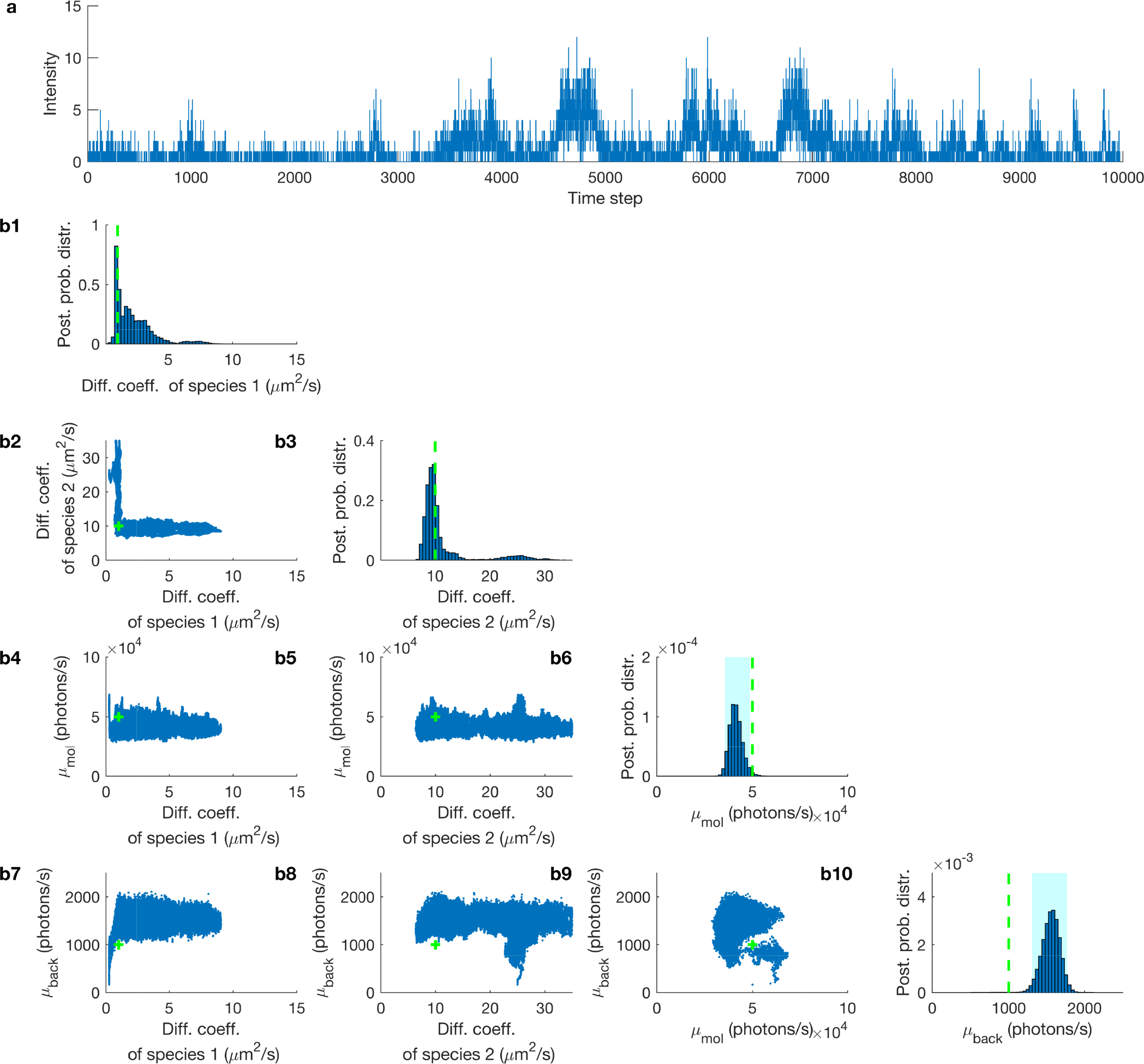
Estimated joint posterior probability distribution of multiple diffusive species. **(a)** A mixed fluorescent intensity trace was obtained by combining the traces from two different synthetic signals with molecular brightness and background emission rates of 5 × 10^4^ and 10^3^ photons/s, respectively, and diffusion coefficients of 1 and 10 *µm*^2^/*s*. **(b1)** The posterior probability distribution of the diffusion coefficient of diffusive species 1. **(b2)** The joint probability distribution of the diffusion coefficient for diffusive species 1 and diffusive species 2. **(b3)** The posterior probability distribution of the diffusion coefficient of diffusive species 2. **(b4)** The joint probability distribution of diffusion coefficient of diffusive species 1 along with the molecular brightness. **(b5)** The joint probability distribution of diffusion coefficient of diffusive species 2 along with the molecular brightness. **(b6)** The posterior probability distribution of the molecular brightness. **(b7)** The joint probability distribution of diffusion coefficient for diffusive species 1 along with the background photon emission rate. **(b8)** The joint probability distribution of the diffusion coefficient of diffusive species 2 and the background photon emission rate. **(b9)** The joint probability distribution of the molecular brightness and background photon emission rate. **(b10)** The posterior probability distribution of the background photon emission rate. The trace is binned at 100 *µ*s with a total trace duration of 1 s. The exact values of the parameters are shown by green dashed lines and the 95% confidence intervals of the posteriors are highlighted in cyan.

**FIG. S12.**
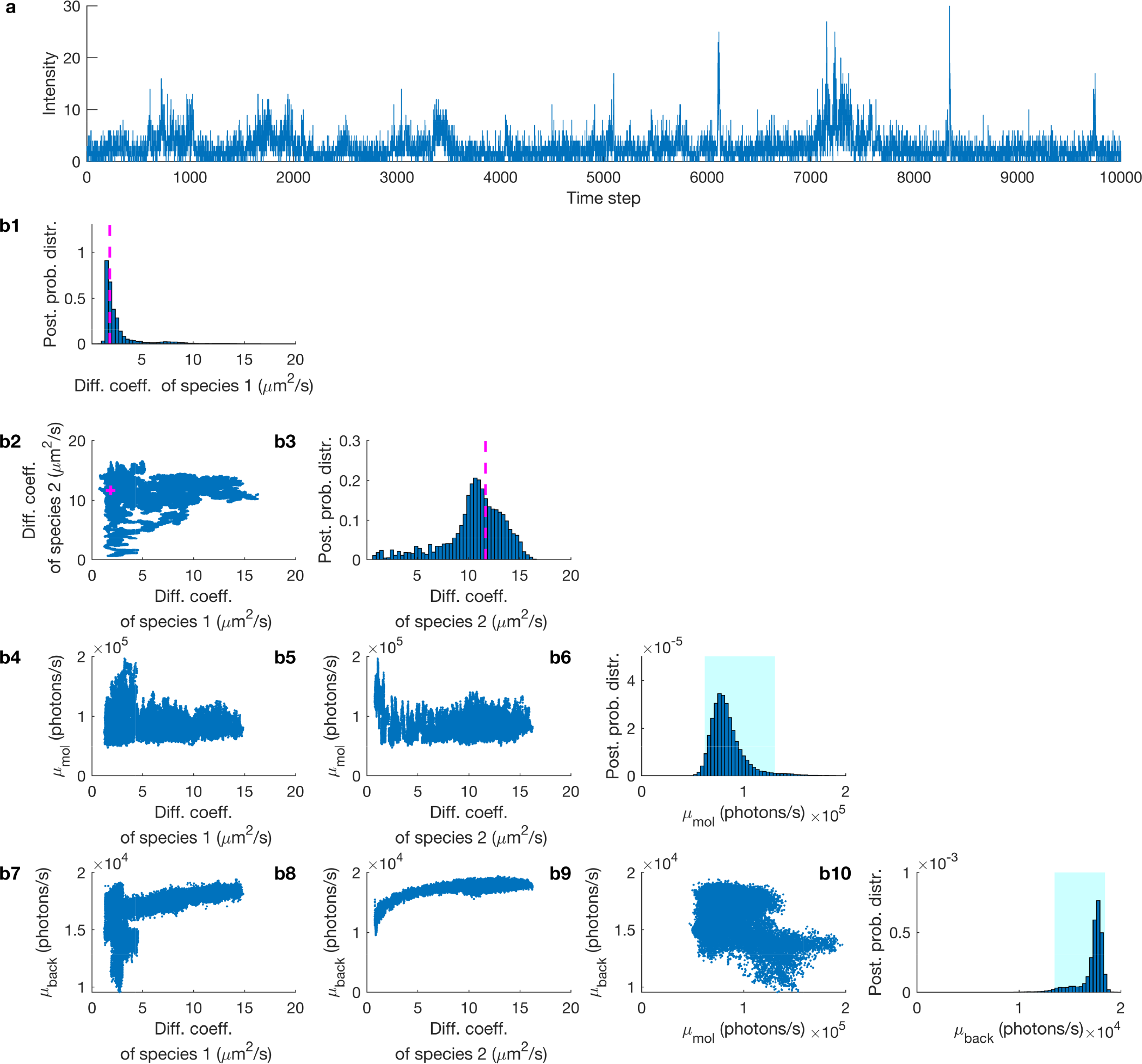
Estimated joint posterior probability distribution of multiple diffusive species. **(a)** Experimental fluorescent intensity trace used in Fig. 7 a3 with length 10^4^ data points and step 100*µs*. **(b1)** The posterior probability distribution of the diffusion coefficient of diffusive species 1. **(b2)** The joint probability distribution of diffusion coefficient of diffusive species 1 and diffusive species 2. **(b3)** The posterior probability distribution of the diffusion coefficient of diffusive species 2. **(b4)** The joint probability distribution of diffusion coefficient of diffusive species 1 along with the molecular brightness. **(b5)** The joint probability distribution of diffusion coefficient of diffusive species 2 along with the molecule photon emission rate. **(b6)** The posterior probability distribution of the molecular brightness. **(b7)** The joint probability distribution of diffusion coefficient of diffusive species 1 and background photon emission rate. **(b8)** The joint probability distribution of diffusion coefficient of diffusive species 2 and background photon emission rate. **(b9)** The joint probability distribution of the molecular brightness and background photon emission rate. **(b10)** The posterior probability distribution of the background photon emission rate. The trace is generated by mixing two experimental traces of concentration 1 nM of freely diffusive Cy3 in a water/glycerol mixtures with 94% and 75% glycerol each. The laser power, wavelength, NA and refractive index are 100 *µ*W, 532 nm, 1.42 and 1.4, respectively. The FCS estimates are shown by a magenta dashed lines and the 95% confidence intervals of the posteriors are highlighted in cyan.

## S2. Summary of point estimates

**TABLE S1.**
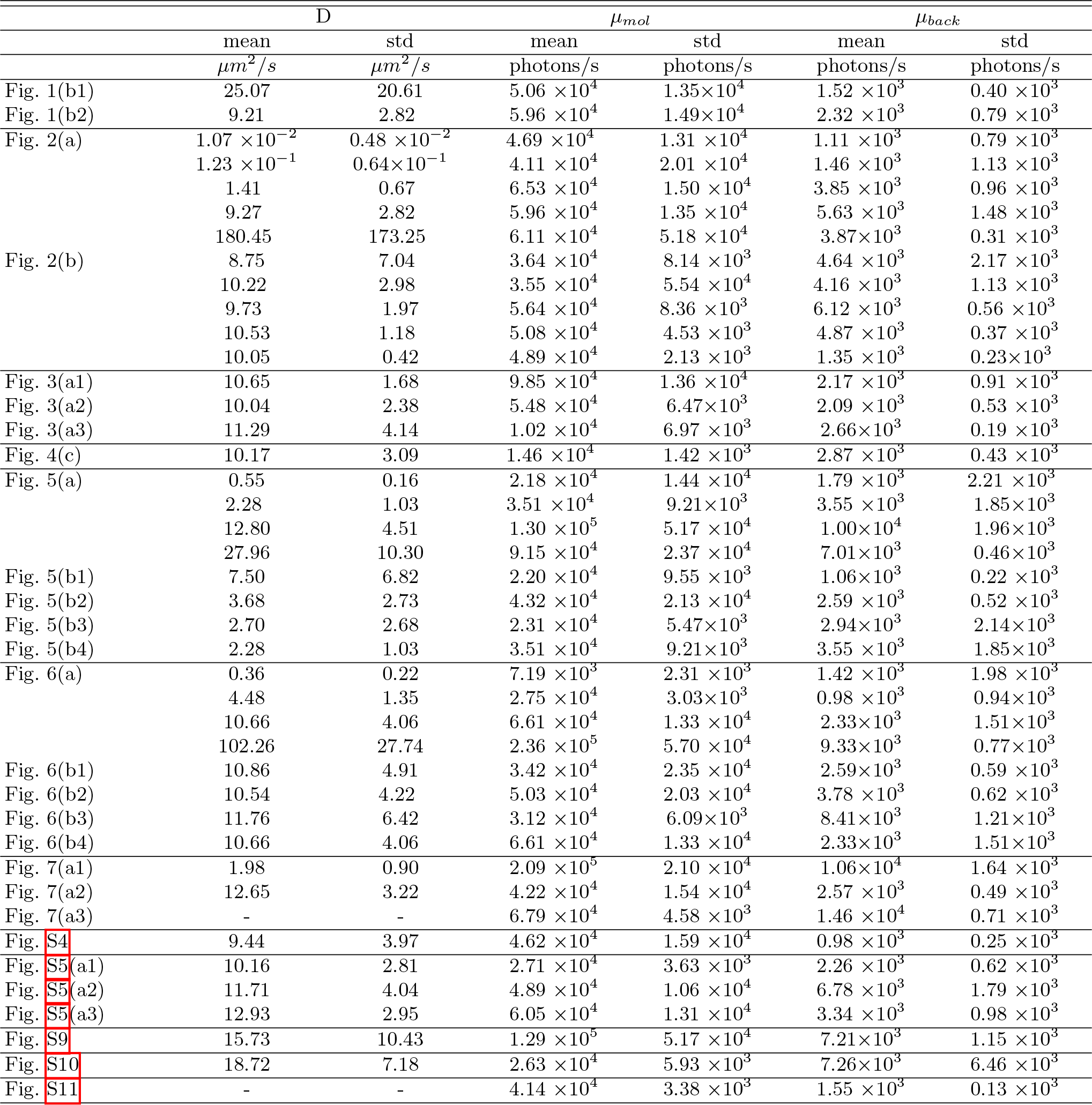
Here we list characteristic values (point estimates) summarizing the posterior probability distributions of this study. Mean and std refer to posterior mean value and standard deviation (i.e. square root of variance). Values are listed according to figures.

## S3. Detailed methods description

### S3.1. Representation of molecular diffusive motion

Consider a particle moving in *1D diffusion*. The probability distribution *p*(*x, t*) of the particle’s location obeys Fick’s second law [1–3] and is given by the diffusion equation

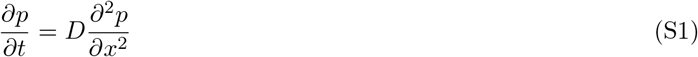

where *D* is the particle’s diffusion coefficient. Assuming the particle is located at *x*_*k*−1_ at a time *t*_*k*−1_, i.e. assuming the initial condition *p*(*x, t*_*k*−1_) = *δ*(*x − x*_*k*−1_), and a free space boundary, i.e. lim_*x*→±∞_*p*(*x, t*) = 0, we can solve this equation to obtain *p*(*x, t*) for any later time *t*. The solution is

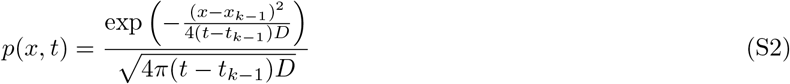

which equals to the probability density of a normal random variable with mean *x*_*k*−1_ and variance 2(*t − t*_*k*−1_)*D*, see Table S5. At time *t* = *t*_*k*_, we therefore have

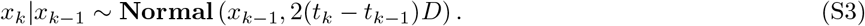

Similarly, solving the diffusion equation for particles following isotropic *3D diffusion* in free space, we have

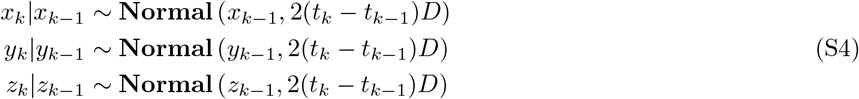

which constitute the molecular motion model used throughout this study.

### S3.2. Description of Stokes-Einstein model

For the experimental data, we benchmark our estimates of the diffusion coefficient against the Stokes-Einstein prediction [2, 3]. Namely, for a spherical particle in a quiescent fluid at uniform temperature

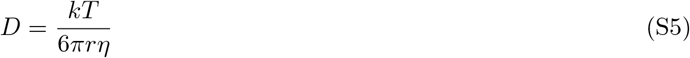

where, *D* is the diffusion coefficient, *k* is Boltzmann’s constant, *T* is the solution’s absolute temperature, *r* is the hydrodynamic radius of the particle [4] and *η* is the solution’s dynamic viscosity [5].

### S3.3. FCS formulation

The formulation we used in this study to autocorrelate the synthetic and experimental time traces is

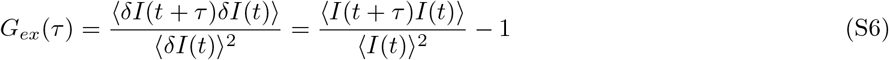

where the *I*(*t*) is the number of detected photons at time *t*. The computational implementation uses the Wiener-Khinchin Theorem [6].

Also, the theoretical function [7–10] used to fit the autocorrelation curves (using a 3DG PSF) is

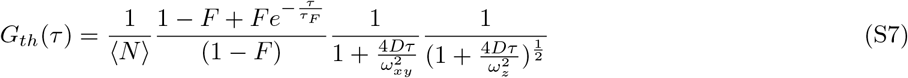

and for the 2DGL PSFs is

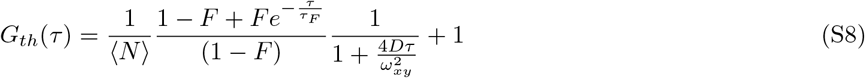

where, 〈*N*〉 is the average number of molecule in the effective volume, *D* is the diffusion coefficient, *τ*_*F*_ is the triplet state relaxation time and *F* is the fraction molecules populating the triplet state.

To find the best fit, we use *χ*^2^ minimization

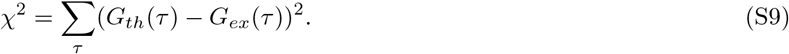

### S3.4. Definition of molecular brightness

As the definition of molecular brightness, we use the *emission rate of detected photons* of a single fluorophore, for example Eq. (2). For a fluorophore located at (*x, y, z*) this is formulated as

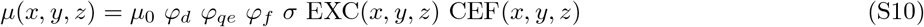

where, *µ*_0_ is the maximum excitation intensity which occurs at the center of the confocal volume, *φ*_*d*_ is the efficiency of the photon collection at the center of the confocal volume, *φ*_*qe*_ is the quantum efficiency of the detector, *φ*_*f*_ is the quantum efficiency of the fluorophore (i.e. quantum yield), *σ* is the absorption cross-section of the fluorophore, EXC(*x, y, z*) is the excitation profile and CEF(*x, y, z*) is the detection profile, i.e. collection efficiency function, which equals the fraction of the detected photons to the total photons emitted by a point source [11].

To obtain Eq. (2), we cast Eq. (S10) in the simplified form

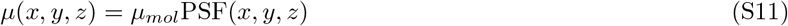

where *µ*_*mol*_ = *µ*_0_ *φ*_*d*_ *φ*_*qe*_ *φ*_*f*_ *σ*, which we term *molecular brightness* at the center of the confocal volume [12], and PSF(*x, y, z*) = EXC(*x, y, z*) CEF(*x, y, z*), which we term the PSFPSF.

To relate the parameter *µ*_*mol*_ to the *average photon count rate*, which is commonly estimated in bulk experiments [13**?**, 14], we consider the spatial average of *µ*(*x, y, z*) as follows

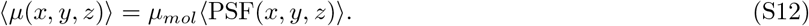

For the specific choice of a 3DG PSF (see below), the average is computed as follows

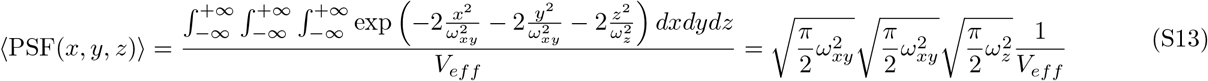

where *V*_*eff*_ denotes the effective volume of 3DG PSF [9, 15] and it is given by

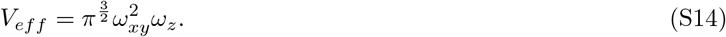

Consequently, Eq. (S13) implies

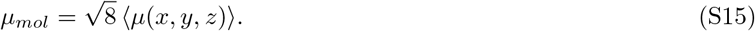

In other words, the molecular brightness is, by definition, approximately 2.8 times larger than the average photon count rate of single molecule [13**?**, 14].

### S3.5. Definition of point spread function models

In this study we use three different point spread functions as approximations to the more realistic Airy function [16–18], namely a 3D-Gaussian (3DG) [19], a 2D-Gaussian-Cylindrical (2DGC) [19] and a 2D-Gaussian-Lorentzian (2DGL) [20–23].

The definition of the PSF for the 3DG case is

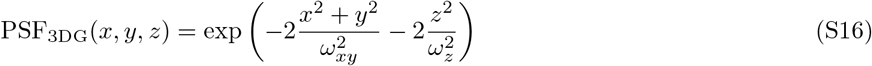

while, the definition of the PSF for the 2DGC case is

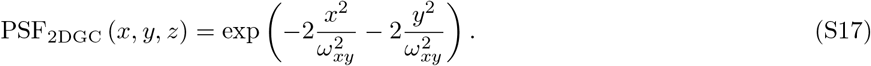

For both cases, *ω*_*xy*_ and *ω*_*z*_ are the semi-axes lateral and parallel to the optical axis. These are represented in terms of the excitation wavelength *λ*_exc_, solution refraction index *n*_sol_, and numerical aperture NA of the microscope as *ω*_*xy*_ = 0.61*λ*_exc_/NA and *ω*_*z*_ = 1.5*n*_sol_*λ*_exc_/NA^2^; for example see [24, 25]. For more realistic representations, *ω*_*xy*_ and *ω*_*z*_ can be estimated directly based on calibration experiments with known diffusion coefficients; for example see [26].

The definition of the PSF for the 2DGL case is

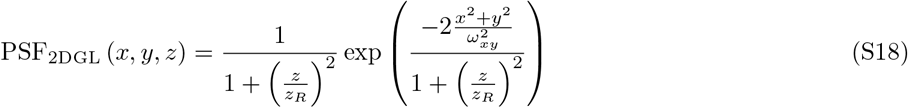

where *ω*_*xy*_, *λ*_exc_, and *n*_sol_ are similar to the 3DG cas or 2DG cases and 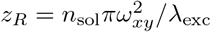.

### S3.6. Description of the data simulation

To generate fluorescence intensity time traces that mimic a realistic confocal setup, we simulate molecules moving [1**?**] through an illuminated 3D volume. The number of moving molecules *N* is prescribed in each simulation. To maintain a relatively stable concentration of molecules near the confocal volume, and so to avoid generating traces where every molecule eventually strays into un-illuminated regions, we impose periodic rectangular boundaries to our volume. The boundaries are placed at ±*L*_*xy*_ perpendicular to the focal plane and ±*L*_*z*_ perpendicular to the optical axis.

We assess the locations of the molecules 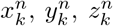, where *k* = 1, …, *K* label time levels and *n* = 1, …, *N* label molecules, at equidistant time intervals *t*_1_, *t*_2_, …, *t*_*K*_. The time interval between successive assessments *δt* = *t*_*k*_ − *t*_*k*−1_, as well as the total trace duration *T*_*total*_ = *t*_*K*_ − *t*_0_, are prescribed.

Molecule locations at the first assessment 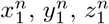 are sampled randomly from a uniform distribution with limits equal to the boundaries ±*L*_*xy*_ and ±*L*_*z*_ of our pre-specified volume. Subsequent locations are generated according to the diffusion model described above under a prescribed diffusion coefficient *D*.

Finally, we obtain individual photon emissions *w*_*k*_ by simulating Bernoulli random variables of success probability *q*_*k*_ = 1 − *e*^−*µ*_*k*_*δt*^, where the rate *µ*_*k*_ gathers single photon contributions from the background and the entire molecule population according to

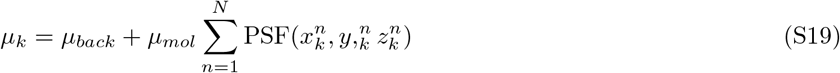

where both background and molecular brightness, *µ*_*back*_ and *µ*_*mol*_, are prescribed.

The PSF model is also prescribed. To avoid artifacts induced by the periodic boundaries we impose in our volume, we ensure that *L*_*xy*_ ≫ *ω*_*xy*_, *L*_*z*_ ≫ *ω*_*z*_, or *L*_*z*_ ≫ *z*_*R*_, where *ω*_*xy*_, *ω*_*z*_ and *z*_*R*_ characterize the geometry of the confocal volume, see Eqs. (S16)–(S18), above.

Detailed parameter choices for all simulations performed are listed in Table S6.

### S3.7. Definition of normalized distance and numbers of molecules

As we need to estimate the positions of the molecules with respect to the center of the confocal volume, which is the point of origin, in order to ultimately estimate the number of molecules as a proxy for molecule concentration, for example Figs. S3 and S8, we must address difficulties associated with symmetries of the confocal PSF with respect to rotations around the optical axis or the focal plane. [27] For this, for a molecule at 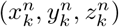, when the 3DG PSF is used, Eq. (S16), we rely on

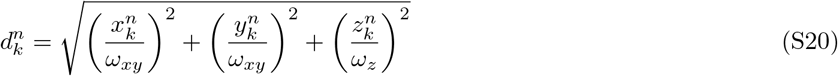

while, when the 2DGL PSF is used, Eq. (S18), we rely on

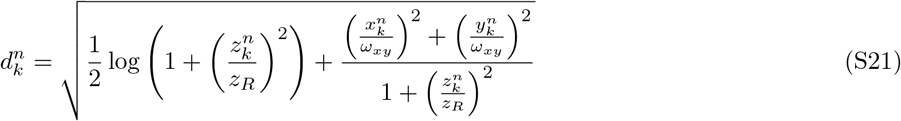

where 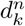 is the normalized distance with respect to the center of the confocal volume of molecule *n* at time *k*. Similarly, when a 2DGC PSF is used, Eq. (S17), we rely on

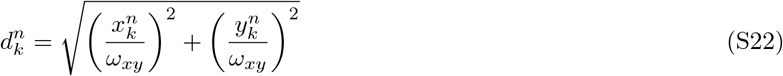

where 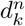 is the normalized distance with respect to the optical axis of molecule *n* at time *k*.

These distances are obtained by setting the respective PSFs equal to 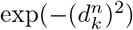 and are unaffected by the aforementioned symmetries, i.e. 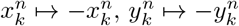, and 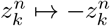.

For a given normalized distance ℓ, we define the number of molecules 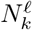 as the number of estimated (active) molecules within the corresponding distance. That is

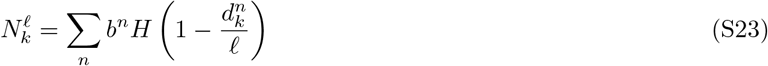

where *H* is the Heaviside step function, *b*^*n*^ is the load of molecule *n*, and *V*_*ℓ*_ is the volume of a designated effective region chosen to agree with the effective volume *V*_*eff*_ used in FCS [9].

### S3.8. Description of the time trace preparation

The initial time trace consists of single photon arrival times which are computationally too expensive to analyze. Our method instead operates on photon intensity traces which are either obtained directly during an experiment or obtained from individual photon arrival time traces after binning. To transform single photon arrival time traces into intensity time traces, we use time bins of fixed size (main size) that typically span multiple photon arrival times. To speed up the computations further, as some bins have none of very few photons, over certain portions of the trace we use larger bins (auxiliary size).

Briefly, the user specifies a minimum number of photons per bin as a lower threshold. As illustrated in Fig. S13, those bins, preselected at the main size, containing fewer photons than the specified threshold are enlarged uniformly in order to achieve an average of at least as many photons as specified by the threshold. This occasional adaptation, from the main to the auxiliary bins, becomes important in the analysis of traces from experiments held near single molecule resolution where molecule concentrations are low so that on average only one molecular passage through the confocal volume happens. Consequently, photon intensities are low, and thus the bulk of computational time otherwise would had been spent processing trace portions of poor quality (i.e. with few or no photons).

**FIG. S13.**
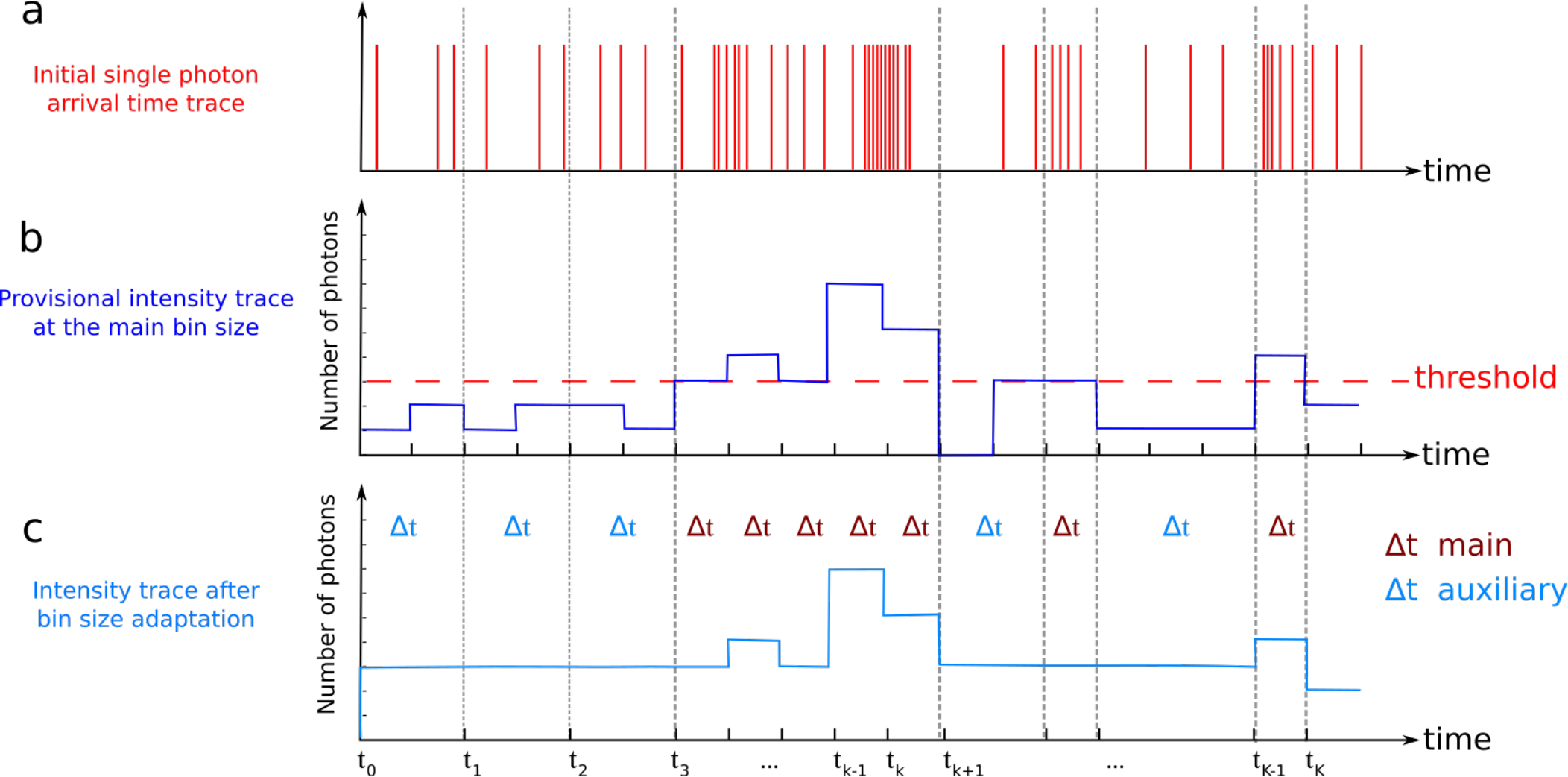
Illustration of time trace preparation. **(a)** Initial trace of single photon arrivals. Here, each vertical line represents the arrival time of a single photon. **(b)** Time trace of photon intensities provisionally binned at the main bin size. The horizontal line denotes the imposed lower threshold on the minimum number of photons in the individual bins. **(c)** Time trace of photon intensities after bin size adaptation. Here, bins, preselected at the main size, with intensities below the imposed threshold are uniformly readjusted to achieve an average intensity similar to the threshold.

To carry our the necessary computations, as we detail shortly, we use the Anscombe transformation [28] to approximate the Poissonian likelihoods of photon intensities (see below). This approximation is robust as long as bins contain on average 4 photons or more. Thus, as a minimum requirement, we also use the aforementioned threshold to ensure the validity of the approximations.

## S4. Detailed description of the inference framework

**FIG. S14.**
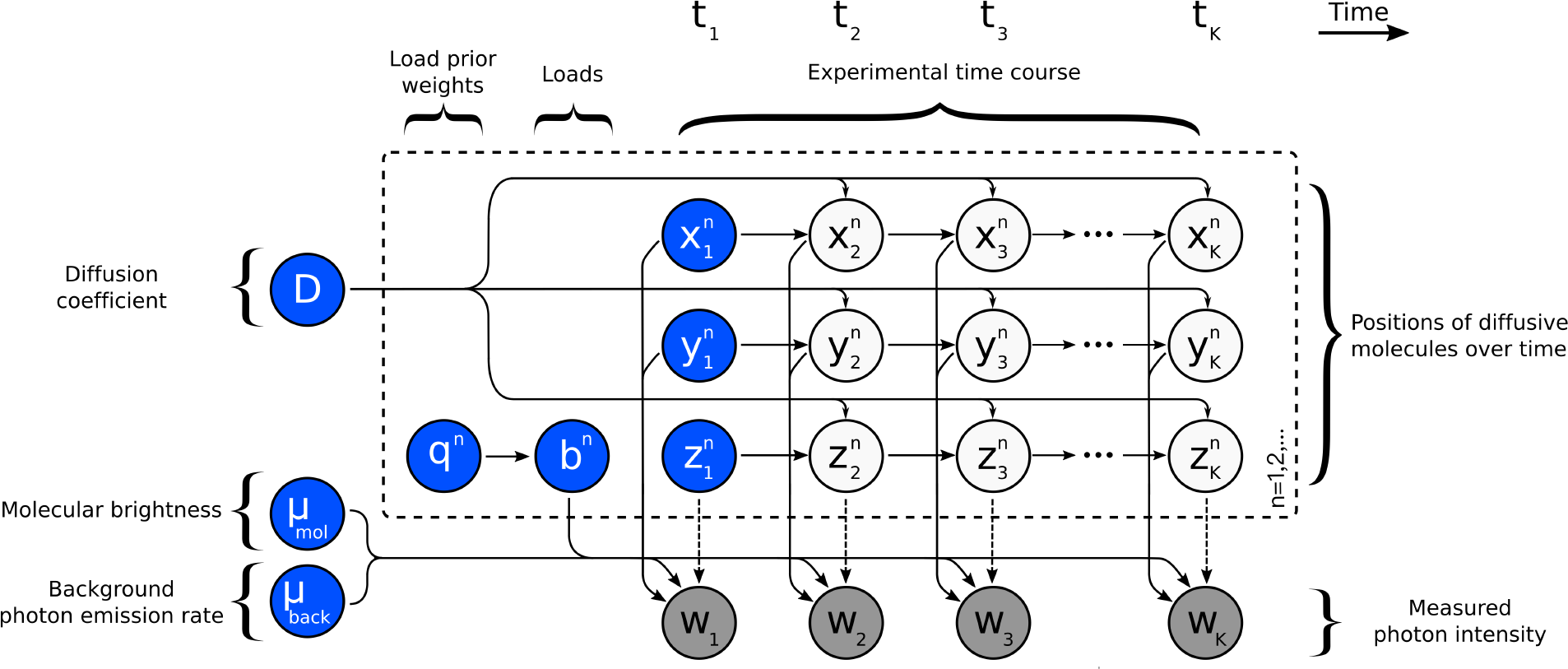
Graphical summary of the framework developed. A population of model molecules, labeled by *n* = 1, 2, …, evolves during the measurement period which is marked by *k* = 1, 2, …, *K*. Here, 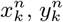 and 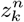 denote the position of molecule *n* at time *t*_*k*_ and *µ*_*mol*_, *µ*_*back*_ denote the molecular brightness and background photon emission rates. The diffusion coefficient *D* determines the evolution of the molecule locations which, in turn, determines the instantaneous photon emission rates and ultimately the recorded photon intensities *w*_*k*_. Load variables *b*^*n*^, with prior weights *q*^*n*^, are introduced to model a molecule population of *a priori* unknown size. Following common machine learning convention, the measurements *w*_*k*_ are dark shaded. Additionally, model variables requiring prior probability distributions are highlighted in blue.

### S4.1. Description of prior probability distributions

The model parameters in our framework that require priors are: the diffusion coefficient *D*; the molecular brightness and background photon emission rates *µ*_*mol*_ and *µ*_*back*_; the initial molecule locations 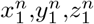; and load prior weights *q*^*n*^. As we already mentioned in the main text, a prior on the population of diffusing molecules is implicitly defined by the prior on both *b*^*n*^ and *q*^*n*^. Our choices are described below.

#### S4.1.1. Prior on the diffusion coefficient

To ensure that the *D* sampled in our formulation attains only positive values, we place an inverse-Gamma prior

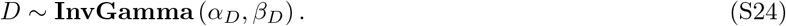

Besides ensuring a positive D, this prior is also conjugate to the motion model we use which facilitates the computations (see below).

#### S4.1.2. Priors on molecular brightness and background photon emission rates

To ensure that *µ*_*mol*_ and *µ*_*back*_ sampled in our formulation attain only positive values, we place Gamma priors on both

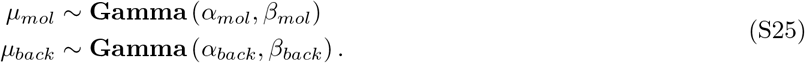

Due to the specific dependencies of the likelihood (that we will discuss shortly) on the photon emission rates, conjugate priors cannot be achieved for *µ*_*mol*_ and *µ*_*back*_. So, the above choice offers no computational advantage (see below) and could be readily replaced with more physically motivated choices if additional information on molecular brightness becomes available.

#### S4.1.3. Priors on initial molecule locations

Due to the symmetries in the confocal PSF, i.e. a molecule at a location (*x, y, z*) emits the same average number of photons as a molecule at locations (±*x*, ±*y*, ±*z*), we are unable to gain insight regarding the octant of the 3D Cartesian space in which each molecule is located. To avoid imposing further assumptions on our framework that may determine each molecule’s octant uniquely, but may limit the framework’s scope to specific experimental setups, we place priors on the initial locations that respect these symmetries. Accordingly, in our framework, at the onset of the measuring period, molecules are equally likely to be located at any of the positions 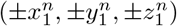.

To facilitate the computations (see below), we place independent symmetric normal distributions, see Table S5, on each Cartesian coordinate of the model molecules

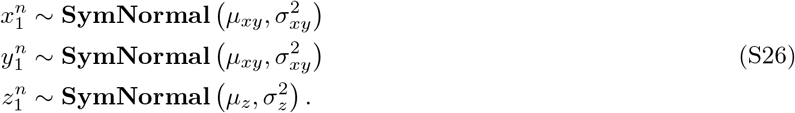

We want to emphasize that the symmetric priors above do not affect our estimates. According to the motion model we employ, no matter where molecules are initiated, they may subsequently move freely and eventually switch to a different octant if warranted by the data. Our symmetric priors merely indicate that for each individual molecular trajectory considered, there are another 7 symmetric trajectories that are equally likely to have occurred.

#### S4.1.4. Priors and hyperpriors for molecule loads

To facilitate the computations (described next), we use a finite, but large, model population consisting of *N* molecules containing both active and inactive molecules. These model molecules are collectively indexed by *n* = 1, 2, …, *N*. As explained in the main text, estimating how many molecules are actually warranted by the data under analysis is equivalent to estimating how many of those *N* molecules are active, i.e. *b*^*n*^ = 1, while the remaining inactive ones, i.e. *b*^*n*^ = 0, have no impact whatsoever and are instantiated only for computational purposes.

To ensure that each load *b*^*n*^ takes only values 0 or 1, we place a Bernoulli prior of weight *q*^*n*^. In turn, on each weight *q*^*n*^, we place a conjugate Beta hyperprior

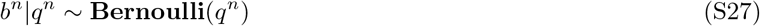

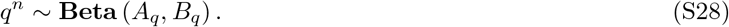

To ensure that the resulting formulation avoids overfitting, we make the specific choices *A*_*q*_ = *α*_*q*_/*N* and *B*_*q*_ = *β*_*q*_(*N* − 1)/*N*. Under these choices [29–32], and in the limit that *N* → ∞; that is, when the assumed molecule population is allowed to be large, this prior/hyperprior converge to a Beta-Bernoulli process. Consequently, for *N* ≫ 1, the posterior remains well defined and becomes independent of the chosen value of *N*. In other words, provided *N* is large enough, its impact on the results is insignificant; while its precise value has only computational implications (see below).

### S4.2. Summary of model equations

For concreteness, below we summarize the entire set of equations used in our framework, including a complete list of priors and hyperpriors

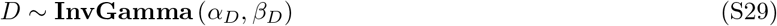

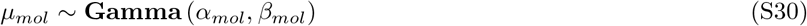

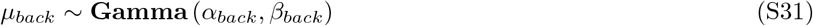

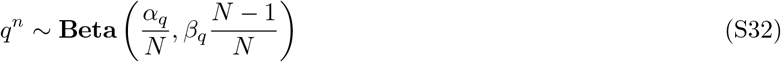

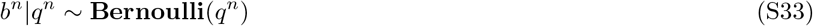

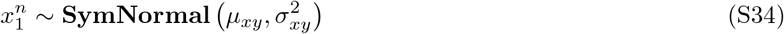

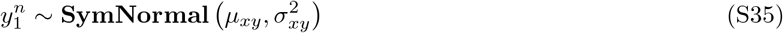

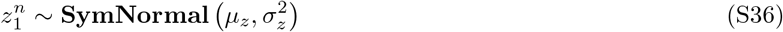

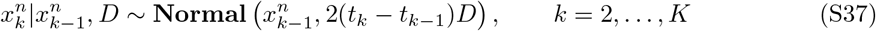

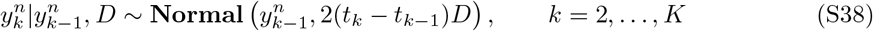

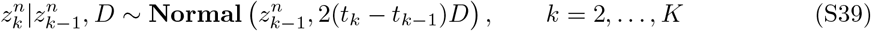

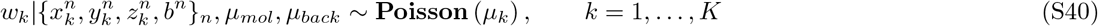

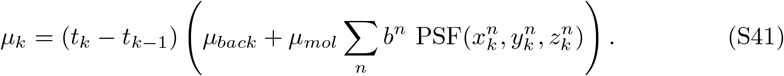

For molecules diffusing in a confocal volume that is extremely elongated over the optical axis, the PSF approaches a cylindrical one. In this case, it is safe to eliminate the 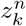 positions from the motion model and simplify Eqs. (S40) and (S41) to

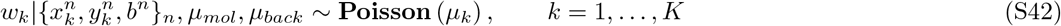

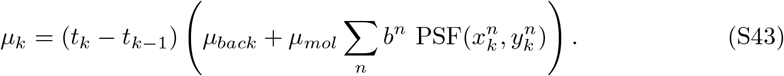

### S4.3. Description of the computational scheme

The joint probability distribution of our framework is 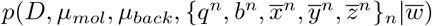, where molecular trajectories and intensities (measurements) are gathered in

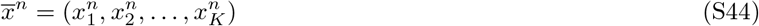

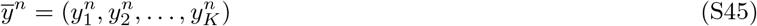

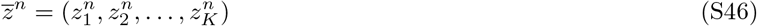

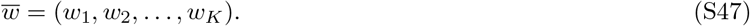

Due to the non-linearities in the PSF and the non-parametric prior on *q*^*n*^ and *b*^*n*^, analytic evaluation or direct sampling of this posterior is impossible. For this reason, we develop a specialized Markov chain Monte Carlo (MCMC) scheme that can be used to generate pseudo-random samples [33–37]. This scheme is explained in detail below.

In order to terminate the MCMC sampler, we need to determine when a representative number of samples has been computed. To do so, we divide the samples already computed into four portions and compare the mean values of the diffusion coefficient of the two last ones

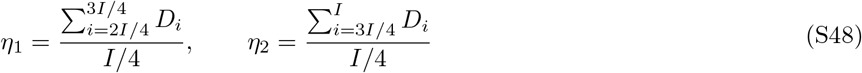

where, *η*_1_ and *η*_2_ are the mean values of the two last portion of the sampled diffusion coefficients denoted *D*_*i*_ and *I* is the total number of computed MCMC samples thus far. Following [33, 34], we terminate the sampler when |*η*_1_ − *η*_2_| < *ε*_thr_, where *ε*_*thr*_ is a pre-specified threshold. Also, to avoid incorporating burn-in samples in the calculations, we ensure a minimum number of iterations *I* of no less than 10^4^.

A working implementation of the resulting scheme in source code and GUI forms, see Fig. S15, are available through the Supporting Material.

**FIG. S15.**
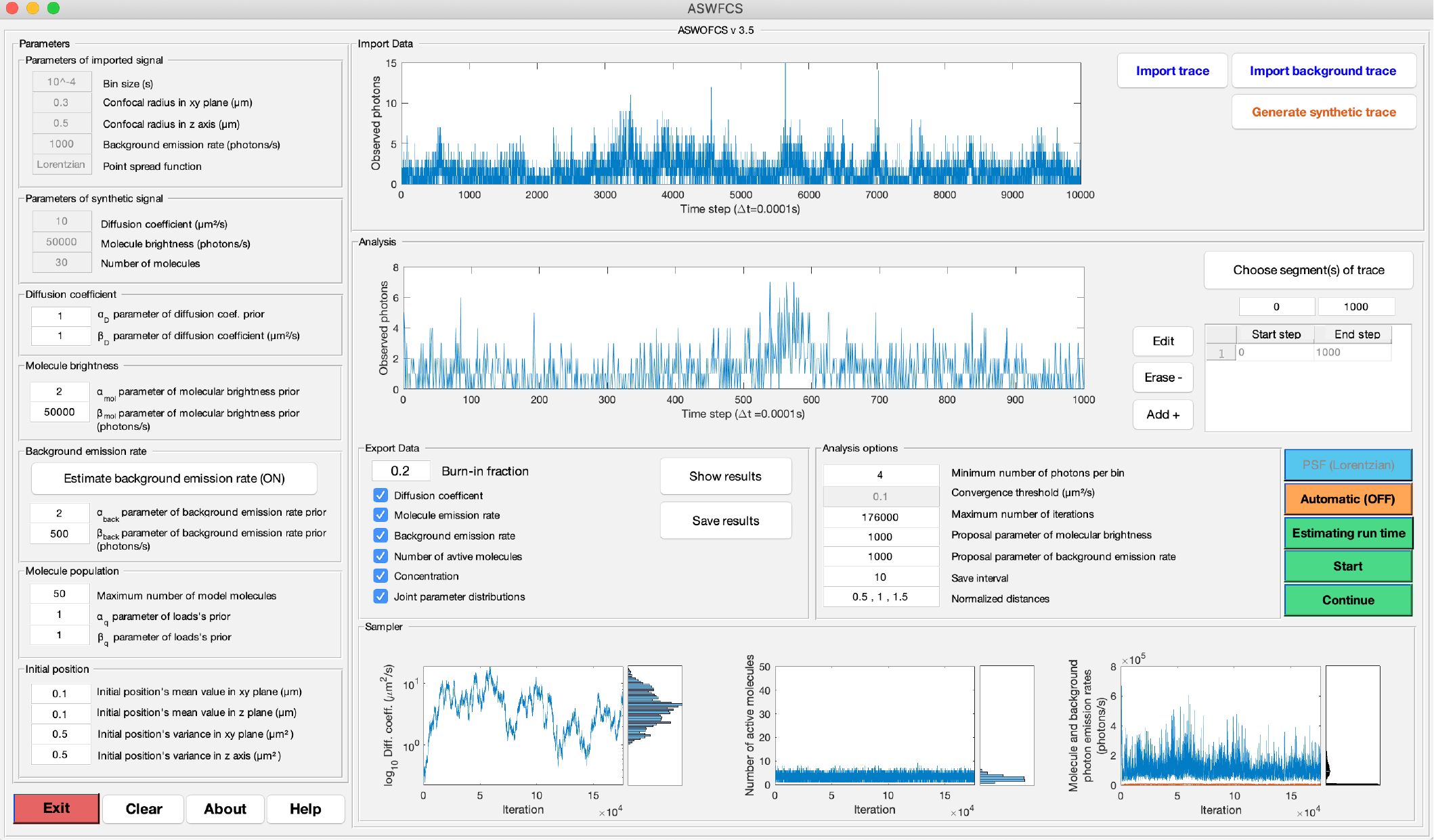
A working implementation of the framework described in this study is available through the Supporting Material. Along with this implementation, we provide a graphical user interface (GUI) that can be used to analyze intensity traces from confocal microscopy.

#### S4.3.1. Overview of the sampling updates

Our MCMC exploits a Gibbs sampling scheme [33–35]. Accordingly, posterior samples are generated by updating each one of the variables involved sequentially by sampling conditioned on all other variables and measurements 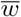.

Conceptually, the steps involved in the generation of each posterior sample 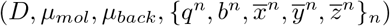 are:

1. For each *n* in the molecule population

i. Update trajectory 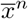 of molecule *n*
ii. Update trajectory 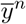 of molecule *n*
iii. Update trajectory 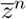 of molecule *n*
2. Update the diffusion coefficient *D*
3. Update jointly the prior weights *q*^*n*^ for all molecules
4. Update jointly the loads *b*^*n*^ for all molecules
5. Update jointly the molecular brightness and background photon emission rates *µ*_*mol*_ and *µ*_*back*_, respectively

Since the locations of the inactive molecules are not associated with the measurements 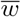, see Fig. S14, and those are updated independently of the locations of the active ones, to make the algorithm computationally more efficient we carry out the above scheme in the equivalent order

1. For each *n* of the *active* molecules

i. Update trajectory 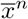 of active molecule *n*
ii. Update trajectory 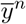 of active molecule *n*
iii. Update trajectory 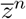 of active molecule *n*
2. Update jointly the trajectories 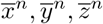 for all *n* of the *inactive* molecules
3. Update the diffusion coefficient *D*
4. Update jointly the prior weights *q*^*n*^ for all model molecules
5. Update jointly the loads *b*^*n*^ for all model molecules
6. Update jointly the molecular brightness and background photon emission rates *µ*_*mol*_ and *µ*_*back*_, respectively

These steps are described in detail below.

#### S4.3.2. Sampling of active molecule trajectories

For a given active molecule *n*, we update the trajectory 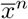 by sampling from the corresponding conditional 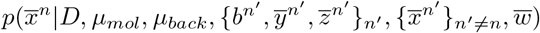, which we achieved through backward sampling [38–40]. To be able to sample a trajectory 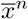 in backward sampling, we factorize the density 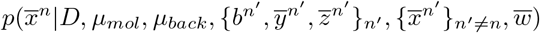 as

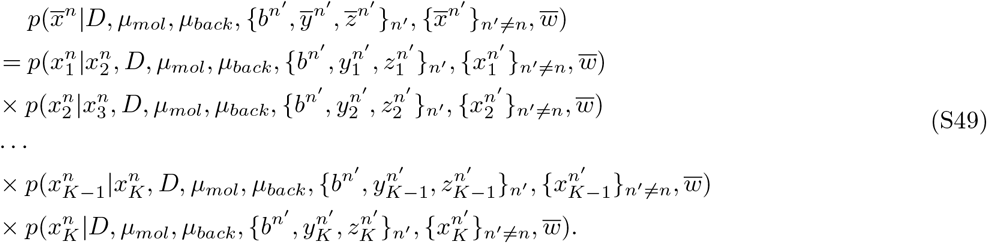

According to this factorization, we sample 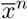, starting from 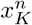 and move backward towards 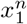. To start the sampling steps, we need to determine each one of the individual densities 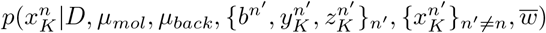 and 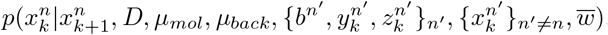. We do this in a forward filtering approach [27, 38–42] which is described in detail below.

##### S4.3.2.a. Forward filtering

By applying Bayes’ rule, each one of the individual densities in Eq. (S49) factorizes as

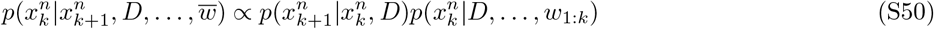

where *w*_1:*k*_ is an abbreviation for *w*_1_, …, *w*_*k*_ and excess parameters are shown by dots. Since the density 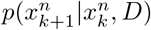 is already known, to sample 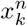 in backward sampling, we only need to determine the filter density 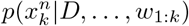.

To be able to apply forward filtering and compute 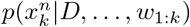 efficiently [43], we use an approximate model [44], where Eq. (S40), is replaced with

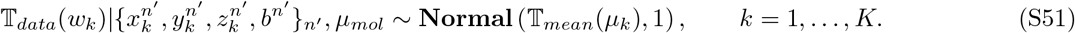

Here, *µ*_*k*_ stems from Eq. (S41) for 3D models and Eq. (S43) for 2D models; while, 𝕋_*data*_(*w*) and 𝕋_*mean*_(*µ*) denote Anscombe transformed [28] variables defined as follows

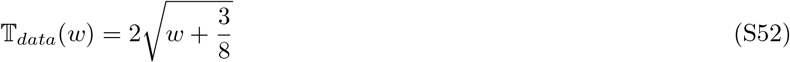

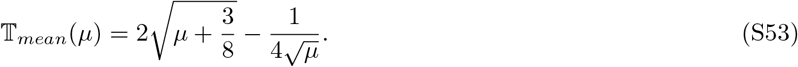

The Anscombe transform exploited here offers a way of transforming Poisson random variables into (approximately) normal ones [28] which facilitates the filtering process described next. The approximation we employ is highly accurate for *µ* ≫ 1, while acceptable accuracy is maintained so long as *µ* > 4 photons.

Under the Anscombe transform, the densities of both the dynamics, Eq. (S37), and observations, Eq. (S51), are normally distributed. So, we can compute the filter distribution 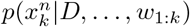 of the approximate model similar to the standard theory underlying nonlinear Kalman filters [27, 45–51].

More specifically, because the mean of the transformed observation distribution, *T*_*mean*_(*µ*_*k*_) is a nonlinear function of the location 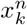, to apply the Kalman filters we need to approximate the transformed observation distribution in such a way that its mean becomes a linear function of the location 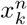. To do so, we use two common approaches: (i) extended Kalman filter (EKF) [45, 52–55], which locally approximate the transformed observation distribution around selected values; and (ii) unscented Kalman filter (UKF) [46–48], which globally approximate the transformed observation distribution.

As explained in detail in [27], the linearization alone is not sufficient to properly approximate the filter. This is because both EKF and UKF assume that the filter is a normal density. This assumption is problematic for our particular case which is symmetric across the origin, i.e. observations provide equal probabilities for the molecule to be in negative or positive side of the center of the PSF, i.e. 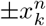. Due to this symmetry across the *yz*-plane, the filtering distribution consists of two modes centered symmetrically across the origin [27]. Therefore, we compute an approximate filter distribution of the form

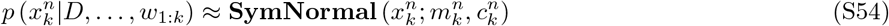

where **SymNormal** 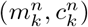 denotes the symmetric normal distribution (see Table S5). The filter’s parameters 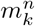 and 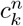 can be computed recursively according to

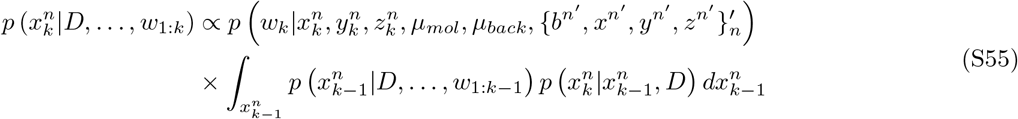

which, for our model, reduces to

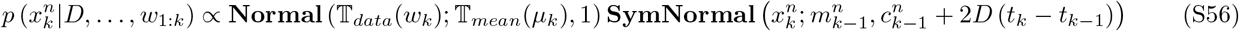

and, in turn, is approximated as

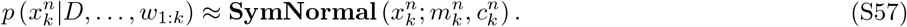

To summarize, in the forward pass of the FFBS, we compute 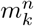 and 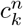 of the filter of the molecule *n*, for all time levels *k* = 1, …, *K*, by linearizing the approximate model around 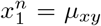 for *k* = 1, and around 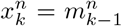 for *k* = 2, …, *K*. Since our observation is nonlinear, to calculate the filter, we opt between two different methods: **(i)** Extended Kalman filter (EKF) and **(ii)** Unscented Kalman filter (UKF).

In the EKF, we linearize the observations to obtain a closed form for the filter (local approximation) and in the UKF we approximate the joint probability distribution of observations and locations with a multivariate normal distribution (global approximation). The reason to use either of these filters is that the EKF is computationally cheaper but less accurate. According to our analysis it may fail to provide unbiased estimates of the background photon emission rate. On the other hand, the UKF is more robust and provides background emission rate estimates, but these benefits come at an increased computational cost.

In this study, we provide both filters and allow the user to choose between them.

###### Extended Kalman filter

Within the EKF approximation, the normal probability distribution preceding the symmetric normal of Eq. (S56) is linearized in order for their product to become a symmetric normal one. In this case, we linearize the mean of the observation density **Normal** (𝕋_*data*_(*w*_*k*_); 𝕋_*mean*_(*µ*_*k*_), 1), around the modes of the filter in the previous time step

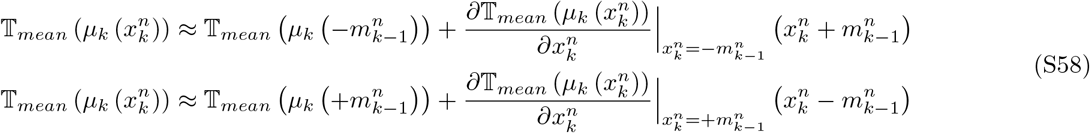

where the first term linearizes around 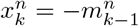 and the second term linearizes around 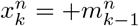. Under these approximations, (S56) attains an analytical solution. In detail

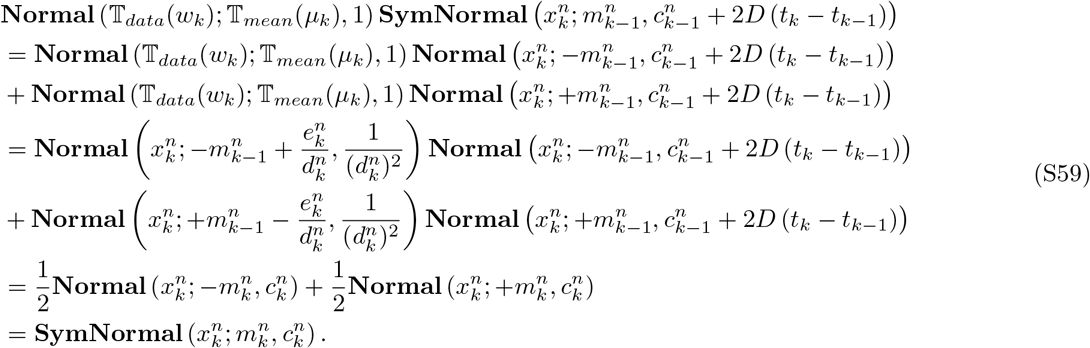

The same calculations apply also for *k* = 1, where the starting density is replaced with the prior of (S26). In this case

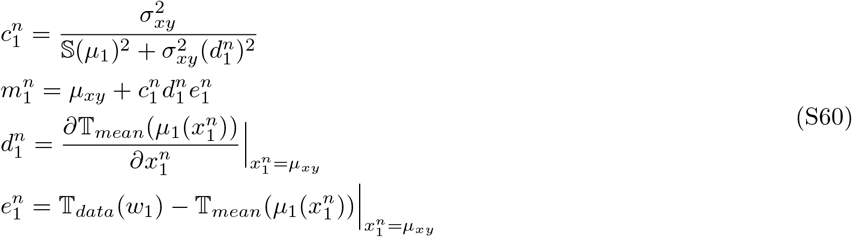

while for *k* = 2, …, *K* are

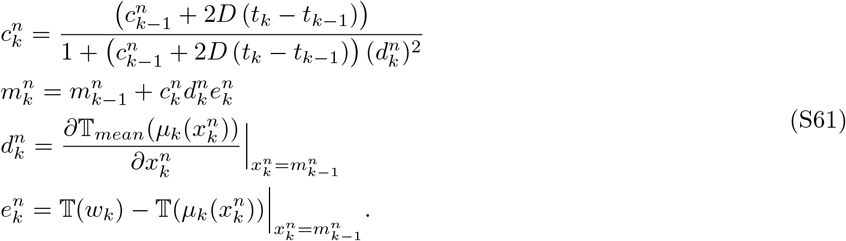

###### Unscented Kalman filter

The unscented Kalman filter [46, 47] tries to fit the joint probability distribution of the observations and locations globally with a multivariate normal distribution to cope with the nonlinearity in Eq. (S56). Specifically the product of Eq. (S56) is approximated as follows

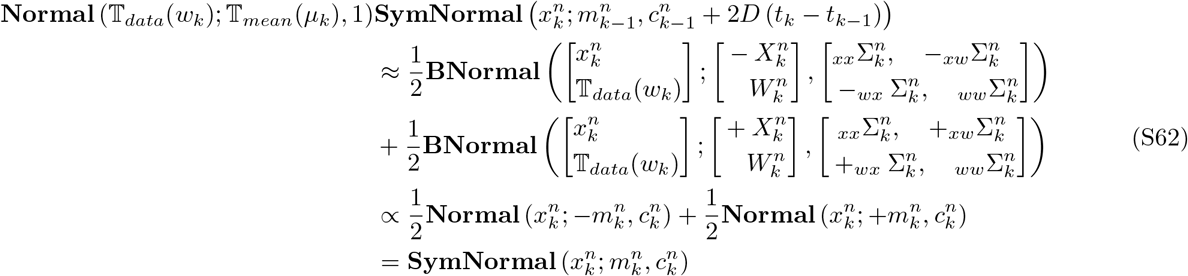

Since we are faced with a filter which has two symmetric modes, we calculate the filter’s mean 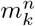 and variance 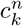 for one of the modes only, while we recover the other mode’s mean and variance by reflection.

**TABLE S2.** Sigma points and corresponding weights for a standard normal according to [59]

| $i$ | 1 | 2 | 3 | 4 | 5 | 6 | 7 | 8 | 9 | 10 | 11 |
| --- | --- | --- | --- | --- | --- | --- | --- | --- | --- | --- | --- |
| $x_i^{sn}$ | -5.1880 | -3.9362 | -2.8651 | -1.8760 | -0.9289 | 0 | 0.9289 | 1.8760 | 2.8651 | 3.9362 | 5.1880 |
| $g_i^*$ | $< 10^{-5}$ | 0.0002 | 0.0067 | 0.0661 | 0.2422 | 0.3694 | 0.2422 | 0.0661 | 0.0067 | 0.0002 | $< 10^{-5}$ |

The means, auto- and cross-covariances in one mode of the (S62) are given by

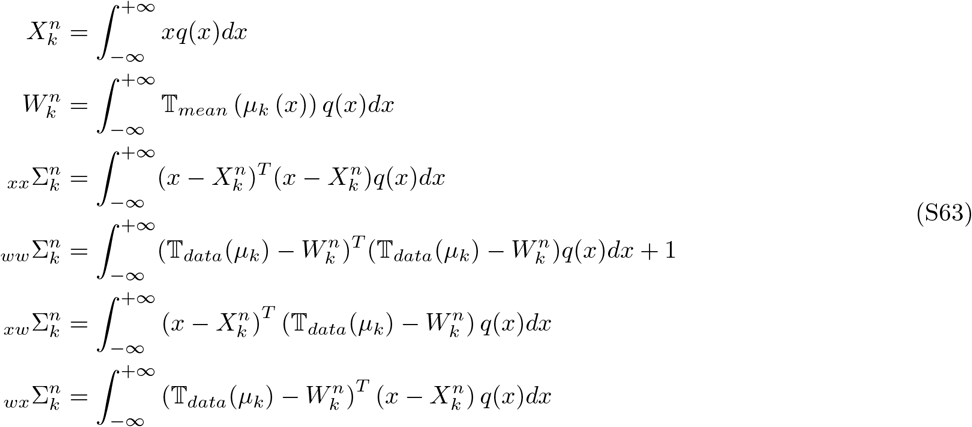

where 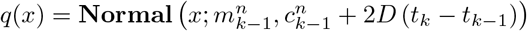 is the probability density of one mode of the filter. The same formula applies to the other mode too.

To calculate the mean value 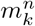 and variance 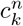 of each normal contributing to the symmetric normal shown above, we need to specify a set of sample points, termed *sigma points* in the UKF literature [46–48, 56–58], to estimate the mean values and covariance matrix of the bivariate normal on which 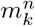 and 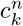 depend. To specify sigma points, we first calculate sigma points 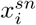 and their weights 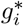 for a standard normal **Normal**(*x*; 0, 1) in Table. S2 according to [59]. We then transform 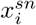 that will be used in this **Normal**(*x*; *m*_*k*−1_, *c*_*k*−1_ + 2*D*(*t*_*k*_ − *t*_*k*−1_)). The transformed sigma points are

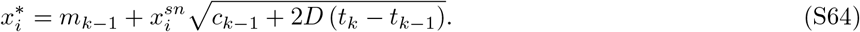

Finally, given 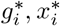, we calculate the mean and covariance of the bivariate normal previously introduced by

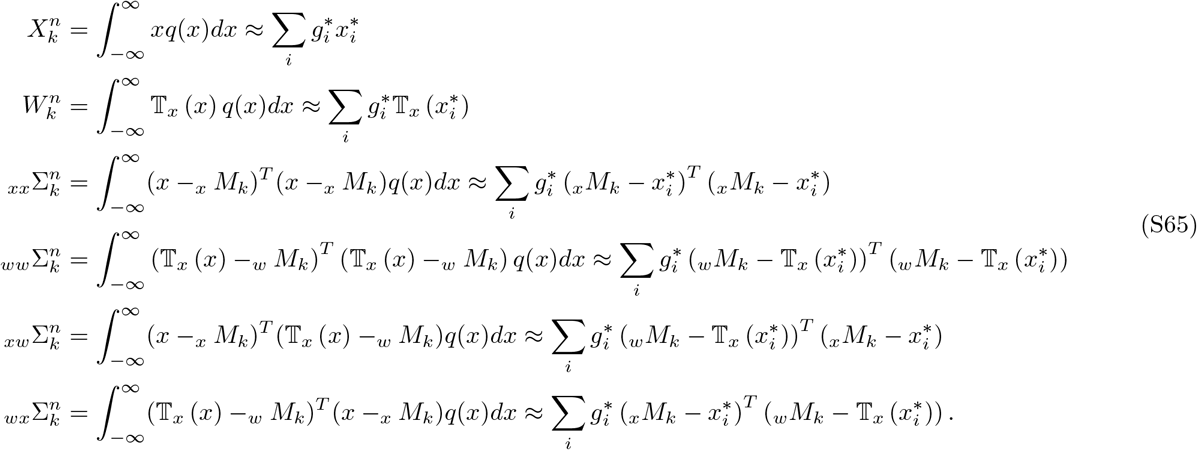

After computing the means 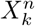 and 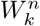 and auto-covariances and cross-covariances _*xx*_Σ_*k*_, _*ww*_Σ_*k*_, _*xw*_Σ_*k*_, _*wx*_Σ_*k*_, the mean and variance of each mode of the filter are given by

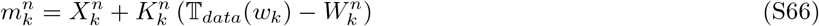

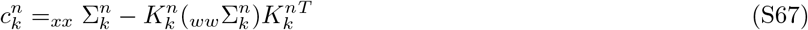

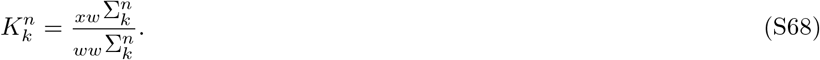

##### S4.3.2.b. Backward sampling

After we compute the filter densities 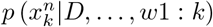 in the forward filtering step, through the EKF or UKF, we are able to sample the location 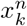 by using backward sampling as in Eq. (S50). Specifically, given a computed filter, we sample sequentially 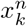 according to

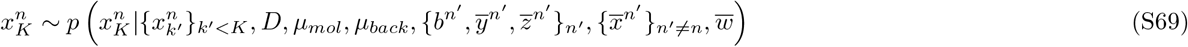

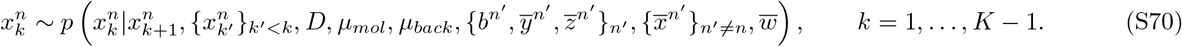

Due to the specific choices of our problem these reduce to

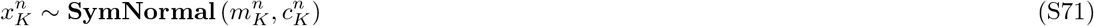

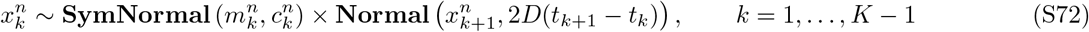

where 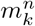 and 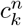 are the parameters of the filter which are calculated in the forward filtering step.

#### S4.3.3. Sampling of inactive molecule trajectories

After updating the trajectories of the active molecules, we update the trajectories of the inactive ones. For this, we sample from the corresponding conditionals 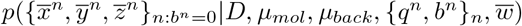. Since the locations of inactive molecules are not associated with the observations in 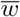 and hyper-priors {*q*^*n*^}_*n*_, these conditionals simplify to 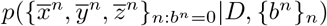 which can be readily simulated jointly in the same manner as standard 3D Brownian motion.

#### S4.3.4. Sampling of the diffusion coefficient

Now that we have updated the locations of active and inactive molecules, we update the diffusion coefficient *D* by sampling from the corresponding conditional 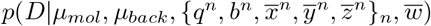, which, due to the specific dependencies of the variables in our formulation, e.g. Eqs. (S24), (S37), (S38) and (S39), simplifies to 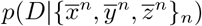. Because of conjugacy, the latter reduces to

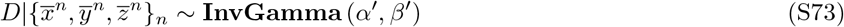

where *α′* and *β′* are given by

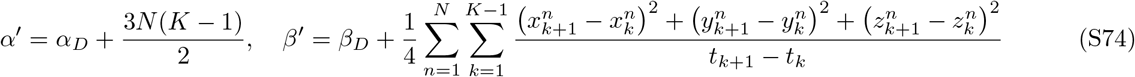

#### S4.3.5. Sampling of the molecule prior weights and loads

We update the load prior weights *q*^*n*^ by sampling from the corresponding conditional 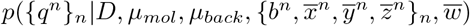, which simplifies to *p*({*q*^*n*^}_*n*_|{*b*^*n*^}_*n*_). For this, we use Eqs. (S33) and (S32), and because of conjugacy, the latter distribution is sampled by sampling each *q*^*n*^ separately according to

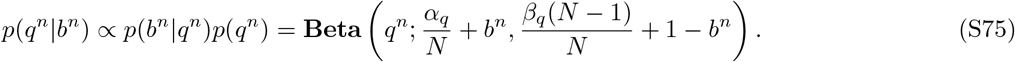

Once the weights *q*^*n*^ are updated, we update the loads *b*^*n*^ by sampling from the corresponding conditional 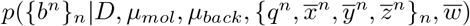. We perform this sampling using a Metropolis-Hastings update with proposals of the form

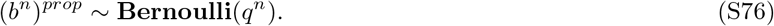

In this case, by choosing the proposal distribution similar to the prior distribution, the acceptance ratio becomes

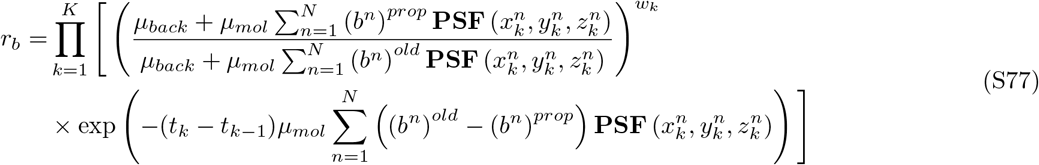

where (*b*^*n*^)^*old*^ denotes the existing sample.

#### S4.3.6. Joint sampling of the molecular brightness and background photon emission rates

Finally, after we updated the locations of molecules, and loads, we update the molecular brightness and background photon emission rates *µ*_*mol*_ and *µ*_*back*_ by sampling from the corresponding conditional 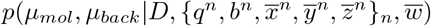, which simplifies to 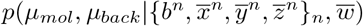. We carry over this sampling using a Metropolis-Hastings update where proposals for *µ*_*mol*_ and *µ*_*back*_ are computed according to

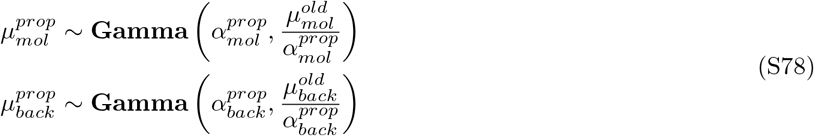

where 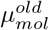 and 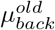 denote the existing samples. The acceptance ratio is

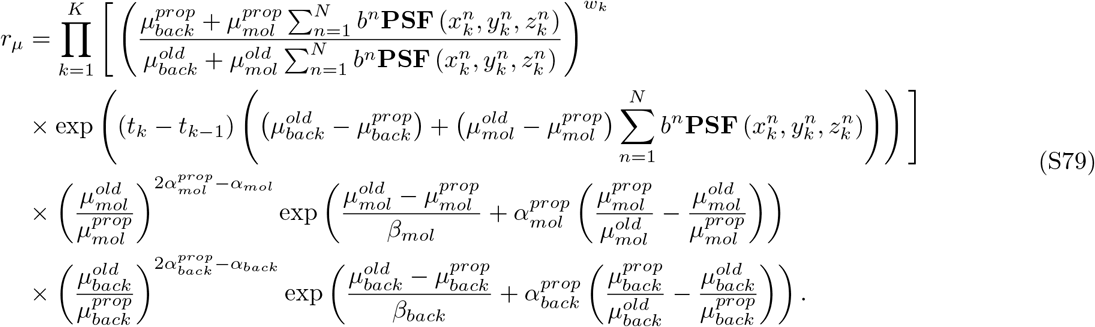

We should emphasize, due to the weakness of the extended Kalman filter as compared to the unscented Kalman filter, we consider the background photon emission rate is fixed for EKF. So, in this case we update the molecular brightness *µ*_*mol*_ only by sampling from the corresponding conditional 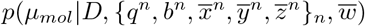, which simplifies to 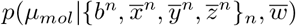. Again, we carry over this sampling using a Metropolis-Hastings update where proposals for *µ*_*mol*_ are computed according to

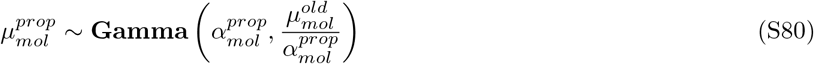

and the acceptance ration will reduces to

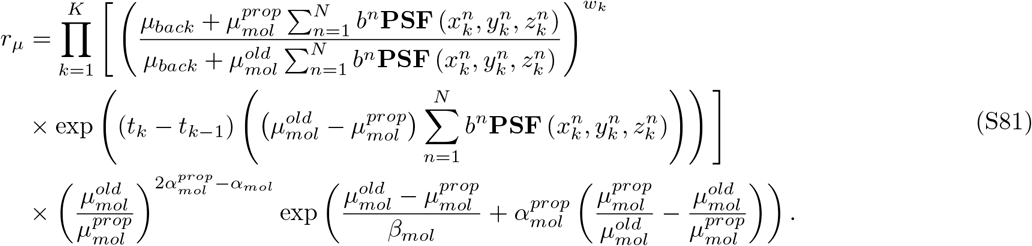

## S5. Extension for multiple diffusive species

In the case of more than one diffusive species, we can readily modify the model to capture multiple diffusion coefficients. To show that our method can be extended, we consider two diffusive species. Namely, the extended formulation is

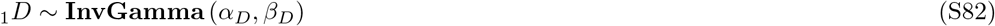

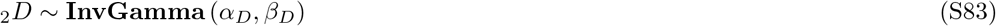

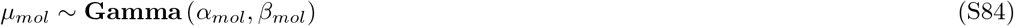

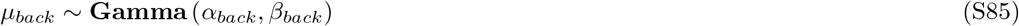

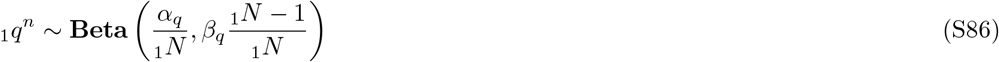

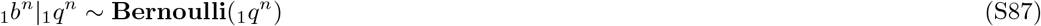

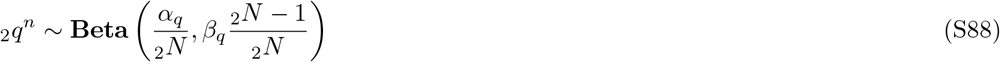

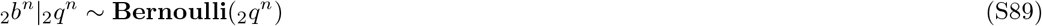

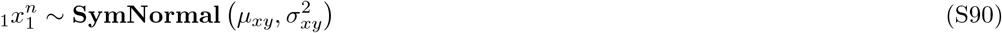

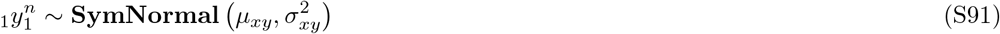

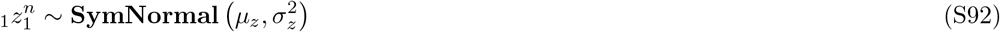

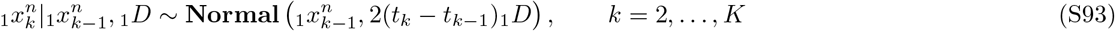

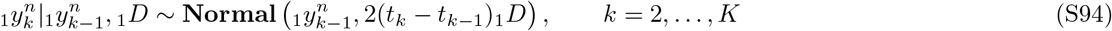

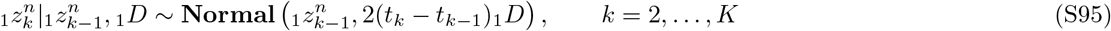

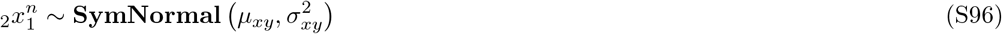

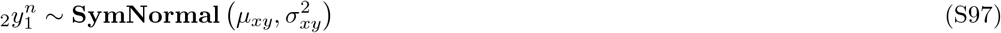

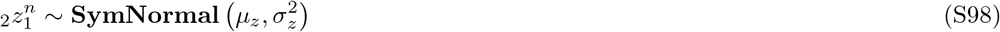

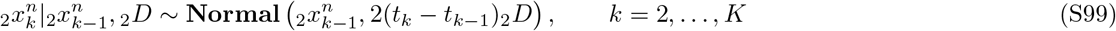

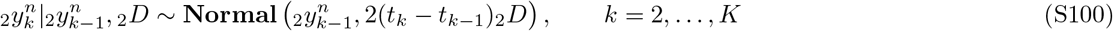

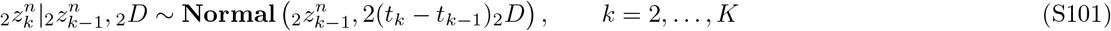

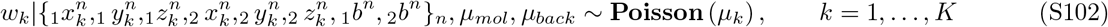

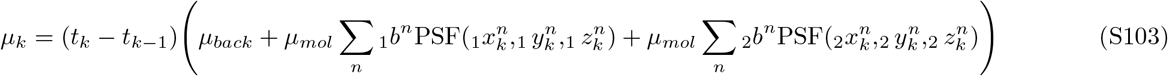

where pre-scripts 1 and 2 are used to distinguish the two species. A graphical summary is show on Fig. S16. We use this formulation for the estimates shown on Figs. S11 and S12.

**FIG. S16.**
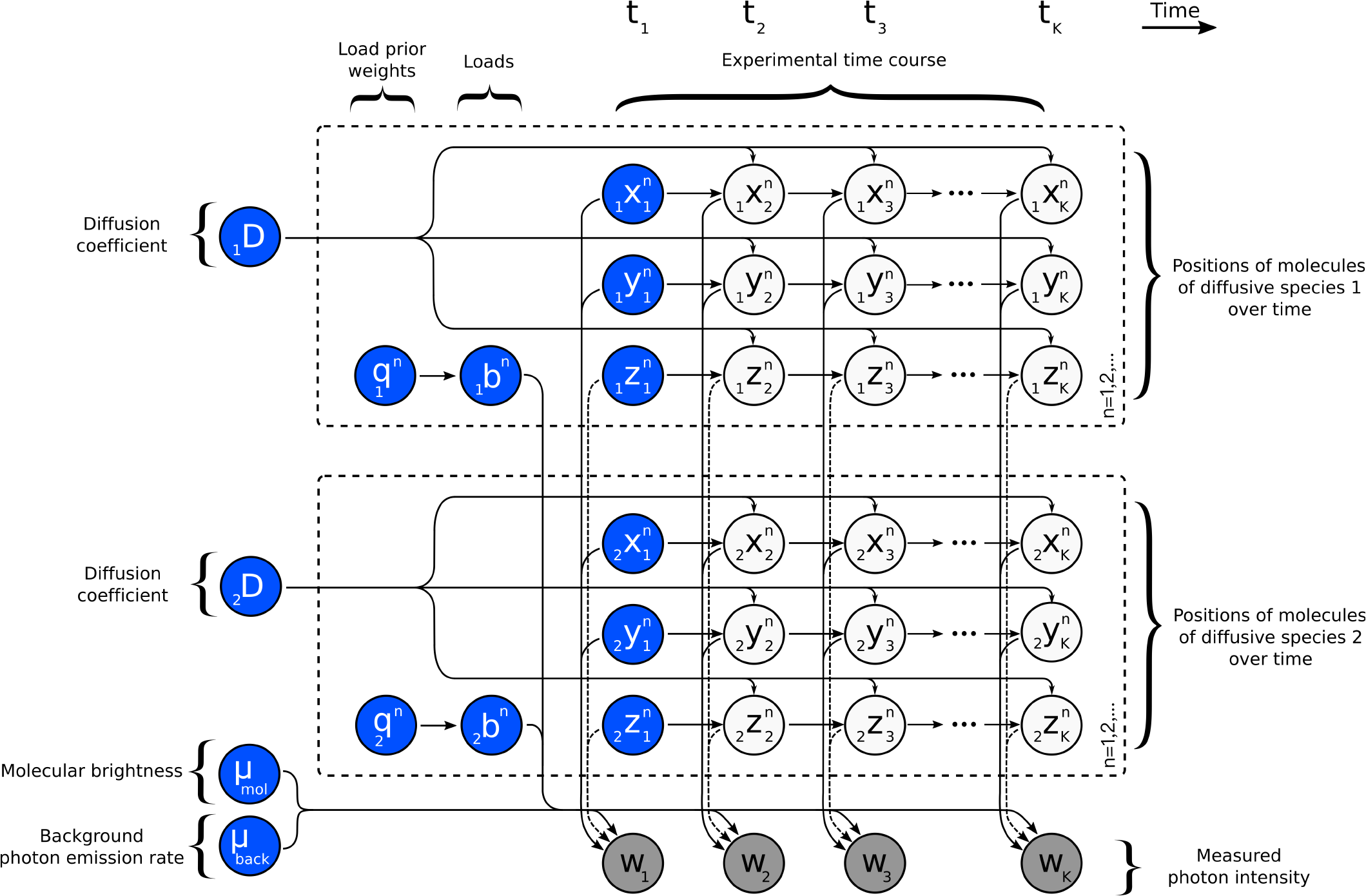
Graphical summary of the framework capturing two independent diffusion coefficients. A multi-species population of model molecules, labeled by *n* = 1, 2, …, evolves during the measurement period which is marked by *k* = 1, 2, …, *K*. Here, 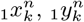 and 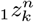 denote the location of molecule *n* at time *t*_*k*_ of species 1, 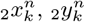 and 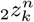 denote the location of molecule *n* at time *t*_*k*_ of species 2, *µ*_*mol*_ and *µ*_*back*_ denote molecular brightness and background photon emission rates. The diffusion coefficient _1_*D* and _2_*D* determine the evolution of the molecule locations of species 1 and 2 which, in turn, determine the instantaneous photon emission rates and ultimately the recorded photon intensities *w*_*k*_. Load variables _1_*b*^*n*^ and _2_*b*^*n*^, with prior weights _1_*q*^*n*^ and _2_*q*^*n*^, respectively, are introduced to model molecule populations of the two species of *a priori* unknown sizes. Following common machine learning convention, the measurements *w*_*k*_ are dark shaded. Additionally, model variables requiring prior probability distributions are highlighted in blue.

## S6. Summary of notation, abbreviations, parameters and other options

**TABLE S3.** Summary of notation.

| Description | Variable | Units |
| --- | --- | --- |
| Diffusion coefficient | $D$ | $\mu\text{m}^2/\text{s}$ |
| $\alpha$ parameter of the diffusion coefficient prior | $\alpha_D$ | - |
| $\beta$ parameter of the diffusion coefficient prior | $\beta_D$ | $\mu\text{m}^2/\text{s}$ |
| Total time trace duration | $T_{total}$ | s |
| Molecular brightness at the center of the confocal volume | $\mu_{mol}$ | photons/s |
| $\alpha$ parameter of the molecular brightness's prior | $\alpha_{mol}$ | - |
| $\beta$ parameter of the molecular brightness's prior | $\beta_{mol}$ | photons/s |
| Proposal parameter of the molecule photon emission rate | $\alpha_{mol}^{prop}$ | - |
| Emission rate of molecule $n$ at time $t_k$ | $\mu_k^n$ | photons/s |
| Combined photon emission rates of all molecules at time $t_k$ | $\mu_k$ | photons/s |
| Background photon emission rate | $\mu_{back}$ | photons/s |
| $\alpha$ parameter of the background photon emission rate's prior | $\alpha_{back}$ | - |
| $\beta$ parameter of the background photon emission rate's prior | $\beta_{back}$ | photons/s |
| Proposal parameter of the background photon emission rate | $\alpha_{back}^{prop}$ | - |
| Minor semi-axis of confocal PSF (focal plane) | $\omega_{xy}$ | $\mu\text{m}$ |
| Major semi-axis of confocal 3DG PSF (optical axis) | $\omega_z$ | $\mu\text{m}$ |
| Major semi-axis of confocal 2DG-L PSF (optical axis) | $z_R$ | $\mu\text{m}$ |
| Laser wavelength | $\lambda_{exc}$ | $\mu\text{m}$ |
| Numerical aperture | NA | - |
| Solution refractive index | $n_{sol}$ | - |
| Location of sigma points | $x^{sn}$ | $\mu\text{m}$ |
| Location of molecule $n$ at time $t_k$ in $x$-coordinate | $x_k^n$ | $\mu\text{m}$ |
| Location of molecule $n$ at time $t_k$ in $y$-coordinate | $y_k^n$ | $\mu\text{m}$ |
| Location of molecule $n$ at time $t_k$ in $z$-coordinate | $z_k^n$ | $\mu\text{m}$ |
| Recorded photon intensity at time $t_k$ | $w_k$ | photons |
| Load variable for molecule $n$ | $b^n$ | - |
| Prior weight for $b_n$ | $q^n$ | - |
| $\alpha$ parameter of prior weight $q^n$ | $\alpha_q$ | - |
| $\beta$ parameter of prior weight $q^n$ | $\beta_q$ | - |
| Upper bound for the number of model molecules | $N$ | - |
| Mean value of initial molecule position's prior in the $xy$-plane | $\mu_{xy}$ | $\mu\text{m}$ |
| Mean value of initial molecule position's prior on the $z$-axis | $\mu_z$ | $\mu\text{m}$ |
| Variance of the initial molecule position's prior in the $xy$-plane | $\sigma_{xy}$ | $\mu\text{m}$ |
| Variance of the initial molecule position's prior on the $z$-axis | $\sigma_z$ | $\mu\text{m}$ |
| Normalized distance of molecule $n$ at time $t_k$ | $d_k^n$ | - |
| Normalized distance for the definition of effective volume | $\ell$ | - |
| Effective volume | $V_\ell$ | $\mu\text{m}^3$ |
| Heaviside function | $H$ | - |
| Periodic boundary in the $xy$-plane (focal plane) | $L_{xy}$ | $\mu\text{m}$ |
| Periodic boundary on the $z$-axis (optical axis) | $L_z$ | $\mu\text{m}$ |
| Convergence threshold | $\epsilon_{thr}$ | $\mu\text{m}^2/\text{s}$ |
| Bin threshold | $\epsilon_{bin}$ | photons |

**TABLE S4.**
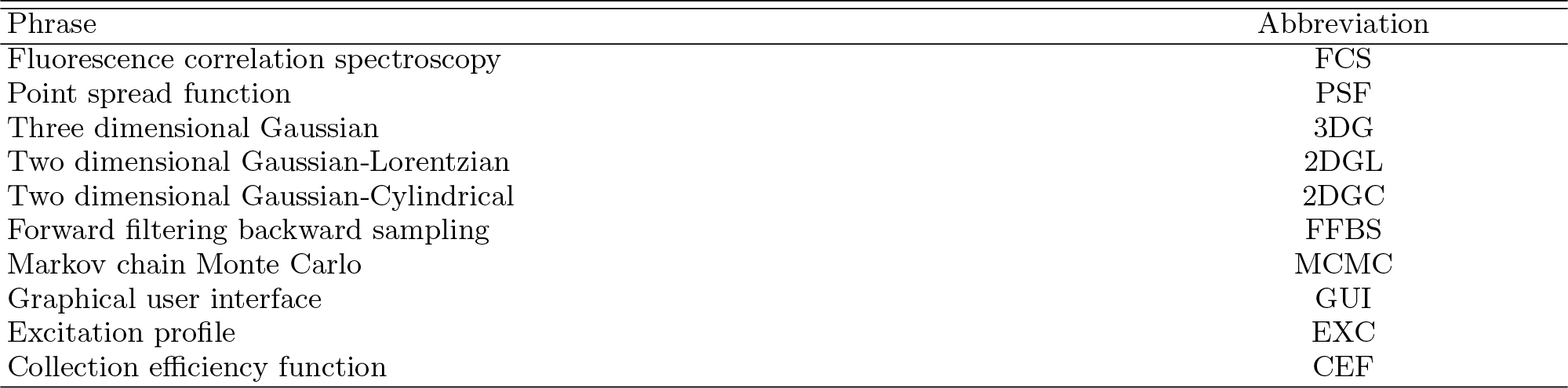
List of abbreviations.

**TABLE S5.**
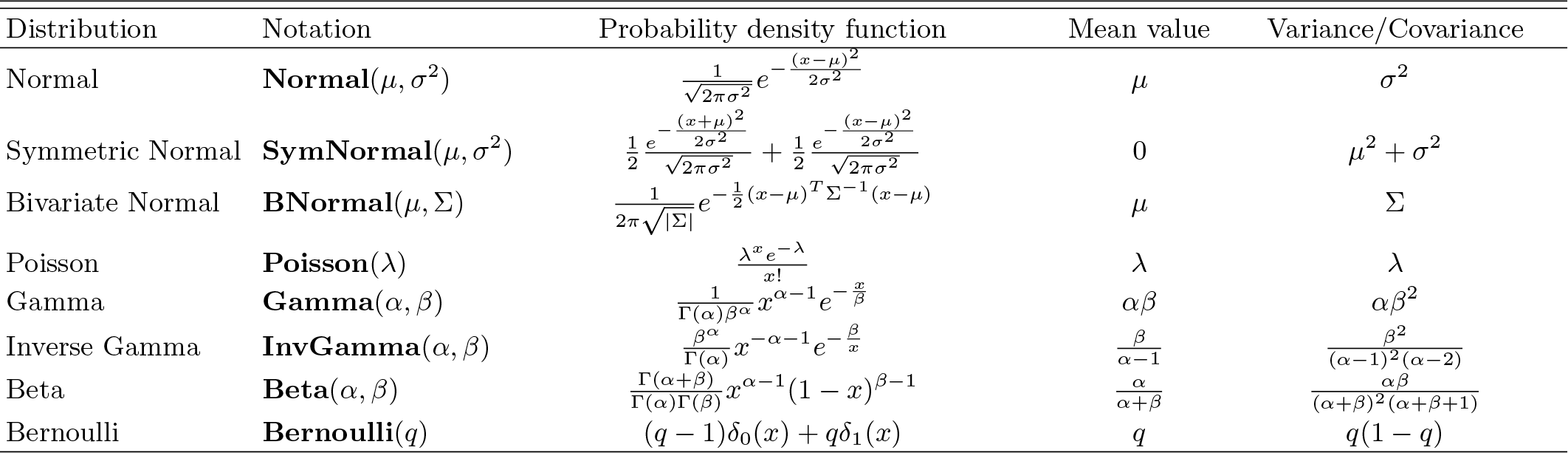
Probability distributions used and their densities. Here, the corresponding random variables are denoted by *x*.

**TABLE S6.** Parameter values used in the generation of the synthetic traces. Choices are listed according to figures.

| | PSF | $L_{xy}$ | $L_z$ | $\omega_{xy}$ | $\omega_{z,zR}$ | $N$ | $D$ | $\mu_{mol}$ | $\mu_{back}$ | $T_{total}$ | $\delta t$ |
| --- | --- | --- | --- | --- | --- | --- | --- | --- | --- | --- | --- |
| Units | | $\mu m$ | $\mu m$ | $\mu m$ | $\mu m$ | - | $\mu m^2/s$ | photons/s | photons/s | s | s |
| Fig. 1(a1) | 3DG | 2 | 3 | 0.30 | 1.50 | 10 | 10 | $5 \times 10^4$ | $10^3$ | 0.1 | $10^{-4}$ |
| Fig. 1(a2) | 3DG | 2 | 3 | 0.30 | 1.50 | 100 | 10 | $5 \times 10^4$ | $10^3$ | 0.1 | $10^{-4}$ |
| Fig. 2(a) | 3DG | 2,2,2,2,4 | 3,3,3,3,7 | 0.30 | 1.50 | $10^2, 10^2, 10^2, 10^2, 10^3$ | $10^{-2}, 10^{-1}, 1, 10, 10^2$ | $5 \times 10^4$ | $10^3$ | 0.1 | $10^{-4}$ |
| Fig. 2(c) | 3DG | 2 | 3 | 0.30 | 1.50 | 150 | 10 | $5 \times 10^4$ | $10^3$ | 10 | $10^{-4}$ |
| Fig. 3(a) | 3DG | 2 | 3 | 0.30 | 1.50 | 150 | 1 | $10^5, 5 \times 10^4, 10^4$ | $10^3$ | 0.1 | $10^{-4}$ |
| Fig. S3 | 3DG | 2 | 3 | 0.30 | 1.50 | 50 | 1 | $5 \times 10^4$ | $10^3$ | 0.1 | $10^{-4}$ |
| Fig. S4(a) | 3DG | 2 | 3 | 0.30 | 1.50 | 50 | 10 | $5 \times 10^4$ | $10^3$ | 100 | $10^{-4}$ |
| Fig. S5(a1) | 3DG | 2 | 3 | 0.30 | 1.50 | 50 | 10 | $5 \times 10^4$ | $10^3$ | 0.1 | $10^{-4}$ |
| Fig. S5(a2) | 2DGL | 2 | 3 | 0.30 | 1.50 | 50 | 10 | $5 \times 10^4$ | $10^3$ | 0.1 | $10^{-4}$ |
| Fig. S5(a3) | 2DGC | 2 | 3 | 0.30 | - | 50 | 10 | $5 \times 10^4$ | $10^3$ | 0.1 | $10^{-4}$ |
| Fig. S11(a) | 3DG | 2 | 3 | 0.30 | 1.50 | 20, 20 | 1,10 | $5 \times 10^4$ | $10^3$ | 1 | $10^{-4}$ |

**TABLE S7.** Parameter values used in the analyses of the traces. Choices are listed according to figures.

| Units | PSF | $\omega_{xy}$<br>$\mu m$ | $\omega_z, z_R$<br>$\mu m$ | $N$<br>- | $\alpha_D$<br>- | $\beta_D$<br>$\mu m^2/s$ | $\alpha_{mol}$<br>- | $\beta_{mol}$<br>pht/s | $\alpha_{mol}^{prop}$<br>- | $\alpha_{back}$<br>- | $\beta_{back}$<br>pht/s | $\alpha_{back}^{prop}$<br>- | $\alpha_q$<br>- | $\beta_q$<br>- | $\mu_{xy}$<br>$\mu m$ | $\mu_z$<br>$\mu m$ | $\sigma_{xy}^2$<br>$\mu m^2$ | $\sigma_z^2$<br>$\mu m^2$ | $\epsilon_{thr}$<br>$\mu m^2/s$ | $\epsilon_{bin}$<br>pht |
| --- | --- | --- | --- | --- | --- | --- | --- | --- | --- | --- | --- | --- | --- | --- | --- | --- | --- | --- | --- | --- |
| Fig. 1(a1) | 3DG | 0.30 | 1.50 | 50 | 1 | 1 | 2 | $10^4$ | $10^3$ | 2 | 500 | $10^3$ | 1 | 1 | 0.2 | 0.2 | 2 | 2 | 0.1 | 4 |
| Fig. 1(a2) | 3DG | 0.30 | 1.50 | 50 | 1 | 1 | 2 | $10^4$ | $10^3$ | 2 | 500 | $10^3$ | 1 | 1 | 0.2 | 0.2 | 2 | 2 | 0.1 | 4 |
| Fig. 2(a) | 3DG | 0.30 | 1.50 | 50 | 1 | 1 | 2 | $10^4$ | $10^3$ | 2 | 500 | $10^3$ | 1 | 1 | 0.2 | 0.2 | 2 | 2 | 0.1 | 4 |
| Fig. 2(b) | 3DG | 0.30 | 1.50 | 50 | 1 | 1 | 2 | $10^4$ | $10^3$ | 2 | 500 | $10^3$ | 1 | 1 | 0.2 | 0.2 | 2 | 2 | 0.1 | 4 |
| Fig. 3 | 3DG | 0.30 | 1.50 | 50 | 1 | 1 | 2 | $10^4$ | $10^3$ | 2 | 500 | $10^3$ | 1 | 1 | 0.2 | 0.2 | 2 | 2 | 0.1 | 4 |
| Fig. 4 | 2DGL | 0.40 | - | 50 | 1 | 1 | 2 | $10^4$ | $10^3$ | 2 | 500 | $10^3$ | 1 | 1 | 0.2 | 0.2 | 2 | 2 | 0.1 | 4 |
| Fig. 5 | 3DG | 0.27 | 4.51 | 50 | 1 | 1 | 2 | $10^4$ | $10^3$ | 2 | 500 | $10^3$ | 1 | 1 | 0.2 | 0.2 | 2 | 2 | 0.1 | 4 |
| Fig. 6 | 3DG | 0.27 | 4.51 | 50 | 1 | 1 | 2 | $10^4$ | $10^3$ | 2 | 500 | $10^3$ | 1 | 1 | 0.2 | 0.2 | 2 | 2 | 0.1 | 4 |
| Fig. 7(a1) | 3DG | 0.27 | 4.51 | 50 | 1 | 1 | 2 | $10^4$ | $10^3$ | 2 | 500 | $10^3$ | 1 | 1 | 0.2 | 0.2 | 2 | 2 | 0.1 | 4 |
| Fig. 7(a2) | 3DG | 0.27 | 4.51 | 50 | 1 | 1 | 2 | $10^4$ | $10^3$ | 2 | 500 | $10^3$ | 1 | 1 | 0.2 | 0.2 | 2 | 2 | 0.1 | 4 |
| Fig. 7(a3) | 3DG | 0.27 | 4.51 | 50 | 1 | 1 | 2 | $10^4$ | $10^3$ | 2 | 500 | $10^3$ | 1 | 1 | 0.2 | 0.2 | 2 | 2 | 0.1 | 4 |
| Fig. S3 | 3DG | 0.30 | 1.50 | 50 | 1 | 1 | 2 | $10^4$ | $10^3$ | 2 | 500 | $10^3$ | 1 | 1 | 0.2 | 0.2 | 2 | 2 | 0.1 | 4 |
| Fig. S4 | 3DG | 0.23 | 0.55 | 50 | 1 | 1 | 2 | $10^4$ | $10^3$ | 2 | 500 | $10^3$ | 1 | 1 | 0.2 | 0.2 | 2 | 2 | 0.1 | 4 |
| Fig. S5(b1) | 3DG | 0.27 | 4.51 | 50 | 1 | 1 | 2 | $10^4$ | $10^3$ | 2 | 500 | $10^3$ | 1 | 1 | 0.2 | 0.2 | 2 | 2 | 0.1 | 4 |
| Fig. S5(b2) | 2DGL | 0.27 | 4.51 | 50 | 1 | 1 | 2 | $10^4$ | $10^3$ | 2 | 500 | $10^3$ | 1 | 1 | 0.2 | 0.2 | 2 | 2 | 0.1 | 4 |
| Fig. S5(b3) | 2DGC | 0.27 | - | 50 | 1 | 1 | 2 | $10^4$ | $10^3$ | 2 | 500 | $10^3$ | 1 | 1 | 0.2 | 0.2 | 2 | 2 | 0.1 | 4 |
| Fig. S8 | 3DG | 0.27 | 4.51 | 50 | 1 | 1 | 2 | $10^4$ | $10^3$ | 2 | 500 | $10^3$ | 1 | 1 | 0.2 | 0.2 | 2 | 2 | 0.1 | 4 |
| Fig. S9 | 3DG | 0.27 | 4.51 | 50 | 1 | 1 | 2 | $10^4$ | $10^3$ | 2 | 500 | $10^3$ | 1 | 1 | 0.2 | 0.2 | 2 | 2 | 0.1 | 4 |
| Fig. S10 | 2DGL | 0.42 | - | 50 | 1 | 1 | 2 | $10^4$ | $10^3$ | 2 | 500 | $10^3$ | 1 | 1 | 0.2 | 0.2 | 2 | 2 | 0.1 | 4 |
| Fig. S11 | 3DG | 0.3 | 1.5 | 50 | 1 | 1 | 2 | $10^4$ | $10^3$ | 2 | 500 | $10^3$ | 1 | 1 | 0.2 | 0.2 | 2 | 2 | 0.1 | 4 |

