## Supplementary material for "An Alternative Framework for Fluorescence Correlation Spectroscopy": Source Code: ASWFCS.m

```
function varargout = ASWFCS(varargin)
% ASWFCS MATLAB code for ASWFCS.fig
%      ASWFCS, by itself, creates a new ASWFCS or raises the existing
%      singleton*.
%
%      H = ASWFCS returns the handle to a new ASWFCS or the handle to
%      the existing singleton*.
%
%      ASWFCS('CALLBACK',hObject,eventData,handles,...) calls the local
%      function named CALLBACK in ASWFCS.M with the given input arguments.
%
%      ASWFCS('Property','Value',...) creates a new ASWFCS or raises the
%      existing singleton*.  Starting from the left, property value pairs are
%      applied to the GUI before ASWFCS_OpeningFcn gets called.  An
%      unrecognized property name or invalid value makes property application
%      stop.  All inputs are passed to ASWFCS_OpeningFcn via varargin.
%
%      *See GUI Options on GUIDE's Tools menu.  Choose "GUI allows only one
%      instance to run (singleton)".
%
% See also: GUIDE, GUIDATA, GUIHANDLES

% Edit the above text to modify the response to help ASWFCS

% Last Modified by GUIDE v2.5 29-Mar-2019 15:04:39

% Begin initialization code - DO NOT EDIT
gui_Singleton = 1;
gui_State = struct('gui_Name',       mfilename, ...
                   'gui_Singleton',  gui_Singleton, ...
                   'gui_OpeningFcn', @ASWFCS_OpeningFcn, ...
                   'gui_OutputFcn',  @ASWFCS_OutputFcn, ...
                   'gui_LayoutFcn',  [] , ...
                   'gui_Callback',   []);
if nargin && ischar(varargin{1})
    gui_State.gui_Callback = str2func(varargin{1});
end

if nargout
    [varargout{1:nargout}] = gui_mainfcn(gui_State, varargin{:});
else
    gui_mainfcn(gui_State, varargin{:});
end
% End initialization code - DO NOT EDIT


% --- Executes just before ASWFCS is made visible.
function ASWFCS_OpeningFcn(hObject, eventdata, handles, varargin)
% This function has no output args, see OutputFcn.
% hObject    handle to figure
% eventdata  reserved - to be defined in a future version of MATLAB
% handles    structure with handles and user data (see GUIDATA)
% varargin   command line arguments to ASWFCS (see VARARGIN)

% Choose default command line output for ASWFCS
handles.output = hObject;

% Update handles structure
guidata(hObject, handles);

 set(handles.text33, 'style', 'text', 'string', ['Diffusion coefficient (',char(hex2dec(strsplit('03BC'))),'m',char(hex2dec(strsplit('00B2'))),'/s)'])
 set(handles.D_beta, 'style', 'text', 'string', [char(hex2dec(strsplit('03B2'))),'   parameter of diffusion coefficient (',char(hex2dec(strsplit('03BC'))),'m',char(hex2dec(strsplit('00B2'))),'/s)'])

 set(handles.text27, 'style', 'text', 'string', ['Initial position',char(hex2dec(strsplit('0027'))),'s variance in xy plane (',char(hex2dec(strsplit('03BC'))),'m',char(hex2dec(strsplit('00B2'))),' )'])
 set(handles.text46, 'style', 'text', 'string', ['Initial position',char(hex2dec(strsplit('0027'))),'s variance in z axis (',char(hex2dec(strsplit('03BC'))),'m',char(hex2dec(strsplit('00B2'))),' )'])
 set(handles.text57, 'style', 'text', 'string', ['Convergence threshold (',char(hex2dec(strsplit('03BC'))),'m',char(hex2dec(strsplit('00B2'))),'/s)'])
 

addpath('sampler_Fun')

% UIWAIT makes ASWFCS wait for user response (see UIRESUME)
% uiwait(handles.figure1);


% --- Outputs from this function are returned to the command line.
function varargout = ASWFCS_OutputFcn(hObject, eventdata, handles) 
% varargout  cell array for returning output args (see VARARGOUT);
% hObject    handle to figure
% eventdata  reserved - to be defined in a future version of MATLAB
% handles    structure with handles and user data (see GUIDATA)

% Get default command line output from handles structure
varargout{1} = handles.output;


% --- Executes on button press in Start.
function Start_Callback(hObject, eventdata, handles)
% hObject    handle to Start (see GCBO)
% eventdata  reserved - to be defined in a future version of MATLAB
% handles    structure with handles and user data (see GUIDATA)


% --- Executes on button press in Stop.
function Stop_Callback(hObject, eventdata, handles)
% hObject    handle to Stop (see GCBO)
% eventdata  reserved - to be defined in a future version of MATLAB
% handles    structure with handles and user data (see GUIDATA)


% --- Executes on button press in partial_signal.
function partial_signal_Callback(hObject, eventdata, handles)
% hObject    handle to partial_signal (see GCBO)
% eventdata  reserved - to be defined in a future version of MATLAB
% handles    structure with handles and user data (see GUIDATA)


function Min_signal_Callback(hObject, eventdata, handles)
% hObject    handle to Min_signal (see GCBO)
% eventdata  reserved - to be defined in a future version of MATLAB
% handles    structure with handles and user data (see GUIDATA)

% Hints: get(hObject,'String') returns contents of Min_signal as text
%        str2double(get(hObject,'String')) returns contents of Min_signal as a double


% --- Executes during object creation, after setting all properties.
function Min_signal_CreateFcn(hObject, eventdata, handles)
% hObject    handle to Min_signal (see GCBO)
% eventdata  reserved - to be defined in a future version of MATLAB
% handles    empty - handles not created until after all CreateFcns called

% Hint: edit controls usually have a white background on Windows.
%       See ISPC and COMPUTER.
if ispc && isequal(get(hObject,'BackgroundColor'), get(0,'defaultUicontrolBackgroundColor'))
    set(hObject,'BackgroundColor','white');
end


function Max_signal_Callback(hObject, eventdata, handles)
% hObject    handle to Max_signal (see GCBO)
% eventdata  reserved - to be defined in a future version of MATLAB
% handles    structure with handles and user data (see GUIDATA)

% Hints: get(hObject,'String') returns contents of Max_signal as text
%        str2double(get(hObject,'String')) returns contents of Max_signal as a double


% --- Executes during object creation, after setting all properties.
function Max_signal_CreateFcn(hObject, eventdata, handles)
% hObject    handle to Max_signal (see GCBO)
% eventdata  reserved - to be defined in a future version of MATLAB
% handles    empty - handles not created until after all CreateFcns called

% Hint: edit controls usually have a white background on Windows.
%       See ISPC and COMPUTER.
if ispc && isequal(get(hObject,'BackgroundColor'), get(0,'defaultUicontrolBackgroundColor'))
    set(hObject,'BackgroundColor','white');
end


% --- Executes on button press in Results_show.
function Results_show_Callback(hObject, eventdata, handles)
% hObject    handle to Results_show (see GCBO)
% eventdata  reserved - to be defined in a future version of MATLAB
% handles    structure with handles and user data (see GUIDATA)


function Refractive_index_Callback(hObject, eventdata, handles)
% hObject    handle to Refractive_index (see GCBO)
% eventdata  reserved - to be defined in a future version of MATLAB
% handles    structure with handles and user data (see GUIDATA)

% Hints: get(hObject,'String') returns contents of Refractive_index as text
%        str2double(get(hObject,'String')) returns contents of Refractive_index as a double


% --- Executes during object creation, after setting all properties.
function Refractive_index_CreateFcn(hObject, eventdata, handles)
% hObject    handle to Refractive_index (see GCBO)
% eventdata  reserved - to be defined in a future version of MATLAB
% handles    empty - handles not created until after all CreateFcns called

% Hint: edit controls usually have a white background on Windows.
%       See ISPC and COMPUTER.
if ispc && isequal(get(hObject,'BackgroundColor'), get(0,'defaultUicontrolBackgroundColor'))
    set(hObject,'BackgroundColor','white');
end


function Laser_Wavelength_Callback(hObject, eventdata, handles)
% hObject    handle to Laser_Wavelength (see GCBO)
% eventdata  reserved - to be defined in a future version of MATLAB
% handles    structure with handles and user data (see GUIDATA)

% Hints: get(hObject,'String') returns contents of Laser_Wavelength as text
%        str2double(get(hObject,'String')) returns contents of Laser_Wavelength as a double


% --- Executes during object creation, after setting all properties.
function Laser_Wavelength_CreateFcn(hObject, eventdata, handles)
% hObject    handle to Laser_Wavelength (see GCBO)
% eventdata  reserved - to be defined in a future version of MATLAB
% handles    empty - handles not created until after all CreateFcns called

% Hint: edit controls usually have a white background on Windows.
%       See ISPC and COMPUTER.
if ispc && isequal(get(hObject,'BackgroundColor'), get(0,'defaultUicontrolBackgroundColor'))
    set(hObject,'BackgroundColor','white');
end


function Objective_magnification_Callback(hObject, eventdata, handles)
% hObject    handle to Objective_magnification (see GCBO)
% eventdata  reserved - to be defined in a future version of MATLAB
% handles    structure with handles and user data (see GUIDATA)

% Hints: get(hObject,'String') returns contents of Objective_magnification as text
%        str2double(get(hObject,'String')) returns contents of Objective_magnification as a double


% --- Executes during object creation, after setting all properties.
function Objective_magnification_CreateFcn(hObject, eventdata, handles)
% hObject    handle to Objective_magnification (see GCBO)
% eventdata  reserved - to be defined in a future version of MATLAB
% handles    empty - handles not created until after all CreateFcns called

% Hint: edit controls usually have a white background on Windows.
%       See ISPC and COMPUTER.
if ispc && isequal(get(hObject,'BackgroundColor'), get(0,'defaultUicontrolBackgroundColor'))
    set(hObject,'BackgroundColor','white');
end


function wxy_Callback(hObject, eventdata, handles)
% hObject    handle to wxy (see GCBO)
% eventdata  reserved - to be defined in a future version of MATLAB
% handles    structure with handles and user data (see GUIDATA)

% Hints: get(hObject,'String') returns contents of wxy as text
%        str2double(get(hObject,'String')) returns contents of wxy as a double


% --- Executes during object creation, after setting all properties.
function wxy_CreateFcn(hObject, eventdata, handles)
% hObject    handle to wxy (see GCBO)
% eventdata  reserved - to be defined in a future version of MATLAB
% handles    empty - handles not created until after all CreateFcns called

% Hint: edit controls usually have a white background on Windows.
%       See ISPC and COMPUTER.
if ispc && isequal(get(hObject,'BackgroundColor'), get(0,'defaultUicontrolBackgroundColor'))
    set(hObject,'BackgroundColor','white');
end


function wz_Callback(hObject, eventdata, handles)
% hObject    handle to wz (see GCBO)
% eventdata  reserved - to be defined in a future version of MATLAB
% handles    structure with handles and user data (see GUIDATA)

% Hints: get(hObject,'String') returns contents of wz as text
%        str2double(get(hObject,'String')) returns contents of wz as a double


% --- Executes during object creation, after setting all properties.
function wz_CreateFcn(hObject, eventdata, handles)
% hObject    handle to wz (see GCBO)
% eventdata  reserved - to be defined in a future version of MATLAB
% handles    empty - handles not created until after all CreateFcns called

% Hint: edit controls usually have a white background on Windows.
%       See ISPC and COMPUTER.
if ispc && isequal(get(hObject,'BackgroundColor'), get(0,'defaultUicontrolBackgroundColor'))
    set(hObject,'BackgroundColor','white');
end


function mu_back_Callback(hObject, eventdata, handles)
% hObject    handle to mu_back (see GCBO)
% eventdata  reserved - to be defined in a future version of MATLAB
% handles    structure with handles and user data (see GUIDATA)

% Hints: get(hObject,'String') returns contents of mu_back as text
%        str2double(get(hObject,'String')) returns contents of mu_back as a double


% --- Executes during object creation, after setting all properties.
function mu_back_CreateFcn(hObject, eventdata, handles)
% hObject    handle to mu_back (see GCBO)
% eventdata  reserved - to be defined in a future version of MATLAB
% handles    empty - handles not created until after all CreateFcns called

% Hint: edit controls usually have a white background on Windows.
%       See ISPC and COMPUTER.
if ispc && isequal(get(hObject,'BackgroundColor'), get(0,'defaultUicontrolBackgroundColor'))
    set(hObject,'BackgroundColor','white');
end


function Delta_t_Callback(hObject, eventdata, handles)
% hObject    handle to Delta_t (see GCBO)
% eventdata  reserved - to be defined in a future version of MATLAB
% handles    structure with handles and user data (see GUIDATA)

% Hints: get(hObject,'String') returns contents of Delta_t as text
%        str2double(get(hObject,'String')) returns contents of Delta_t as a double


% --- Executes during object creation, after setting all properties.
function Delta_t_CreateFcn(hObject, eventdata, handles)
% hObject    handle to Delta_t (see GCBO)
% eventdata  reserved - to be defined in a future version of MATLAB
% handles    empty - handles not created until after all CreateFcns called

% Hint: edit controls usually have a white background on Windows.
%       See ISPC and COMPUTER.
if ispc && isequal(get(hObject,'BackgroundColor'), get(0,'defaultUicontrolBackgroundColor'))
    set(hObject,'BackgroundColor','white');
end


function concen_radious_xy_Callback(hObject, eventdata, handles)
% hObject    handle to concen_radious (see GCBO)
% eventdata  reserved - to be defined in a future version of MATLAB
% handles    structure with handles and user data (see GUIDATA)

% Hints: get(hObject,'String') returns contents of concen_radious as text
%        str2double(get(hObject,'String')) returns contents of concen_radious as a double


% --- Executes during object creation, after setting all properties.
function concen_radious_xy_CreateFcn(hObject, eventdata, handles)
% hObject    handle to concen_radious (see GCBO)
% eventdata  reserved - to be defined in a future version of MATLAB
% handles    empty - handles not created until after all CreateFcns called

% Hint: edit controls usually have a white background on Windows.
%       See ISPC and COMPUTER.
if ispc && isequal(get(hObject,'BackgroundColor'), get(0,'defaultUicontrolBackgroundColor'))
    set(hObject,'BackgroundColor','white');
end


function concen_radious_z_Callback(hObject, eventdata, handles)
% hObject    handle to concen_radious_z (see GCBO)
% eventdata  reserved - to be defined in a future version of MATLAB
% handles    structure with handles and user data (see GUIDATA)

% Hints: get(hObject,'String') returns contents of concen_radious_z as text
%        str2double(get(hObject,'String')) returns contents of concen_radious_z as a double


% --- Executes during object creation, after setting all properties.
function concen_radious_z_CreateFcn(hObject, eventdata, handles)
% hObject    handle to concen_radious_z (see GCBO)
% eventdata  reserved - to be defined in a future version of MATLAB
% handles    empty - handles not created until after all CreateFcns called

% Hint: edit controls usually have a white background on Windows.
%       See ISPC and COMPUTER.
if ispc && isequal(get(hObject,'BackgroundColor'), get(0,'defaultUicontrolBackgroundColor'))
    set(hObject,'BackgroundColor','white');
end


function mu_prior_z_Callback(hObject, eventdata, handles)
% hObject    handle to mu_prior_z (see GCBO)
% eventdata  reserved - to be defined in a future version of MATLAB
% handles    structure with handles and user data (see GUIDATA)

% Hints: get(hObject,'String') returns contents of mu_prior_z as text
%        str2double(get(hObject,'String')) returns contents of mu_prior_z as a double


% --- Executes during object creation, after setting all properties.
function mu_prior_z_CreateFcn(hObject, eventdata, handles)
% hObject    handle to mu_prior_z (see GCBO)
% eventdata  reserved - to be defined in a future version of MATLAB
% handles    empty - handles not created until after all CreateFcns called

% Hint: edit controls usually have a white background on Windows.
%       See ISPC and COMPUTER.
if ispc && isequal(get(hObject,'BackgroundColor'), get(0,'defaultUicontrolBackgroundColor'))
    set(hObject,'BackgroundColor','white');
end


function var_prior_xy_Callback(hObject, eventdata, handles)
% hObject    handle to var_prior_xy (see GCBO)
% eventdata  reserved - to be defined in a future version of MATLAB
% handles    structure with handles and user data (see GUIDATA)

% Hints: get(hObject,'String') returns contents of var_prior_xy as text
%        str2double(get(hObject,'String')) returns contents of var_prior_xy as a double


% --- Executes during object creation, after setting all properties.
function var_prior_xy_CreateFcn(hObject, eventdata, handles)
% hObject    handle to var_prior_xy (see GCBO)
% eventdata  reserved - to be defined in a future version of MATLAB
% handles    empty - handles not created until after all CreateFcns called

% Hint: edit controls usually have a white background on Windows.
%       See ISPC and COMPUTER.
if ispc && isequal(get(hObject,'BackgroundColor'), get(0,'defaultUicontrolBackgroundColor'))
    set(hObject,'BackgroundColor','white');
end


function mu_prior_xy_Callback(hObject, eventdata, handles)
% hObject    handle to mu_prior_xy (see GCBO)
% eventdata  reserved - to be defined in a future version of MATLAB
% handles    structure with handles and user data (see GUIDATA)

% Hints: get(hObject,'String') returns contents of mu_prior_xy as text
%        str2double(get(hObject,'String')) returns contents of mu_prior_xy as a double


% --- Executes during object creation, after setting all properties.
function mu_prior_xy_CreateFcn(hObject, eventdata, handles)
% hObject    handle to mu_prior_xy (see GCBO)
% eventdata  reserved - to be defined in a future version of MATLAB
% handles    empty - handles not created until after all CreateFcns called

% Hint: edit controls usually have a white background on Windows.
%       See ISPC and COMPUTER.
if ispc && isequal(get(hObject,'BackgroundColor'), get(0,'defaultUicontrolBackgroundColor'))
    set(hObject,'BackgroundColor','white');
end


function alpha_load_Callback(hObject, eventdata, handles)
% hObject    handle to alpha_load (see GCBO)
% eventdata  reserved - to be defined in a future version of MATLAB
% handles    structure with handles and user data (see GUIDATA)

% Hints: get(hObject,'String') returns contents of alpha_load as text
%        str2double(get(hObject,'String')) returns contents of alpha_load as a double


% --- Executes during object creation, after setting all properties.
function alpha_load_CreateFcn(hObject, eventdata, handles)
% hObject    handle to alpha_load (see GCBO)
% eventdata  reserved - to be defined in a future version of MATLAB
% handles    empty - handles not created until after all CreateFcns called

% Hint: edit controls usually have a white background on Windows.
%       See ISPC and COMPUTER.
if ispc && isequal(get(hObject,'BackgroundColor'), get(0,'defaultUicontrolBackgroundColor'))
    set(hObject,'BackgroundColor','white');
end


function beta_load_Callback(hObject, eventdata, handles)
% hObject    handle to beta_load (see GCBO)
% eventdata  reserved - to be defined in a future version of MATLAB
% handles    structure with handles and user data (see GUIDATA)

% Hints: get(hObject,'String') returns contents of beta_load as text
%        str2double(get(hObject,'String')) returns contents of beta_load as a double


% --- Executes during object creation, after setting all properties.
function beta_load_CreateFcn(hObject, eventdata, handles)
% hObject    handle to beta_load (see GCBO)
% eventdata  reserved - to be defined in a future version of MATLAB
% handles    empty - handles not created until after all CreateFcns called

% Hint: edit controls usually have a white background on Windows.
%       See ISPC and COMPUTER.
if ispc && isequal(get(hObject,'BackgroundColor'), get(0,'defaultUicontrolBackgroundColor'))
    set(hObject,'BackgroundColor','white');
end


function Number_particles_Callback(hObject, eventdata, handles)
% hObject    handle to Number_particles (see GCBO)
% eventdata  reserved - to be defined in a future version of MATLAB
% handles    structure with handles and user data (see GUIDATA)

% Hints: get(hObject,'String') returns contents of Number_particles as text
%        str2double(get(hObject,'String')) returns contents of Number_particles as a double


% --- Executes during object creation, after setting all properties.
function Number_particles_CreateFcn(hObject, eventdata, handles)
% hObject    handle to Number_particles (see GCBO)
% eventdata  reserved - to be defined in a future version of MATLAB
% handles    empty - handles not created until after all CreateFcns called

% Hint: edit controls usually have a white background on Windows.
%       See ISPC and COMPUTER.
if ispc && isequal(get(hObject,'BackgroundColor'), get(0,'defaultUicontrolBackgroundColor'))
    set(hObject,'BackgroundColor','white');
end


function mu_init_Callback(hObject, eventdata, handles)
% hObject    handle to mu_init (see GCBO)
% eventdata  reserved - to be defined in a future version of MATLAB
% handles    structure with handles and user data (see GUIDATA)

% Hints: get(hObject,'String') returns contents of mu_init as text
%        str2double(get(hObject,'String')) returns contents of mu_init as a double


% --- Executes during object creation, after setting all properties.
function mu_init_CreateFcn(hObject, eventdata, handles)
% hObject    handle to mu_init (see GCBO)
% eventdata  reserved - to be defined in a future version of MATLAB
% handles    empty - handles not created until after all CreateFcns called

% Hint: edit controls usually have a white background on Windows.
%       See ISPC and COMPUTER.
if ispc && isequal(get(hObject,'BackgroundColor'), get(0,'defaultUicontrolBackgroundColor'))
    set(hObject,'BackgroundColor','white');
end


function mu_alpha_Callback(hObject, eventdata, handles)
% hObject    handle to mu_alpha (see GCBO)
% eventdata  reserved - to be defined in a future version of MATLAB
% handles    structure with handles and user data (see GUIDATA)

% Hints: get(hObject,'String') returns contents of mu_alpha as text
%        str2double(get(hObject,'String')) returns contents of mu_alpha as a double


% --- Executes during object creation, after setting all properties.
function mu_alpha_CreateFcn(hObject, eventdata, handles)
% hObject    handle to mu_alpha (see GCBO)
% eventdata  reserved - to be defined in a future version of MATLAB
% handles    empty - handles not created until after all CreateFcns called

% Hint: edit controls usually have a white background on Windows.
%       See ISPC and COMPUTER.
if ispc && isequal(get(hObject,'BackgroundColor'), get(0,'defaultUicontrolBackgroundColor'))
    set(hObject,'BackgroundColor','white');
end


function mu_beta_Callback(hObject, eventdata, handles)
% hObject    handle to mu_beta (see GCBO)
% eventdata  reserved - to be defined in a future version of MATLAB
% handles    structure with handles and user data (see GUIDATA)

% Hints: get(hObject,'String') returns contents of mu_beta as text
%        str2double(get(hObject,'String')) returns contents of mu_beta as a double


% --- Executes during object creation, after setting all properties.
function mu_beta_CreateFcn(hObject, eventdata, handles)
% hObject    handle to mu_beta (see GCBO)
% eventdata  reserved - to be defined in a future version of MATLAB
% handles    empty - handles not created until after all CreateFcns called

% Hint: edit controls usually have a white background on Windows.
%       See ISPC and COMPUTER.
if ispc && isequal(get(hObject,'BackgroundColor'), get(0,'defaultUicontrolBackgroundColor'))
    set(hObject,'BackgroundColor','white');
end


function D_init_Callback(hObject, eventdata, handles)
% hObject    handle to D_init (see GCBO)
% eventdata  reserved - to be defined in a future version of MATLAB
% handles    structure with handles and user data (see GUIDATA)

% Hints: get(hObject,'String') returns contents of D_init as text
%        str2double(get(hObject,'String')) returns contents of D_init as a double


% --- Executes during object creation, after setting all properties.
function D_init_CreateFcn(hObject, eventdata, handles)
% hObject    handle to D_init (see GCBO)
% eventdata  reserved - to be defined in a future version of MATLAB
% handles    empty - handles not created until after all CreateFcns called

% Hint: edit controls usually have a white background on Windows.
%       See ISPC and COMPUTER.
if ispc && isequal(get(hObject,'BackgroundColor'), get(0,'defaultUicontrolBackgroundColor'))
    set(hObject,'BackgroundColor','white');
end


function D_alpha_prior_Callback(hObject, eventdata, handles)
% hObject    handle to D_alpha_prior (see GCBO)
% eventdata  reserved - to be defined in a future version of MATLAB
% handles    structure with handles and user data (see GUIDATA)

% Hints: get(hObject,'String') returns contents of D_alpha_prior as text
%        str2double(get(hObject,'String')) returns contents of D_alpha_prior as a double


% --- Executes during object creation, after setting all properties.
function D_alpha_prior_CreateFcn(hObject, eventdata, handles)
% hObject    handle to D_alpha_prior (see GCBO)
% eventdata  reserved - to be defined in a future version of MATLAB
% handles    empty - handles not created until after all CreateFcns called

% Hint: edit controls usually have a white background on Windows.
%       See ISPC and COMPUTER.
if ispc && isequal(get(hObject,'BackgroundColor'), get(0,'defaultUicontrolBackgroundColor'))
    set(hObject,'BackgroundColor','white');
end


function D_beta_prior_Callback(hObject, eventdata, handles)
% hObject    handle to D_beta_prior (see GCBO)
% eventdata  reserved - to be defined in a future version of MATLAB
% handles    structure with handles and user data (see GUIDATA)

% Hints: get(hObject,'String') returns contents of D_beta_prior as text
%        str2double(get(hObject,'String')) returns contents of D_beta_prior as a double


% --- Executes during object creation, after setting all properties.
function D_beta_prior_CreateFcn(hObject, eventdata, handles)
% hObject    handle to D_beta_prior (see GCBO)
% eventdata  reserved - to be defined in a future version of MATLAB
% handles    empty - handles not created until after all CreateFcns called

% Hint: edit controls usually have a white background on Windows.
%       See ISPC and COMPUTER.
if ispc && isequal(get(hObject,'BackgroundColor'), get(0,'defaultUicontrolBackgroundColor'))
    set(hObject,'BackgroundColor','white');
end


function Diff_synth_Callback(hObject, eventdata, handles)
% hObject    handle to Diff_synth (see GCBO)
% eventdata  reserved - to be defined in a future version of MATLAB
% handles    structure with handles and user data (see GUIDATA)

% Hints: get(hObject,'String') returns contents of Diff_synth as text
%        str2double(get(hObject,'String')) returns contents of Diff_synth as a double


% --- Executes during object creation, after setting all properties.
function Diff_synth_CreateFcn(hObject, eventdata, handles)
% hObject    handle to Diff_synth (see GCBO)
% eventdata  reserved - to be defined in a future version of MATLAB
% handles    empty - handles not created until after all CreateFcns called

% Hint: edit controls usually have a white background on Windows.
%       See ISPC and COMPUTER.
if ispc && isequal(get(hObject,'BackgroundColor'), get(0,'defaultUicontrolBackgroundColor'))
    set(hObject,'BackgroundColor','white');
end


function mu_synth_Callback(hObject, eventdata, handles)
% hObject    handle to mu_synth (see GCBO)
% eventdata  reserved - to be defined in a future version of MATLAB
% handles    structure with handles and user data (see GUIDATA)

% Hints: get(hObject,'String') returns contents of mu_synth as text
%        str2double(get(hObject,'String')) returns contents of mu_synth as a double


% --- Executes during object creation, after setting all properties.
function mu_synth_CreateFcn(hObject, eventdata, handles)
% hObject    handle to mu_synth (see GCBO)
% eventdata  reserved - to be defined in a future version of MATLAB
% handles    empty - handles not created until after all CreateFcns called

% Hint: edit controls usually have a white background on Windows.
%       See ISPC and COMPUTER.
if ispc && isequal(get(hObject,'BackgroundColor'), get(0,'defaultUicontrolBackgroundColor'))
    set(hObject,'BackgroundColor','white');
end


function Num_synth_Callback(hObject, eventdata, handles)
% hObject    handle to Num_synth (see GCBO)
% eventdata  reserved - to be defined in a future version of MATLAB
% handles    structure with handles and user data (see GUIDATA)

% Hints: get(hObject,'String') returns contents of Num_synth as text
%        str2double(get(hObject,'String')) returns contents of Num_synth as a double


% --- Executes during object creation, after setting all properties.
function Num_synth_CreateFcn(hObject, eventdata, handles)
% hObject    handle to Num_synth (see GCBO)
% eventdata  reserved - to be defined in a future version of MATLAB
% handles    empty - handles not created until after all CreateFcns called

% Hint: edit controls usually have a white background on Windows.
%       See ISPC and COMPUTER.
if ispc && isequal(get(hObject,'BackgroundColor'), get(0,'defaultUicontrolBackgroundColor'))
    set(hObject,'BackgroundColor','white');
end


function edit35_Callback(hObject, eventdata, handles)
% hObject    handle to edit35 (see GCBO)
% eventdata  reserved - to be defined in a future version of MATLAB
% handles    structure with handles and user data (see GUIDATA)

% Hints: get(hObject,'String') returns contents of edit35 as text
%        str2double(get(hObject,'String')) returns contents of edit35 as a double


% --- Executes during object creation, after setting all properties.
function edit35_CreateFcn(hObject, eventdata, handles)
% hObject    handle to edit35 (see GCBO)
% eventdata  reserved - to be defined in a future version of MATLAB
% handles    empty - handles not created until after all CreateFcns called

% Hint: edit controls usually have a white background on Windows.
%       See ISPC and COMPUTER.
if ispc && isequal(get(hObject,'BackgroundColor'), get(0,'defaultUicontrolBackgroundColor'))
    set(hObject,'BackgroundColor','white');
end


function edit36_Callback(hObject, eventdata, handles)
% hObject    handle to edit36 (see GCBO)
% eventdata  reserved - to be defined in a future version of MATLAB
% handles    structure with handles and user data (see GUIDATA)

% Hints: get(hObject,'String') returns contents of edit36 as text
%        str2double(get(hObject,'String')) returns contents of edit36 as a double


% --- Executes during object creation, after setting all properties.
function edit36_CreateFcn(hObject, eventdata, handles)
% hObject    handle to edit36 (see GCBO)
% eventdata  reserved - to be defined in a future version of MATLAB
% handles    empty - handles not created until after all CreateFcns called

% Hint: edit controls usually have a white background on Windows.
%       See ISPC and COMPUTER.
if ispc && isequal(get(hObject,'BackgroundColor'), get(0,'defaultUicontrolBackgroundColor'))
    set(hObject,'BackgroundColor','white');
end


% --- Executes on button press in checkbox1.
function checkbox1_Callback(hObject, eventdata, handles)
% hObject    handle to checkbox1 (see GCBO)
% eventdata  reserved - to be defined in a future version of MATLAB
% handles    structure with handles and user data (see GUIDATA)

% Hint: get(hObject,'Value') returns toggle state of checkbox1


% --- Executes on button press in checkbox4.
function checkbox4_Callback(hObject, eventdata, handles)
% hObject    handle to checkbox4 (see GCBO)
% eventdata  reserved - to be defined in a future version of MATLAB
% handles    structure with handles and user data (see GUIDATA)

% Hint: get(hObject,'Value') returns toggle state of checkbox4


% --- Executes on button press in checkbox3.
function checkbox2_Callback(hObject, eventdata, handles)
% hObject    handle to checkbox3 (see GCBO)
% eventdata  reserved - to be defined in a future version of MATLAB
% handles    structure with handles and user data (see GUIDATA)

% Hint: get(hObject,'Value') returns toggle state of checkbox3


% --- Executes on button press in checkbox3.
function checkbox3_Callback(hObject, eventdata, handles)
% hObject    handle to checkbox3 (see GCBO)
% eventdata  reserved - to be defined in a future version of MATLAB
% handles    structure with handles and user data (see GUIDATA)

% Hint: get(hObject,'Value') returns toggle state of checkbox3


function edit44_Callback(hObject, eventdata, handles)
% hObject    handle to edit44 (see GCBO)
% eventdata  reserved - to be defined in a future version of MATLAB
% handles    structure with handles and user data (see GUIDATA)

% Hints: get(hObject,'String') returns contents of edit44 as text
%        str2double(get(hObject,'String')) returns contents of edit44 as a double


% --- Executes during object creation, after setting all properties.
function edit44_CreateFcn(hObject, eventdata, handles)
% hObject    handle to edit44 (see GCBO)
% eventdata  reserved - to be defined in a future version of MATLAB
% handles    empty - handles not created until after all CreateFcns called

% Hint: edit controls usually have a white background on Windows.
%       See ISPC and COMPUTER.
if ispc && isequal(get(hObject,'BackgroundColor'), get(0,'defaultUicontrolBackgroundColor'))
    set(hObject,'BackgroundColor','white');
end


% --- Executes on button press in pushbutton11.
function pushbutton11_Callback(hObject, eventdata, handles)
% hObject    handle to pushbutton11 (see GCBO)
% eventdata  reserved - to be defined in a future version of MATLAB
% handles    structure with handles and user data (see GUIDATA)
    load('results.mat');
    
    handles.edit44.Enable                        ='off';
    handles.checkbox1.Enable                     ='off';
    handles.checkbox2.Enable                     ='off';
    handles.checkbox3.Enable                     ='off';
    handles.checkbox4.Enable                     ='off';
    handles.checkbox7.Enable                     ='off';
    handles.checkbox8.Enable                     ='off';

    
    MAXX=floor(length(Data.D)*str2num(handles.edit44.String));
    
if handles.checkbox1.Value==1
figure(1);
      subplot(2,1,1)
      plot(1:1:MAXX,Data.D(1:MAXX),'color',[1 0 0])
      hold on
      plot(MAXX:1:str2num(handles.edit39.String)+1,Data.D(MAXX:end),'color',[0 0 1])
      xlabel('Iteration')
      ylabel('Diff. coeff. (\mum^2/s) in Log scale')
      set(gca,'yscale','log')

      subplot(2,1,2)

      bnd = logspace(log10(min(Data.D(MAXX:end)))-1,log10(max(Data.D(MAXX:end)))+1,100);
      histogram(Data.D(MAXX:end),bnd,'Normalization','pdf')
      set(gca,'xscale','log')
      xlabel('Diff. coeff. (\mum^2/s)');
      ylabel('Posterior');
end
    
      
if handles.checkbox2.Value==1
figure(3)
      subplot(2,1,1)
      plot(1:1:MAXX,Data.mu(1:MAXX),'color',[1 0 0])
      hold on
      plot(MAXX:1:str2num(handles.edit39.String)+1,Data.mu(MAXX:end),'color',[0 0 1])
      xlabel('Iteration')
      ylabel({'Molecule emission rate' ;'(photons/s)'})
      
      subplot(2,1,2)
      histogram(Data.mu(MAXX:end),'Normalization','pdf')
      hold on
      xlim([0 max(Data.mu(MAXX:end))])
      xlabel('Molecule emission rate (photons/s)')
      ylabel('Posterior')
      
end

if handles.checkbox3.Value==1
figure(2)

maxx_size_signals=size(Data.b,1);
      for k=1:maxx_size_signals
      subplot(2,maxx_size_signals,k)
      plot(1:1:MAXX,reshape(sum(Data.b(k,:,1:MAXX),2),1,MAXX),'color',[1 0 0])
      hold on
      plot(MAXX:1:str2num(handles.edit39.String)+1,reshape(sum(Data.b(k,:,MAXX:end),2),1,size(Data.b,3)-MAXX+1),'color',[0 0 1])
      xlabel('Iteration')
      ylabel('Active molecules')
      ylim([0 str2num(handles.Number_particles.String)])
      title(['Signal ',num2str(k)])
      
      subplot(2,maxx_size_signals,maxx_size_signals+k)
      histogram(reshape(sum(Data.b(k,:,MAXX:end),2),1,size(Data.b,3)-MAXX+1),'Normalization','pdf')
      xlim([0 str2num(handles.Number_particles.String)])
      xlabel('Active molecules')
      ylabel('Posterior')
      end
end

if handles.checkbox4.Value==1
    
   wxy    = str2double(handles.wxy.String);
   wz     = str2double(handles.wz.String);
   radiii = str2num(handles.concen_radious.String);
   ell    = char(hex2dec(strsplit('2113')));
   
   figure(4)
 
      maxx_size_signals=size(Data.concentration,1);
      len=size(Data.concentration,2);
     
   if  sum(handles.edit51.String)  == sum('Cylindrical')
       PSF_TYPE = 2;
   else
       PSF_TYPE = 1;
   end
   
   
      for i=1:maxx_size_signals
          mean25_stairs=[];
          mean75_stairs=[];
          timee=[];
          time=Data.Time_partial{i}./str2num(handles.Delta_t.String);
          
      for l=1:length(radiii)
          concenn25=[];
          concenn50=[];
          concenn75=[];
          
          switch PSF_TYPE
              
          case 1
              
              for k = 1 : length(Data.concentration(i,1,1:length(Data.Time_partial{i}),1))
              concenn25(1,k)=quantile(reshape(Data.concentration(i,:,k,l),len,1),0.025) ;
              concenn50(1,k)=quantile(reshape(Data.concentration(i,:,k,l),len,1),0.50) ;
              concenn75(1,k)=quantile(reshape(Data.concentration(i,:,k,l),len,1),0.975) ;
              end 
         otherwise
              
              for k = 1 : length(Data.concentration(i,1,1:length(Data.Time_partial{i}),1))
              concenn25(1,k)=quantile(reshape(Data.concentration(i,:,k,l),len,1),0.025) ;
              concenn50(1,k)=quantile(reshape(Data.concentration(i,:,k,l),len,1),0.50) ;
              concenn75(1,k)=quantile(reshape(Data.concentration(i,:,k,l),len,1),0.975) ;
              end     
              
          end
          
          
          
          for k=1:length(Data.concentration(i,1,1:length(Data.Time_partial{i}),1))
              mean25_stairs(2*k-1:2*k)=concenn25(k);
              mean75_stairs(2*k-1:2*k)=concenn75(k);
    
              if  k<length(Data.concentration(i,1,1:length(Data.Time_partial{i}),1))
            
                  mean75_stairs(2*k)=concenn75(k+1);
                  mean25_stairs(2*k)=concenn25(k+1);
              else
                  mean75_stairs(2*k)=concenn75(k);
                  mean25_stairs(2*k)=concenn25(k);
              end
    
              timee(2*k-1:2*k)=time(k);
              timee(2*k)=0;
          end
       
       

          subplot(length(radiii) ,maxx_size_signals ,(l-1)*maxx_size_signals+i)
          
          x2 = [cumsum(timee),fliplr(cumsum(timee))];
          inBetween = [mean25_stairs, fliplr(mean75_stairs)];
          h2=fill(x2, inBetween, 'm','facealpha', 0.2,'edgecolor', 'none');
          
          hold on
          h1=stairs(cumsum([0 time]),[0 concenn50],'b');
          xlim([0 sum(time)])
          ylim([0 unique(max(concenn75))*1.5+0.0001])
          xlabel(['Time step (',num2str(str2num(handles.Delta_t.String)),' s)'])
          if l==1
              title(['signal ',num2str(i)])
          end
          
          if i==1
              ylabel([{['Concentration N_k^{',ell,'}']};{[ell,'=',num2str(radiii(l))]}],'Fontsize',11)
          end
          box off
          
          if i==maxx_size_signals
          [BL,BLicons]=legend([h1,h2],'Median of posterior','95% confidence interval','location','Eastoutside');
          PatchInLegend = findobj(BLicons(end), 'type', 'patch');
          set(PatchInLegend, 'facea', 0.2)
          set(BL,'box','off')
          end
          
          
      end
      
      end
      

end


if handles.checkbox8.Value==1
   figure(5)
   num_signals=size(Data.b,1);
   if sum(handles.pushbutton22.String)==sum('Estimate background emission rate (ON)')
      Results=[Data.D(MAXX:end)   ;   Data.mu(MAXX:end)   ;   Data.mu_back(MAXX:end);...
            reshape(sum(Data.b(:,:,MAXX:end),2),num_signals,size(Data.D(MAXX:end),2))+0.2*rand(size(Data.D(MAXX:end)));...
            reshape(sum(Data.q(:,:,MAXX:end),2),num_signals,size(Data.D(MAXX:end),2))]';
      numParams=size(Results,2);
      Titles={'D','\mu_{mol}','\mu_{back}'};
      for k=1:num_signals
          Titles(k+3)={['b' , num2str(k)]};
          Titles(k+num_signals+3)={['q' , num2str(k)]};
      end
   else
    
      Results=[Data.D(MAXX:end)   ;   Data.mu(MAXX:end)    ;...
            reshape(sum(Data.b(:,:,MAXX:end),2),num_signals,size(Data.D(MAXX:end),2))+0.2*rand(size(Data.D(MAXX:end)));...
            reshape(sum(Data.q(:,:,MAXX:end),2),num_signals,size(Data.D(MAXX:end),2))]';
      numParams=size(Results,2);
    
      Titles={'D','\mu_{mol}'};
      for k=1:num_signals
          Titles(k+2)={['b' , num2str(k)]};
          Titles(k+num_signals+2)={['q' , num2str(k)]};
      end
   end
   for i=1:numParams
       for j=i:numParams
           subplot(numParams,numParams,numParams*(i-1)+i+(j-i));
           if i~=j
            
            histogram2(Results(:,i),Results(:,j),'Normalization','Probability');
            title([Titles{i},' , ',Titles{j}])
            ylabel(Titles{j})
            xlabel(Titles{i})
            zlabel('Joint Posterior')
            hold on
           end
           if i==j
           histogram(Results(:,j),'Normalization','pdf')
           ylabel('Posterior')
           xlabel(Titles{i})
           box off
           hold on 
           end
       end
   end
end


if sum(handles.pushbutton22.String)==sum('Estimate background emission rate (ON)')  
   if handles.checkbox7.Value==1
      figure(6)
      
      subplot(2,1,1)
      plot(1:1:MAXX,Data.mu_back(1:MAXX),'color',[1 0 0])
      hold on
      plot(MAXX:1:str2num(handles.edit39.String)+1,Data.mu_back(MAXX:end),'color',[0 0 1])
      xlabel('Iteration')
      ylabel({'Background photon emission rate' ;'(photons/s)'})
      
      subplot(2,1,2)
      histogram(Data.mu_back(MAXX:end),'Normalization','pdf')
      hold on
      xlim([0 max(Data.mu_back(MAXX:end))])
      xlabel('Background photon emission rate (photons/s)')
      ylabel('Posterior')
   end
   handles.checkbox7.Enable                     ='on';
end


    drawnow
    handles.edit44.Enable                        ='on';
    handles.checkbox1.Enable                     ='on';
    handles.checkbox2.Enable                     ='on';
    handles.checkbox3.Enable                     ='on';
    handles.checkbox4.Enable                     ='on';
    handles.checkbox8.Enable                     ='on';


% --- Executes on button press in pushbutton10.
function pushbutton10_Callback(hObject, eventdata, handles)
% hObject    handle to pushbutton10 (see GCBO)
% eventdata  reserved - to be defined in a future version of MATLAB
% handles    structure with handles and user data (see GUIDATA)


    
    handles.D_alpha_prior.Enable           ='off';
    handles.D_beta_prior.Enable            ='off';
    handles.mu_alpha.Enable                ='off';
    handles.mu_beta.Enable                 ='off';
    handles.Number_particles.Enable        ='off';
    handles.alpha_load.Enable              ='off';
    handles.beta_load.Enable               ='off';
    handles.concen_radious.Enable          ='off';
    handles.mu_prior_xy.Enable             ='off';
    handles.mu_prior_z.Enable              ='off';
    handles.var_prior_xy.Enable            ='off';
    handles.var_prior_z.Enable             ='off';
    handles.pushbutton11.Enable            ='off';
    handles.pushbutton16.Enable            ='off';
    handles.pushbutton22.Enable            ='off';
    handles.pushbutton17.Enable            ='off';
    handles.pushbutton23.Enable            ='off';
    handles.pushbutton20.Enable            ='off';
    handles.pushbutton8.Enable             ='off';
    handles.checkbox1.Enable               ='off';
    handles.checkbox2.Enable               ='off';
    handles.checkbox3.Enable               ='off';
    handles.checkbox4.Enable               ='off';
    handles.checkbox7.Enable               ='off';
    handles.checkbox8.Enable               ='off';
    handles.edit44.Enable                  ='off';
    handles.edit52.Enable                  ='off';
    handles.edit39.Enable                  ='off';
    handles.edit40.Enable                  ='off';
    handles.edit41.Enable                  ='off';
    handles.edit46.Enable                  ='off';
    handles.edit47.Enable                  ='off';
    handles.edit48.Enable                  ='off';
    handles.edit53.Enable                  ='off';
    
    
    cla(handles.axes7)
    cla(handles.axes8)
    cla(handles.axes15)
    cla(handles.axes16)
    cla(handles.axes18)
    cla(handles.axes19)
    cla(handles.axes20)
    
    
    
    min = str2num(get(handles.edit35,'String')) ;
    max = str2num(get(handles.edit36,'String')) ;
   
    load('results.mat');
    
    signal=Data.Trace_uniform_bin;
    
    while isempty(max-min) || max-min<=0 || max<0 || min<0 || max>length(signal)-1

              opts.Interpreter = 'tex';
              answer = inputdlg({'Minimum Time step','Maximum Time step'},...
                       'Please choose some reasonable vlues!',[1 60;1 60],...
                       {'0',num2str(length(signal)-1)},opts);
                   
              min = str2num(answer{1}) ;
              max = str2num(answer{2}) ;
    end

    handles.edit35.String  = num2str(min);
    handles.edit36.String  = num2str(max);
    
    
    signal_partial = signal(min+1:max+1);
    

    plot(handles.axes7,min:1:max,signal_partial)
    hold (handles.axes7,'off')

    xlabel(handles.axes7,['Time step (\Deltat =', num2str(str2num(handles.Delta_t.String)), 's)'])
    ylabel(handles.axes7,'Observed photons')
    xlim(handles.axes7,[min max])

    if length(handles.uitable5.Data)>0
       handles.pushbutton28.Enable             ='on' ;
    else
        handles.pushbutton28.Enable            ='off';
        handles.pushbutton30.Enable            ='off';
        handles.pushbutton31.Enable            ='off';
    end
    
    handles.pushbutton29.Enable            ='on';
    handles.edit53.Enable                  ='on';
    
    handles.text36.Enable                  ='on';
    handles.text37.Enable                  ='on';
    
    
    
    
    
    
% --- Executes on button press in Import_signal.
function Import_signal_Callback(hObject, eventdata, handles)
% hObject    handle to Import_signal (see GCBO)
% eventdata  reserved - to be defined in a future version of MATLAB
% handles    structure with handles and user data (see GUIDATA)

    handles.D_alpha_prior.Enable           ='off';
    handles.D_beta_prior.Enable            ='off';
    handles.mu_alpha.Enable                ='off';
    handles.mu_beta.Enable                 ='off';
    handles.Number_particles.Enable        ='off';
    handles.alpha_load.Enable              ='off';
    handles.beta_load.Enable               ='off';
    handles.concen_radious.Enable          ='off';
    handles.mu_prior_xy.Enable             ='off';
    handles.mu_prior_z.Enable              ='off';
    handles.var_prior_xy.Enable            ='off';
    handles.var_prior_z.Enable             ='off';
    
    handles.pushbutton8.Enable             ='off';
    handles.pushbutton10.Enable            ='off';
    handles.pushbutton11.Enable            ='off';
    handles.pushbutton16.Enable            ='off';
    handles.pushbutton17.Enable            ='off';
    handles.pushbutton22.Enable            ='off';
    handles.pushbutton23.Enable            ='off';
    handles.pushbutton28.Enable            ='off';
    handles.pushbutton20.Enable            ='off';
    handles.pushbutton29.Enable            ='off';
    handles.pushbutton30.Enable            ='off';
    handles.pushbutton31.Enable            ='off';
    handles.pushbutton32.Enable            ='off';
    handles.pushbutton33.Enable            ='off';
    
    handles.edit35.Enable                  ='off';
    handles.edit36.Enable                  ='off';
    handles.edit39.Enable                  ='off';
    handles.edit40.Enable                  ='off';
    handles.edit41.Enable                  ='off';
    handles.edit46.Enable                  ='off';
    handles.edit44.Enable                  ='off';
    handles.edit47.Enable                  ='off';
    handles.edit48.Enable                  ='off';
    handles.edit52.Enable                  ='off';
    handles.edit53.Enable                  ='off';
    
    handles.checkbox1.Enable               ='off';
    handles.checkbox2.Enable               ='off';
    handles.checkbox3.Enable               ='off';
    handles.checkbox4.Enable               ='off';
    handles.checkbox7.Enable               ='off';
    handles.checkbox8.Enable               ='off';
    
    cla(handles.Original_signal)
    cla(handles.axes7)
    cla(handles.axes8)
    cla(handles.axes15)
    cla(handles.axes16)
    cla(handles.axes18)
    cla(handles.axes19)
    cla(handles.axes20)
    handles.uitable5.Data                         =  [] ;
    
    handles.Import_signal.Enable                  ='off';
    handles.Generate_Synthetic_signal.Enable      ='off';
    handles.Measure_Background.Enable             ='off';

    
    opts.Interpreter = 'tex';
    answer = inputdlg({['Background photon emission rate (photons/s)',newline,...
                        'If you already import a background trace, leave this part!!'],...
                        'Confocal radius in xy plane (\mu m)','Confocal radius in z axis (\mu m)'},...
                        'Parameters',[ 1 60 ; 1 60 ; 1 60 ],...
                       {handles.mu_back.String,'0.3','1.5'},opts);
                   
        
                   
  
    if length(answer)==3
    while isempty(str2num(answer{2})) || str2num(answer{2})==0 ||...
          isempty(str2num(answer{3})) || str2num(answer{3})==0 
          
          
          opts.Interpreter = 'tex';
          answer = inputdlg({['Background photon emission rate (photons/s)',newline,...
                              'If you already import a background trace, leave this part!!'],...
                              'Confocal radius in xy plane (\mu m)','Confocal radius in z axis (\mu m)'},...
                              'Parameters',[ 1 60 ; 1 60 ; 1 60],...
                          {handles.mu_back.String,'0.3','1.5'},opts);
    end
    
    wxy   = str2num(answer{2});
    wz    = str2num(answer{3});
   
    
    if  length(answer{1})==length('-')
        handles.mu_back.String = 1000;
    else
        handles.mu_back.String                 = answer{1};
    end
    handles.wxy.String                     = num2str(wxy);
    handles.wz.String                      = num2str(wz);
   
    
    
    
    choice = questdlg({'Do you want to import a single photon arrival time trace?'},...
                       'Sample analysis','Yes','No','Yes');
    
    if  strcmp(choice,'Yes')
  
    opts.Interpreter = 'tex';
    answer = inputdlg({'Bin Size (s)   "Smaller Bin size -> longer calculation time"'},...
                           'Parameters',[1 60],{handles.Delta_t.String},opts);
  
    while isempty(str2num(answer{1})) || str2num(answer{1})==0
          opts.Interpreter = 'tex';
          answer = inputdlg({'Bin Size (s)   "smaller Bin size -> longer calculation time"'},...
                                 'Parameters',[1 60],{handles.Delta_t.String},opts);
    end
    Delta_t = str2num(answer{1});
    handles.Delta_t.String=answer{1};

% Find the file from the directory
    [filename, pathname] = uigetfile( { '*.txt','Text-files (*.txt)';...
                                        '*.mat','MAT-files (*.mat)' }, ...
                                        'Import a time series', ...
                                        'MultiSelect', 'off');

% Check the filename                             
    if ~isequal(filename,0)
        
% Load the file
       Trace_singel_photon = load(filename,pathname);

% Bin the single phton arrival trace based on the bin size Delta_t
       edge=cumsum([0,repmat(Delta_t,1,floor(Trace_singel_photon(end)/Delta_t)+1)]);
       [Trace_uniform_bin,~] = histcounts(Trace_singel_photon,edge);

% plot data
       plot(handles.Original_signal,0:1:length(Trace_uniform_bin)-1,Trace_uniform_bin)
       xlabel(handles.Original_signal,['Time step (\Deltat=' num2str(Delta_t) 's)'])
       ylabel(handles.Original_signal,'Observed photons')

% Save the imported trace in forms of single photon arrival times and bined trace
       Data.Trace_singel_photon = Trace_singel_photon ;
       Data.Trace_uniform_bin   = Trace_uniform_bin   ;
       Data.Trace_type          = 'Time_arrival'      ;
       
       save('results','Data')
    end   
    
    
    else
        choice1 = questdlg({'Do you want to import a binned signal?'},'Sample analysis','Yes','No','Yes');
        if  strcmp(choice1,'Yes')
            
            opts.Interpreter = 'tex';
            answer = inputdlg({'Bin Size(s)'});
  
            while isempty(str2num(answer{1})) || str2num(answer{1})==0
                  opts.Interpreter = 'tex';
                  answer = inputdlg({'Bin Size(s)'});
            end
            Delta_t = str2num(answer{1});
            handles.Delta_t.String=answer{1};
            
            [filename, pathname] = uigetfile( { '*.txt','Text-files (*.txt)'  ;...
                                                '*.mat','MAT-files (*.mat)' } , ...
                                                'Import a time series'        , ...
                                                'MultiSelect'                 , 'off');
            
             if ~isequal(filename,0)
                Trace_uniform_bin = load(filename,pathname);
    
                
                % plot data
                plot(handles.Original_signal,0:1:length(Trace_uniform_bin)-1,Trace_uniform_bin)
                xlabel(handles.Original_signal,['Time step (\Deltat=' num2str(Delta_t) 's)'])
                ylabel(handles.Original_signal,'Observed photons')
                
               % Save the imported trace in forms of single photon arrival times and bined trace
                Data.Trace_uniform_bin   = Trace_uniform_bin   ;
                Data.Trace_type          = 'Binn_Trace'        ;
                save('results','Data');
   
             end
         end
   

    end
    
    % Set the handles
    handles.pushbutton10.Enable                   ='on';
    handles.edit53.Enable                         ='on';
           
    handles.edit35.String                         ='0' ;
    handles.edit36.String                         ='0' ;
            
    handles.edit35.Enable                         ='on';
    handles.edit36.Enable                         ='on';
   
    handles.Import_signal.Enable                  ='on';
    handles.Generate_Synthetic_signal.Enable      ='on';
    handles.Measure_Background.Enable             ='on';
    handles.pushbutton32.Enable                   ='on';
    handles.pushbutton33.Enable                   ='on';
    else
    handles.Import_signal.Enable                  ='on';
    handles.Generate_Synthetic_signal.Enable      ='on';
    handles.Measure_Background.Enable             ='on';
    handles.pushbutton32.Enable                   ='on';
    handles.pushbutton33.Enable                   ='on';
    end
    

    
        
        
        


    
    
% --- Executes on button press in Generate_Synthetic_signal.
function Generate_Synthetic_signal_Callback(hObject, eventdata, handles)
% hObject    handle to Generate_Synthetic_signal (see GCBO)
% eventdata  reserved - to be defined in a future version of MATLAB
% handles    structure with handles and user data (see GUIDATA)
    
    S = dir('*.mat');                   % Collect directories
    N = {S.name};                       % Collect Names of the text files
    for i=1:length(N)                   % A For loop for read data
        delete(N{i});           % Import data from the file
    end
    clear('S','N')
    
    handles.Import_signal.Enable                ='off';
    handles.Generate_Synthetic_signal.Enable    ='off';
    handles.Measure_Background.Enable           ='off';

     
    handles.uitable5.Data                       =   [];
       
    handles.edit35.Enable                       ='off';
    handles.edit36.Enable                       ='off';
    handles.edit39.Enable                       ='off';
    handles.edit40.Enable                       ='off';
    handles.edit41.Enable                       ='off';
    handles.edit44.Enable                       ='off';
    handles.edit46.Enable                       ='off';
    handles.edit47.Enable                       ='off';
    handles.edit48.Enable                       ='off';
    handles.edit52.Enable                       ='off';
    handles.edit53.Enable                       ='off';
    
    handles.concen_radious.Enable               ='off';
    handles.D_alpha_prior.Enable                ='off';
    handles.D_beta_prior.Enable                 ='off'; 
    handles.mu_alpha.Enable                     ='off';
    handles.mu_beta.Enable                      ='off'; 
    handles.Number_particles.Enable             ='off'; 
    handles.alpha_load.Enable                   ='off';
    handles.beta_load.Enable                    ='off';
    handles.mu_prior_xy.Enable                  ='off';
    handles.mu_prior_z.Enable                   ='off';
    handles.var_prior_xy.Enable                 ='off';
    handles.var_prior_z.Enable                  ='off';
    
    handles.pushbutton8.Enable                  ='off'; 
    handles.pushbutton10.Enable                 ='off';
    handles.pushbutton11.Enable                 ='off';
    handles.pushbutton16.Enable                 ='off';
    handles.pushbutton17.Enable                 ='off';
    handles.pushbutton19.Enable                 ='off';
    handles.pushbutton20.Enable                 ='off';
    handles.pushbutton21.Enable                 ='off';
    handles.pushbutton22.Enable                 ='off';
    handles.pushbutton23.Enable                 ='off';
    handles.pushbutton24.Enable                 ='off';
    handles.pushbutton25.Enable                 ='off';
    handles.pushbutton26.Enable                 ='off';
    handles.pushbutton28.Enable                 ='off';
    handles.pushbutton29.Enable                 ='off';
    handles.pushbutton30.Enable                 ='off';
    handles.pushbutton31.Enable                 ='off';
    handles.pushbutton32.Enable                 ='off';
    handles.pushbutton33.Enable                 ='off';
    
    handles.edit44.Enable                       ='off';
    handles.checkbox1.Enable                    ='off';
    handles.checkbox2.Enable                    ='off';
    handles.checkbox3.Enable                    ='off';
    handles.checkbox4.Enable                    ='off';
    handles.checkbox7.Enable                    ='off';
    handles.checkbox8.Enable                    ='off';

    cla(handles.Original_signal)
    cla(handles.axes7)
    cla(handles.axes8)
    cla(handles.axes15)
    cla(handles.axes16)
    cla(handles.axes18)
    cla(handles.axes19)
    cla(handles.axes20)
    
answer={};

    opts.Interpreter = 'tex';
    answer = inputdlg({'Length of the Trace (Time step)',...
                       'Minimum Data acquisition time (s)','Bin size (s)',...
                       'Confocal radius in xy plane (\mum)',...
                       'Confocal radius in z axis (\mum)',...
                       'Diffusion coefficent (\mum^2/s)',...
                       'Background photon emission rate (photons/s)',...
                       'Molecule photon emission rate (photons/s)',...
                       'Radius of the periodic boundary in xy plane (\mum)',...
                       'Radius of the periodic boundary in z axis (\mum)',...
                       'Number of molecules'},...
                       'Parameters',...
                       [1 50;1 50; 1 50; 1 50; 1 50; 1 50; 1 50; 1 50; 1 50; 1 50; 1 50],...
                       {'10^7','10^-7','10^-4','0.3','3.5','10','1000','50000','2','3','30'},opts);
                  
              
if length(answer)==11
    while isempty(str2num(answer{1})) || str2num(answer{1})==0 ||...
          isempty(str2num(answer{2})) || str2num(answer{2})==0 ||...
          isempty(str2num(answer{3})) || str2num(answer{3})==0 ||...
          isempty(str2num(answer{4})) || str2num(answer{4})==0 ||...
          isempty(str2num(answer{5})) || str2num(answer{5})==0 ||...
          isempty(str2num(answer{6})) || str2num(answer{6})==0 ||...
          isempty(str2num(answer{7})) || str2num(answer{7})==0 ||...
          isempty(str2num(answer{8})) || str2num(answer{8})==0 ||...
          isempty(str2num(answer{9})) || str2num(answer{9})==0 ||...
          isempty(str2num(answer{10}))|| str2num(answer{10})==0||...
          isempty(str2num(answer{11}))|| str2num(answer{11})==0
          
          opts.Interpreter = 'tex';
          answer = inputdlg({'Length of the Trace (Time step)',...
                             'Minimum Data acquisition time (s)',...
                             'Bin size (s)',...
                             'Confocal radius in xy plane (\mum)',...
                             'Confocal radius in z axis (\mum)',...
                             'Diffusion coefficent (\mum^2/s)',...
                             'Background photon emission rate(photons/s)',...
                             'Molecule photon emission rate (photons/s)',...
                             'Radius of the periodic boundary in xy plane (\mum)',...
                             'Radius of the periodic boundary in z axis (\mum)',...
                             'Number of molecules'},...
                             'Parameters',...
                             [1 50;1 50; 1 50; 1 50; 1 50; 1 50; 1 50; 1 50; 1 50; 1 50; 1 50],...
                             {'10^7','10^-7','10^-4','0.3','3.5','10','1000','50000','2','3','30'},opts);
    end

    Synthetic_lenght   = str2num(answer{1});
    deltat             = str2num(answer{2});
    Delta_t            = str2num(answer{3});
    wxy                = str2num(answer{4});
    wz                 = str2num(answer{5});
    Diffusion          = str2num(answer{6});
    mu_back            = str2num(answer{7});
    mu                 = str2num(answer{8});
    Lxy                = str2num(answer{9});
    Lz                 = str2num(answer{10});
    N                  = str2num(answer{11});
   

    
    choice = questdlg('Please choose the Point Spread Function of interest:','Point Spread Function','Gaussian','Lorentzian','Cylindrical','');
    
    if     strcmp(choice,'Gaussian')
           handles.edit51.String = 'Gaussian' ;
           
    elseif strcmp(choice,'Lorentzian')
        
           handles.edit51.String = 'Lorentzian' ;
           
    elseif strcmp(choice,'Cylindrical')
        
           wz            = 100;
           handles.edit51.String = 'Cylindrical' ;
    else
           handles.edit51.String = 'NAN' ;
           handles.Import_signal.Enable                  ='on';
           handles.Generate_Synthetic_signal.Enable      ='on';
           handles.Measure_Background.Enable             ='on';
    end
    
    
  
    
    handles.deltat.String                  = answer{2};
    handles.Delta_t.String                 = answer{3};
    handles.Diff_synth.String              = answer{6};
    handles.mu_back.String                 = answer{7};  
    handles.mu_synth.String                = answer{8};
    handles.Num_synth.String               = answer{11};
    handles.wxy.String                     = num2str(wxy);
    handles.wz.String                      = num2str(wz);
    
    
    

    
    
% sample generator function, you should direct it to the same folder
[ Trace_singel_photon ] = Sample_Generator( ...
...
...
Lxy , Lz      , Synthetic_lenght      , N         , ...
wxy , wz      , Diffusion             , deltat    , ...
mu  , mu_back , handles.edit51.String , handles  );


if length(Trace_singel_photon)<1   
    text(handles.Original_signal,0.4 ,0.5 ,{'No Photon is detected';'Please redo the generation with lorger time steps'})
else
    handles.pushbutton10.Enable            ='on';
    handles.edit53.Enable                  ='on';
    
    handles.edit35.String                  ='0';
    handles.edit36.String                  ='0';
    
    handles.edit35.Enable                  ='on';
    handles.edit36.Enable                  ='on';


    
% Bin the single phton arrival trace based on the bin size Delta_t
edge=cumsum([0,repmat(Delta_t,1,floor(max(Trace_singel_photon)/Delta_t)+1)]);
[Trace_uniform_bin,~] = histcounts(Trace_singel_photon,edge);


Data.Trace_singel_photon = Trace_singel_photon  ;
Data.Trace_uniform_bin   = Trace_uniform_bin    ;
Data.Trace_type          = 'Time_arrival'       ;

save('results','Data')

% set(handles.Plot_signal,signal)
plot(handles.Original_signal,0:1:length(Trace_uniform_bin)-1,Trace_uniform_bin)
xlabel(handles.Original_signal,['Time step (\Deltat=' num2str(Delta_t) 's)'])
ylabel(handles.Original_signal,'Observed photons')
end

   handles.pushbutton19.Enable                   ='on';
end

   handles.Import_signal.Enable                  ='on';
   handles.Generate_Synthetic_signal.Enable      ='on';
   handles.Measure_Background.Enable             ='on';
   handles.pushbutton32.Enable                   ='on';
   handles.pushbutton33.Enable                   ='on';


     


% --- Executes on button press in Measure_Background.
function Measure_Background_Callback(hObject, eventdata, handles)
% hObject    handle to Measure_Background (see GCBO)
% eventdata  reserved - to be defined in a future version of MATLAB
% handles    structure with handles and user data (see GUIDATA)

    handles.pushbutton19.Enable            ='on';
    cla(handles.Original_signal)
    cla(handles.axes7)
    cla(handles.axes8)
    cla(handles.axes15)
    cla(handles.axes16)
    cla(handles.axes18)
    cla(handles.axes19)
    cla(handles.axes20)
    handles.uitable5.Data                    =  [];
     
    handles.Refractive_index.String          = '-';
    handles.Delta_t.String                   = '-';
    handles.Laser_Wavelength.String          = '-';
    handles.Objective_magnification.String   = '-';
    handles.wxy.String                       = '-';
    handles.wz.String                        = '-';
    handles.mu_back.String                   = '-';
    handles.Diff_synth.String                = '-';
    handles.mu_synth.String                  = '-';
    handles.Num_synth.String                 = '-';
    
    handles.edit35.String                    = '0'   ;
    handles.edit36.String                    = '0'   ;
    handles.edit39.String                    = '1000';
    handles.edit40.String                    = '1000';
    handles.edit41.String                    = '10'  ;
    
    
    handles.D_alpha_prior.Enable             ='off';
    handles.D_beta_prior.Enable              ='off';
    handles.mu_alpha.Enable                  ='off';
    handles.mu_beta.Enable                   ='off';
    handles.Number_particles.Enable          ='off';
    handles.Measure_Background.Enable        ='off';
    handles.Generate_Synthetic_signal.Enable ='off';
    handles.Import_signal.Enable             ='off';
    handles.alpha_load.Enable                ='off';
    handles.beta_load.Enable                 ='off';
    handles.concen_radious.Enable            ='off';
    handles.mu_prior_xy.Enable               ='off';
    handles.mu_prior_z.Enable                ='off';
    handles.var_prior_xy.Enable              ='off';
    handles.var_prior_z.Enable               ='off';
    handles.pushbutton10.Enable              ='off';
    handles.edit35.Enable                    ='off';
    handles.edit36.Enable                    ='off';
    handles.edit39.Enable                    ='off';
    handles.edit40.Enable                    ='off';
    handles.edit41.Enable                    ='off';
    handles.edit53.Enable                    ='off';
    handles.pushbutton8.Enable               ='off';
    handles.pushbutton11.Enable              ='off';
    handles.pushbutton17.Enable              ='off';
    handles.pushbutton20.Enable              ='off';
    handles.pushbutton21.Enable              ='off';
    handles.pushbutton22.Enable              ='off';
    handles.pushbutton23.Enable              ='off';
    handles.pushbutton24.Enable              ='off';
    handles.pushbutton25.Enable              ='off';
    handles.pushbutton26.Enable              ='off';
    handles.pushbutton32.Enable              ='off';
    handles.pushbutton33.Enable              ='off';
    
    handles.edit44.Enable                    ='off';
    handles.checkbox1.Enable                 ='off';
    handles.checkbox2.Enable                 ='off';
    handles.checkbox3.Enable                 ='off';
    handles.checkbox4.Enable                 ='off';
    handles.checkbox7.Enable                 ='off';
    handles.checkbox8.Enable                 ='off';
    

    
    
    choice = questdlg({'Do you want to import a single photon arrival time signal?'},'Sample analysis','Yes','No','Yes');
    
    if  strcmp(choice,'Yes')
    
        
        opts.Interpreter = 'tex';
        answer = inputdlg({'Bin Size (s)   "smaller Bin size=longer calculation time"'},...
                           'Parameters',[1 60],{handles.Delta_t.String},opts);
  
        while isempty(str2num(answer{1})) || str2num(answer{1})==0
              opts.Interpreter = 'tex';
              answer = inputdlg({'Bin Size (s)   "smaller Bin size=longer calculation time"'},...
                                 'Parameters',[1 60],{handles.Delta_t.String},opts);
        end
        Delta_t = str2num(answer{1});
        handles.Delta_t.String=answer{1};
        
        [filename, pathname] = uigetfile( { '*.txt','Text-files (*.txt)';...
                                        '*.mat','MAT-files (*.mat)' }, ...
                                        'Import a time series', ...
                                        'MultiSelect', 'off');
            
     if ~isequal(filename,0)
            b_signal = load(filename,pathname);
            
            edge=cumsum([0,repmat(Delta_t,1,floor(b_signal(end)/Delta_t)+1)]);
            [background_signal,~] = histcounts(b_signal,edge);

            % plot data
            plot(handles.Original_signal,0:1:length(background_signal)-1,background_signal)
            xlabel(handles.Original_signal,['Time step (\Deltat=' num2str(Delta_t) 's)'])
            ylabel(handles.Original_signal,'Observed photons')
   
            mu_back_alpha_prior =  1              ;
            mu_back_beta_prior  = 10              ;
            Max_iter            = 1000            ;
            mu_back_sample      = nan(Max_iter,1) ;
            for i=1:Max_iter
                mu_back_sample(i) = gamrnd(mu_back_alpha_prior+sum(background_signal),...
                        1/(Delta_t*length(background_signal)+(1/mu_back_beta_prior)));
            end
            mu_back=floor(mean(mu_back_sample));
            handles.mu_back.String=num2str(mu_back);
    end
             
    handles.Import_signal.Enable                  ='on';
    handles.Generate_Synthetic_signal.Enable      ='on';
    handles.Measure_Background.Enable             ='on';
    handles.pushbutton32.Enable                   ='on';
    handles.pushbutton33.Enable                   ='on';
    else
        
        
        
     choice1 = questdlg({'Do you want to import a binned signal?'},'Sample analysis','Yes','No','Yes');
        if  strcmp(choice1,'Yes')
            
            opts.Interpreter = 'tex';
            answer = inputdlg({'Bin Size (s)'});
  
            while isempty(str2num(answer{1})) || str2num(answer{1})==0
                  opts.Interpreter = 'tex';
                  answer = inputdlg({'Bin Size(s)'});
            end
            Delta_t = str2num(answer{1});
            handles.Delta_t.String=answer{1};
            
            [filename, pathname] = uigetfile( { '*.txt','Text-files (*.txt)';...
                                        '*.mat','MAT-files (*.mat)' }, ...
                                        'Import a time series', ...
                                        'MultiSelect', 'off');
            
             if ~isequal(filename,0)
                 background_signal = load(filename,pathname);
                % plot data
            plot(handles.Original_signal,0:1:length(background_signal)-1,background_signal)
            xlabel(handles.Original_signal,['Time step (\Deltat=' num2str(Delta_t) 's)'])
            ylabel(handles.Original_signal,'Observed photons')
   
            mu_back_alpha_prior =  1              ;
            mu_back_beta_prior  = 10              ;
            Max_iter            = 1000            ;
            mu_back_sample      = nan(Max_iter,1) ;
            for i=1:Max_iter
                mu_back_sample(i) = gamrnd(mu_back_alpha_prior+sum(background_signal),...
                        1/(Delta_t*length(background_signal)+(1/mu_back_beta_prior)));
            end
            mu_back=floor(mean(mu_back_sample));
            handles.mu_back.String=num2str(mu_back);
                
             end
         end   
        
        
        
        
        
        
        
        
        
        
        
        
        
    handles.Import_signal.Enable                  ='on';
    handles.Generate_Synthetic_signal.Enable      ='on';
    handles.Measure_Background.Enable             ='on'; 
    handles.pushbutton32.Enable                   ='on';
    handles.pushbutton33.Enable                   ='on';
    end


% --- Executes on button press in pushbutton8.
function pushbutton8_Callback(hObject, eventdata, handles)
% hObject    handle to pushbutton8 (see GCBO)
% eventdata  reserved - to be defined in a future version of MATLAB
% handles    structure with handles and user data (see GUIDATA)
     
     
    handles.D_alpha_prior.Enable           ='off';
    handles.D_beta_prior.Enable            ='off';
    handles.mu_alpha.Enable                ='off';
    handles.mu_beta.Enable                 ='off';
    handles.Number_particles.Enable        ='off';
    handles.alpha_load.Enable              ='off';
    handles.beta_load.Enable               ='off';
    handles.concen_radious.Enable          ='off';
    handles.mu_prior_xy.Enable             ='off';
    handles.mu_prior_z.Enable              ='off';
    handles.var_prior_xy.Enable            ='off';
    handles.var_prior_z.Enable             ='off';
    
    handles.pushbutton8.String             = 'Start';
    
    handles.Generate_Synthetic_signal.Enable = 'off';
    handles.Measure_Background.Enable      ='off';
    handles.Import_signal.Enable           ='off';
    
    handles.edit35.Enable                  ='off';
    handles.edit36.Enable                  ='off';
    handles.edit39.Enable                  ='off';
    handles.edit40.Enable                  ='off';
    handles.edit41.Enable                  ='off';
    handles.edit46.Enable                  ='off';
    handles.edit47.Enable                  ='off';
    handles.edit48.Enable                  ='off';
    handles.edit52.Enable                  ='off';
    handles.edit53.Enable                  ='off';
    
    handles.pushbutton8.Enable             ='off';
    handles.pushbutton10.Enable            ='off';
    handles.pushbutton11.Enable            ='off';
    handles.pushbutton16.Enable            ='off';
    handles.pushbutton17.Enable            ='off';
    handles.pushbutton18.Enable            ='off';
    handles.pushbutton19.Enable            ='off';
    handles.pushbutton20.Enable            ='off';
    handles.pushbutton21.Enable            ='off';
    handles.pushbutton22.Enable            ='off';
    handles.pushbutton23.Enable            ='off';
    handles.pushbutton24.Enable            ='off';
    handles.pushbutton25.Enable            ='off';
    handles.pushbutton26.Enable            ='off';
    handles.pushbutton28.Enable            ='off';
    handles.pushbutton29.Enable            ='off';
    handles.pushbutton30.Enable            ='off';
    handles.pushbutton31.Enable            ='off';
    handles.pushbutton32.Enable            ='off';
    handles.pushbutton33.Enable            ='off';
    
    handles.edit44.Enable                  ='off';
    handles.checkbox1.Enable               ='off';
    handles.checkbox2.Enable               ='off';
    handles.checkbox3.Enable               ='off';
    handles.checkbox4.Enable               ='off';
    handles.checkbox7.Enable               ='off';
    handles.checkbox8.Enable               ='off';

    handles.pushbutton8.Enable             = 'off';

     load('results.mat');

     
     
     len = length(Data.Trace_partial)                                                     ;
     for k=1:len
         sign_siz(k)   = length(Data.Trace_partial{k})                                     ; 
         Data.x{k}     = zeros(3*str2num(handles.Number_particles.String),sign_siz(k))+.1  ;
     end
     
    
% Set the automaic adaptivity of the MCMC chain     
     if  sum(handles.pushbutton23.String)  == sum('Automatic (ON)')
         Automatic=1;
     else
         Automatic=0;
     end
     
     
% Set the Filter type to Extended Kalman Filter(EKF) & Unscented Kalman Filter(UKF)
     if  sum(handles.pushbutton22.String)==sum('Estimate background emission rate (OFF)')
         Filter_typ=1;
     else
         Filter_typ=2;
     end

% Set the PSF function (Gaussina, Lorentzian or Dual focus FCS)
     if      sum(handles.pushbutton28.String)==sum('PSF (Gaussian)')          
             PSF_Func=1;     
     elseif  sum(handles.pushbutton28.String)==sum('PSF (Lorentzian)')            
             PSF_Func=2;        
     elseif  sum(handles.pushbutton28.String)==sum('PSF (Cylindrical)') 
             PSF_Func=3;
     end
     
     
     
     
     
     
%%%%%%%%%%%%%%%%%%%%%%%%%%%% Initial vslues %%%%%%%%%%%%%%%%%%%%%%%%%%%%%%%
     
     Data.D     = 1/gamrnd(str2num(handles.D_alpha_prior.String) ,str2num(handles.D_beta_prior.String) )   ;
     Data.mu    = gamrnd(str2num(handles.mu_alpha.String) ,str2num(handles.mu_beta.String) )   ;
    
     
     if  sum(handles.pushbutton22.String)==sum('Estimate background emission rate (OFF)')
         Data.mu_back     = str2num(handles.mu_back.String)                                                ;
         handles.edit46.String = num2str(str2num(handles.mu_back.String)*10*sum(sign_siz)) ;
         handles.edit47.String = num2str(1/(10*sum(sign_siz))) ;
     else
         Data.mu_back     = gamrnd(str2num(handles.edit46.String) ,str2num(handles.edit47.String) )     ;
     end
     
     
     Data.b           = zeros(len,str2num(handles.Number_particles.String),1)                              ; 
     Data.q           =zeros(len,str2num(handles.Number_particles.String),1);
     Data.concentration=zeros(len, 1000 ,max(sign_siz),length(str2num(handles.concen_radious.String)))     ;

     
    [ Data.D             , Data.mu                 , Data.mu_back       , Data.b  , ...
      Data.concentration , Data.acceptance_rate_mu , Data.acceptance_b  , Data.x  , ...
      Data.q         ] = Gibbs_sampler(...
      ...
      ...
      Data.Trace_partial                        ,...
      Data.Time_partial                         ,...
      str2num(handles.D_alpha_prior.String)     ,...
      str2num(handles.D_beta_prior.String)      ,...
      str2num(handles.edit41.String)            ,...
      str2num(handles.Number_particles.String)  ,...
      str2num(handles.edit39.String)            ,...
      str2num(handles.wxy.String)               ,...
      str2num(handles.wz.String)                ,...
      str2num(handles.edit40.String)            ,...
      str2num(handles.mu_alpha.String)          ,...
      str2num(handles.mu_beta.String)           ,...
      str2num(handles.edit48.String)            ,...
      str2num(handles.edit46.String)            ,...
      str2num(handles.edit47.String)            ,...
      str2num(handles.alpha_load.String)        ,...
      str2num(handles.beta_load.String)         ,...
      str2num(handles.mu_prior_xy.String)       ,...
      str2num(handles.mu_prior_z.String)        ,...
      str2num(handles.var_prior_xy.String)      ,...
      str2num(handles.var_prior_z.String)       ,...
      Data.mu_back                              ,...
      20                                        ,...
      str2num(handles.concen_radious.String)    ,...
      Data.x                                    ,...
      handles                                   ,...
      Data.D                                    ,...
      Data.mu                                   ,...
      Data.b                                    ,...
      Data.q                                    ,...
      Data.concentration                        ,...
      Automatic                                 ,...
      PSF_Func                                  ,...
      Filter_typ                                ,...
      str2num(handles.edit52.String)           );   
         
  
      
     if sum(handles.pushbutton22.String)==sum('Estimate background emission rate (OFF)')
      
         plot(handles.axes16,1:1:length(Data.mu),Data.mu)
         histogram(handles.axes20,Data.mu,'Normalization','pdf','Orientation','horizontal')
     else
         plot(handles.axes16,1:1:length(Data.mu),Data.mu,1:1:length(Data.mu),Data.mu_back)
      
         histogram(handles.axes20,Data.mu,'Normalization','pdf','Orientation','horizontal')
         hold (handles.axes20,'on')
         histogram(handles.axes20,Data.mu_back,'Normalization','pdf','Orientation','horizontal')
         hold (handles.axes20,'off')
         
         handles.edit46.Enable                  ='on';
         handles.edit47.Enable                  ='on';
         handles.edit48.Enable                  ='on';
         handles.checkbox7.Enable               ='on';
         handles.checkbox7.Value                = 1  ;
     end
     
     
     
     
     
    save('results' , 'Data'   )
             
    plot(handles.axes8,1:1:length(Data.mu),Data.D)
    xlabel(handles.axes8,'Iteration')
    ylabel(handles.axes8,'log_{10} Diff. coeff. (\mum^2/s)')
    xlim(handles.axes8,[0 length(Data.mu)])
    
    
    
    for k=1:len
        plot(handles.axes15,1:1:length(Data.mu),sum(reshape(Data.b(k,:,:),size(Data.b,2),size(Data.b,3)),1))
        hold (handles.axes15,'on')
    end
    hold (handles.axes15,'off')
    
    xlabel(handles.axes15,'Iteration')
    ylabel(handles.axes15,'Number of active molecules')
    ylim(handles.axes15,[0 str2num(handles.Number_particles.String)])
    xlim(handles.axes15,[0 length(Data.mu)])
    

    
    xlabel(handles.axes16,'Iteration')
    ylabel(handles.axes16,{'Molecule emission rate';'(photons/s)'})
    xlim(handles.axes16,[0 length(Data.mu)])
    
    
    
    histogram(handles.axes19,Data.D,'Normalization','pdf','Orientation','horizontal')
    handles.axes19.XAxisLocation = 'top';
    ylim(handles.axes19,[0 max(Data.D)])
    handles.axes19.YAxisLocation = 'right';
    set(handles.axes19,'yscale','log') 
    
    
    for k=1:len
        histogram(handles.axes18,sum(reshape(Data.b(k,:,:),size(Data.b,2),size(Data.b,3)),1),...
                  'Normalization','pdf','Orientation','horizontal')
        hold (handles.axes18,'on')
    end
    hold (handles.axes18,'off')
    handles.axes18.YTick = [];
    handles.axes18.XAxisLocation = 'top';
    ylim(handles.axes18,[0 str2num(handles.Number_particles.String)])
    

    handles.axes20.YTick = [];
    handles.axes20.XAxisLocation = 'top';
    ylim(handles.axes20,[0 max(Data.mu)])
    
    drawnow
    
    handles.edit44.Enable                        ='on';
    handles.checkbox1.Enable                     ='on';
    handles.checkbox2.Enable                     ='on';
    handles.checkbox3.Enable                     ='on';
    handles.checkbox4.Enable                     ='on';
    handles.checkbox8.Enable                     ='on';
    
    handles.Generate_Synthetic_signal.Enable     ='on';
    handles.Measure_Background.Enable            ='on';
    handles.Import_signal.Enable                 ='on';
    
    handles.edit35.Enable                        ='on';
    handles.edit36.Enable                        ='on';
    handles.edit40.Enable                        ='on';
    handles.edit53.Enable                        ='on';
    
    handles.pushbutton8.Enable                   ='on';
    handles.pushbutton10.Enable                  ='on';
    handles.pushbutton11.Enable                  ='on';
    handles.pushbutton16.Enable                  ='on';
    handles.pushbutton18.Enable                  ='on';
    handles.pushbutton19.Enable                  ='on';
    handles.pushbutton21.Enable                  ='on';
    handles.pushbutton22.Enable                  ='on';
    handles.pushbutton23.Enable                  ='on';
    handles.pushbutton24.Enable                  ='on';
    handles.pushbutton25.Enable                  ='on';
    handles.pushbutton26.Enable                  ='on';
    handles.pushbutton29.Enable                  ='on';
    handles.pushbutton30.Enable                  ='on';
    handles.pushbutton31.Enable                  ='on';
    handles.pushbutton32.Enable                  ='on';
    handles.pushbutton33.Enable                  ='on';
    
    if  sum(handles.pushbutton23.String) == sum('Automatic (OFF)')
        handles.pushbutton20.Enable              ='on';
        handles.pushbutton17.Enable              ='on';
        handles.edit39.Enable                    ='on';
        handles.edit41.Enable                    ='on';
    else
        handles.edit52.Enable                    ='on';
    end
    

    handles.D_alpha_prior.Enable                 ='on';
    handles.D_beta_prior.Enable                  ='on';
    handles.mu_alpha.Enable                      ='on';
    handles.mu_beta.Enable                       ='on';
    handles.Number_particles.Enable              ='on';
    handles.alpha_load.Enable                    ='on';
    handles.beta_load.Enable                     ='on';
    handles.concen_radious.Enable                ='on';
    handles.mu_prior_xy.Enable                   ='on';
    handles.mu_prior_z.Enable                    ='on';
    handles.var_prior_xy.Enable                  ='on';
    handles.var_prior_z.Enable                   ='on';

function edit41_Callback(hObject, eventdata, handles)
% hObject    handle to edit41 (see GCBO)
% eventdata  reserved - to be defined in a future version of MATLAB
% handles    structure with handles and user data (see GUIDATA)

% Hints: get(hObject,'String') returns contents of edit41 as text
%        str2double(get(hObject,'String')) returns contents of edit41 as a double


% --- Executes during object creation, after setting all properties.
function edit41_CreateFcn(hObject, eventdata, handles)
% hObject    handle to edit41 (see GCBO)
% eventdata  reserved - to be defined in a future version of MATLAB
% handles    empty - handles not created until after all CreateFcns called

% Hint: edit controls usually have a white background on Windows.
%       See ISPC and COMPUTER.
if ispc && isequal(get(hObject,'BackgroundColor'), get(0,'defaultUicontrolBackgroundColor'))
    set(hObject,'BackgroundColor','white');
end


function edit40_Callback(hObject, eventdata, handles)
% hObject    handle to edit40 (see GCBO)
% eventdata  reserved - to be defined in a future version of MATLAB
% handles    structure with handles and user data (see GUIDATA)

% Hints: get(hObject,'String') returns contents of edit40 as text
%        str2double(get(hObject,'String')) returns contents of edit40 as a double


% --- Executes during object creation, after setting all properties.
function edit40_CreateFcn(hObject, eventdata, handles)
% hObject    handle to edit40 (see GCBO)
% eventdata  reserved - to be defined in a future version of MATLAB
% handles    empty - handles not created until after all CreateFcns called

% Hint: edit controls usually have a white background on Windows.
%       See ISPC and COMPUTER.
if ispc && isequal(get(hObject,'BackgroundColor'), get(0,'defaultUicontrolBackgroundColor'))
    set(hObject,'BackgroundColor','white');
end


function edit39_Callback(hObject, eventdata, handles)
% hObject    handle to edit39 (see GCBO)
% eventdata  reserved - to be defined in a future version of MATLAB
% handles    structure with handles and user data (see GUIDATA)

% Hints: get(hObject,'String') returns contents of edit39 as text
%        str2double(get(hObject,'String')) returns contents of edit39 as a double


% --- Executes during object creation, after setting all properties.
function edit39_CreateFcn(hObject, eventdata, handles)
% hObject    handle to edit39 (see GCBO)
% eventdata  reserved - to be defined in a future version of MATLAB
% handles    empty - handles not created until after all CreateFcns called

% Hint: edit controls usually have a white background on Windows.
%       See ISPC and COMPUTER.
if ispc && isequal(get(hObject,'BackgroundColor'), get(0,'defaultUicontrolBackgroundColor'))
    set(hObject,'BackgroundColor','white');
end


% --- Executes on button press in pushbutton16.
function pushbutton16_Callback(hObject, eventdata, handles)
% hObject    handle to pushbutton16 (see GCBO)
% eventdata  reserved - to be defined in a future version of MATLAB
% handles    structure with handles and user data (see GUIDATA)
    load('results.mat');
    
    handles.edit44.Enable                        ='off';
    handles.checkbox1.Enable                     ='off';
    handles.checkbox2.Enable                     ='off';
    handles.checkbox3.Enable                     ='off';
    handles.checkbox4.Enable                     ='off';
    handles.checkbox7.Enable                     ='off';
    handles.checkbox8.Enable                     ='off';
    handles.pushbutton11.Enable                  ='off';
    
    current_location=cd;
    
    mkdir Saved_Results
    cd Saved_Results
    
    MAXX=floor(length(Data.D)*str2num(handles.edit44.String));
    
    
    choice = questdlg({'In what format do you want to save the results?'},'SAVE RESULTS','Text(.txt)','Matlab(.mat)','Cancel');
    

    if  strcmp(choice,'Text(.txt)')
        formatSpec = ' %4.2f ';
        if  handles.checkbox1.Value==1
            fid1 = fopen( 'Diffusion_coefficient.txt', 'wt' );        
            fprintf(fid1,formatSpec,Data.D(MAXX:end));
        end
        if  handles.checkbox2.Value==1
            fid2 = fopen( 'Emission_rate.txt', 'wt' );
            fprintf(fid2,formatSpec,Data.mu(MAXX:end));
        end
        if  handles.checkbox3.Value==1
            fid3 = fopen( 'Number_of_Acive_molecules (loads).txt', 'wt' );
            for i=1:size(Data.b,1)
                fprintf(fid3,['\n','Trace',num2str(i),'\n']);
                fprintf(fid3,'%g\t',sum(Data.b(i,:,MAXX:end)));
            end
        end

        if  handles.checkbox4.Value==1
            concenn50=[];
            fid4 = fopen( 'Concentration.txt', 'wt' );
            for i=1:size(Data.b,1)
                fprintf(fid4,['\n','Trace',num2str(i),'\n']);
                
                for l=1:length(str2num(handles.concen_radious.String))
                    concenn50=[];
                    concenn75=[];
                    concenn25=[];
                    for k = 1 : length(Data.Trace_partial{i})
                        concenn50(1,k)=quantile(reshape(Data.concentration(i,:,k,l),1,1000),0.50)    ;
                        concenn25(1,k)=quantile(reshape(Data.concentration(i,:,k,l),1,1000),0.025)    ;
                        concenn75(1,k)=quantile(reshape(Data.concentration(i,:,k,l),1,1000),0.975)    ;
                    end
                    fprintf(fid4,['\n','Nomalized distance ',num2str(l),'\n','Median of posterior                ']);
                    fprintf(fid4,'%g\t',concenn50);
                    fprintf(fid4,['\n','25 percent quantile of posterior   ']);
                    fprintf(fid4,'%g\t',concenn25);
                    fprintf(fid4,['\n','75 percent quantile of posterior   ']);
                    fprintf(fid4,'%g\t',concenn75);
                    fprintf(fid4,'\n');
                end
            end
        end
        
        if  handles.checkbox7.Value==1
            fid5 = fopen( 'Diffusion_coefficient.txt', 'wt' );        
            fprintf(fid5,formatSpec,Data.mu_back(MAXX:end));
            
        end
      

        fid5 = fopen('Signal.txt', 'wt' );
        for i = 1 : length(Data.Trace_partial)
            fprintf(fid5,['\n','Trace',num2str(i),'\n']);
            fprintf(fid5,'%g\t',Data.Trace_partial{i});
        end
        
%         fprintf(fid5,formatSpec,Data.Trace_partial);
%         
        fid6 = fopen('Time.txt', 'wt' );
        for i = 1 : length(Data.Time_partial)
            fprintf(fid6,['\n','Trace',num2str(i),'\n']);
            fprintf(fid6,'%g\t',Data.Time_partial{i});
        end
        
    else
        if  strcmp(choice,'Matlab(.mat)')
            if  handles.checkbox1.Value==1
                Diffusion=Data.D(MAXX:end);
            else
                Diffusion=[];
            end
            if  handles.checkbox2.Value==1
                Emission=Data.mu(MAXX:end);
            else
                Emission=[];
            end
            if  handles.checkbox7.Value==1
                Background_Emission=Data.mu_back(MAXX:end);
            else
                Background_Emission=[];
            end
            if  handles.checkbox3.Value==1
                for i=1:size(Data.b,1)
                loads(i,:)=sum(Data.b(i,:,MAXX:end));
                end
            else
                loads=[];
            end
            if  handles.checkbox4.Value==1
                Concentration=Data.concentration;
            else
                Concentration=[];
            end
            Signal=Data.Trace_partial;
            Time=Data.Time_partial;
            
            if sum(handles.pushbutton22.String)==sum('Estimate background emission rate (OFF)')
               save('Results','Diffusion','Emission','loads','Concentration','Signal','Time')
            else
               save('Results','Diffusion','Emission','Background_Emission','loads','Concentration','Signal','Time')
            end
        end
    end
    

    
    drawnow
    handles.edit44.Enable                        ='on';
    handles.checkbox1.Enable                     ='on';
    handles.checkbox2.Enable                     ='on';
    handles.checkbox3.Enable                     ='on';
    handles.checkbox4.Enable                     ='on';
    handles.checkbox8.Enable                     ='on';
    handles.pushbutton11.Enable                  ='on';
    
    if sum(handles.pushbutton22.String)=='Estimate background emission rate (ON)'
            handles.checkbox7.Enable             ='on';
    end
    
    cd(current_location)
    clear all
    clc
    
   


% --- Executes on button press in pushbutton17.
function pushbutton17_Callback(hObject, eventdata, handles)
% hObject    handle to pushbutton17 (see GCBO)
% eventdata  reserved - to be defined in a future version of MATLAB
% handles    structure with handles and user data (see GUIDATA)


    handles.Generate_Synthetic_signal.Enable    =  'off';
    handles.Measure_Background.Enable           =  'off';
    handles.Import_signal.Enable                =  'off';
    
    
    
    handles.pushbutton8.Enable                  =  'off';
    handles.pushbutton10.Enable                 =  'off';
    handles.pushbutton11.Enable                 =  'off';
    handles.pushbutton16.Enable                 =  'off';
    handles.pushbutton17.Enable                 =  'off';
    handles.pushbutton19.Enable                 =  'off';
    handles.pushbutton20.Enable                 =  'off';
    handles.pushbutton22.Enable                 =  'off';
    handles.pushbutton23.Enable                 =  'off';
    handles.pushbutton28.Enable                 =  'off';
    handles.pushbutton29.Enable                 =  'off';
    handles.pushbutton32.Enable                 =  'off';
    handles.pushbutton33.Enable                 =  'off';
    
     load('results.mat');
     
     len = length(Data.Trace_partial)                                     ;
     for k=1:len
         sign_siz(k)   = length(Data.Trace_partial{k})                    ; 
         Data.x{k}     = zeros(3*str2num(handles.Number_particles.String),sign_siz(k))+.1  ;
     end
     
     
     iter_b_mu   =     20         ;

%%%%%%%%%%%%%%%%%%%%%%%%%%%%%%%%% Initial vslues %%%%%%%%%%%%%%%%%%%%%%%%%%%%%%%%%%%
     Data.D       = 1/gamrnd(str2num(handles.D_alpha_prior.String) ,str2num(handles.D_beta_prior.String) ) ;
     Data.mu      = gamrnd(str2num(handles.mu_alpha.String) ,str2num(handles.mu_beta.String) )             ;
     Data.b       = zeros(len,str2num(handles.Number_particles.String),1)                                ;
     
     
     
%      Data.q             = zeros(len,1)                              ;   
     Data.q           =zeros(len,str2num(handles.Number_particles.String),1);          
     Data.concentration =zeros(len , 1000 ,max(sign_siz) , length(str2num(handles.concen_radious.String)) ) ;

   
     if      sum(handles.pushbutton28.String)==sum('PSF (Gaussian)')          
             PSF_Func=1;     
     elseif  sum(handles.pushbutton28.String)==sum('PSF (Lorentzian)')            
             PSF_Func=2;        
     elseif  sum(handles.pushbutton28.String)==sum('PSF (Cylindrical)')
             PSF_Func=3;
     end
     
     if  sum(handles.pushbutton22.String)==sum('Estimate background emission rate (OFF)')
         Filter_typ=1;
         Data.mu_back     = str2num(handles.mu_back.String)                                 ;
         handles.edit46.String = num2str(str2num(handles.mu_back.String)*10*sum(sign_siz)) ;
         handles.edit47.String = num2str(1/(10*sum(sign_siz))) ;
     else
         Filter_typ=2;
         Data.mu_back     = gamrnd(str2num(handles.edit46.String) ,str2num(handles.edit47.String) )     ;
     end
      
     
     
     tic
       [ ~ , ~ , ~ , ~ , ~ , ~ , ~ , ~ , ~ ] = Gibbs_sampler(...
      ...
      ...
     Data.Trace_partial                        ,...
     Data.Time_partial                         ,...
     str2num(handles.D_alpha_prior.String)     ,...
     str2num(handles.D_beta_prior.String)      ,...
     10                                        ,...
     str2num(handles.Number_particles.String)  ,...
     20                                        ,...
     str2num(handles.wxy.String)               ,...
     str2num(handles.wz.String)                ,...
     1000                                      ,...
     str2num(handles.mu_alpha.String)          ,...
     str2num(handles.mu_beta.String)           ,...
     1000                                      ,...
     str2num(handles.edit46.String)            ,...
     str2num(handles.edit47.String)            ,...
     str2num(handles.alpha_load.String)        ,...
     str2num(handles.beta_load.String)         ,...
     str2num(handles.mu_prior_xy.String)       ,...
     str2num(handles.mu_prior_z.String)        ,...
     str2num(handles.var_prior_xy.String)      ,...
     str2num(handles.var_prior_z.String)       ,...
     Data.mu_back                              ,...
     iter_b_mu                                 ,...
     str2num(handles.concen_radious.String)    ,...
     Data.x                                    ,...
     handles                                   ,...
     Data.D                                    ,...
     Data.mu                                   ,...
     Data.b                                    ,...
     Data.q                                    ,...
     Data.concentration                        ,...
     0                                         ,...
     PSF_Func                                  ,...
     Filter_typ                                ,...
     str2num(handles.edit52.String)           );

     timmer=toc;

     
     Data.timmer=(timmer/19);
    
    save('results','Data');

    timmer=floor(str2num(handles.edit39.String)*(timmer/19));
    
    if  timmer>=3600
        timmer=floor(timmer/3600);
        estimate_time=['Start (' num2str(timmer) 'hr(s))'];
    else
        if timmer>=60
           timmer=floor(timmer/60);
           estimate_time=['Start (' num2str(timmer) 'min)'];
        else
            estimate_time=['Start (' num2str(timmer) 's)'];
        end
    end
   
    
    handles.Generate_Synthetic_signal.Enable    =  'on';
    handles.Measure_Background.Enable           =  'on';
    handles.Import_signal.Enable                =  'on';
    
    handles.pushbutton8.Enable                  =  'on';
    handles.pushbutton10.Enable                 =  'on';
    handles.pushbutton17.Enable                 =  'on';
    handles.pushbutton19.Enable                 =  'on';
    handles.pushbutton22.Enable                 =  'on';
    handles.pushbutton23.Enable                 =  'on';
    handles.pushbutton28.Enable                 =  'on';
    handles.pushbutton29.Enable                 =  'on';
    handles.pushbutton32.Enable                 =  'on';
    handles.pushbutton33.Enable                 =  'on';
    
    handles.pushbutton8.String                  = estimate_time;

    


    cla(handles.axes8)
    cla(handles.axes15)
    cla(handles.axes16)
    cla(handles.axes18)
    cla(handles.axes19)
    cla(handles.axes20)


% --- Executes on button press in pushbutton18.
function pushbutton18_Callback(hObject, eventdata, handles)
% hObject    handle to pushbutton18 (see GCBO)
% eventdata  reserved - to be defined in a future version of MATLAB
% handles    structure with handles and user data (see GUIDATA)

choice = questdlg({'All the files in the directory will be removed. Please save them if you did not!'...
                   },'Are you sure?','Yes','No','Yes');
    
if  strcmp(choice,'Yes')

     S = dir('*.mat');                   % Collect directories
     N = {S.name};                       % Collect Names of the text files
     for i=1:length(N)                   % A For loop for read data
         delete(N{i});           % Import data from the file
     end


     clear all;
     clc;
     close all;  
end


% --- Executes on button press in pushbutton19.
function pushbutton19_Callback(hObject, eventdata, handles)
% hObject    handle to pushbutton19 (see GCBO)
% eventdata  reserved - to be defined in a future version of MATLAB
% handles    structure with handles and user data (see GUIDATA)

 clc;
    S = dir('*.mat');                   % Collect directories
    N = {S.name};                       % Collect Names of the text files
    for i=1:length(N)                   % A For loop for read data
        delete(N{i});           % Import data from the file
    end
    clear('S','N')
 
    handles.Refractive_index.String        = '-';
    handles.Delta_t.String                 = '-';
    handles.Laser_Wavelength.String        = '-';
    handles.Objective_magnification.String = '-';
    handles.wxy.String                     = '-';
    handles.wz.String                      = '-';
    handles.mu_back.String                 = '-';
    handles.Diff_synth.String              = '-';
    handles.mu_synth.String                = '-';
    handles.Num_synth.String               = '-';
    
    handles.edit35.String                  = '0';
    handles.edit36.String                  = '0';
    handles.edit39.String                  = '1000';
    handles.edit40.String                  = '1000';
    handles.edit41.String                  = '10';
    handles.edit48.String                  = '1000';
    handles.edit52.String                  = '0.1';
    
    handles.D_alpha_prior.Enable           ='off';
    handles.D_beta_prior.Enable            ='off';
    handles.mu_alpha.Enable                ='off';
    handles.mu_beta.Enable                 ='off';
    handles.Number_particles.Enable        ='off';
    handles.alpha_load.Enable              ='off';
    handles.beta_load.Enable               ='off';
    handles.concen_radious.Enable          ='off';
    handles.mu_prior_xy.Enable             ='off';
    handles.mu_prior_z.Enable              ='off';
    handles.var_prior_xy.Enable            ='off';
    handles.var_prior_z.Enable             ='off';
    
    handles.pushbutton8.Enable             ='off';
    handles.pushbutton10.Enable            ='off';
    handles.pushbutton11.Enable            ='off';
    handles.pushbutton17.Enable            ='off';
    handles.pushbutton16.Enable            ='off';
    handles.pushbutton19.Enable            ='off';
    handles.pushbutton20.Enable            ='off';
    handles.pushbutton22.Enable            ='off';
    handles.pushbutton23.Enable            ='off';
    handles.pushbutton24.Enable            ='off';
    handles.pushbutton25.Enable            ='off';
    handles.pushbutton26.Enable            ='off';
    handles.pushbutton28.Enable            ='off';
    handles.pushbutton29.Enable            ='off';
    handles.pushbutton30.Enable            ='off';
    handles.pushbutton31.Enable            ='off';
    
    handles.text36.Enable                  ='off';
    handles.text37.Enable                  ='off';
    
    handles.uitable5.Data                  =[];
    handles.uitable5.Enable                ='off';
    
    handles.edit35.Enable                  ='off';
    handles.edit36.Enable                  ='off';
    handles.edit39.Enable                  ='off';
    handles.edit40.Enable                  ='off';
    handles.edit41.Enable                  ='off';
    handles.edit44.Enable                  ='off';
    handles.edit46.Enable                  ='off';
    handles.edit47.Enable                  ='off';
    handles.edit48.Enable                  ='off';
    handles.edit52.Enable                  ='off';
    handles.edit53.Enable                  ='off';
    
    handles.checkbox1.Enable               ='off';
    handles.checkbox2.Enable               ='off';
    handles.checkbox3.Enable               ='off';
    handles.checkbox4.Enable               ='off';
    handles.checkbox7.Enable               ='off';
    handles.checkbox8.Enable               ='off';
    
    handles.Generate_Synthetic_signal.Enable   ='on';
    handles.Measure_Background.Enable          ='on';
    handles.Import_signal.Enable               ='on';
    
    cla(handles.Original_signal)
    cla(handles.axes7)
    cla(handles.axes8)
    cla(handles.axes15)
    cla(handles.axes16)
    cla(handles.axes18)
    cla(handles.axes19)
    cla(handles.axes20)
    
    
    handles.pushbutton22.String ='Estimate background emission rate (OFF)';
    handles.pushbutton8.String  ='Start';
    
    clear all;
 
 
 
 
 


% --- Executes on button press in pushbutton20.
function pushbutton20_Callback(hObject, eventdata, handles)
% hObject    handle to pushbutton20 (see GCBO)
% eventdata  reserved - to be defined in a future version of MATLAB
% handles    structure with handles and user data (see GUIDATA)


    handles.pushbutton8.Enable                   ='off';
    handles.pushbutton10.Enable                  ='off';
    handles.pushbutton11.Enable                  ='off';
    handles.pushbutton16.Enable                  ='off';
    handles.pushbutton17.Enable                  ='off';
    handles.pushbutton20.Enable                  ='off';
    handles.pushbutton22.Enable                  ='off';
    handles.pushbutton23.Enable                  ='off';
    handles.pushbutton29.Enable                  ='off';
    handles.pushbutton30.Enable                  ='off';
    handles.pushbutton31.Enable                  ='off';
    handles.pushbutton32.Enable                  ='off';
    handles.pushbutton33.Enable                  ='off';
    
    handles.checkbox1.Enable                     ='off';
    handles.checkbox2.Enable                     ='off';
    handles.checkbox3.Enable                     ='off';
    handles.checkbox4.Enable                     ='off';
    handles.checkbox7.Enable                     ='off';
    handles.checkbox8.Enable                     ='off';

    handles.Generate_Synthetic_signal.Enable     ='off';
    handles.Measure_Background.Enable            ='off';
    handles.Import_signal.Enable                 ='off';
    
    handles.edit35.Enable                        ='off';
    handles.edit36.Enable                        ='off';
    handles.edit39.Enable                        ='off';
    handles.edit40.Enable                        ='off';
    handles.edit41.Enable                        ='off';
    handles.edit44.Enable                        ='off';
    handles.edit46.Enable                        ='off';
    handles.edit47.Enable                        ='off';
    handles.edit48.Enable                        ='off';
    handles.edit53.Enable                        ='off';

    handles.D_alpha_prior.Enable                 ='off';
    handles.D_beta_prior.Enable                  ='off';
    handles.mu_alpha.Enable                      ='off';
    handles.mu_beta.Enable                       ='off';
    handles.Number_particles.Enable              ='off';
    handles.alpha_load.Enable                    ='off';
    handles.beta_load.Enable                     ='off';
    handles.concen_radious.Enable                ='off';
    handles.mu_prior_xy.Enable                   ='off';
    handles.mu_prior_z.Enable                    ='off';
    handles.var_prior_xy.Enable                  ='off';
    handles.var_prior_z.Enable                   ='off';
    
    
     
    load('results.mat');
   
answer={};

    opts.Interpreter = 'tex';
    answer = inputdlg({'Number of extra iterations'},...
                        'Please consider the Running time!',...
                        [1 60],{num2str(length(Data.D)-1)},opts);
                    
  if length(answer)==1
    while isempty(str2num(answer{1})) || str2num(answer{1})==0
           opts.Interpreter = 'tex';
           answer = inputdlg({'Number of extra iterations'},...
                              'Please choose a number greater than 0!',...
                              [1 60],{num2str(length(Data.D)-1)},opts);
    end
    
    
    
    handles.edit39.String = num2str(length(Data.D)-1+str2num(answer{1}));

    
    if      sum(handles.pushbutton28.String)==sum('PSF (Gaussian)')          
             PSF_Func=1;     
     elseif  sum(handles.pushbutton28.String)==sum('PSF (Lorentzian)')            
             PSF_Func=2;        
     elseif  sum(handles.pushbutton28.String)==sum('PSF (Cylindrical)')
             PSF_Func=3;
    end
     
     
    len = length(Data.Trace_partial)                                     ;
    for k=1:len
         sign_siz(k)   = length(Data.Trace_partial{k})                    ;
    end
    
    if  sum(handles.pushbutton22.String)==sum('Estimate background emission rate (OFF)')
         Filter_typ=1;
         handles.edit46.String = num2str(str2num(handles.mu_back.String)*10*sum(sign_siz)) ;
         handles.edit47.String = num2str(1/(10*sum(sign_siz))) ;
    else
         Filter_typ=2;
    end

     
    
        
      [ Data.D              , Data.mu                  , Data.mu_back      , Data.b , ...
        Data.concentration  , Data.acceptance_rate_mu  , Data.acceptance_b , Data.x , ...
        Data.q           ] =  Gibbs_sampler(...
      ...
      ...
      Data.Trace_partial                        ,...
      Data.Time_partial                         ,...
      str2num(handles.D_alpha_prior.String)     ,...
      str2num(handles.D_beta_prior.String)      ,...
      str2num(handles.edit41.String)            ,...
      str2num(handles.Number_particles.String)  ,...
      str2num(answer{1})                        ,...
      str2num(handles.wxy.String)               ,...
      str2num(handles.wz.String)                ,...
      str2num(handles.edit40.String)            ,...
      str2num(handles.mu_alpha.String)          ,...
      str2num(handles.mu_beta.String)           ,...
      str2num(handles.edit48.String)            ,...
      str2num(handles.edit46.String)            ,...
      str2num(handles.edit47.String)            ,...
      str2num(handles.alpha_load.String)        ,...
      str2num(handles.beta_load.String)         ,...
      str2num(handles.mu_prior_xy.String)       ,...
      str2num(handles.mu_prior_z.String)        ,...
      str2num(handles.var_prior_xy.String)      ,...
      str2num(handles.var_prior_z.String)       ,...
      Data.mu_back                              ,...
      20                                        ,...
      str2num(handles.concen_radious.String)    ,...
      Data.x                                    ,...
      handles                                   ,...
      Data.D                                    ,...
      Data.mu                                   ,...
      Data.b                                    ,...
      Data.q                                    ,...
      Data.concentration                        ,...
      0                                         ,...
      PSF_Func                                  ,...
      Filter_typ                                ,...
      str2num(handles.edit52.String)           );
  
  
  
     if  sum(handles.pushbutton22.String)==sum('Estimate background emission rate (OFF)')
         histogram(handles.axes20,Data.mu,'Normalization','pdf','Orientation','horizontal')
         plot(handles.axes16,1:1:str2num(handles.edit39.String)+1,Data.mu)
     else
         plot(handles.axes16,1:1:str2num(handles.edit39.String)+1,Data.mu,1:1:str2num(handles.edit39.String)+1,Data.mu_back)
         histogram(handles.axes20,Data.mu,'Normalization','pdf','Orientation','horizontal')
         hold (handles.axes20,'on')
         histogram(handles.axes20,Data.mu_back,'Normalization','pdf','Orientation','horizontal')
         hold (handles.axes20,'off')
         handles.checkbox7.Enable                     ='on';
     end
      
  
      
    
    
    save('results' , 'Data' )
                
    plot(handles.axes8,1:1:str2num(handles.edit39.String)+1,Data.D)
    xlabel(handles.axes8,['Iteration'])
    ylabel(handles.axes8,'log_{10} Diff. coeff. (\mum^2/s)')
    xlim(handles.axes8,[0 str2num(handles.edit39.String)+1])
    ylim(handles.axes8,[0 max(Data.D)])
    set(handles.axes8,'yscale','log') 
    
    for k=1:length(Data.Trace_partial)
        plot(handles.axes15,1:1:str2num(handles.edit39.String)+1,sum(reshape(Data.b(k,:,:),size(Data.b,2),size(Data.b,3)),1))
        hold (handles.axes15,'on')
    end
    hold (handles.axes15,'off')
    xlabel(handles.axes15,['Iteration'])
    ylabel(handles.axes15,'Number of active molecules')
    ylim(handles.axes15,[0 str2num(handles.Number_particles.String)])
    xlim(handles.axes15,[0 str2num(handles.edit39.String)+1])
    
    xlabel(handles.axes16,'Iteration')
    if  sum(handles.pushbutton22.String)==sum('Estimate background emission rate (OFF)')
        ylabel(handles.axes16,{'Molecule photon emission rate' ;'(photons/s)'})
    else
        ylabel(handles.axes16,{'Molecule and background';'photon emission rates' ;'(photons/s)'})
    end
    xlim(handles.axes16,[0 str2num(handles.edit39.String)+1])
    
    bnd = logspace(log10(min(Data.D))-1,log10(max(Data.D))+1,100);
    
    
    histogram(handles.axes19,Data.D,bnd,'Normalization','pdf','Orientation','horizontal')
    handles.axes19.XAxisLocation = 'top';
    ylim(handles.axes19,[0 max(Data.D)])
    handles.axes19.YTick = [];
    handles.axes19.XTick = [];
    handles.axes19.YAxisLocation = 'right';
    set(handles.axes19,'yscale','log') 
    
    
    for k=1:length(Data.Trace_partial)
        histogram(handles.axes18,sum(reshape(Data.b(k,:,:),size(Data.b,2),size(Data.b,3)),1),'Normalization','Probability','Orientation','horizontal')
        hold (handles.axes18,'on')
    end
    hold (handles.axes18,'off')
    handles.axes18.YTick = [];
    handles.axes18.XTick = [];
    handles.axes18.XAxisLocation = 'top';
    ylim(handles.axes18,[0 str2num(handles.Number_particles.String)])
    
    
    handles.axes20.YTick = [];
    handles.axes20.XTick = [];
    handles.axes20.XAxisLocation = 'top';
    ylim(handles.axes20,[0 max(Data.mu)])
    
    drawnow
    
    handles.pushbutton8.Enable                   ='on';
    handles.pushbutton10.Enable                  ='on';
    handles.pushbutton17.Enable                  ='on';
    handles.pushbutton11.Enable                  ='on';
    handles.pushbutton16.Enable                  ='on';
    handles.pushbutton20.Enable                  ='on';
    handles.pushbutton22.Enable                  ='on';
    
    if sum(handles.pushbutton22.String)==sum('Estimate background emission rate (OFF)')
        handles.edit46.Enable                        ='off';
        handles.edit47.Enable                        ='off';
    else
        handles.edit46.Enable                        ='on';
        handles.edit47.Enable                        ='on';
    end
    handles.pushbutton23.Enable                  ='on';
    handles.pushbutton29.Enable                  ='on';
    handles.pushbutton30.Enable                  ='on';
    handles.pushbutton31.Enable                  ='on';
    handles.pushbutton32.Enable                  ='on';
    handles.pushbutton33.Enable                  ='on';
    
    handles.checkbox1.Enable                     ='on';
    handles.checkbox2.Enable                     ='on';
    handles.checkbox3.Enable                     ='on';
    handles.checkbox4.Enable                     ='on';
    handles.checkbox8.Enable                     ='on';

    handles.Generate_Synthetic_signal.Enable     ='on';
    handles.Measure_Background.Enable            ='on';
    handles.Import_signal.Enable                 ='on';
    
    handles.edit35.Enable                        ='on';
    handles.edit36.Enable                        ='on';
    handles.edit39.Enable                        ='on';
    handles.edit40.Enable                        ='on';
    handles.edit41.Enable                        ='on';
    handles.edit44.Enable                        ='on';
    handles.edit53.Enable                        ='on';

    if sum(handles.pushbutton22.String)==sum('Estimate background emission rate (ON)')
        handles.pushbutton22.Enable              ='on';
        handles.edit46.Enable                    ='on';
        handles.edit47.Enable                    ='on';
        handles.edit48.Enable                    ='on';
    end
    
    handles.D_alpha_prior.Enable                 ='on';
    handles.D_beta_prior.Enable                  ='on';
    handles.mu_alpha.Enable                      ='on';
    handles.mu_beta.Enable                       ='on';
    handles.Number_particles.Enable              ='on';
    handles.alpha_load.Enable                    ='on';
    handles.beta_load.Enable                     ='on';
    handles.concen_radious.Enable                ='on';
    handles.mu_prior_xy.Enable                   ='on';
    handles.mu_prior_z.Enable                    ='on';
    handles.var_prior_xy.Enable                  ='on';
    handles.var_prior_z.Enable                   ='on';
    else
        
    handles.pushbutton8.Enable                   ='on';
    handles.pushbutton10.Enable                  ='on';
    handles.pushbutton17.Enable                  ='on';
    handles.pushbutton11.Enable                  ='on';
    handles.pushbutton16.Enable                  ='on';
    handles.pushbutton20.Enable                  ='on';
    handles.pushbutton22.Enable                  ='on';
    
    if sum(handles.pushbutton22.String)==sum('Estimate background emission rate (OFF)')
        handles.edit46.Enable                        ='off';
        handles.edit47.Enable                        ='off';
    else
        handles.edit46.Enable                        ='on';
        handles.edit47.Enable                        ='on';
    end
    handles.pushbutton23.Enable                  ='on';
    handles.pushbutton29.Enable                  ='on';
    handles.pushbutton30.Enable                  ='on';
    handles.pushbutton31.Enable                  ='on';
    handles.pushbutton32.Enable                  ='on';
    handles.pushbutton33.Enable                  ='on';
    
    handles.checkbox1.Enable                     ='on';
    handles.checkbox2.Enable                     ='on';
    handles.checkbox3.Enable                     ='on';
    handles.checkbox4.Enable                     ='on';
    handles.checkbox8.Enable                     ='on';

    handles.Generate_Synthetic_signal.Enable     ='on';
    handles.Measure_Background.Enable            ='on';
    handles.Import_signal.Enable                 ='on';
    
    handles.edit35.Enable                        ='on';
    handles.edit36.Enable                        ='on';
    handles.edit39.Enable                        ='on';
    handles.edit40.Enable                        ='on';
    handles.edit41.Enable                        ='on';
    handles.edit44.Enable                        ='on';
    handles.edit53.Enable                        ='on';

    if sum(handles.pushbutton22.String)==sum('Estimate background emission rate (ON)')
        handles.pushbutton22.Enable              ='on';
        handles.edit46.Enable                    ='on';
        handles.edit47.Enable                    ='on';
        handles.edit48.Enable                    ='on';
    end
    
    handles.D_alpha_prior.Enable                 ='on';
    handles.D_beta_prior.Enable                  ='on';
    handles.mu_alpha.Enable                      ='on';
    handles.mu_beta.Enable                       ='on';
    handles.Number_particles.Enable              ='on';
    handles.alpha_load.Enable                    ='on';
    handles.beta_load.Enable                     ='on';
    handles.concen_radious.Enable                ='on';
    handles.mu_prior_xy.Enable                   ='on';
    handles.mu_prior_z.Enable                    ='on';
    handles.var_prior_xy.Enable                  ='on';
    handles.var_prior_z.Enable                   ='on';    
        
        
    end


function var_prior_z_Callback(hObject, eventdata, handles)
% hObject    handle to var_prior_z (see GCBO)
% eventdata  reserved - to be defined in a future version of MATLAB
% handles    structure with handles and user data (see GUIDATA)

% Hints: get(hObject,'String') returns contents of var_prior_z as text
%        str2double(get(hObject,'String')) returns contents of var_prior_z as a double


% --- Executes during object creation, after setting all properties.
function var_prior_z_CreateFcn(hObject, eventdata, handles)
% hObject    handle to var_prior_z (see GCBO)
% eventdata  reserved - to be defined in a future version of MATLAB
% handles    empty - handles not created until after all CreateFcns called

% Hint: edit controls usually have a white background on Windows.
%       See ISPC and COMPUTER.
if ispc && isequal(get(hObject,'BackgroundColor'), get(0,'defaultUicontrolBackgroundColor'))
    set(hObject,'BackgroundColor','white');
end


% --- Executes on selection change in concen_radious.
function concen_radious_Callback(hObject, eventdata, handles)
% hObject    handle to concen_radious (see GCBO)
% eventdata  reserved - to be defined in a future version of MATLAB
% handles    structure with handles and user data (see GUIDATA)

% Hints: contents = cellstr(get(hObject,'String')) returns concen_radious contents as cell array
%        contents{get(hObject,'Value')} returns selected item from concen_radious


% --- Executes during object creation, after setting all properties.
function concen_radious_CreateFcn(hObject, eventdata, handles)
% hObject    handle to concen_radious (see GCBO)
% eventdata  reserved - to be defined in a future version of MATLAB
% handles    empty - handles not created until after all CreateFcns called

% Hint: listbox controls usually have a white background on Windows.
%       See ISPC and COMPUTER.
if ispc && isequal(get(hObject,'BackgroundColor'), get(0,'defaultUicontrolBackgroundColor'))
    set(hObject,'BackgroundColor','white');
end


% --- Executes on button press in pushbutton22.
function pushbutton22_Callback(hObject, eventdata, handles)
% hObject    handle to pushbutton22 (see GCBO)
% eventdata  reserved - to be defined in a future version of MATLAB
% handles    structure with handles and user data (see GUIDATA)


    handles.pushbutton8.Enable                  ='off';
    
    
    handles.pushbutton20.Enable                 ='off';
    handles.pushbutton17.Enable                 ='off';
    
    if  sum(handles.pushbutton22.String)==sum('Estimate background emission rate (ON)')
        handles.pushbutton22.String                 ='Estimate background emission rate (OFF)';
        handles.edit46.Enable                       ='off';
        handles.edit47.Enable                       ='off';
        handles.edit48.Enable                       ='off';
        if  sum(handles.pushbutton23.String)==sum('Automatic (OFF)')
                handles.pushbutton17.Enable                 ='on';
            else
                handles.pushbutton17.Enable                 ='off';
                handles.pushbutton8.Enable                  ='on';
        end
    else
        choice = questdlg('To estimate the background photon emission rate, a different filter known as Unscented Kalman filter (UKF) is used which is slower than the default filter (Extended Kalman Filter). Do you still want to continue?','Warning!','YES','NO','NO');
    
        if  strcmp(choice,'YES')
            handles.pushbutton22.String                 ='Estimate background emission rate (ON)';
            handles.edit46.Enable                       ='on';
            handles.edit47.Enable                       ='on';
            handles.edit48.Enable                       ='on';
            
            if  sum(handles.pushbutton23.String)==sum('Automatic (OFF)')
                handles.pushbutton17.Enable                 ='on';
            else
                handles.pushbutton17.Enable                 ='off';
                handles.pushbutton8.Enable                  ='on';
            end
                
        end
        
    end
    
    
    
    
    
    
function edit47_Callback(hObject, eventdata, handles)
% hObject    handle to edit47 (see GCBO)
% eventdata  reserved - to be defined in a future version of MATLAB
% handles    structure with handles and user data (see GUIDATA)

% Hints: get(hObject,'String') returns contents of edit47 as text
%        str2double(get(hObject,'String')) returns contents of edit47 as a double


% --- Executes during object creation, after setting all properties.
function edit47_CreateFcn(hObject, eventdata, handles)
% hObject    handle to edit47 (see GCBO)
% eventdata  reserved - to be defined in a future version of MATLAB
% handles    empty - handles not created until after all CreateFcns called

% Hint: edit controls usually have a white background on Windows.
%       See ISPC and COMPUTER.
if ispc && isequal(get(hObject,'BackgroundColor'), get(0,'defaultUicontrolBackgroundColor'))
    set(hObject,'BackgroundColor','white');
end


function edit46_Callback(hObject, eventdata, handles)
% hObject    handle to edit46 (see GCBO)
% eventdata  reserved - to be defined in a future version of MATLAB
% handles    structure with handles and user data (see GUIDATA)

% Hints: get(hObject,'String') returns contents of edit46 as text
%        str2double(get(hObject,'String')) returns contents of edit46 as a double


% --- Executes during object creation, after setting all properties.
function edit46_CreateFcn(hObject, eventdata, handles)
% hObject    handle to edit46 (see GCBO)
% eventdata  reserved - to be defined in a future version of MATLAB
% handles    empty - handles not created until after all CreateFcns called

% Hint: edit controls usually have a white background on Windows.
%       See ISPC and COMPUTER.
if ispc && isequal(get(hObject,'BackgroundColor'), get(0,'defaultUicontrolBackgroundColor'))
    set(hObject,'BackgroundColor','white');
end


% --- Executes on button press in checkbox7.
function checkbox7_Callback(hObject, eventdata, handles)
% hObject    handle to checkbox7 (see GCBO)
% eventdata  reserved - to be defined in a future version of MATLAB
% handles    structure with handles and user data (see GUIDATA)

% Hint: get(hObject,'Value') returns toggle state of checkbox7


function edit48_Callback(hObject, eventdata, handles)
% hObject    handle to edit48 (see GCBO)
% eventdata  reserved - to be defined in a future version of MATLAB
% handles    structure with handles and user data (see GUIDATA)

% Hints: get(hObject,'String') returns contents of edit48 as text
%        str2double(get(hObject,'String')) returns contents of edit48 as a double


% --- Executes during object creation, after setting all properties.
function edit48_CreateFcn(hObject, eventdata, handles)
% hObject    handle to edit48 (see GCBO)
% eventdata  reserved - to be defined in a future version of MATLAB
% handles    empty - handles not created until after all CreateFcns called

% Hint: edit controls usually have a white background on Windows.
%       See ISPC and COMPUTER.
if ispc && isequal(get(hObject,'BackgroundColor'), get(0,'defaultUicontrolBackgroundColor'))
    set(hObject,'BackgroundColor','white');
end


% --- Executes on button press in pushbutton23.
function pushbutton23_Callback(hObject, eventdata, handles)
% hObject    handle to pushbutton23 (see GCBO)
% eventdata  reserved - to be defined in a future version of MATLAB
% handles    structure with handles and user data (see GUIDATA)


    
    handles.checkbox1.Enable                     ='off';
    handles.checkbox2.Enable                     ='off';
    handles.checkbox3.Enable                     ='off';
    handles.checkbox4.Enable                     ='off';
    
    
    handles.pushbutton11.Enable                  ='off';
    handles.pushbutton16.Enable                  ='off';
    
    handles.pushbutton20.Enable                  ='off';
     
    handles.pushbutton23.Enable                  ='off';
    handles.pushbutton24.Enable                  ='off';
    handles.pushbutton25.Enable                  ='off';
    handles.pushbutton26.Enable                  ='off';
    
    
    
   if  sum(handles.pushbutton23.String)  == sum('Automatic (ON)')
       handles.pushbutton23.String                  =    'Automatic (OFF)' ;
       handles.edit39.Enable                        =    'on'             ;
       handles.edit41.Enable                        =    'on'             ;
       handles.pushbutton8.Enable                   =    'off'            ;
       handles.pushbutton17.Enable                  =    'on'             ;
       handles.edit52.Enable                        =    'off'            ;
       if  sum(handles.pushbutton22.String)  == sum('Estimate background emission rate (OFF)')
           
           handles.edit46.Enable                    =    'off'            ;
           handles.edit47.Enable                    =    'off'            ;
           handles.edit48.Enable                    =    'off'            ;
           
       else
           handles.edit46.Enable                    =    'on'             ;
           handles.edit47.Enable                    =    'on'             ;
           handles.edit48.Enable                    =    'on'             ;
       end
   else
      
    choice = questdlg({['Automatic process will follows the convergence to ',...
                       'the stationary distribution of the diffusion coefficient. '...
                       'Computational time in this mode depends on many factors. '...
                       'So, this process could take ',...
                       'a while, depends on the imported signal. For the '...
                       'safty, results of the MCMC chain will save and show '...
                       'after every 10000 iterations. If you still want to '...
                       'continue, choose YES and click on the start button']},...
                       'Warning!','Yes','No','Yes');
    
    if  strcmp(choice,'Yes')
        handles.pushbutton8.String                   =    'Start'           ;
        handles.pushbutton23.String                  =    'Automatic (ON)'   ;
        handles.edit39.Enable                        =    'off'             ;
        handles.edit41.Enable                        =    'off'             ;
        handles.pushbutton8.Enable                   =    'on'              ;
        handles.pushbutton17.Enable                  =    'off'             ;
        handles.edit52.Enable                        =    'on'              ;
    else
        handles.edit39.Enable                        =    'on'              ;
        handles.edit41.Enable                        =    'on'              ;
        handles.pushbutton8.Enable                   =    'off'             ;
        handles.pushbutton17.Enable                  =    'on'              ;
        handles.edit52.Enable                        =    'off'             ;
        
        if  sum(handles.pushbutton22.String)  == sum('Estimate background emission rate (OFF)')
            handles.edit46.Enable                    =    'off'             ;
            handles.edit47.Enable                    =    'off'             ;
            handles.edit48.Enable                    =    'off'             ;
        else
            handles.edit46.Enable                    =    'on'             ;
            handles.edit47.Enable                    =    'on'             ;
            handles.edit48.Enable                    =    'on'             ;
        end
    end 
    
   end


    handles.edit35.Enable                        ='on';
    handles.edit36.Enable                        ='on';
    handles.edit40.Enable                        ='on';
    
    
    if sum(handles.pushbutton22.String)==sum('Estimate background emission rate (ON)')
        handles.edit48.Enable                        ='on';
    end
    handles.pushbutton10.Enable                  ='on';
    handles.pushbutton22.Enable                  ='on';
    handles.pushbutton23.Enable                  ='on';
    handles.pushbutton24.Enable                  ='on';
    handles.pushbutton25.Enable                  ='on';
    handles.pushbutton26.Enable                  ='on';
    
    handles.D_alpha_prior.Enable                 ='on';
    
    
    
%%%%%%%%%%%%%%%%%%%%%%%%%%%%%%%%% Initial vslues %%%%%%%%%%%%%%%%%%%%%%%%%%%%%%%%%%%
     load('results.mat');
     
     len = length(Data.Trace_partial)                                     ;
     for k=1:len
         sign_siz(k)   = length(Data.Trace_partial{k})                    ; 
         Data.x{k}     = zeros(3*str2num(handles.Number_particles.String),sign_siz(k))+.1  ;
     end
     
     
     iter_b_mu   =     20         ;
     
     Data.D             = 1                                                                                 ;
     Data.mu            = 300000                                                                            ;
     Data.b             = zeros(len,str2num(handles.Number_particles.String),1)                              ;
     
     
     
     Data.q             = zeros(len,str2num(handles.Number_particles.String),1)                              ;                                                  ;          
     Data.concentration =zeros(len , 1000 ,max(sign_siz) , length(str2num(handles.concen_radious.String)) ) ;

   
     if      sum(handles.pushbutton28.String)==sum('PSF (Gaussian)')          
             PSF_Func=1;     
     elseif  sum(handles.pushbutton28.String)==sum('PSF (Lorentzian)')            
             PSF_Func=2;        
     else
             PSF_Func=0;
     end
     
     if  sum(handles.pushbutton22.String)==sum('Estimate background emission rate (OFF)')
         Filter_typ=1;
         Data.mu_back     = str2num(handles.mu_back.String)                                 ;
         handles.edit46.String = num2str(str2num(handles.mu_back.String)*10*sum(sign_siz)) ;
         handles.edit47.String = num2str(1/(10*sum(sign_siz))) ;
     else
         Filter_typ=2;
         Data.mu_back     = 1000                                                            ;
     end
    
    
    


% --- Executes on button press in checkbox8.
function checkbox8_Callback(hObject, eventdata, handles)
% hObject    handle to checkbox8 (see GCBO)
% eventdata  reserved - to be defined in a future version of MATLAB
% handles    structure with handles and user data (see GUIDATA)

% Hint: get(hObject,'Value') returns toggle state of checkbox8


% --- Executes on button press in pushbutton25.
function pushbutton25_Callback(hObject, eventdata, handles)
% hObject    handle to pushbutton25 (see GCBO)
% eventdata  reserved - to be defined in a future version of MATLAB
% handles    structure with handles and user data (see GUIDATA)


% --- Executes on button press in pushbutton24.
function pushbutton24_Callback(hObject, eventdata, handles)
% hObject    handle to pushbutton24 (see GCBO)
% eventdata  reserved - to be defined in a future version of MATLAB
% handles    structure with handles and user data (see GUIDATA)


% --- Executes on button press in pushbutton26.
function pushbutton26_Callback(hObject, eventdata, handles)
% hObject    handle to pushbutton26 (see GCBO)
% eventdata  reserved - to be defined in a future version of MATLAB
% handles    structure with handles and user data (see GUIDATA)


% --- Executes on button press in pushbutton28.
function pushbutton28_Callback(hObject, eventdata, handles)
% hObject    handle to pushbutton28 (see GCBO)
% eventdata  reserved - to be defined in a future version of MATLAB
% handles    structure with handles and user data (see GUIDATA)

choice = questdlg('Please choose the Point Spread Function of interest:','Point Spread Function','Gaussian','Lorentzian','Cylindrical','');
    
    if  strcmp(choice,'Gaussian')
        handles.pushbutton28.String='PSF (Gaussian)';
        handles.pushbutton23.Enable = 'on';
        
        if  sum(handles.pushbutton23.String)==sum('Automatic (OFF)')
            handles.pushbutton17.Enable                  ='on';
            handles.edit52.Enable                        ='off';
        else
            handles.pushbutton17.Enable                  ='off';
            handles.pushbutton8.Enable                   ='on';
            handles.edit52.Enable                        ='on';
        end
        
        handles.pushbutton22.Enable                  ='on';
        handles.edit39.Enable                        ='on';
        handles.edit40.Enable                        ='on';
        handles.edit41.Enable                        ='on';
        
        
        
        if  sum(handles.pushbutton22.String)==sum('Estimate background emission rate (ON)')
            handles.edit46.Enable                        ='on';
            handles.edit47.Enable                        ='on';
            handles.edit48.Enable                        ='on';
        else
            handles.edit46.Enable                        ='off';
            handles.edit47.Enable                        ='off';
            handles.edit48.Enable                        ='off';
        end
        
        handles.D_alpha_prior.Enable                 ='on';
        handles.D_beta_prior.Enable                  ='on';
        handles.mu_alpha.Enable                      ='on';
        handles.mu_beta.Enable                       ='on';
        handles.Number_particles.Enable              ='on';
        handles.alpha_load.Enable                    ='on';
        handles.beta_load.Enable                     ='on';
        handles.concen_radious.Enable                ='on';
        handles.mu_prior_xy.Enable                   ='on';
        handles.mu_prior_z.Enable                    ='on';
        handles.var_prior_xy.Enable                  ='on';
        handles.var_prior_z.Enable                   ='on';
    
    
    elseif strcmp(choice,'Lorentzian')
           handles.pushbutton28.String                  ='PSF (Lorentzian)';
           handles.pushbutton23.Enable                  ='on';
          
           if  sum(handles.pushbutton23.String)==sum('Automatic (OFF)')
               handles.pushbutton17.Enable                  ='on';
               handles.edit52.Enable                        ='off';
               handles.pushbutton8.Enable                   ='off';
           else
               handles.pushbutton8.Enable                   ='on';
               handles.pushbutton17.Enable                  ='off';
               handles.edit52.Enable                        ='on';
           end
           
           handles.pushbutton22.Enable                  ='on';
           handles.edit39.Enable                        ='on';
           handles.edit40.Enable                        ='on';
           handles.edit41.Enable                        ='on';
           
           if  sum(handles.pushbutton22.String)==sum('Estimate background emission rate (ON)')
               handles.edit46.Enable                        ='on';
               handles.edit47.Enable                        ='on';
               handles.edit48.Enable                        ='on';
           else
               handles.edit46.Enable                        ='off';
               handles.edit47.Enable                        ='off';
               handles.edit48.Enable                        ='off';
           end
           
           handles.D_alpha_prior.Enable                 ='on';
           handles.D_beta_prior.Enable                  ='on';
           handles.mu_alpha.Enable                      ='on';
           handles.mu_beta.Enable                       ='on';
           handles.Number_particles.Enable              ='on';
           handles.alpha_load.Enable                    ='on';
           handles.beta_load.Enable                     ='on';
           handles.concen_radious.Enable                ='on';
           handles.mu_prior_xy.Enable                   ='on';
           handles.mu_prior_z.Enable                    ='on';
           handles.var_prior_xy.Enable                  ='on';
           handles.var_prior_z.Enable                   ='on';
     
    elseif strcmp(choice,'Cylindrical')
           handles.pushbutton28.String                  ='PSF (Cylindrical)';
           handles.pushbutton23.Enable                  ='on';
          
           if  sum(handles.pushbutton23.String)==sum('Automatic (OFF)')
               handles.pushbutton17.Enable                  ='on';
               handles.edit52.Enable                        ='off';
               handles.pushbutton8.Enable                   ='off';
           else
               handles.pushbutton8.Enable                   ='on';
               handles.pushbutton17.Enable                  ='off';
               handles.edit52.Enable                        ='on';
           end
           
           handles.pushbutton22.Enable                  ='on';
           handles.edit39.Enable                        ='on';
           handles.edit40.Enable                        ='on';
           handles.edit41.Enable                        ='on';
           
           if  sum(handles.pushbutton22.String)==sum('Estimate background emission rate (ON)')
               handles.edit46.Enable                        ='on';
               handles.edit47.Enable                        ='on';
               handles.edit48.Enable                        ='on';
           else
               handles.edit46.Enable                        ='off';
               handles.edit47.Enable                        ='off';
               handles.edit48.Enable                        ='off';
           end
           
           handles.D_alpha_prior.Enable                 ='on';
           handles.D_beta_prior.Enable                  ='on';
           handles.mu_alpha.Enable                      ='on';
           handles.mu_beta.Enable                       ='on';
           handles.Number_particles.Enable              ='on';
           handles.alpha_load.Enable                    ='on';
           handles.beta_load.Enable                     ='on';
           handles.concen_radious.Enable                ='on';
           handles.mu_prior_xy.Enable                   ='on';
           handles.mu_prior_z.Enable                    ='on';
           handles.var_prior_xy.Enable                  ='on';
           handles.var_prior_z.Enable                   ='on';
     
    else
          handles.pushbutton28.String='PSF (NAN)';
          handles.pushbutton23.Enable                  ='off';
          handles.pushbutton17.Enable                  ='off';
          handles.pushbutton22.Enable                  ='off';
          handles.edit39.Enable                        ='off';
          handles.edit40.Enable                        ='off';
          handles.edit41.Enable                        ='off';
          
          handles.edit46.Enable                        ='off';
          handles.edit47.Enable                        ='off';
          handles.edit48.Enable                        ='off';
          
          handles.D_alpha_prior.Enable                 ='off';
          handles.D_beta_prior.Enable                  ='off';
          handles.mu_alpha.Enable                      ='off';
          handles.mu_beta.Enable                       ='off';
          handles.Number_particles.Enable              ='off';
          handles.alpha_load.Enable                    ='off';
          handles.beta_load.Enable                     ='off';
          handles.concen_radious.Enable                ='off';
          handles.mu_prior_xy.Enable                   ='off';
          handles.mu_prior_z.Enable                    ='off';
          handles.var_prior_xy.Enable                  ='off';
          handles.var_prior_z.Enable                   ='off';
    end
    handles.pushbutton18.Enable                  ='on';
    handles.pushbutton19.Enable                  ='on';
    
    handles.checkbox1.Enable                     ='off';
    handles.checkbox2.Enable                     ='off';
    handles.checkbox3.Enable                     ='off';
    handles.checkbox4.Enable                     ='off';
    handles.checkbox7.Enable                     ='off';
    handles.checkbox8.Enable                     ='off';
        


function edit51_Callback(hObject, eventdata, handles)
% hObject    handle to edit51 (see GCBO)
% eventdata  reserved - to be defined in a future version of MATLAB
% handles    structure with handles and user data (see GUIDATA)

% Hints: get(hObject,'String') returns contents of edit51 as text
%        str2double(get(hObject,'String')) returns contents of edit51 as a double


% --- Executes during object creation, after setting all properties.
function edit51_CreateFcn(hObject, eventdata, handles)
% hObject    handle to edit51 (see GCBO)
% eventdata  reserved - to be defined in a future version of MATLAB
% handles    empty - handles not created until after all CreateFcns called

% Hint: edit controls usually have a white background on Windows.
%       See ISPC and COMPUTER.
if ispc && isequal(get(hObject,'BackgroundColor'), get(0,'defaultUicontrolBackgroundColor'))
    set(hObject,'BackgroundColor','white');
end


function edit52_Callback(hObject, eventdata, handles)
% hObject    handle to edit52 (see GCBO)
% eventdata  reserved - to be defined in a future version of MATLAB
% handles    structure with handles and user data (see GUIDATA)

% Hints: get(hObject,'String') returns contents of edit52 as text
%        str2double(get(hObject,'String')) returns contents of edit52 as a double


% --- Executes during object creation, after setting all properties.
function edit52_CreateFcn(hObject, eventdata, handles)
% hObject    handle to edit52 (see GCBO)
% eventdata  reserved - to be defined in a future version of MATLAB
% handles    empty - handles not created until after all CreateFcns called

% Hint: edit controls usually have a white background on Windows.
%       See ISPC and COMPUTER.
if ispc && isequal(get(hObject,'BackgroundColor'), get(0,'defaultUicontrolBackgroundColor'))
    set(hObject,'BackgroundColor','white');
end


% --- Executes on button press in pushbutton29.
function pushbutton29_Callback(hObject, eventdata, handles)
% hObject    handle to pushbutton29 (see GCBO)
% eventdata  reserved - to be defined in a future version of MATLAB
% handles    structure with handles and user data (see GUIDATA)

    load('results.mat');
    
    trace_bin           = Data.Trace_uniform_bin   ;
    
    
    minn = str2num(get(handles.edit35,'String')) ;
    maxx = str2num(get(handles.edit36,'String')) ;
    
    
     while isempty(maxx-minn) || maxx-minn<=0 || maxx<0 || minn<0 || maxx>length(trace_bin)-1

              opts.Interpreter = 'tex';
              answer = inputdlg({'Minimum Time step','Maximum Time step'},...
                       'Please choose reasonable vlues!',[1 60;1 60],...
                       {'0',num2str(length(Data.Trace_partial)-1)},opts);
                   
              minn = str2num(answer{1}) ;
              maxx = str2num(answer{2}) ;
    end

    handles.edit35.String  = num2str(minn);
    handles.edit36.String  = num2str(maxx);

    
    
    if     sum(str2num(char(handles.uitable5.Data)))==0
           Size_signal=0;
           
    elseif sum(handles.uitable5.Data{1,1})==0
           Size_signal=0;
    else
        Trace_partial = Data.Trace_partial;
        Time          = Data.Time_partial;
        Size_signal   = length(Trace_partial);
    end
    
    handles.uitable5.Data{Size_signal+1,1}=num2str(minn);
    handles.uitable5.Data{Size_signal+1,2}=num2str(maxx);
    

    if length(Data.Trace_type) == length('Time_arrival')
    
    trace_singel_photon = Data.Trace_singel_photon ;
    
    mminn=min(find(trace_singel_photon>=(minn-1)*str2num(handles.Delta_t.String)));
    mmaxx=max(find(trace_singel_photon<=maxx*str2num(handles.Delta_t.String)));
    edge=cumsum(...
        [0,repmat(...
           str2num(handles.Delta_t.String),1,...
           floor(trace_singel_photon(mmaxx)/str2num(handles.Delta_t.String))-...
           floor(trace_singel_photon(mminn)/str2num(handles.Delta_t.String))...
           )...
           ]);
       
    if  mminn==1
        sss=0;
    else
        sss=trace_singel_photon(mminn);
    end
    
    
    % Consider the portion of the singel photon time arrivals trace
    single_trace=trace_singel_photon(mminn:mmaxx)-sss;
    
    [Trace_first,~] = histcounts(single_trace,edge);


    
  
    % Calculate the adaptive binned trace
    [Trace_partial{Size_signal+1},Time{Size_signal+1}] = ...
     ...
     Adaptive_bin(str2num(handles.edit53.String)    ,...
                  Trace_first                       ,...
                  single_trace                      ,...
                  str2num(handles.Delta_t.String)  ); 
    
    Data.Trace_partial   =    Trace_partial  ;
    Data.Time_partial    =    Time           ;
    
    
    save('results','Data')

    if  length(Data.Trace_partial)>0
        handles.pushbutton28.Enable = 'on';
        handles.pushbutton30.Enable = 'on';
        handles.pushbutton31.Enable = 'on';
    else
        handles.pushbutton28.Enable = 'off';
        handles.pushbutton30.Enable = 'off';
        handles.pushbutton31.Enable = 'off';
    end
    
    else
      
        signald   = trace_bin(minn+1:maxx)'     ;
        Timee     = ones(1,length(signald))  ;

endloop=0;
i=1;
for n=1:4
    endloop=0;
    i=2;
while endloop==0
    if signald(i-1) < str2num(handles.edit53.String)
       signald(i-1) = signald(i-1)+signald(i)    ;
       Timee(i-1)    = Timee(i-1)+Timee(i)          ;
    
    if  i      <= length(signald)-2
        signald = [signald(1:i-1),signald(i+1:end)];
        Timee    = [Timee(1:i-1),Timee(i+1:end)];
        i       = i+1;
    else
        signald = [signald(1:i-1),signald(i+1:end)];
        Timee    = [Timee(1:i-1),Timee(i+1:end)];
        endloop = 1;
    end
    else
        if  i<=length(signald)-1
            i=i+1;
        else
            endloop=1;
        end
    end
        
end
end

Trace_partial{Size_signal+1}=signald;
Time{Size_signal+1}  = Timee*str2num(handles.Delta_t.String);
        Data.Trace_partial   =    Trace_partial  ;
        Data.Time_partial    =    Time           ;
    
    
        save('results','Data')

        if  length(Data.Trace_partial)>0
            handles.pushbutton28.Enable = 'on';
            handles.pushbutton30.Enable = 'on';
            handles.pushbutton31.Enable = 'on';
        else
            handles.pushbutton28.Enable = 'off';
            handles.pushbutton30.Enable = 'off';
            handles.pushbutton31.Enable = 'off';
        end
        
        
    end
    
    
    
    

% --- Executes on button press in pushbutton30.
function pushbutton30_Callback(hObject, eventdata, handles)
% hObject    handle to pushbutton30 (see GCBO)
% eventdata  reserved - to be defined in a future version of MATLAB
% handles    structure with handles and user data (see GUIDATA)

    load('results.mat');

    num_signals=length(Data.Trace_partial);

    opts.Interpreter = 'tex';
    answer = inputdlg({'Which trace you wish to delete?'},...
                       'Delete trace',[1 60],...
                       {''},opts);
  
    while sum(find(str2num(answer{1})==[1:num_signals]))==0
          
          opts.Interpreter = 'tex';
          answer = inputdlg({['Please choose values between 1 to ',num2str(num_signals)]},...
                       'Delete trace',[1 60],...
                       {''},opts);
    end
    
    Data.Trace_partial(str2num(answer{1}))=[];
    handles.uitable5.Data(str2num(answer{1}),:)=[];

    
    if   sum(cell2mat(handles.uitable5.Data))==0
        
         handles.pushbutton28.Enable = 'off';
         handles.pushbutton30.Enable = 'off';
         handles.pushbutton31.Enable = 'off';
    end
        
    save('results','Data')

    
    
    
    

% --- Executes on button press in pushbutton31.
function pushbutton31_Callback(hObject, eventdata, handles)
% hObject    handle to pushbutton31 (see GCBO)
% eventdata  reserved - to be defined in a future version of MATLAB
% handles    structure with handles and user data (see GUIDATA)


    load('results.mat');
    
    trace_bin           = Data.Trace_uniform_bin   ;
    trace_singel_photon = Data.Trace_singel_photon ;
    
    
    minn = str2num(get(handles.edit35,'String')) ;
    maxx = str2num(get(handles.edit36,'String')) ;
    
    
     while isempty(maxx-minn) || maxx-minn<=0 || maxx<0 || minn<0 || maxx>length(trace_bin)-1

              opts.Interpreter = 'tex';
              answer = inputdlg({'Minimum Time step','Maximum Time step'},...
                       'Please choose reasonable values!',[1 60;1 60],...
                       {'0',num2str(length(Data.Trace_partial)-1)},opts);
                   
              minn = str2num(answer{1}) ;
              maxx = str2num(answer{2}) ;
    end

    handles.edit35.String  = num2str(minn);
    handles.edit36.String  = num2str(maxx);
    
    
     if     sum(str2num(char(handles.uitable5.Data)))==0
           Size_signal=0;
           
    elseif sum(handles.uitable5.Data{1,1})==0
           Size_signal=0;
    else
        Trace_partial = Data.Trace_partial;
        Time          = Data.Time_partial;
        Size_signal   = length(Trace_partial);
    end
    
    
    mminn=min(find(trace_singel_photon>=(minn-1)*str2num(handles.Delta_t.String)));
    mmaxx=max(find(trace_singel_photon<=maxx*str2num(handles.Delta_t.String)));
    edge=cumsum(...
        [0,repmat(...
           str2num(handles.Delta_t.String),1,...
           floor(trace_singel_photon(mmaxx)/str2num(handles.Delta_t.String))-...
           floor(trace_singel_photon(mminn)/str2num(handles.Delta_t.String))...
           )...
           ]);
       
    if  mminn==1
        sss=0;
    else
        sss=trace_singel_photon(mminn);
    end
    
    
    % Consider the portion of the singel photon time arrivals trace
    single_trace=trace_singel_photon(mminn:mmaxx)-sss;
    
    [Trace_first,edges] = histcounts(single_trace,edge);


    
  
    
    opts.Interpreter = 'tex';
    answer = inputdlg({'Which trace you wish to change?'},...
                       'change trace',[1 60],...
                       {''},opts);
  
    while sum(find(str2num(answer{1})==[1:Size_signal]))==0
          
          opts.Interpreter = 'tex';
          answer = inputdlg({['Please choose values between 1 to ',num2str(num_Traces)]},...
                       'change trace',[1 60],...
                       {''},opts);
    end
    
    
     % Calculate the adaptive binned trace
    [Trace_partial{str2num(answer{1})},Time{str2num(answer{1})}] = ...
     ...
     Adaptive_bin(str2num(handles.edit53.String)    ,...
                  Trace_first                       ,...
                  single_trace                      ,...
                  str2num(handles.Delta_t.String)  ); 
    
    Data.Trace_partial   =    Trace_partial  ;
    Data.Time_partial    =    Time           ;
    
    
    handles.uitable5.Data(str2num(answer{1}),1:2)=[{handles.edit35.String} ,{handles.edit36.String}];
    
    save('results','Data')


% --- Executes on button press in pushbutton32.
function pushbutton32_Callback(hObject, eventdata, handles)
% hObject    handle to pushbutton32 (see GCBO)
% eventdata  reserved - to be defined in a future version of MATLAB
% handles    structure with handles and user data (see GUIDATA)
hFig = figure('Menubar','none', 'Toolbar','none','numbertitle','off','name','ASWFCS README');
set(hFig,'Units','Characters','Position',get(hFig,'Position').*[1 1 0 0.12]+[0 0 80 0]);
uicontrol(hFig, 'Style','edit',...
    'Min',0, 'Max',2, 'HorizontalAlignment','left', ...
    'Units','normalized', 'Position',[0 0 1 1], ...
    'String',fileread('README.txt'));
movegui(hFig,'center');


function edit53_Callback(hObject, eventdata, handles)
% hObject    handle to edit53 (see GCBO)
% eventdata  reserved - to be defined in a future version of MATLAB
% handles    structure with handles and user data (see GUIDATA)

% Hints: get(hObject,'String') returns contents of edit53 as text
%        str2double(get(hObject,'String')) returns contents of edit53 as a double


% --- Executes during object creation, after setting all properties.
function edit53_CreateFcn(hObject, eventdata, handles)
% hObject    handle to edit53 (see GCBO)
% eventdata  reserved - to be defined in a future version of MATLAB
% handles    empty - handles not created until after all CreateFcns called

% Hint: edit controls usually have a white background on Windows.
%       See ISPC and COMPUTER.
if ispc && isequal(get(hObject,'BackgroundColor'), get(0,'defaultUicontrolBackgroundColor'))
    set(hObject,'BackgroundColor','white');
end


% --- Executes on button press in pushbutton33.
function pushbutton33_Callback(hObject, eventdata, handles)
% hObject    handle to pushbutton33 (see GCBO)
% eventdata  reserved - to be defined in a future version of MATLAB
% handles    structure with handles and user data (see GUIDATA)
open('Help.pdf')

% --- Executes when figure1 is resized.
function figure1_SizeChangedFcn(hObject, eventdata, handles)
% hObject    handle to figure1 (see GCBO)
% eventdata  reserved - to be defined in a future version of MATLAB
% handles    structure with handles and user data (see GUIDATA)
```
