## Supplementary material for "An Alternative Framework for Fluorescence Correlation Spectroscopy": Source Code: Help.pdf

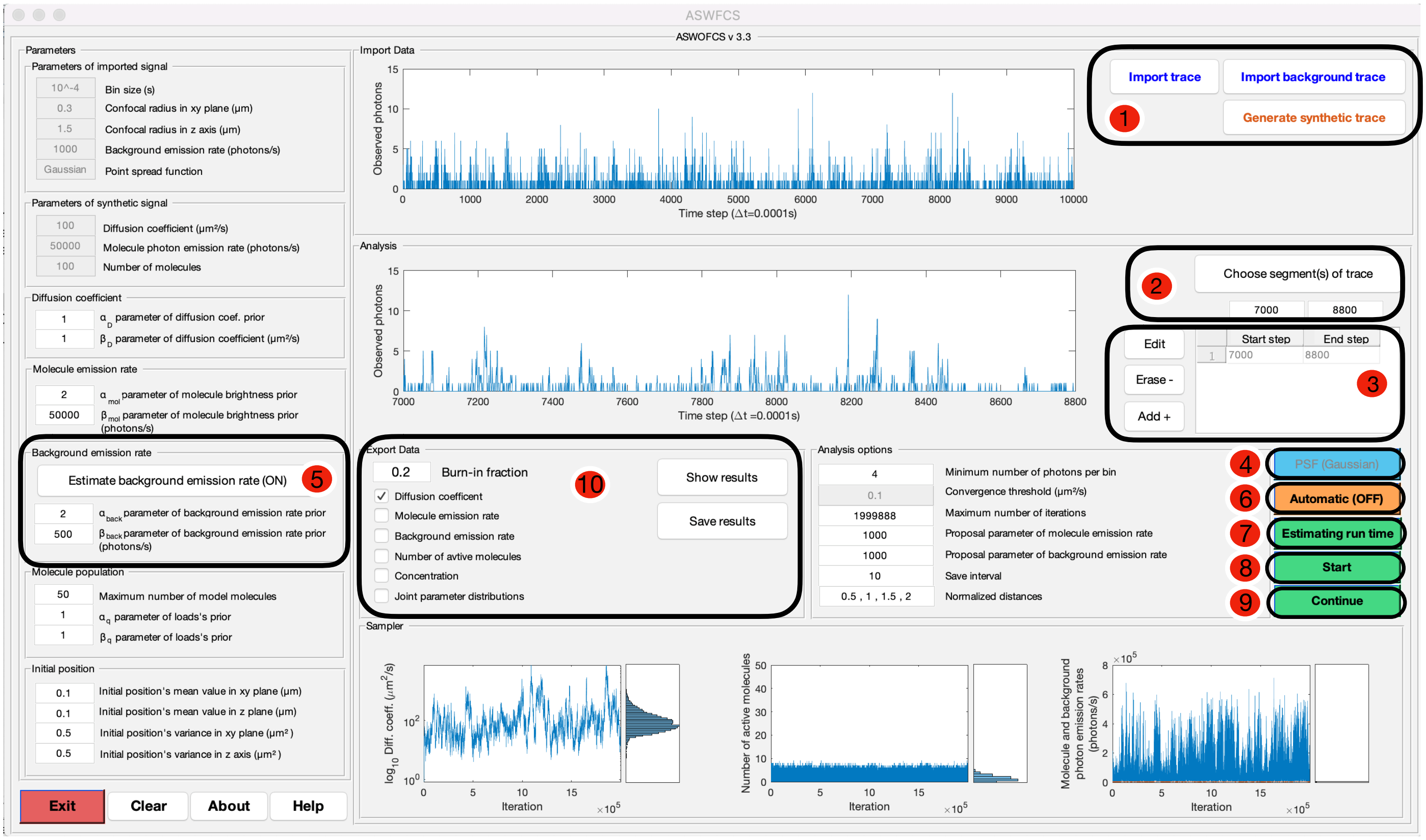


1: First, Import the trace or generate a trace

- If you have a background trace, import it first to measure the background.
- If you do not provide a background trace, the value of the background emission rate will be asked.
- If you do not provide a background emission rate, you need to turn on the UKF by clicking on number 5 when it is available to you.

2: Choose a segment of the trace – the chosen trace will be shown in the left panel.

3: After the segment is chosen, click on Add+. You are able to add as many segments as you wish.

- You can erase or edit the chosen traces.

4: Choose the PSF type (Gaussian, Lorentzian or Cylindrical).

5: You are able to change the Filter type (EKF or UKF).

- If you choose EKF, it assumes that the background emission rate is known.
- If you choose UKF, it learns the background, but is slower. There will be more options available.

6: You can set the sampler to terminate the MCMC chain, automatically, after the convergence threshold condition have been met. Please be aware that automatic processing time depends on the imported trace(s). To prevent any conflict and confusion, all buttons will be off during this process.

7: If automatic mode is turned off, an estimated run time of the code (based on the number of iterations) will be provided.

8: By clicking on the Start button, the code will begin to run. This process will take time, and based on the chosen save window sizes the samples will be shown and saved. To prevent any conflict and confusion, all buttons will be off during this process.

9: After execution, if needed, the user can increase the number iterations.

10: There are buttons to show or save the results.
