## Supplementary material for "An Alternative Framework for Fluorescence Correlation Spectroscopy": Source Code: README.txt

===========================================================================
Matlab software implementation for:
    AN ALTERNATIVE FRAMEWORK FOR FLUORESCENCE CORRELATION SPECTROSCOPY
    Nature Communication (2019)
===========================================================================
 
Copyright (C) 2019, Sina Jazani, and Steve Press�

 
This software package includes several Matlab scripts and auxiliary
functions, which implement the computational algorithms for the method
ASWFCS described in the following publication:
 
    An Alterantive Framework for Fluorescence Correlation Spectroscopy
    Sina Jazani, Ioannis Sgouralis, Omer M. Shafraz, Marcia Levitus, Sanjeevi Sivasankar 
    and Steve Presse
    Nature Communication (2019)


Please cite the papers above in any publications using this software.

 
===========================================================================
Setup and Usage
===========================================================================
  
1) To use this software in GUI form run:
    ASWFCS
in Matlab's command window.
 
 
2) For further details or to report bugs contact:
   

3) This software (source code and GUI) has been developed in Matlab R2018b.

 
===========================================================================
Copyright and License
===========================================================================
 
Copyright (C) 2019, Sina Jazani, and Steve Press�
 
 
Permission is granted for anyone to copy, use, or modify these programs 
and accompanying documents for purposes of research or education, provided
this copyright notice is retained, and note is made of any changes that
have been made.
 
These programs and documents are distributed without any warranty, express 
or implied.  As the programs were written for research purposes only, they 
have not been tested to the degree that would be advisable in any important
application.  All use of these programs is entirely at the user's own risk.
