## Supplementary material for "An Alternative Framework for Fluorescence Correlation Spectroscopy": Source Code: Adaptive_bin.m

```
function[Trace , Time] = Adaptive_bin( Threshold , Trace_uniform_bin , Trace_single_photon , Deltat)

%%%%%%%%%%%%%%%%%%%%%%%%%%%%%%%%%%%%%%%%%%%%%%%%%%%%%%%%%%%%%%%%%%%%%%%%%%%
%%%%%%%%%%%%%%%%%%%%%%%%%%%%%% INPUT %%%%%%%%%%%%%%%%%%%%%%%%%%%%%%%%%%%%%%
%%%%%%%%%%%%%%%%%%%%%%%%%%%%%%%%%%%%%%%%%%%%%%%%%%%%%%%%%%%%%%%%%%%%%%%%%%%
     % Threshold                :  Defined threshold to adapt the bin sizes
     % Trace_uniform_bin        :  Uniformly binned trace
     % Trace_single_photon      :  Original single photon arrival time
     % Deltat                   :  Variance of the previous filter
     
%%%%%%%%%%%%%%%%%%%%%%%%%%%%%%%%%%%%%%%%%%%%%%%%%%%%%%%%%%%%%%%%%%%%%%%%%%%
%%%%%%%%%%%%%%%%%%%%%%%%%%%%%% OUTPUT %%%%%%%%%%%%%%%%%%%%%%%%%%%%%%%%%%%%%
%%%%%%%%%%%%%%%%%%%%%%%%%%%%%%%%%%%%%%%%%%%%%%%%%%%%%%%%%%%%%%%%%%%%%%%%%%%
     % Time                     :  Time vector
     % Trace                    :  Adaptive binned trace
     
     
% Pre-calculation of trace length
Trace_len=length(Trace_uniform_bin);

% Pre-specify the Time vector
Time=ones(1,Trace_len);

% Set the end loop label for while loop
end_loop=0;

% Starting the while loop on the adaptive bin sizes
 while end_loop==0

% Set the counter on adaptive bins at step kk to zero
i=1;

% Set the end loop label for bin sizes while loop
end_loop_bin=0;

% Starting the while loop on bin sizes
    while end_loop_bin==0
        
        % Combine to bins if one of them is less than the threshold
        if  Trace_uniform_bin(i)<Threshold
            if sum(Time(1:i))>=Trace_len
               end_loop_bin=1;
            else
                Trace_uniform_bin(i+1)=[];
                Time(i)=Time(i)+Time(i+1);
                Time(i+1)=[];
            end
        end
        
        % End the inner while loop if we reached to the end of the signal
        if sum(Time(1:i))>=Trace_len
           end_loop_bin=1;
        end
        
        % count the number of the bins
        i=i+1;
        
    % End of the inner while loop   
    end
    
    
    
% count the number of the photons in the new bins
    [Trace_uniform_bin,~] = histcounts(Trace_single_photon,cumsum([0 Time].*Deltat) );
    
    
  Time=Time(1:end-1);
  Trace_len=sum(Time);
  Trace_uniform_bin=Trace_uniform_bin(1:end-1);
  
% End the outer while loop if there were no bins with less photons than the threshold
    if length(find(Trace_uniform_bin<Threshold))==0
       end_loop=1;
    end

% End of the outer while loop
 end
 

%%%%%%%%%%%%%%%%%%%%%%%%%%%%%% Outputs %%%%%%%%%%%%%%%%%%%%%%%%%%%%%%%%%%%%
 Trace=Trace_uniform_bin;
 Time=Time*Deltat;
end
```
