## Supplementary material for "An Alternative Framework for Fluorescence Correlation Spectroscopy": Source Code: EKF.m

```
function [XX,VT] = EKF( x        , c         , constant1   , I    , ...
                        Y        , Z         , observation , vdif , ...
                        f)

%%%%%%%%%%%%%%%%%%%%%%%%%%%%%%%%%%%%%%%%%%%%%%%%%%%%%%%%%%%%%%%%%%%%%%%%%%%
%%%%%%%%%%%%%%%%%%%%%%%%%%%%%% INPUT %%%%%%%%%%%%%%%%%%%%%%%%%%%%%%%%%%%%%%
%%%%%%%%%%%%%%%%%%%%%%%%%%%%%%%%%%%%%%%%%%%%%%%%%%%%%%%%%%%%%%%%%%%%%%%%%%%
     % x                        :  Mean value of the previous filter
     % c                        :  Variance of the previous filter
     % constant1                :  Effect of the other molecules on the observation
     % I                        :  Molecule emission rate
     % Y                        :  Y dimension of the molecule (It could be X or Y dimension depends on the FFBS function)
     % Z                        :  Z dimension of the molecule (It could be X or Y dimension depends on the FFBS function)
     % observation              :  Observation (Transformed trace by Anscombe)
     % vdif                     :  Variance of the kinetics (vdif=2* D* Delta_t)
     
%%%%%%%%%%%%%%%%%%%%%%%%%%%%%%%%%%%%%%%%%%%%%%%%%%%%%%%%%%%%%%%%%%%%%%%%%%%
%%%%%%%%%%%%%%%%%%%%%%%%%%%%%% OUTPUT %%%%%%%%%%%%%%%%%%%%%%%%%%%%%%%%%%%%%
%%%%%%%%%%%%%%%%%%%%%%%%%%%%%%%%%%%%%%%%%%%%%%%%%%%%%%%%%%%%%%%%%%%%%%%%%%%
     % XX                       :  Mean value of the filter
     % VT                       :  Variance of the filter


     constant =  I*f(x,Y,Z)                                               ;
     m        =  constant1+constant                                       ;
     A        =  sqrt( m + 0.375 )                                        ;
     B        =  sqrt( m  )                                               ;
     d        =  x*constant*( -4/A -0.5/(B^3)  )  +eps                    ;
                
% Prediction of the variace
    VTT    =  c+vdif                                                      ;
% Variance of the filter
    VT     =  VTT/(VTT*(d^2)+1)                                           ;
% Mean value of the filter
    XX     =  x + d  *VT*( observation -  2*A +0.25/B  )                  ;
    
    
end
```
