## Supplementary material for "An Alternative Framework for Fluorescence Correlation Spectroscopy": Source Code: EKF_2DGL_3.m

```
function [XX,VT] = EKF_2DGL_3( x        , c         , constant1   , I    , ...
                               Y        , Z         , observation , vdif , ...
                               f)

%%%%%%%%%%%%%%%%%%%%%%%%%%%%%%%%%%%%%%%%%%%%%%%%%%%%%%%%%%%%%%%%%%%%%%%%%%%
%%%%%%%%%%%%%%%%%%%%%%%%%%%%%% INPUT %%%%%%%%%%%%%%%%%%%%%%%%%%%%%%%%%%%%%%
%%%%%%%%%%%%%%%%%%%%%%%%%%%%%%%%%%%%%%%%%%%%%%%%%%%%%%%%%%%%%%%%%%%%%%%%%%%
     % x                        :  Mean value of the previous filter
     % c                        :  Variance of the previous filter
     % constant1                :  Effect of the other molecules on the observation
     % I                        :  Molecule emission rate
     % Y                        :  Y dimension of the molecule (It could be X or Y dimension depends on the FFBS function)
     % Z                        :  Z dimension of the molecule (It could be X or Y dimension depends on the FFBS function)
     % observation              :  Observation (Transformed trace by Anscombe)
     % vdif                     :  Variance of the kinetics (vdif=2* D* Delta_t)
     
%%%%%%%%%%%%%%%%%%%%%%%%%%%%%%%%%%%%%%%%%%%%%%%%%%%%%%%%%%%%%%%%%%%%%%%%%%%
%%%%%%%%%%%%%%%%%%%%%%%%%%%%%% OUTPUT %%%%%%%%%%%%%%%%%%%%%%%%%%%%%%%%%%%%%
%%%%%%%%%%%%%%%%%%%%%%%%%%%%%%%%%%%%%%%%%%%%%%%%%%%%%%%%%%%%%%%%%%%%%%%%%%%
     % XX                       :  Mean value of the filter
     % VT                       :  Variance of the filter


    const =  I.*f(x,Y,Z)                                                  ;
    m     =  constant1 + const                                            ;
    A     =  sqrt( m + 0.375 )                                            ;
    B     =  sqrt( m )                                                    ;
    d     = -x*(1 - 2*(Z^2+Y^2)./((1 + x^2)^3) )*const*(2/A + 0.25/(B^3)) ;
   
% Prediction of the variace
    VTT   =  c + vdif                                                     ;
% Variance of the filter
    VT    =  VTT/( VTT*(d^2) + 1 )                                        ;
% Mean value of the filter
    XX    =  x + d*VT*( observation -  2*A + 0.25/B  )                    ;
    
    
end
```
