## Supplementary material for "An Alternative Framework for Fluorescence Correlation Spectroscopy": Source Code: FFBS.m

```
%%%%%%%%%%%%%%%%%%%%%%%%%%%%%%%%%%%%%%%%%%%%%%%%%%%%%%%%%%%%%%%%%%%%%%%%%%%
%%%%%%%%%%%%%%%%%%%%%%%%%% Extended Kalman Filter %%%%%%%%%%%%%%%%%%%%%%%%%
%%%%%%%%%%%%%%%%%%%%%%%%%%%%%%%%%%%%%%%%%%%%%%%%%%%%%%%%%%%%%%%%%%%%%%%%%%%

function[initial_positions]  =  FFBS( ...
    ...
    ...
observation      , I                  , I_b       , vdif      , ...
Number_molecules , initial_positions  , num_par   , mu_prior  , ...
var_prior        , b                  , sign_siz  , PSF_func  , ...
Filter_typ       , yy                 , p         , f_3DG     , ...
f_2DGL_D1_2      , f_2DGL_D3          , f_2DGC    , AAA       ) 

%%%%%%%%%%%%%%%%%%%%%%%%%%%%%%%%%%%%%%%%%%%%%%%%%%%%%%%%%%%%%%%%%%%%%%%%%%%
%%%%%%%%%%%%%%%%%%%%%%%%%%%%%% INPUT %%%%%%%%%%%%%%%%%%%%%%%%%%%%%%%%%%%%%%
%%%%%%%%%%%%%%%%%%%%%%%%%%%%%%%%%%%%%%%%%%%%%%%%%%%%%%%%%%%%%%%%%%%%%%%%%%%
     % observation              :  Anscombe transformed trace
     % I                        :  Vector of molecle photon emission rates at any time step
     % I_b                      :  Vector of background photon emission rates at any time step
     % vdif                     :  Vector of the kinetic variances at each time step and coordinates
     % Number_molecules         :  Maximum number of molecules
     % initial_positions        :  Locations of all molecules
     % num_par                  :  The tag for targeted molecule
     % mu_prior                 :  Mean value for prior of X0 in normalized scale for x, y and z coordintes
     % var_prior                :  Variance for prior of X0 in normalized scale for x, y and z coordintes
     % b                        :  Loads
     % sign_siz                 :  length of the trace
     % PSF_func                 :  Point Spread Function type ( 'Gaussian' or 'Lorentzian Gaussian')
     % Filter_typ               :  Filter Type ('EKF' or 'UKF')
     % yy                       :  Location of the sigma points used in UKF
     % p                        :  Weight of the sigma points used in UKF
     
%%%%%%%%%%%%%%%%%%%%%%%%%%%%%%%%%%%%%%%%%%%%%%%%%%%%%%%%%%%%%%%%%%%%%%%%%%%
%%%%%%%%%%%%%%%%%%%%%%%%%%%%%% OUTPUT %%%%%%%%%%%%%%%%%%%%%%%%%%%%%%%%%%%%%
%%%%%%%%%%%%%%%%%%%%%%%%%%%%%%%%%%%%%%%%%%%%%%%%%%%%%%%%%%%%%%%%%%%%%%%%%%%
     % initial_positions        :  Updated position of targeted molecules


%%%%%%%%%%%%%%%%%%%%%%%%%%%%%%%%%%%%%%%%%%%%%%%%%%%%%%%%%%%%%%%%%%%%%%%%%%%
%%%%%%%%%%%%%%%%%%%%%%%%%%  Precalculations %%%%%%%%%%%%%%%%%%%%%%%%%%%%%%%
%%%%%%%%%%%%%%%%%%%%%%%%%%%%%%%%%%%%%%%%%%%%%%%%%%%%%%%%%%%%%%%%%%%%%%%%%%%

vt              =  nan(1,sign_siz+1)   ;  % Forward Filter Varinace
x               =  nan(1,sign_siz+1)   ;  % Mean value of the Forward Filter

% Pre-calculations 
nump=num_par*3;

% Calculate the effect of the other molecule on the likelihood
b(num_par)=0;


% Choose the function for PSF
switch  PSF_func
    
%%%%%%%%%%%%%%%%%%%%%%%%%%%%%%%%%%%%%%%%%%%%%%%%%%%%%%%%%%%%%%%%%%%%%%%%%%%
%%%%%%%%%%%%%%%%%%%%%%%%%%  3D Gaussian PSF  %%%%%%%%%%%%%%%%%%%%%%%%%%%%%%
%%%%%%%%%%%%%%%%%%%%%%%%%%%%%%%%%%%%%%%%%%%%%%%%%%%%%%%%%%%%%%%%%%%%%%%%%%%
    case 1   
          constant1 = I_b +I .* ( b * exp(-2*(initial_positions(AAA(1,:),:).^2 ...
                                             +initial_positions(AAA(2,:),:).^2 ...
                                             +initial_positions(AAA(3,:),:).^2 )));                         
          AA=[ 1 , 2 , 3 ; ...
               2 , 1 , 3 ; ...
               3 , 1 , 2];
%%%%%%%%%%%%%%%%%%%%%%%%%%%%%%%%%%%%%%%%%%%%%%%%%%%%%%%%%%%%%%%%%%%%%%%%%%%
%%%%%%%%%%%%%%%%%%%%%  nn=1,2,3 as X, Y , Z  %%%%%%%%%%%%%%%%%%%%%%%%%%%%%%    
%%%%%%%%%%%%%%%%%%%%%%%%%%%%%%%%%%%%%%%%%%%%%%%%%%%%%%%%%%%%%%%%%%%%%%%%%%%
for nn = 1:3  
    
    
x(1)    =  ((-1)^(floor(2*rand())))*mu_prior(AA(nn,1))       ;  % Set the initial Mean value of the Forward Filter
vt(1)   =  var_prior(AA(nn,1))      ;  % Set the initial Variance of the Forward Filter

%%%%%%%%%%%%%%%%%%%%%%%%%%% Forward Filter %%%%%%%%%%%%%%%%%%%%%%%%%%%%%%%%
% Choose the filter type
switch Filter_typ
    case 1    % EKF
         for t=1:sign_siz  
             [x(t+1),vt(t+1)] = EKF( x(t)                                      , ...
                                     vt(t)                                     , ...
                                     constant1(t)                              , ...
                                     I(t)                                      , ...
                                     initial_positions(nump-(3-AA(nn,2)) ,t)   , ...
                                     initial_positions(nump-(3-AA(nn,3)) ,t)   , ...
                                     observation(t)                            , ...
                                     vdif(AA(nn,1),t)                          , ...
                                     f_3DG                                    );
         end
    case 2    % UKF
         for t=1:sign_siz  
             [x(t+1),vt(t+1)] = UKF( x(t)                                      , ...
                                     vt(t)                                     , ...
                                     constant1(t)                              , ...
                                     I(t)                                      , ...
                                     initial_positions(nump-(3-AA(nn,2)) ,t)   , ...
                                     initial_positions(nump-(3-AA(nn,3)) ,t)   , ...
                                     observation(t)                            , ...
                                     vdif(AA(nn,1),t)                          , ...
                                     yy                                        , ...
                                     p                                         , ...
                                     f_3DG                                    );
         end
end
        

%%%%%%%%%%%%%%%%%%%%%%%%% Backward sampling %%%%%%%%%%%%%%%%%%%%%%%%%%%%%%%

% Sample the last point directly from the likelihood(filter)
if  rand()>0.5
    initial_positions(nump-(3-AA(nn,1)),sign_siz)  =  x(sign_siz+1) + sqrt(vt(sign_siz+1))*randn()         ;
else
    initial_positions(nump-(3-AA(nn,1)),sign_siz)  = -x(sign_siz+1) + sqrt(vt(sign_siz+1))*randn()         ;
end


% Sample the rest of the points from the likelihood times the kinetics
vtt  = vt(1:sign_siz)+vdif(AA(nn,1),:)  ;
vvt  = sqrt(vt(1:sign_siz).*vdif(AA(nn,1),:)./vtt)    ;
for i = sign_siz:-1:2            
    if rand() < 1/(1+exp(-2*initial_positions(nump-(3-AA(nn,1)),i)*x(i)/vtt(i)))
        initial_positions(nump-(3-AA(nn,1)),i-1) = ...
            ( initial_positions(nump-(3-AA(nn,1)),i)*vt(i)+vdif(AA(nn,1),i)*x(i) )/vtt(i) + vvt(i)*randn()        ;
    else         
        initial_positions(nump-(3-AA(nn,1)),i-1) = ...
            ( initial_positions(nump-(3-AA(nn,1)),i)*vt(i)-vdif(AA(nn,1),i)*x(i) )/vtt(i) + vvt(i)*randn()        ;
    end
    
end
     

end

%%%%%%%%%%%%%%%%%%%%%%%%%%%%%%%%%%%%%%%%%%%%%%%%%%%%%%%%%%%%%%%%%%%%%%%%%%%
%%%%%%%%%%%%%%%%%%%%% Lorentzian-Gaussian PSF %%%%%%%%%%%%%%%%%%%%%%%%%%%%%
%%%%%%%%%%%%%%%%%%%%%%%%%%%%%%%%%%%%%%%%%%%%%%%%%%%%%%%%%%%%%%%%%%%%%%%%%%%
    case 2     
              constant1 = I_b+ I .* (b * (exp(-2*(initial_positions(AAA(1,:),:).^2 ...
                                                 +initial_positions(AAA(2,:),:).^2) ./...
                                               (1+initial_positions(AAA(3,:),:).^2))./...
                                               (1+initial_positions(AAA(3,:),:).^2)) );

          if  rand()<0.5
              AA=[1 , 3 , 2 ; ...
                  2 , 1 , 3 ; ...
                  3 , 2 , 1];
          elseif rand()<0.5
               AA=[3 , 1 , 2 ; ...
                   2 , 3 , 1 ; ...
                   1 , 2 , 3];
          else
               AA=[2 , 1 , 3 ; ...
                   3 , 2 , 1 ; ...
                   1 , 3 , 2];
          end
                                         
                                           
%%%%%%%%%%%%%%%%%%%%%%%%%%%%%%%%%%%%%%%%%%%%%%%%%%%%%%%%%%%%%%%%%%%%%%%%%%%
%%%%%%%%%%%%%%%%%%%%%  nn=1,2,3 as X, Y , Z  %%%%%%%%%%%%%%%%%%%%%%%%%%%%%%    
%%%%%%%%%%%%%%%%%%%%%%%%%%%%%%%%%%%%%%%%%%%%%%%%%%%%%%%%%%%%%%%%%%%%%%%%%%%
for nn=1:3  
    
    
x(1)    =  mu_prior(AA(nn,1))       ;  % Set the initial Mean value of the Forward Filter
vt(1)   =  var_prior(AA(nn,1))      ;  % Set the initial Variance of the Forward Filter


%%%%%%%%%%%%%%%%%%%%%%%%%%% Forward Filter %%%%%%%%%%%%%%%%%%%%%%%%%%%%%%%%
% Choose the filter type
switch AA(nn,1)
       case 3
            switch Filter_typ
                   case 1    % EKF
                        for t=1:sign_siz  
                            [x(t+1),vt(t+1)] = EKF_2DGL_3( x(t)                                      , ...
                                                           vt(t)                                     , ...
                                                           constant1(t)                              , ...
                                                           I(t)                                      , ...
                                                           initial_positions(nump-(3-AA(nn,2)) ,t)   , ...
                                                           initial_positions(nump-(3-AA(nn,3)) ,t)   , ...
                                                           observation(t)                            , ...
                                                           vdif(AA(nn,1),t)                          , ...
                                                           f_2DGL_D3                                );
                        end
                   case 2    % UKF
                        for t=1:sign_siz  
                            [x(t+1),vt(t+1)] = UKF_2DGL_3( x(t)                                      , ...
                                                           vt(t)                                     , ...
                                                           constant1(t)                              , ...
                                                           I(t)                                      , ...
                                                           initial_positions(nump-(3-AA(nn,2)) ,t)   , ...
                                                           initial_positions(nump-(3-AA(nn,3)) ,t)   , ...
                                                           observation(t)                            , ...
                                                           vdif(AA(nn,1),t)                          , ...
                                                           yy                                        , ...
                                                           p                                         , ...
                                                           f_2DGL_D3                                );
                        end
            end
    otherwise 
            switch Filter_typ
                   case 1   % EKF
                        for t=1:sign_siz  
                            [x(t+1),vt(t+1)] = EKF_2DGL_12( x(t)                                      , ...
                                                            vt(t)                                     , ...
                                                            constant1(t)                              , ...
                                                            I(t)                                      , ...
                                                            initial_positions(nump-(3-AA(nn,2)) ,t)   , ...
                                                            initial_positions(nump-(3-AA(nn,3)) ,t)   , ...
                                                            observation(t)                            , ...
                                                            vdif(AA(nn,1),t)                          , ...
                                                            f_2DGL_D1_2                                );
                        end
                       
                   case 2   % UKF
                        for t=1:sign_siz  
                            [x(t+1),vt(t+1)] = UKF( x(t)                                      , ...
                                                    vt(t)                                     , ...
                                                    constant1(t)                              , ...
                                                    I(t)                                      , ...
                                                    initial_positions(nump-(3-AA(nn,2)) ,t)   , ...
                                                    initial_positions(nump-(3-AA(nn,3)) ,t)   , ...
                                                    observation(t)                            , ...
                                                    vdif(AA(nn,1),t)                          , ...
                                                    yy                                        , ...
                                                    p                                         , ...
                                                    f_2DGL_D1_2                              );
                        end
            end
end
        

%%%%%%%%%%%%%%%%%%%%%%%%% Backward sampling %%%%%%%%%%%%%%%%%%%%%%%%%%%%%%%

% Sample the last point directly from the likelihood(filter)
if  rand()>0.5
    initial_positions(nump-(3-AA(nn,1)),sign_siz)  =  x(sign_siz+1) + sqrt(vt(sign_siz+1))*randn()         ;
else
    initial_positions(nump-(3-AA(nn,1)),sign_siz)  = -x(sign_siz+1) + sqrt(vt(sign_siz+1))*randn()         ;
end


% Sample the rest of the points from the likelihood times the kinetics
vtt  = vt(1:sign_siz)+vdif(AA(nn,1),:)  ;
vvt  = sqrt(vt(1:sign_siz).*vdif(AA(nn,1),:)./vtt)    ;
for i = sign_siz:-1:2            
    if rand() < 1/(1+exp(-2*initial_positions(nump-(3-AA(nn,1)),i)*x(i)/vtt(i)))
        initial_positions(nump-(3-AA(nn,1)),i-1) = ...
            ( initial_positions(nump-(3-AA(nn,1)),i)*vt(i)+vdif(AA(nn,1),i)*x(i) )/vtt(i) + vvt(i)*randn()        ;
    else         
        initial_positions(nump-(3-AA(nn,1)),i-1) = ...
            ( initial_positions(nump-(3-AA(nn,1)),i)*vt(i)-vdif(AA(nn,1),i)*x(i) )/vtt(i) + vvt(i)*randn()        ;
    end
    
end
     

end
                                           
                                           
%%%%%%%%%%%%%%%%%%%%%%%%%%%%%%%%%%%%%%%%%%%%%%%%%%%%%%%%%%%%%%%%%%%%%%%%%%%
%%%%%%%%%%%%%%%%%%%%%%%%%% Cylindrical PSF %%%%%%%%%%%%%%%%%%%%%%%%%%%%%%%%
%%%%%%%%%%%%%%%%%%%%%%%%%%%%%%%%%%%%%%%%%%%%%%%%%%%%%%%%%%%%%%%%%%%%%%%%%%%
    otherwise
              constant1 = I_b +I .* ( b * exp(-2*(initial_positions(AAA(1,:),:).^2 ...
                                                 +initial_positions(AAA(2,:),:).^2 )));
                                             
 AA=[1 , 2 ; ...
     2 , 1 ];

%%%%%%%%%%%%%%%%%%%%%%%%%%%%%%%%%%%%%%%%%%%%%%%%%%%%%%%%%%%%%%%%%%%%%%%%%%%
%%%%%%%%%%%%%%%%%%%%%%%  nn=1,2  as X, Y   %%%%%%%%%%%%%%%%%%%%%%%%%%%%%%%%    
%%%%%%%%%%%%%%%%%%%%%%%%%%%%%%%%%%%%%%%%%%%%%%%%%%%%%%%%%%%%%%%%%%%%%%%%%%%
for nn=1:2  
    
    
x(1)    =  mu_prior(AA(nn,1))       ;  % Set the initial Mean value of the Forward Filter
vt(1)   =  var_prior(AA(nn,1))      ;  % Set the initial Variance of the Forward Filter


%%%%%%%%%%%%%%%%%%%%%%%%%%% Forward Filter %%%%%%%%%%%%%%%%%%%%%%%%%%%%%%%%
% Choose the filter type
switch Filter_typ
    
    case 1    % EKF
         for t=1:sign_siz  
             [x(t+1),vt(t+1)] = EKF( x(t)                                      , ...
                                     vt(t)                                     , ...
                                     constant1(t)                              , ...
                                     I(t)                                      , ...
                                     initial_positions(nump-(3-AA(nn,2)) ,t)   , ...
                                     initial_positions(nump              ,t)   , ...
                                     observation(t)                            , ...
                                     vdif(AA(nn,1),t)                          , ...
                                     f_2DGC                                   );
         end
    case 2    % UKF
         for t=1:sign_siz  
             [x(t+1),vt(t+1)] = UKF( x(t)                                      , ...
                                     vt(t)                                     , ...
                                     constant1(t)                              , ...
                                     I(t)                                      , ...
                                     initial_positions(nump-(3-AA(nn,2)) ,t)   , ...
                                     initial_positions(nump              ,t)   , ...
                                     observation(t)                            , ...
                                     vdif(AA(nn,1),t)                          , ...
                                     yy                                        , ...
                                     p                                         , ...
                                     f_2DGC                                   );
         end
end
        

%%%%%%%%%%%%%%%%%%%%%%%%% Backward sampling %%%%%%%%%%%%%%%%%%%%%%%%%%%%%%%

% Sample the last point directly from the likelihood(filter)
if  rand()>0.5
    initial_positions(nump-(3-AA(nn,1)),sign_siz)  =  x(sign_siz+1) + sqrt(vt(sign_siz+1))*randn()         ;
else
    initial_positions(nump-(3-AA(nn,1)),sign_siz)  = -x(sign_siz+1) + sqrt(vt(sign_siz+1))*randn()         ;
end


% Sample the rest of the points from the likelihood times the kinetics
vtt  = vt(1:sign_siz)+vdif(AA(nn,1),:)  ;
vvt  = sqrt(vt(1:sign_siz).*vdif(AA(nn,1),:)./vtt)    ;
for i = sign_siz:-1:2            
    if rand() < 1/(1+exp(-2*initial_positions(nump-(3-AA(nn,1)),i)*x(i)/vtt(i)))
        initial_positions(nump-(3-AA(nn,1)),i-1) = ...
            ( initial_positions(nump-(3-AA(nn,1)),i)*vt(i)+vdif(AA(nn,1),i)*x(i) )/vtt(i) + vvt(i)*randn()        ;
    else         
        initial_positions(nump-(3-AA(nn,1)),i-1) = ...
            ( initial_positions(nump-(3-AA(nn,1)),i)*vt(i)-vdif(AA(nn,1),i)*x(i) )/vtt(i) + vvt(i)*randn()        ;
    end
end

end

%%%%%%%%%%%%%%%%%%%%%%%%%%%%%%%%%%%%%%%%%%%%%%%%%%%%%%%%%%%%%%%%%%%%%%%%%%%
%%%%%%%%%%%%%%%  Z coordinate from random walk  %%%%%%%%%%%%%%%%%%%%%%%%%%%    
%%%%%%%%%%%%%%%%%%%%%%%%%%%%%%%%%%%%%%%%%%%%%%%%%%%%%%%%%%%%%%%%%%%%%%%%%%%
initial_positions(nump,:) =  cumsum([((-1)^(floor(2*rand())))*mu_prior(3) + sqrt(var_prior(3))*randn(),...
                                              sqrt(vdif(3,2:end)).*randn(1,sign_siz-1)],2)                          ;
end


end
```
