## Supplementary material for "An Alternative Framework for Fluorescence Correlation Spectroscopy": Source Code: Gibbs_sampler.m

```
function [ D              , mu                  , mu_back            , b   , ...
           concentration  , acceptance_rate_mu  , acceptance_rate_b  , x   , ...
           q             ] = Gibbs_sampler(...
...
...
Trace              , Time                 , D_alpha_prior  , ...
D_beta_prior       , save_size            , Number_particles , Max_iter       , ...
wxy                , wz                   , mu_proposal      , mu_alpha       , ...
mu_beta            , mu_back_proposal     , mu_back_alpha    , mu_back_beta   , ...
alpha_q            , beta_q               , mu_prior_xy      , mu_prior_z     , ...
var_prior_xy       , var_prior_z          , mu_back          , iter_b_mu      , ...
concen_radious     , x                    , handles          , D              , ...
mu                 , b                    , q                , concentration  , ...
Automatics         , PSF_Func             , Filter_typ       , percentage_dif )

%%%%%%%%%%%%%%%%%%%%%%%%%%%%%%%%%%%%%%%%%%%%%%%%%%%%%%%%%%%%%%%%%%%%%%%%%%%
%%%%%%%%%%%%%%%%%%%%%%%%%%%%%% INPUT %%%%%%%%%%%%%%%%%%%%%%%%%%%%%%%%%%%%%%
%%%%%%%%%%%%%%%%%%%%%%%%%%%%%%%%%%%%%%%%%%%%%%%%%%%%%%%%%%%%%%%%%%%%%%%%%%%
     % Trace_uniform_bin             :  Uniformly binned traces
     % Trace_singel_photon           :  Original traces (Single photon arrival times)
     % Delta_t                       :  Data acquisition time
     % D_alpha_prior                 :  Alpha parameter of inverse gamma prob. dis. for prior related to Diffusion
     % D_beta_prior                  :  Beta parameter of inverse gamma prob. dis. for prior related to Diffusion
     % Max_iter                      :  Maximum number of MCMC chains
     % wxy                           :  Minor Semi-axis of the confocal volume (1/e of the laser beam wast)
     % wz                            :  Major Semi-axis of the confocal volume
     % mu_proposal                   :  Proposal parameter for molecule photon emission rate
     % mu_alpha                      :  Alpha parameter of molecule photon emission rate prior
     % mu_beta                       :  Beta parameter of molecule photon emission rate prior
     % mu_back_proposal              :  Proposal parameter for background photon emission rate
     % mu_back_alpha                 :  Alpha parameter of backgrouund photon emission rate prior
     % mu_back_beta                  :  Beta parameter of background photon emission rate  prior
     % alpha_q                       :  Alpha parameter for weight of load b     ( Gamma distribution)
     % beta_q                        :  Beta parameter for weight of load b      ( Gamma distribution)
     % mu_prior_xy                   :  Mean value of X0 prior in normalized scale for x and y coordintes
     % mu_prior_z                    :  Mean value of X0 prior in normalized scale for z coordinte
     % var_prior_xy                  :  Variance of X0 prior in normalized scale for x and y coordintes
     % var_prior_z                   :  Variance of X0 prior in normalized scale for z coordinte
     % iter_b_mu                     :  Number of iterations for metropolis-hastings step
     % concen_radious                :  Define normalize distances to measure the concentrations
     % D                             :  Sampled diffusion coefficient
     % x                             :  Updated locations of molecules
     % handles                       :  GUI information
     % mu                            :  Sampled molecule emission rate
     % mu_back                       :  Sampled background emission rate
     % b                             :  Sampled loads
     % q                             :  Sampled weights of the loads
     % concentration                 :  Updated concentrations of molecules in the ROI
     % Automatics                    :  Automatic command
     % PSF_Func                      :  Point Spread Function type ( 'Gaussian' or 'Lorentzian Gaussian')
     % Filter_typ                    :  Filter Type ('EKF' or 'UKF')
     % percentage_dif                :  Convergance threshold
     
     
%%%%%%%%%%%%%%%%%%%%%%%%%%%%%%%%%%%%%%%%%%%%%%%%%%%%%%%%%%%%%%%%%%%%%%%%%%%
%%%%%%%%%%%%%%%%%%%%%%%%%%%%%% OUTPUT %%%%%%%%%%%%%%%%%%%%%%%%%%%%%%%%%%%%%
%%%%%%%%%%%%%%%%%%%%%%%%%%%%%%%%%%%%%%%%%%%%%%%%%%%%%%%%%%%%%%%%%%%%%%%%%%%
     % D                             :  An array of sampeled Diffusion coefficients
     % mu                            :  An array of sampeled molecule emission rates
     % mu_back                       :  An array of sampeled background emission rates
     % b                             :  A matrix of sampeled loads
     % q                             :  A matrix of sampeled weights of the loads
     % concentration                 :  Concentration of molecules in ROI
     % acceptance_rate_mu            :  Acceptance rate of the emission rate Metropolis
     % acceptance_rate_b             :  Acceptance rate of the loads Metropolis
     % x                             :  Export the last sampled positions to use as the initial value of next run

     
% Shuffle the random generator
    rng('shuffle')

% Calculate the sigma points and their weights to be used for the unscented Kalman filter (UKF)
    [yy, pp] = nwspgr('GQN', 1 , 13)                                      ;
    yy=yy'                                                                ;
   
% Calculate the number of traces
    Number_trace = length(Trace)                                          ;


% Calculate the length of each trace
    for k=1:Number_trace
       Trace_size(k)   = length(Trace{k})                                 ; 
    end
    
% Normalize the mean value and variance of the prior over initial locations
    mu_prior  = [ mu_prior_xy/wxy  , mu_prior_xy/wxy , mu_prior_z/wz ]                     ;
    var_prior = [ var_prior_xy/wxy^2, var_prior_xy/wxy^2 , var_prior_z/wz^2 ]                  ;

    
% Calculate the size of the previous sample
    OLD_Iteration_size=length(D)                                          ;

% Set the initial iteration number
    i=OLD_Iteration_size                                                  ;
    
% Check the Automatic command
    if  Automatics==1
        save_size=0                                                       ;
        Max_iter=2                                                        ;
    end
    
% Set the parameters' vectors and initial values
    acceptance_rate_mu    = 0                                             ;
    acceptance_rate_b     = 0                                             ;


% Transform the original trace to Guassian by using Anscombe Transformation
observation         = Anscombe(Trace)                                     ;

% Set the end loop command to zero
endloop=0                                                                 ;


% Precalculation of normalized distances lengths
len_concen_radius = length(concen_radious)                                ;


if  Automatics==0
% Initialize the time bar
    cla(handles.axes23)
    set(handles.axes23,'Units','pixels','XLim',[0 1],'YLim',[0 1],...
        'XTick',[],'YTick',[],'Color','w','XColor','w','YColor','w')      ;
    
    patch(handles.axes23,[0 0 0 0],[0 1 1 0],'b')                         ;
    handles.text80.String ='Initializing ...'                             ; 
    time_loop             = 0                                             ;
    save_bar              = 1                                             ;
    drawnow
    tic
end
    
% Definition of the PSFs
f_3DG       =@(x,y,z) exp( -2*(x.^2 + y.^2 + z.^2))                       ;  % 3DG
f_2DGL_D1_2 =@(x,y,z) exp( -2*(x.^2 + y.^2)./(1 + z.^2))./(1 + z.^2)      ;  % 2DG-Lorentzian, x and y demensions
f_2DGL_D3   =@(x,y,z) exp( -2*(y.^2 + z.^2)./(1 + x.^2))./(1 + x.^2)      ;  % 2DG-Lorentzian, z demension
f_2DGC      =@(x,y,z) exp( -2*(x.^2 + y.^2))                              ;  % 2DG-cylindrical


% Pre calculation
AAA(1,:) = 1:3:3*Number_particles                                         ;
AAA(2,:) = 2:3:3*Number_particles                                         ;
AAA(3,:) = 3:3:3*Number_particles                                         ;

% Start the Gibbs sampling iteration loop
while endloop==0

% iteration counter
i = i+1                                                                   ;


% Initiate square dicplacements of the molecules to zero
kk_diffusion=0                                                            ;

% Start the FOR loop on the number of imported signals
for k=1:Number_trace
    
    % Pre-calculation of the variance of the kinetics at each time step and coordiante
    
    vdif{k}      =   2*D(i-1)*[0 Time{k}(2:end)]./[wxy^2 ; wxy^2 ; wz^2]      ; 
    
  
    % Pre-calculation of the molecule and background emission rates times data acquisition time (Time)
    I_b          =  mu_back(i-1) * Time{k}                                ; 
    I            =  mu(i-1)      * Time{k}                                ; 

    % Start the FOR loop on the number of molecules
    for num_par=1:Number_particles
        
        % Check if the molecule num_par is active (b=1)
        if  b(k,num_par,i-1)==1
%%%%%%%%%%%%%%%%%%%%%%%%%%%%%%%%%%%%%%%%%%%%%%%%%%%%%%%%%%%%%%%%%%%%%%%%%%%
% Sampling ACTIVE trajectories for particle num_par in  imported signal k %
%%%%%%%%%%%%%%%%%%%%%%%%%%%%%%%%%%%%%%%%%%%%%%%%%%%%%%%%%%%%%%%%%%%%%%%%%%%
            x{k}  = FFBS( observation{k}    , I           , I_b           , vdif{k}   , ...
                          Number_particles  , x{k}        , num_par       , mu_prior  , ...
                          var_prior         , b(k,:,i-1)  , Trace_size(k) , PSF_Func  , ...
                          Filter_typ        , yy          , pp            , f_3DG     , ...
                          f_2DGL_D1_2       , f_2DGL_D3   , f_2DGC        , AAA);
        end
    end


%%%%%%%%%%%%%%%%%%%%%%%%%%%%%%%%%%%%%%%%%%%%%%%%%%%%%%%%%%%%%%%%%%%%%%%%%%%
%%%%%%%% Sampling INACTIVE trajectories for imported signal k  %%%%%%%%%%%%
%%%%%%%%%%%%%%%%%%%%%%%%%%%%%%%%%%%%%%%%%%%%%%%%%%%%%%%%%%%%%%%%%%%%%%%%%%%
    x{k}  = Inactive_molecules( x{k}      , Trace_size(k) , Time{k} , b(k,:,i-1) , ...
                                mu_prior  , var_prior     , wxy     , wz         , ...
                                D(i-1)   );


%%%%%%%%%%%%%%%%%%%%%%%%%%%%%%%%%%%%%%%%%%%%%%%%%%%%%%%%%%%%%%%%%%%%%%%%%%%
%%% Compute concetration of the molecules in imported signal k in the ROI %
%%%%%%%%%%%%%%%%%%%%%%%%%%%%%%%%%%%%%%%%%%%%%%%%%%%%%%%%%%%%%%%%%%%%%%%%%%%
    xx=x{k};
    
       switch  PSF_Func
            case 1 % 3D Gaussian
                  Constant{k} =exp(-2*(xx(AAA(1,:),:).^2 +xx(AAA(2,:),:).^2 +xx(AAA(3,:),:).^2) );
                               
                  for l=1:len_concen_radius
          
                      concentration(k,i-(floor((i-1)/10000)*10000),1:Trace_size(k),l) =...
                          ...
                      b(k,:,i-1)*(sqrt( (xx(AAA(1,:),:).^2+xx(AAA(2,:),:).^2+xx(AAA(3,:),:).^2 ))<=concen_radious(l));
                  end 
                  
            case 2 % 2D Gaussian-Lorentzian
                  Constant{k} = exp(-2*(xx(AAA(1,:),:).^2+xx(AAA(2,:),:).^2)./(1+xx(AAA(3,:),:).^2))./(1+xx(AAA(3,:),:).^2);
                                                             
                  for l=1:len_concen_radius
                      concentration(k,i-(floor((i-1)/10000)*10000),1:Trace_size(k),l) =...
                          ...
                      b(k,:,i-1)*(sqrt( 0.5.*log(1+xx(AAA(3,:),:).^2)+((xx(AAA(1,:),:).^2)+(xx(AAA(2,:),:).^2))./(1+xx(AAA(3,:),:).^2) )<=concen_radious(l));
                  end
                  
       otherwise   % 2D Gaussian-Cylindrical
                  Constant{k} =exp(-2*(xx(AAA(1,:),:).^2 +xx(AAA(2,:),:).^2 ) );
                  
                  for l=1:len_concen_radius
                  concentration(k,i-(floor((i-1)/10000)*10000),1:Trace_size(k),l) =...
                          ...
                      b(k,:,i-1)*(sqrt( (xx(AAA(1,:),:).^2 + xx(AAA(2,:),:).^2 ))<=concen_radious(l));
                  end
                                     
       end
   
   

% Calculate square diplacements of the molecules for each imported signal
% We will use it to sample the diffusion coefficient
   kk_diffusion=kk_diffusion+(repmat([wxy wxy wz].^2,1,Number_particles)*...
                           ((xx(:,2:end-1)-xx(:,3:end)).^2))*(Time{k}(3:end).^-1)';
end


%%%%%%%%%%%%%%%%%%%%%%%%%%%%%%%%%%%%%%%%%%%%%%%%%%%%%%%%%%%%%%%%%%%%%%%%%%%
%%%%%%%%%%%%%%%%%%%%% Sample the diffusion coefficient %%%%%%%%%%%%%%%%%%%%
%%%%%%%%%%%%%%%%%%%%%%%%%%%%%%%%%%%%%%%%%%%%%%%%%%%%%%%%%%%%%%%%%%%%%%%%%%%
  D(i)   =    1/gamrnd(D_alpha_prior+((sum(Trace_size-2))*1.5*Number_particles) , ...
                       1/( D_beta_prior+0.25*kk_diffusion ) ); 


               
              
%%%%%%%%%%%%%%%%%%%%%%%%%%%%%%%%%%%%%%%%%%%%%%%%%%%%%%%%%%%%%%%%%%%%%%%%%%%%%%%%%%%%%%%%%%%%%%%%
% Sample the loads, hyper prior on the loads, molecule and background photon emission rates %%%%
%%%%%%%%%%%%%%%%%%%%%%%%%%%%%%%%%%%%%%%%%%%%%%%%%%%%%%%%%%%%%%%%%%%%%%%%%%%%%%%%%%%%%%%%%%%%%%%

% Set the new samples equal to the old ones
mu(i)        = mu(i-1)      ;
mu_back(i)   = mu_back(i-1) ;
b(:,:,i)     = b(:,:,i-1)   ;

% Since we sample mu and mu_back through a metropolis-hastings, we sample
% them several times to have a better mixture.


   
       for j=1:iter_b_mu
           
           % Update the emission rates
           [ mu(i)  , mu_back(i) ,acceptance_rate_mu ] = Sample_emission_rates( ...
           ...
           ...
           mu_proposal   , mu_alpha            , mu_beta          , mu(i)          , ...
           Trace_size    , acceptance_rate_mu  , observation      , mu_back(i)     , ...
           Time          , b(:,:,i)            , mu_back_proposal , mu_back_alpha  , ...
           mu_back_beta  , Constant            , Filter_typ      );
             
           %%%%%%%%%%%%%%%%%%%%%%%%%%%%%%%%%%%%%%%%%%%%%%%%%%%%%%%%%%%%%%%%%%%%%%%%%%%
           %%%%%%%%%%%%%%% Update the weight over loads (hyper priors) %%%%%%%%%%%%%%%
           %%%%%%%%%%%%%%%%%%%%%%%%%%%%%%%%%%%%%%%%%%%%%%%%%%%%%%%%%%%%%%%%%%%%%%%%%%%

           
           
           q(:,:,i) = betarnd(...
                   alpha_q/Number_particles+b(:,:,i),...
                   (beta_q*(Number_particles-1)/Number_particles)+1-b(:,:,i)...
                   );
          
           %%%%%%%%%%%%%%%%%%%%%%%%%%%%%%%%%%%%%%%%%%%%%%%%%%%%%%%%%%%%%%%%%%%%%%%%%%%
           %%%%%%%%%%%%%%%%%%%%%%%%%% Update the loads  %%%%%%%%%%%%%%%%%%%%%%%%%%%%%%
           %%%%%%%%%%%%%%%%%%%%%%%%%%%%%%%%%%%%%%%%%%%%%%%%%%%%%%%%%%%%%%%%%%%%%%%%%%%
           for k=1:Number_trace

               [ b(k,:,i) , acceptance_rate_b ] = Sample_Loads( ...
                ...
                ...
               b(k,:,i)     , mu(i)             , observation{k}  , acceptance_rate_b  , ...
               Time{k}      , Number_particles  , mu_back(i)      , q(k,:,i)           , ...
               Constant{k} ); 
           end
       end

       
%%%%%%%%%%%%%%%%%%%%%%%%%%%%%%%%%%%%%%%%%%%%%%%%%%%%%%%%%%%%%%%%%%%%%%%%%%% 
%%%%%%%%%%%%%%%%%%%%%%% Save and plot the data %%%%%%%%%%%%%%%%%%%%%%%%%%%%
%%%%%%%%%%%%%%%%%%%%%%%%%%%%%%%%%%%%%%%%%%%%%%%%%%%%%%%%%%%%%%%%%%%%%%%%%%% 

       if find((OLD_Iteration_size+floor((1:1:save_size).*Max_iter./save_size))==i)>=1
                  
                 % Plot the remaining time
                 if  Automatics==0
                     time_loop=time_loop+toc;
                     remaining_time=floor((time_loop/(save_bar.*Max_iter./save_size))*((save_size-save_bar).*Max_iter./save_size)*10)/10;
                 
                     handles.text80.String=['Remaining time = ', num2str(remaining_time),' s'];
                     patch(handles.axes23,[0 0 save_bar/save_size save_bar/save_size],[0 1 1 0],'b');
                     save_bar=save_bar+1;
                     tic
                 end
           
           
                 % Calculate the acceptance rates
                 acceptance_rate_muu  = acceptance_rate_mu / (iter_b_mu*(i-1))  ;
                 acceptance_rate_bb   = acceptance_rate_b  / (iter_b_mu*Number_trace*(i-1))  ;

                 % Save the results based on the save size
                 save('results' , 'D'                   , 'mu'                 , 'mu_back'   ...
                                , 'b'                   , 'concentration'      , 'x'         ...
                                , 'acceptance_rate_muu' , 'acceptance_rate_bb' , 'q'         ...
                                , 'Time'                , 'Trace'              )
                          
                 % Plot the sampler results at each save size interval
                 if  handles.pushbutton8.Value==1   ||  handles.pushbutton20.Value==1     
                 plot(handles.axes8,1:1:i,D(1:i))
                 xlabel(handles.axes8,'Iteration')
                 ylabel(handles.axes8,'log_{10} Diff. coef. (\mum^2/s)')
                 xlim(handles.axes8,[0 i])
                 ylim(handles.axes8,[0 max(D(1:i))])
                 set(handles.axes8,'yscale','log') 
                  
                 for k=1:Number_trace
                     plot(handles.axes15,1:1:i,sum(reshape(b(k,:,:),size(b,2),size(b,3)),1))
                     hold (handles.axes15,'on')
                 end
                 hold (handles.axes15,'off')
                 xlabel(handles.axes15,'Iteration')
                 ylabel(handles.axes15,'Number of active molecules')
                 ylim(handles.axes15,[0 Number_particles])
                 xlim(handles.axes15,[0 i])
    
               if  Filter_typ ==1
                   plot(handles.axes16, 1:1:i ,mu(1:i)    )
                   histogram(handles.axes20,mu(1:i),'Normalization','Probability','Orientation','horizontal')
               else
                   plot(handles.axes16, 1:1:i ,mu(1:i)    ,1:1:i ,mu_back(1:i)  )
                   
                   histogram(handles.axes20,mu(1:i),'Normalization','Probability','Orientation','horizontal')
                   hold (handles.axes20,'on')
                   histogram(handles.axes20,mu_back(1:i),'Normalization','Probability','Orientation','horizontal')
                   hold (handles.axes20,'off')
               end
                 
                 xlabel(handles.axes16,'Iteration')
                 if  sum(handles.pushbutton22.String)==sum('Estimate background emission rate (OFF)')
                     ylabel(handles.axes16,{'Molecule photon emission rate' ;'(photons/s)'})
                 else
                     ylabel(handles.axes16,{'Molecule and background';'photon emission rates' ;'(photons/s)'})
                 end
                 xlim(handles.axes16,[0 i])
    
                 
                 
                 bnd = logspace(log10(min(D(1:i)))-1,log10(max(D(1:i)))+1,100);

                 histogram(handles.axes19,D(1:i),bnd,'Normalization','pdf','Orientation','horizontal')
                 handles.axes19.XAxisLocation = 'top';
                 ylim(handles.axes19,[0 max(D)])
                 handles.axes19.YTick = [];
                 handles.axes19.XTick = [];
                 handles.axes19.YAxisLocation = 'right';
                 set(handles.axes19,'yscale','log') 
                 
                 
                 
                for k=1:Number_trace
                    histogram(handles.axes18,sum(reshape(b(k,:,1:i),size(b,2),size(b,3)),1),...
                                       'Normalization','Probability','Orientation','horizontal')
                    hold (handles.axes18,'on')
                end
                 hold (handles.axes18,'off')
                 handles.axes18.YTick = [];
                 handles.axes18.XTick = [];
                 handles.axes18.XAxisLocation = 'top';
                 ylim(handles.axes18,[0 Number_particles])
                 
    
                 handles.axes20.YTick = [];
                 handles.axes20.XTick = [];
                 handles.axes20.XAxisLocation = 'top';
                 ylim(handles.axes20,[0 max(mu(1:i))])
                 
                 
                 drawnow
                 end         
       end

%%%%%%%%%%%%%%%%%%%%%%%%%%%%%%%%%%%%%%%%%%%%%%%%%%%%%%%%%%%%%%%%%%%%%%%%%%% 
%%%%%%%%%%%%%%%%%%%%%%%%%%%%% Convergance %%%%%%%%%%%%%%%%%%%%%%%%%%%%%%%%%
%%%%%%%%%%%%%%%%%%%%%%%%%%%%%%%%%%%%%%%%%%%%%%%%%%%%%%%%%%%%%%%%%%%%%%%%%%% 

% In order to stop the MCMC chain automatically, we applied a convergance 
% method based on the mean values of the last two quarters of the sampled 
% diffusion coefficient and comparing these values with the convergence threshold

% For an adequate sample size, we consider 10000 iterations to be the minimum sample size

if  Automatics==0
    if i==Max_iter+OLD_Iteration_size
       endloop=1;
    end
else
    if  save_size<floor(i/10000)    
        save_size=floor(i/10000);
        handles.edit41.String=save_size;
        Max_iter=i;
        handles.edit39.String=Max_iter;
    end
    if floor(i/10000)==i/10000
       siz=floor(i/4);
       m75=mean(D(2*siz:3*siz)) ;
       m100=mean(D(3*siz:i));
      
       if (abs(m75-m100)/m75)*100<percentage_dif
           endloop=1;
       end  
    end
end
 

end

% End of the iteration loop and show "Done!" as a note.
if  Automatics==0
    handles.text80.String='Done!';
end


% Normalize the acceptance rates
acceptance_rate_mu  = acceptance_rate_mu / (iter_b_mu*(i-length(D)-1))     ;
acceptance_rate_b   = acceptance_rate_b  / (Number_trace*(i-length(D)-1))  ;

end
```
