## Supplementary material for "An Alternative Framework for Fluorescence Correlation Spectroscopy": Source Code: Inactive_molecules.m

```
function[ x ] = Inactive_molecules( ...
...
...
x        , sign_siz   , Time  , b    , ...
mu_prior , var_prior  , wxy              ,wz        , D     )   


%%%%%%%%%%%%%%%%%%%%%%%%%%%%%%%%%%%%%%%%%%%%%%%%%%%%%%%%%%%%%%%%%%%%%%%%%%%
%%%%%%%%%%%%%%%%%%%%%%%%%%%%%% INPUT %%%%%%%%%%%%%%%%%%%%%%%%%%%%%%%%%%%%%%
%%%%%%%%%%%%%%%%%%%%%%%%%%%%%%%%%%%%%%%%%%%%%%%%%%%%%%%%%%%%%%%%%%%%%%%%%%%
     % x                             :  Positions of the molecules (Normalized)
     % sign_siz                      :  Length of the observation trace
     % Time                          :  Data acquisition time
     % b                             :  Loads for active molecules
     % mu_prior                      :  Mean value of initial positions prior  ( Normal distribution)
     % var_prior                     :  Variance of initial positions prior    ( Normal distribution)
     % wxy                           :  Radius of the confocal in xy plane
     % wz                            :  Radius of the confocal in z axis
     % D                             :  Old sampled diffusion coefficent

%%%%%%%%%%%%%%%%%%%%%%%%%%%%%%%%%%%%%%%%%%%%%%%%%%%%%%%%%%%%%%%%%%%%%%%%%%%
%%%%%%%%%%%%%%%%%%%%%%%%%%%%%% OUTPUT %%%%%%%%%%%%%%%%%%%%%%%%%%%%%%%%%%%%%
%%%%%%%%%%%%%%%%%%%%%%%%%%%%%%%%%%%%%%%%%%%%%%%%%%%%%%%%%%%%%%%%%%%%%%%%%%%
     % x                             :  sampled inactive particle trajectories


% Pre-calculation of the kinetic at each time step

% Check if the load on molecule is off
     active_molecules = find(b==0);

% Measure the nmber of inactive molecuules
     size             = length(active_molecules);

% Calculate the kinetics
     kxy              = repmat(sqrt(2*D*Time(2:end))./wxy,size,1)   ;
     kz               = kxy.*(wxy/wz)                               ;


% Sample the locations of molecules through brownian motion
     x(3*active_molecules-2,:) = cumsum([((-1)^(floor(2*rand())))*mu_prior(1) + sqrt(var_prior(1))*randn(size,1),...
                                             kxy.*randn(size,sign_siz-1)],2)                            ;
     x(3*active_molecules-1,:) = cumsum([((-1)^(floor(2*rand())))*mu_prior(1) + sqrt(var_prior(1))*randn(size,1),...
                                             kxy.*randn(size,sign_siz-1)],2)                            ;
     x(3*active_molecules  ,:) = cumsum([((-1)^(floor(2*rand())))*mu_prior(2) + sqrt(var_prior(2))*randn(size,1),...
                                              kz.*randn(size,sign_siz-1)],2)                            ;

end
```
