## Supplementary material for "An Alternative Framework for Fluorescence Correlation Spectroscopy": Source Code: Sample_emission_rates.m

```
function[ MU , MU_BACK ,mu_acceptance_rate ] = Sample_emission_rates( ...
...
...
mu_proposal   ,  mu_alpha            , mu_beta           ,  MU_OLD          , ...
sign_siz      ,  mu_acceptance_rate  , trace             ,  MU_BACK_OLD     , ...
Time          ,  b                   , mu_back_proposal  ,  mu_back_alpha   , ...
mu_back_beta  ,  Constant            , Filter_typ) 


%%%%%%%%%%%%%%%%%%%%%%%%%%%%%%%%%%%%%%%%%%%%%%%%%%%%%%%%%%%%%%%%%%%%%%%%%%%
%%%%%%%%%%%%%%%%%%%%%%%%%%%%%% INPUT %%%%%%%%%%%%%%%%%%%%%%%%%%%%%%%%%%%%%%
%%%%%%%%%%%%%%%%%%%%%%%%%%%%%%%%%%%%%%%%%%%%%%%%%%%%%%%%%%%%%%%%%%%%%%%%%%%
     % mu_proposal                   :  Proposal parameter for MU           ( Gamma distribution)
     % mu_alpha                      :  Alpha parameter of MU prior         ( Gamma distribution)
     % mu_beta                       :  Beta parameter of MU prior          ( Gamma distribution)
     % MU_OLD                        :  Previous sampled molecule photon emission rate
     % sign_siz                      :  Length of the observation signal
     % Number_molecules              :  Number of defined molecules
     % mu_acceptance_rate            :  Previous acceptance rate 
     % trace                         :  Original trace
     % MU_BACK_OLD                   :  Previous background emission rate
     % Time                          :  Data acqisition time
     % b                             :  Loads for active molecules
     % mu_back_proposal              :  Proposal parameter for MU_back           ( Gamma distribution)
     % mu_back_alpha                 :  Alpha parameter of MU_back prior         ( Gamma distribution)
     % mu_back_beta                  :  Beta parameter of MU_back prior          ( Gamma distribution)
     
%%%%%%%%%%%%%%%%%%%%%%%%%%%%%%%%%%%%%%%%%%%%%%%%%%%%%%%%%%%%%%%%%%%%%%%%%%%
%%%%%%%%%%%%%%%%%%%%%%%%%%%%%% OUTPUT %%%%%%%%%%%%%%%%%%%%%%%%%%%%%%%%%%%%%
%%%%%%%%%%%%%%%%%%%%%%%%%%%%%%%%%%%%%%%%%%%%%%%%%%%%%%%%%%%%%%%%%%%%%%%%%%%
     % MU                            :  Updated MU
     % MU_BACK                       :  Updated MU Background
     % mu_acceptance_rate            :  Updated acceptance rate


% Propose MU from a gamma distribution
         MU_NEW        =    (MU_OLD/mu_proposal)*randg(mu_proposal);
         MU_BACK_NEW      =  (MU_BACK_OLD/mu_back_proposal)*randg(mu_back_proposal)  ;
          
         
% Calculate the likelihood

         constant1=0;
         for k  = 1:length(sign_siz)
             Tnew = sqrt(4*(MU_BACK_NEW+MU_NEW.*(b(k,:)*Constant{k})).*Time{k}+1.5)-...
                 0.25./sqrt((MU_BACK_NEW+MU_NEW.*(b(k,:)*Constant{k})).*Time{k})  ;   
             
             Told = sqrt(4*(MU_BACK_OLD+MU_OLD.*(b(k,:)*Constant{k})).*Time{k}+1.5)-...
                 0.25./sqrt((MU_BACK_OLD+MU_OLD.*(b(k,:)*Constant{k})).*Time{k})  ; 
                      
             constant1 = constant1 + sum( (Told.^2 - Tnew.^2)./2 + trace{k}.*(Tnew-Told) );
         end

% Calculate the logarithm of posterior ratio                           
         logr = constant1 + ((2*mu_proposal-mu_alpha)*log(MU_OLD/MU_NEW))+...
                ((MU_OLD-MU_NEW)/mu_beta)+...
                (mu_proposal*((MU_NEW/MU_OLD)-(MU_OLD/MU_NEW)))+...
                ((2*mu_back_proposal-mu_back_alpha)*log(MU_BACK_OLD/MU_BACK_NEW))+...   
                ((MU_BACK_OLD-MU_BACK_NEW)/mu_back_beta)+...
                (mu_back_proposal*((MU_BACK_NEW/MU_BACK_OLD)-(MU_BACK_OLD/MU_BACK_NEW)));
         
% Accept or reject the proposals
         if  logr>(-exprnd(1))
             MU      = MU_NEW      ;
             MU_BACK = MU_BACK_NEW ;
             mu_acceptance_rate = mu_acceptance_rate+1;
         else
             MU      = MU_OLD      ;
             MU_BACK = MU_BACK_OLD ;
         end
         
end
```
