## Supplementary material for "An Alternative Framework for Fluorescence Correlation Spectroscopy": Source Code: Sample_Generator.m

```
function [ single_photon_arrivals ] = Sample_Generator( ...
...
...
Lxy                , Lz                   , Length_signal  , Number_molecules , ...
wxy                , wz                   , D              , tau              , ...
Molecule_emission  , Background_emission  , PSF_func       , handles)
  
%%%%%%%%%%%%%%%%%%%%%%%%%%%%%%%%%%%%%%%%%%%%%%%%%%%%%%%%%%%%%%%%%%%%%%%%%%%
%%%%%%%%%%%%%%%%%%%%%%%%%%%%%% INPUT %%%%%%%%%%%%%%%%%%%%%%%%%%%%%%%%%%%%%%
%%%%%%%%%%%%%%%%%%%%%%%%%%%%%%%%%%%%%%%%%%%%%%%%%%%%%%%%%%%%%%%%%%%%%%%%%%%
     % Lxy                   :  Periodic boundary in  x and y coordinates
     % Lz                    :  Periodic boundary in  z coordinate
     % Length_signal         :  Length of the trace
     % Number_molecules      :  Number of molecules for each species
     % wxy                   :  Minor Semi-axis of the confocal volume
     % wz                    :  Major Semi-axis of the confocal volume
     % D                     :  Diffusion coeffcient(s)
     % tau                   :  Data acquisition time
     % Molecule_emission     :  Molecule photon emission rates
     % Background_emission   :  Backgrouund photon emission rates
     % PSF_func              :  Point spread function type (3DG = 1 , 2DG-L =2 and Cylindrical = 3)

           
%%%%%%%%%%%%%%%%%%%%%%%%%%%%%%%%%%%%%%%%%%%%%%%%%%%%%%%%%%%%%%%%%%%%%%%%%%%
%%%%%%%%%%%%%%%%%%%%%%%%%%%%%% OUTPUT %%%%%%%%%%%%%%%%%%%%%%%%%%%%%%%%%%%%%
%%%%%%%%%%%%%%%%%%%%%%%%%%%%%%%%%%%%%%%%%%%%%%%%%%%%%%%%%%%%%%%%%%%%%%%%%%%
     % single_photon_arrivals  :  Single photon arrival time trace
     
% Set the PSF
if sum(PSF_func)==sum('Gaussian')
    PSF = 1;   % 3D Gaussian
else
    if  sum(PSF_func)==sum('Lorentzian')
        PSF = 2;   % 2DG-Lorentzian
    else 
        PSF = 3;   % Cylindrical
    end
end

% Create an empty vector for the intensities over time steps
Emit_rate_pos = zeros(1,Length_signal+1); 

% Measure the kinetic step size
k=sqrt(2*D*tau);

% consider the space size for locations
x = zeros ( 2 , Number_molecules );
y = zeros ( 2 , Number_molecules );
z = zeros ( 2 , Number_molecules );


% Initialize the time bar
cla(handles.axes23)
set(handles.axes23,'Units','pixels','XLim',[0 1],'YLim',[0 1],...
        'XTick',[],'YTick',[],'Color','w','XColor','w','YColor','w');
    
patch(handles.axes23,[0 0 0 0],[0 1 1 0],'b');
handles.text80.String='Initializing ...';
drawnow
time_loop =0 ;
save_bar=1;
tic
    
    

% Randomly sample the location of the molecules from the uniform probability
for jk1=1:Number_molecules
    
    x(1,jk1) = Lxy*(1-2*rand());
    y(1,jk1) = Lxy*(1-2*rand());
    z(1,jk1) = Lz *(1-2*rand());
    
    switch PSF
        case 1
             Emit_rate_pos(1)=Emit_rate_pos(1)+exp(-2*( (x(1,jk1)/wxy)^2+(y(1,jk1)/wxy)^2+(z(1,jk1)/wz)^2) );
        case 2
             Emit_rate_pos(1)=Emit_rate_pos(1)+exp(-(2*((x(1,jk1)^2)+(y(1,jk1)^2))/(wxy^2))/(1+(z(1,jk1)/wz)^2))/(1+(z(1,jk1)/wz)^2);
        otherwise
             Emit_rate_pos(1)=Emit_rate_pos(1)+exp(-2*( (x(1,jk1)/wxy)^2+(y(1,jk1)/wxy)^2) );
    end
    
end
  

for i=2:Length_signal+1  
    for j1  = 1:Number_molecules
        
        % Sample the locations baswed on brownian motion
        x(2,j1) = x(1,j1)+k*randn();
        y(2,j1) = y(1,j1)+k*randn();
        z(2,j1) = z(1,j1)+k*randn();
       
        % Periodoc boundaries
        if  x(2,j1) >= Lxy
            x(2,j1)  = x(2,j1)-2*Lxy;
        end  
        if  x(2,j1) <= -Lxy
            x(2,j1)  = x(2,j1)+2*Lxy;
        end 
        if  y(2,j1) >= Lxy
            y(2,j1)  = y(2,j1)-2*Lxy;
        end  
        if  y(2,j1) <= -Lxy
            y(2,j1)  = y(2,j1)+2*Lxy;
        end 
        if  z(2,j1) >= Lz
            z(2,j1)  = z(2,j1)-2*Lz;
        end  
        if  z(2,j1) <= -Lz
            z(2,j1)  = z(2,j1)+2*Lz;
        end 
        
        switch PSF
            case 1
                  Emit_rate_pos(i)=Emit_rate_pos(i)+exp(-2*( (x(2,j1)/wxy)^2+(y(2,j1)/wxy)^2 +(z(2,j1)/wz)^2 )); 
            case 2
                  Emit_rate_pos(i)=Emit_rate_pos(i)+exp(-(2*((x(2,j1)^2)+(y(2,j1)^2))/(wxy^2))/(1+(z(2,j1)/wz)^2))/(1+(z(2,j1)/wz)^2); 
            otherwise
                  Emit_rate_pos(i)=Emit_rate_pos(i)+exp(-2*( (x(2,j1)/wxy)^2+(y(2,j1)/wxy)^2 )); 
        end
    end
    
    x(1,:)=x(2,:);
    y(1,:)=y(2,:);
    z(1,:)=z(2,:);
    
    
    if find((floor((1:1:10).*(Length_signal+1)./10))==i)>=1
      
       time_loop=time_loop+toc;
       remaining_time=floor((time_loop/(save_bar.*(Length_signal+1)./10))*((10-save_bar).*...
           (Length_signal+1)./10)*10)/10;
       handles.text80.String=['Remaining time = ', num2str(remaining_time),' s'];
       patch(handles.axes23,[0  0  save_bar/10  save_bar/10],[0 1 1 0],'b');
       save_bar=save_bar+1;
       drawnow
       tic
    end
    
                     
end % end of time loop
handles.text80.String='Done!';

% sample the photons from a bernoulli distribution
   photons= binornd(ones(1,length(Emit_rate_pos)) , 1-exp(-tau*(Background_emission+Molecule_emission.*Emit_rate_pos)) );

% Find the arrival times of observed photons
   single_photon_arrivals=find(photons>0)*tau;
end
```
