## Supplementary material for "An Alternative Framework for Fluorescence Correlation Spectroscopy": Source Code: Sample_Loads.m

```
function[ b , acceptance_rate_b ]    =  Sample_Loads( ...
...
...
b_old               ,   I         ,   trace             , ...
acceptance_rate_b   ,   Time      ,   Number_moleclues  , I_back    , ...
q                   ,   Constant    ) 

%%%%%%%%%%%%%%%%%%%%%%%%%%%%%%%%%%%%%%%%%%%%%%%%%%%%%%%%%%%%%%%%%%%%%%%%%%%
%%%%%%%%%%%%%%%%%%%%%%%%%%%%%% INPUT %%%%%%%%%%%%%%%%%%%%%%%%%%%%%%%%%%%%%%
%%%%%%%%%%%%%%%%%%%%%%%%%%%%%%%%%%%%%%%%%%%%%%%%%%%%%%%%%%%%%%%%%%%%%%%%%%%
     % b_old                         :  Old sampled load 
     % I                             :  Moelcule photon emission rate
     % trace                         :  Original trace
     % x                             :  Positions of the molecules (Normalized)
     % acceptance_rate_b             :  Previous acceptance rate 
     % Time                          :  Data acquisition time
     % Number_molecules              :  Number of defined molecules
     % I_back                        :  Background photon emission rate
     % q                             :  Weight of loads
     % PSF_Func                      :  (PSF_Func=1 -> Gaussian , PSF_Func=2 -> Lorentzian-Gaussian)
     
%%%%%%%%%%%%%%%%%%%%%%%%%%%%%%%%%%%%%%%%%%%%%%%%%%%%%%%%%%%%%%%%%%%%%%%%%%%
%%%%%%%%%%%%%%%%%%%%%%%%%%%%%% OUTPUT %%%%%%%%%%%%%%%%%%%%%%%%%%%%%%%%%%%%%
%%%%%%%%%%%%%%%%%%%%%%%%%%%%%%%%%%%%%%%%%%%%%%%%%%%%%%%%%%%%%%%%%%%%%%%%%%%
     % b                             :  Updated load b
     % acceptance_rate_b             :  Updated acceptance rate
   
     
   % Propose the new loads
    b_new = rand(1,Number_moleclues)<q;
 

    
   Tnew = sqrt(4*(I_back + I.*(b_new*Constant)).*Time + 1.5)-0.25./sqrt((I_back + I.*(b_new*Constant)).*Time )  ;   
   Told = sqrt(4*(I_back + I.*(b_old*Constant)).*Time + 1.5)-0.25./sqrt((I_back + I.*(b_old*Constant)).*Time )   ;
         
   % Logarithm of the Posterior ratio
   
   logr = sum( (Told.^2 - Tnew.^2)./2 + trace.*(Tnew-Told) );

   
   % Check the acceptance
   if  logr>(-exprnd(1))
       b = b_new ;   %% Accept %%
       acceptance_rate_b = acceptance_rate_b + 1 ;
   else
       b = b_old ;   %% Reject %%
   end
   
end
```
