## Supplementary material for "An Alternative Framework for Fluorescence Correlation Spectroscopy": Source Code: UKF.m

```
function [XX,VT] = UKF( x  , c            , constant , I   , Y  , ...
                        Z  , observation  , vdif     , yy  , p  , ...
                        f  , PSF          )


%%%%%%%%%%%%%%%%%%%%%%%%%%%%%%%%%%%%%%%%%%%%%%%%%%%%%%%%%%%%%%%%%%%%%%%%%%%
%%%%%%%%%%%%%%%%%%%%%%%%%%%%%% INPUT %%%%%%%%%%%%%%%%%%%%%%%%%%%%%%%%%%%%%%
%%%%%%%%%%%%%%%%%%%%%%%%%%%%%%%%%%%%%%%%%%%%%%%%%%%%%%%%%%%%%%%%%%%%%%%%%%%
     % x                        :  Mean value of the previous filter
     % c                        :  Variance of the previous filter
     % constant                 :  Effect of the other molecules and background on the observation
     % I                        :  Molecule emission rate
     % Y                        :  Y dimension of the molecule (It could be X or Y dimension depends on the FFBS function)
     % Z                        :  Z dimension of the molecule (It could be X or Y dimension depends on the FFBS function)
     % observation              :  Observation (Transformed signal by Anscombe)
     % vdif                     :  variance of the kinetics (vdif=2* D* Time/wxy^2)
     % yy                       :  Location of the sigma points used in UKF
     % p                        :  Weight of the sigma point used in UKF
     
%%%%%%%%%%%%%%%%%%%%%%%%%%%%%%%%%%%%%%%%%%%%%%%%%%%%%%%%%%%%%%%%%%%%%%%%%%%
%%%%%%%%%%%%%%%%%%%%%%%%%%%%%% OUTPUT %%%%%%%%%%%%%%%%%%%%%%%%%%%%%%%%%%%%%
%%%%%%%%%%%%%%%%%%%%%%%%%%%%%%%%%%%%%%%%%%%%%%%%%%%%%%%%%%%%%%%%%%%%%%%%%%%
     % XX                       :  Mean value of the filter in the positive side
     % VT                       :  Variance of the filter in the positive side

     
% Since we have a symmetric filter,first we calculate the mean and variance
% of the one mode and due to the symmetry the other mode has the same variace
% and symmetric mean value.


% Find the sigma points based on the mean valuue and variance of the
% previous filter and kinetic
         xx= x + yy*sqrt(c+vdif);

         MM= f(xx,Y,Z);
         MM(xx<0) = f(0,Y,Z);
     
         mm    = constant +I.*MM ;
         
         % Calculate the obvervation at sigma points
         vv    = sqrt( 4*mm + 1.5) - 0.25./sqrt(mm);
         
         % Prediction Step
         X     = xx*p;
         V     = vv*p;

         % Calculate the covariance matrix elements
         pvv   = ( (vv-V).^2  )*p +1;

         % Calculate the gain
         Kgain = (((xx-X).*(vv-V))*p)/pvv;          % To speed up the code, we avoid pxv = ((xx-X).*(vv-V))*p.
         
         % Correction Step
         XX    = abs(X+Kgain*(observation-V));
         VT    = ((xx-X).^2)*p - Kgain*pvv*Kgain;   % To speed up code, we avoid calculating pxx = ((xx-X).^2)*p.


end
```
