## Supplementary material for "An Alternative Framework for Fluorescence Correlation Spectroscopy": Source Code: Anscombe.m

```
function [observation] = Anscombe(trace)
% %% INPUT
     % trace               : Original poissonian nise trace
     
% %% OUTPUT
     % observation         : Transformed to normal distribution 

     observation = sqrt( 4*trace+1.5 ) ;
%       observation = trace ;
end
```
