## Supplementary material for "An Alternative Framework for Fluorescence Correlation Spectroscopy": Source Code: Diffusion.m

```
function [ D ]= Diffusion( x              , Number_molecules , wxy      , wz    , ...
                           D_alpha_prior  , D_beta_prior     , sign_siz , Time  , ...
                           Max_Species      )

%%%%%%%%%%%%%%%%%%%%%%%%%%%%%%%%%%%%%%%%%%%%%%%%%%%%%%%%%%%%%%%%%%%%%%%%%%%
%%%%%%%%%%%%%%%%%%%%%%%%%%%%%% INPUT %%%%%%%%%%%%%%%%%%%%%%%%%%%%%%%%%%%%%%
%%%%%%%%%%%%%%%%%%%%%%%%%%%%%%%%%%%%%%%%%%%%%%%%%%%%%%%%%%%%%%%%%%%%%%%%%%%
     % x                        :  Locations of all molecules
     % Number_molecules         :  Maximum number of molecule for each species
     % wxy                      :  Minor Semi-axis of the confocal volume
     % wz                       :  Major Semi-axis of the confocal volume
     % D_alpha_prior            :  Alpha for inverse gamma prob. dis. for prior related to Diffusion
     % D_beta_prior             :  Beta for  inverse gamma prob. dis. for prior related to Diffusion
     % sign_size                :  Length of the observation trace
     % Time                     :  Vector of bin sizes
     % Max_Species              :  Total number of species
     
%%%%%%%%%%%%%%%%%%%%%%%%%%%%%%%%%%%%%%%%%%%%%%%%%%%%%%%%%%%%%%%%%%%%%%%%%%%
%%%%%%%%%%%%%%%%%%%%%%%%%%%%%% OUTPUT %%%%%%%%%%%%%%%%%%%%%%%%%%%%%%%%%%%%%
%%%%%%%%%%%%%%%%%%%%%%%%%%%%%%%%%%%%%%%%%%%%%%%%%%%%%%%%%%%%%%%%%%%%%%%%%%%
     % D                        :  Sampled diffusion coefficients of species


% creat an empty column
D=nan(Max_Species,1);

% For loop for number of species
       for m=1:Max_Species
% Sample diffusion coefficient of species m
          D(m,1)  =  1/gamrnd(...
                       D_alpha_prior+((sign_siz-2)*1.5*Number_molecules) , ...
                      1/(D_beta_prior+0.25*(repmat([wxy wxy wz].^2,1,Number_molecules)*...
                      ((x(3*Number_molecules*(m-1)+1:3*Number_molecules*m,2:end-1)...
                       -x(3*Number_molecules*(m-1)+1:3*Number_molecules*m,3:end )).^2))*(Time(3:end).^-1)')) ;
       end
       
end
```
