## Supplementary material for "An Alternative Framework for Fluorescence Correlation Spectroscopy": Source Code: FFBS.m

```
%%%%%%%%%%%%%%%%%%%%%%%%%%%%%%%%%%%%%%%%%%%%%%%%%%%%%%%%%%%%%%%%%%%%%%%%%%%
%%%%%%%%%%%%%%%%%%%%%%%%%% Extended Kalman Filter %%%%%%%%%%%%%%%%%%%%%%%%%
%%%%%%%%%%%%%%%%%%%%%%%%%%%%%%%%%%%%%%%%%%%%%%%%%%%%%%%%%%%%%%%%%%%%%%%%%%%

function[initial_positions ]  =  FFBS( ...
    ...
    ...
observation        , I           , I_b         , Max              , ...
initial_positions  , num_par     , muu         , varr             , ...
yy                 , pp          , m           , PSF_func         , ...
vdif               , b           , Max_Species , Number_molecules , ...
f_3DG              , f_2DGL_D1_2 , f_2DGL_D3   , f_2DGC           ) 

%%%%%%%%%%%%%%%%%%%%%%%%%%%%%%%%%%%%%%%%%%%%%%%%%%%%%%%%%%%%%%%%%%%%%%%%%%%
%%%%%%%%%%%%%%%%%%%%%%%%%%%%%% INPUT %%%%%%%%%%%%%%%%%%%%%%%%%%%%%%%%%%%%%%
%%%%%%%%%%%%%%%%%%%%%%%%%%%%%%%%%%%%%%%%%%%%%%%%%%%%%%%%%%%%%%%%%%%%%%%%%%%
     % observation              :  Anscombe transformed trace
     % I                        :  Vector of molecule photon emission rates at any time step
     % I_b                      :  Vector of background photon emission rates at any time step
     % Max                      :  length of the trace
     % initial_positions        :  Locations of all molecules
     % num_par                  :  The tag for targeted molecule
     % muu                      :  Mean values for X0 prior in normalized scale for x, y and z coordintes
     % varr                     :  Variance for X0 prior in normalized scale for x, y and z coordintes 
     % yy                       :  Location of the sigma points used in UKF
     % pp                       :  Weight of the sigma points used in UKF
     % m                        :  The tag for targeted species
     % llk                      :  The sequence of coordinates
     % PSF_func                 :  Point Spread Function type ( 'Gaussian' = 1 or 'Lorentzian Gaussian' = 2)
     % vdif                     :  Vectore of the kinetic variances at each time step and coordinates
     % b                        :  Loads of all molecules and species
     % Max_Species              :  Total number of species
     % Number_molecules         :  Maximum number of molecules for each species
    

%%%%%%%%%%%%%%%%%%%%%%%%%%%%%%%%%%%%%%%%%%%%%%%%%%%%%%%%%%%%%%%%%%%%%%%%%%%
%%%%%%%%%%%%%%%%%%%%%%%%%%%%%% OUTPUT %%%%%%%%%%%%%%%%%%%%%%%%%%%%%%%%%%%%%
%%%%%%%%%%%%%%%%%%%%%%%%%%%%%%%%%%%%%%%%%%%%%%%%%%%%%%%%%%%%%%%%%%%%%%%%%%%
     % initial_positions        :  Updated locations of targeted molecule


%%%%%%%%%%%%%%%%%%%%%%%%%%%%%%%%%%%%%%%%%%%%%%%%%%%%%%%%%%%%%%%%%%%%%%%%%%%
%%%%%%%%%%%%%%%%%%%%%%%%%%  Precalculations %%%%%%%%%%%%%%%%%%%%%%%%%%%%%%%
%%%%%%%%%%%%%%%%%%%%%%%%%%%%%%%%%%%%%%%%%%%%%%%%%%%%%%%%%%%%%%%%%%%%%%%%%%%
vt        =  nan(1,Max+1)               ;  % Forward Filter Varinace
x         =  nan(1,Max+1)               ;  % Mean value of the Forward Filter


% calculate the effect of the other particels on the likelihood
b(Number_molecules*(m-1)+num_par)=0;

% Precalculation
nump=3*Number_molecules*(m-1)+num_par*3;

%%%%%%%%%%%%%%%%%%%%%%%%%%%%%%%%%%%%%%%%%%%%%%%%%%%%%%%%%%%%%%%%%%%%%%%%%%%
%%%%%%%%%%%%%%%%%%%%%  nn=1,2,3 as X, Y , Z  %%%%%%%%%%%%%%%%%%%%%%%%%%%%%%    
%%%%%%%%%%%%%%%%%%%%%%%%%%%%%%%%%%%%%%%%%%%%%%%%%%%%%%%%%%%%%%%%%%%%%%%%%%%
switch PSF_func
       case 1    % 3D Gaussian PSF
           % calculate the effect of the other particels on the likelihood
           constant1=b'*(repelem(I,Number_molecules,1).*...
                  exp(-2*(initial_positions(1:3:3*Number_molecules*Max_Species,:).^2+...
                          initial_positions(2:3:3*Number_molecules*Max_Species,:).^2+...
                          initial_positions(3:3:3*Number_molecules*Max_Species,:).^2)));
           % Contribution of backgrond and all of active molecules except the current one in the trace
           constant1  = I_b+ constant1;
           
           
            AA = [ 1 , 2 , 3  ; ...
                   2 , 1 , 3  ; ...
                   3 , 1 , 2 ];

            for nn = 1:3
                x(1)    =  muu(AA(nn,1))       ;  % Set the initial Mean value of the Forward Filter
                vt(1)   =  varr(AA(nn,1))      ;  % Set the initial Variance of the Forward Filter
                
                for t = 1:Max
                    [x(t+1),vt(t+1)] = UKF( x(t)                                    , ...
                                            vt(t)                                   , ...
                                            constant1(t)                            , ...
                                            I(m,t)                                    , ...
                                            initial_positions(nump-(3-AA(nn,2)),t)  , ...
                                            initial_positions(nump-(3-AA(nn,3)),t)  , ...
                                            observation(t)                          , ...
                                            vdif(AA(nn,1),t)                        , ...
                                            yy                                      , ...
                                            pp                                      , ...
                                            f_3DG                                  );
                end
           
            
            
%%%%%%%%%%%%%%%%%%%%%%%%% Backward sampling %%%%%%%%%%%%%%%%%%%%%%%%%%%%%%%
              % Sample the last point directly form the likelihood(filter)
              if  rand()>0.5
                  initial_positions(nump-(3-AA(nn,1)),Max)     =  x(Max+1) + sqrt(vt(Max+1))*randn()         ;
              else
                  initial_positions(nump-(3-AA(nn,1)),Max)     = -x(Max+1) + sqrt(vt(Max+1))*randn()         ;
              end

% Sample the rest of points form the likelihood times the kinetics
% Sample the rest of the points from the likelihood times the kinetics
              vtt  = vt(1:Max)+vdif(AA(nn,1),:)  ;
              vvt  = sqrt(vt(1:Max).*vdif(AA(nn,1),:)./vtt)    ;
              for i = Max:-1:2            
                  if rand() < 1/(1+exp(-2*initial_positions(nump-(3-AA(nn,1)),i)*x(i)/vtt(i)))
                  initial_positions(nump-(3-AA(nn,1)),i-1) = ...
                    ( initial_positions(nump-(3-AA(nn,1)),i)*vt(i)+vdif(AA(nn,1),i)*x(i) )/vtt(i) + vvt(i)*randn()   ;
                  else         
                  initial_positions(nump-(3-AA(nn,1)),i-1) = ...
                    ( initial_positions(nump-(3-AA(nn,1)),i)*vt(i)-vdif(AA(nn,1),i)*x(i) )/vtt(i) + vvt(i)*randn()   ;
                  end
              end  
            end

       case 2   % 2D Gaussian Lorentzian PSF
           % calculate the effect of the other particels on the likelihood
           constant1=b'*(repelem(I,Number_molecules,1).*...
                  (exp(-2*(initial_positions(1:3:3*Number_molecules*Max_Species,:).^2+...
                           initial_positions(2:3:3*Number_molecules*Max_Species,:).^2)./...
                        (1+initial_positions(3:3:3*Number_molecules*Max_Species,:).^2))./...
                        (1+initial_positions(3:3:3*Number_molecules*Max_Species,:).^2)));       
           % Contribution of backgrond and all of active molecules except the current one in the trace
           constant1  = I_b+ constant1;         
                    
                    
            AA=[1 , 2 , 3 ; ...
                2 , 1 , 3 ; ...
                3 , 1 , 2]; 
            
           for nn=1:3 
               x(1)    =  muu(AA(nn,1))       ;  % Set the initial Mean value of the Forward Filter
               vt(1)   =  varr(AA(nn,1))      ;  % Set the initial Variance of the Forward Filter
               switch dimension
                     case AA(nn,1) 
                          for t=1:Max
                              [x(t+1),vt(t+1)] = UKF( x(t)                                    , ...
                                                      vt(t)                                   , ...
                                                      constant1(t)                            , ...
                                                      I(m,t)                                    , ...
                                                      initial_positions(nump-(3-AA(nn,2)),t)  , ...
                                                      initial_positions(nump-(3-AA(nn,3)),t)  , ...
                                                      observation(t)                          , ...
                                                      vdif(AA(nn,1),t)                        , ...
                                                      yy                                      , ...
                                                      pp                                      , ...
                                                      f_2DGL_D1_2                            );
                           end   
                         
                  case AA(nn,1)
                       for t=1:Max
                           [x(t+1),vt(t+1)] = UKF( x(t)                                    , ...
                                                   vt(t)                                   , ...
                                                   constant1(t)                            , ...
                                                   I(m,t)                                    , ...
                                                   initial_positions(nump-(3-AA(nn,2)),t)  , ...
                                                   initial_positions(nump-(3-AA(nn,3)),t)  , ...
                                                   observation(t)                          , ...
                                                   vdif(AA(nn,1),t)                        , ...
                                                   yy                                      , ...
                                                   pp                                      , ...
                                                   f_2DGL_D1_2                            );
                       end   
                      
                  otherwise 
                       for t=1:Max
                           [x(t+1),vt(t+1)] = UKF( x(t)                                    , ...
                                                   vt(t)                                   , ...
                                                   constant1(t)                            , ...
                                                   I(m,t)                                    , ...
                                                   initial_positions(nump-(3-AA(nn,2)),t)  , ...
                                                   initial_positions(nump-(3-AA(nn,3)),t)  , ...
                                                   observation(t)                          , ...
                                                   vdif(AA(nn,1),t)                        , ...
                                                   yy                                      , ...
                                                   pp                                      , ...
                                                   f_2DGL_D3                              );
                       end   
              end
           
           
%%%%%%%%%%%%%%%%%%%%%%%%% Backward sampling %%%%%%%%%%%%%%%%%%%%%%%%%%%%%%%
              % Sample the last point directly form the likelihood(filter)
              if  rand()>0.5
                  initial_positions(nump-(3-AA(nn,1)),Max)     =  x(Max+1) + sqrt(vt(Max+1))*randn()         ;
              else
                  initial_positions(nump-(3-AA(nn,1)),Max)     = -x(Max+1) + sqrt(vt(Max+1))*randn()         ;
              end

% Sample the rest of points form the likelihood times the kinetics
% Sample the rest of the points from the likelihood times the kinetics
              vtt  = vt(1:Max)+vdif(AA(nn,1),:)  ;
              vvt  = sqrt(vt(1:Max).*vdif(AA(nn,1),:)./vtt)    ;
              for i = Max:-1:2            
                  if rand() < 1/(1+exp(-2*initial_positions(nump-(3-AA(nn,1)),i)*x(i)/vtt(i)))
                  initial_positions(nump-(3-AA(nn,1)),i-1) = ...
                    ( initial_positions(nump-(3-AA(nn,1)),i)*vt(i)+vdif(AA(nn,1),i)*x(i) )/vtt(i) + vvt(i)*randn()   ;
                  else         
                  initial_positions(nump-(3-AA(nn,1)),i-1) = ...
                    ( initial_positions(nump-(3-AA(nn,1)),i)*vt(i)-vdif(AA(nn,1),i)*x(i) )/vtt(i) + vvt(i)*randn()   ;
                  end
    
              end 
           end

    otherwise    % 2D Gaussian Cylindrical PSF
            % calculate the effect of the other particels on the likelihood
            constant1=b'*(repelem(I,Number_molecules,1).*exp(-2*(initial_positions(1:3:3*Number_molecules*Max_Species,:).^2+...
                          initial_positions(2:3:3*Number_molecules*Max_Species,:).^2)));  
           % Contribution of backgrond and all of active molecules except the current one in the trace
           constant1  = I_b+ constant1;         
                      
            AA=[1 , 2  ; ...
                2 , 1 ]; 
            
           for nn=1:2 
               x(1)    =  muu(AA(nn,1))       ;  % Set the initial Mean value of the Forward Filter
               vt(1)   =  varr(AA(nn,1))      ;  % Set the initial Variance of the Forward Filter
               for t=1:Max
                   [x(t+1),vt(t+1)] = UKF( x(t)                                      , ...
                                           vt(t)                                     , ...
                                           constant1(t)                              , ...
                                           I(m,t)                                      , ...
                                           initial_positions(nump-(3-AA(nn,2)),t)    , ...
                                           initial_positions(nump-(3-AA(nn,2)),t)    , ...
                                           observation(t)                            , ...
                                           vdif(AA(nn,1),t)                          , ...
                                           AA(nn,1)                                  , ...
                                           yy                                        , ...
                                           pp                                        , ...
                                           f_2DGC                                    );
               end
         
           
%%%%%%%%%%%%%%%%%%%%%%%%% Backward sampling %%%%%%%%%%%%%%%%%%%%%%%%%%%%%%
               % Sample the last point directly form the likelihood(filter)
               if  rand()>0.5
                   initial_positions(nump-(3-AA(nn,1)),Max)     =  x(Max+1) + sqrt(vt(Max+1))*randn()         ;
               else
                   initial_positions(nump-(3-AA(nn,1)),Max)     = -x(Max+1) + sqrt(vt(Max+1))*randn()         ;
               end


               % Sample the rest of points form the likelihood times the kinetics
               % Sample the rest of the points from the likelihood times the kinetics
               vtt  = vt(1:Max)+vdif(AA(nn,1),:)  ;
               vvt  = sqrt(vt(1:Max).*vdif(AA(nn,1),:)./vtt)    ;
               for i = Max:-1:2            
                   if rand() < 1/(1+exp(-2*initial_positions(nump-(3-AA(nn,1)),i)*x(i)/vtt(i)))
                                 initial_positions(nump-(3-AA(nn,1)),i-1) = ...
                                   ( initial_positions(nump-(3-AA(nn,1)),i)*vt(i)+vdif(AA(nn,1),i)*x(i) )/vtt(i) + vvt(i)*randn()   ;
                   else         
                                 initial_positions(nump-(3-AA(nn,1)),i-1) = ...
                                   ( initial_positions(nump-(3-AA(nn,1)),i)*vt(i)-vdif(AA(nn,1),i)*x(i) )/vtt(i) + vvt(i)*randn()   ;
                   end
               end
           end
end

end
```
