## Supplementary material for "An Alternative Framework for Fluorescence Correlation Spectroscopy": Source Code: Gibbs_sampler.m

```
function [ D                   , mu                  , mu_back  ,  b    , ...
           acceptance_rate_mu  , acceptance_rate_b   , x        ,  q    , ...
           concentration] = ...
...
...
Gibbs_sampler(...
...
...
trace            , Time          , D_alpha_prior    , D_beta_prior     , ...
D                , mu            , mu_back          , Number_molecules , ...
Max_iter         , wxy           , wz               , mu_proposal      , ...
mu_back_proposal , mu_alpha      , mu_beta          , mu_back_alpha    , ...
mu_back_beta     , alpha_q       , beta_q           , mu_prior_xy      , ...
mu_prior_z       , var_prior_xy  , var_prior_z      , b                , ...
x                , q             , concen_radious   , concentration    , ...
Max_Species      , PSF_func      , save_size        )


%%%%%%%%%%%%%%%%%%%%%%%%%%%%%%%%%%%%%%%%%%%%%%%%%%%%%%%%%%%%%%%%%%%%%%%%%%%
%%%%%%%%%%%%%%%%%%%%%%%%%%%%%% INPUT %%%%%%%%%%%%%%%%%%%%%%%%%%%%%%%%%%%%%%
%%%%%%%%%%%%%%%%%%%%%%%%%%%%%%%%%%%%%%%%%%%%%%%%%%%%%%%%%%%%%%%%%%%%%%%%%%%
     % trace                         :  Original trace (Number of observed photons per bin)
     % Time                          :  The time arrivel time differences
     % D_alpha_prior                 :  Alpha for inverse gamma prob. dis. for prior related to Diffusion
     % D_beta_prior                  :  Beta for  inverse gamma prob. dis. for prior related to Diffusion
     % D                             :  Values for Diffusion coefficient
     % mu                            :  Values for molecule photon emission rate
     % mu_back                       :  Values for background photon emission rate
     % Number_molecules              :  Number of defined molecules for each species
     % Max_iter                      :  Number of Gibbs sampler iterations
     % wxy                           :  Minor Semi-axis of the confocal volume
     % wz                            :  Major Semi-axis of the confocal volume
     % mu_proposal                   :  Proposal for sampling emission rate of molecules in Metropolis_mu
     % mu_back_proposal              :  Proposal for sampling background emission rate in Metropolis_mu
     % mu_alpha                      :  Alpha value for Emission rate of molecules, prior
     % mu_beta                       :  Beta value for Emission rate of molecules, prior
     % mu_back_alpha                 :  Alpha value for Background emission rate's prior 
     % mu_back_beta                  :  Beta value for Background emission rate's prior 
     % alpha_q                       :  Alpha parameter of load b prior         ( Gamma distribution)
     % beta_q                        :  Beta parameter of load b prior          ( Gamma distribution)
     % mu_prior_xy                   :  Mean value for X0 prior in x and y coordintes
     % mu_prior_z                    :  Mean value for X0 prior in z coordinte
     % var_prior_xy                  :  Variance for X0 prior in x and y coordintes
     % var_prior_z                   :  Variance for X0 prior in z coordintes
     % b                             :  Sampled loads
     % x                             :  Updated locations of molecules
     % q                             :  Sampled weights of loads
     % concen_radious                :  Semi minor axis of ellipsoid to consider as the effective region
     % concen_test                   :  Number of species
     % concentration                 :  Updated concentrations of molecules in the ROI
     % Max_Species                   :
     % PSF_func                      :  Point Spread Function type ( 'Gaussian' or 'Lorentzian Gaussian')
     % iter_b_mu                     :  Number of iteration for emission rates and loads Metropolis 
     % save_size                     :  Lenght of save intervals


%%%%%%%%%%%%%%%%%%%%%%%%%%%%%%%%%%%%%%%%%%%%%%%%%%%%%%%%%%%%%%%%%%%%%%%%%%%
%%%%%%%%%%%%%%%%%%%%%%%%%%%%%% OUTPUT %%%%%%%%%%%%%%%%%%%%%%%%%%%%%%%%%%%%%
%%%%%%%%%%%%%%%%%%%%%%%%%%%%%%%%%%%%%%%%%%%%%%%%%%%%%%%%%%%%%%%%%%%%%%%%%%%

     % D                             :  An array of sampeled Diffusion coefficient
     % mu                            :  An array of sampeled molecule emission rate
     % mu_back                       :  An array of sampeled background emission rate
     % b                             :  A matrix of sampeled loads
     % q                             :  A matrix of sampeled weights of loads
     % concentration                 :  Concentration of molecules in ROI
     % acceptance_rate_mu            :  Acceptance rate of the emission rates Metropolis
     % acceptance_rate_b             :  Acceptance rate of loads Metropolis
     % x                             :  Export the last sampled positions to use as the initial value of next run


% Shuffle the random generator
    rng('shuffle')
    
% Calculate the sigma points and their weigths to be used in UKF
     [yy, pp] = nwspgr('GQN', 1 , 5);
     yy=yy';

          
% Calculate the length of the trace
    sign_siz   = length(trace)                                            ; 

% Set the acceptance rates to zero
    acceptance_rate_mu  = 0                                               ;
    acceptance_rate_b   = 0                                               ;


% Normalize the mean value and variance of the prior
    X0   = [mu_prior_xy mu_prior_xy mu_prior_z]                           ;
    V0   = [var_prior_xy var_prior_xy var_prior_z]                        ;
    
% Transform the original trace to guassian by using Anscombe Transformation
    observation         =  Anscombe(trace)                                ;
    
% Calculate the size of the previous sample size 
    size_iter_old = length(mu_back)                                            ;
    
% Initialize the time bar
    hbar = waitbar(0,'Initializing ...')                                  ;
    time_loop =0 ;
    save_bar=1;
    tic
    set(hbar,'CloseRequestFcn','');
    % closereq
    
% Definition of the PSFs
f_3DG       =@(x,y,z) exp( -2*(x.^2 + y.^2 + z.^2));                     % 3DG

f_2DGL_D1_2 =@(x,y,z) exp( -2*(x.^2 + y.^2)./(1 + z.^2))./(1 + z.^2);    % 2DG-Lorentzian, x and y demensions
f_2DGL_D3   =@(x,y,z) exp( -2*(y.^2 + z.^2)./(1 + x.^2))./(1 + x.^2);    % 2DG-Lorentzian, z demension

f_2DGC      =@(x,y,z) exp( -2*(x.^2 + y.^2));                            % 2DG-cylindrical


% Start the Gibbs sampling iteration loop
    for i = 1+size_iter_old : Max_iter+size_iter_old
      
%%%%%%%%%%%%%%%%%%%%%%%%%%%%%%%%%%%%%%%%%%%%%%%%%%%%%%%%%%%%%%%%%%%%%%%%%%%
%%%%%%%%%%%%%%%%%%%%%%%%%  Pre-calculations %%%%%%%%%%%%%%%%%%%%%%%%%%%%%%%
%%%%%%%%%%%%%%%%%%%%%%%%%%%%%%%%%%%%%%%%%%%%%%%%%%%%%%%%%%%%%%%%%%%%%%%%%%%

    I_b    =  mu_back(i-1) .* Time       ;
    I      =  mu(:,i-1)    .* Time       ;

    for num_par=1:Number_molecules
% Start the FOR loop on the number of species
       for m=1:Max_Species
% Pre-calculation of the variance of the kinetics at each time step and coordiantes
            vdif   =  2*D(m,i-1)*[0 Time(2:end)]./[wxy^2;wxy^2;wz^2]      ;
%%%%%%%%%%%%%%%%%%%%%%%%%%%%%%%%%%%%%%%%%%%%%%%%%%%%%%%%%%%%%%%%%%%%%%%%%%%
%%%%% Sampling ACTIVE trajectorie for molecule num_par and species m %%%%%%
%%%%%%%%%%%%%%%%%%%%%%%%%%%%%%%%%%%%%%%%%%%%%%%%%%%%%%%%%%%%%%%%%%%%%%%%%%%
               if  b(Number_molecules*(m-1)+num_par,i-1)==1
                   x    = FFBS( observation  , I           , I_b          , sign_siz           , ...
                                x            , num_par     , X0           , V0                 , ...
                                yy           , pp          , m            , PSF_func           , ...
                                vdif         , b(:,i-1)    , Max_Species  ,  Number_molecules , ...
                                f_3DG        , f_2DGL_D1_2 , f_2DGL_D3    , f_2DGC           );      
               end
       end
    end
       
% Start the FOR loop on the number of species
    
%%%%%%%%%%%%%%%%%%%%%%%%%%%%%%%%%%%%%%%%%%%%%%%%%%%%%%%%%%%%%%%%%%%%%%%%%%%
%%%%%%%%%%% Compute concetration of the molecules in the ROI %%%%%%%%%%%%%%
%%%%%%%%%%%%%%%%%%%%%%%%%%%%%%%%%%%%%%%%%%%%%%%%%%%%%%%%%%%%%%%%%%%%%%%%%%%
     switch PSF_func
            case 1     % 3D Gaussian PSF
                 for l=1:length(concen_radious)
                      concentration(l,:,i-(floor((i-1)/1000)*1000)) = ...
                            b(:,i-1)'*(sqrt( x(1:3:3*Number_molecules*Max_Species,:).^2+...
                                             x(2:3:3*Number_molecules*Max_Species,:).^2+...
                                             x(3:3:3*Number_molecules*Max_Species,:).^2) <=concen_radious(l) );
                 end
             case 2      % 2D Gaussian-Lorentzian PSF
                  for l=1:length(concen_radious)
                     concentration(l,:,i-(floor((i-1)/1000)*1000)) = b(:,i-1)'*...
                     (sqrt( 0.5.*log(1+x(3:3:3*Number_molecules*Max_Species,:).^2)+...
                                ((x(1:3:3*Number_molecules*Max_Species,:).^2)+...
                                (x(2:3:3*Number_molecules*Max_Species,:).^2))...
                             ./(1+x(3:3:3*Number_molecules*Max_Species,:).^2) )<=concen_radious(l));
                  end
         otherwise       % 2D Gaussian-Cylindrical PSF
                  for l=1:length(concen_radious)
                      concentration(l,:,i-(floor((i-1)/1000)*1000)) = ...
                            b(:,i-1)'*(sqrt( x(1:3:3*Number_molecules*Max_Species,:).^2+...
                                             x(2:3:3*Number_molecules*Max_Species,:).^2) <=concen_radious(l) );
                  end
     end
     
       
%%%%%%%%%%%%%%%%%%%%%%%%%%%%%%%%%%%%%%%%%%%%%%%%%%%%%%%%%%%%%%%%%%%%%%%%%%%
%%%%%%%%%%%%%%%%%% Sampling INACTIVE trajectories  %%%%%%%%%%%%%%%%%%%%%%%%
%%%%%%%%%%%%%%%%%%%%%%%%%%%%%%%%%%%%%%%%%%%%%%%%%%%%%%%%%%%%%%%%%%%%%%%%%%%                          
       x  = Inactive_molecules( x            , sign_siz      , Number_molecules , Time        , ...
                                b(:,i-1)   , mu_prior_xy   , mu_prior_z       , var_prior_xy   , ...
                                var_prior_z  , wxy           , wz               , D(:,i-1)       , ...
                                Max_Species  );


%%%%%%%%%%%%%%%%%%%%%%%%%%%%%%%%%%%%%%%%%%%%%%%%%%%%%%%%%%%%%%%%%%%%%%%%%%%
%%%%%%%%%%%%%%%%%%%% Sampling the Diffusion coefficient %%%%%%%%%%%%%%%%%%%
%%%%%%%%%%%%%%%%%%%%%%%%%%%%%%%%%%%%%%%%%%%%%%%%%%%%%%%%%%%%%%%%%%%%%%%%%%%
       D(:,i)= Diffusion( x             , Number_molecules  , wxy      , wz      , ...
                          D_alpha_prior , D_beta_prior      , sign_siz , Time , ...
                          Max_Species     ) ;    
   


%%%%%%%%%%%%%%%%%%%%%%%%%%%%%%%%%%%%%%%%%%%%%%%%%%%%%%%%%%%%%%%%%%%%%%%%%%%%%%%%%
% Sample loads, weights of loads, molecule and background photon emission rates %
%%%%%%%%%%%%%%%%%%%%%%%%%%%%%%%%%%%%%%%%%%%%%%%%%%%%%%%%%%%%%%%%%%%%%%%%%%%%%%%%%

% Set the new samples equal to the old ones
mu(:,i)=mu(:,i-1);
mu_back(i)=mu_back(i-1);
b(:,i)=b(:,i-1);

% Since we sample mu and mu_back through a metropolis-hastings, we sample
% them several times to have a better mixture.
switch PSF_func
         
    case 1     % 3D Gaussian PSF
                   constant= exp(-2*(x(1:3:3*Number_molecules*Max_Species,:).^2+...
                                     x(2:3:3*Number_molecules*Max_Species,:).^2+...
                                     x(3:3:3*Number_molecules*Max_Species,:).^2));
    case 2      % 2D Gaussian Lorentzian PSF
                   constant= exp(-2*(x(1:3:3*Number_molecules*Max_Species,:).^2+...
                                     x(2:3:3*Number_molecules*Max_Species,:).^2)./...
                                  (1+x(3:3:3*Number_molecules*Max_Species,:).^2))./...
                                  (1+x(3:3:3*Number_molecules*Max_Species,:).^2) ;
    otherwise     % Cylindrical
                   constant= exp(-2*(x(1:3:3*Number_molecules*Max_Species,:).^2+...
                                     x(2:3:3*Number_molecules*Max_Species,:).^2));
end       

           for k=1:100    
           %%%%%%%%%%%%%%%%%%%%%%%%%%%%%%%%%%%%%%%%%%%%%%%%%%%%%%%%%%%%%%%%
           %%% Sampling molecule and background photon emission rates %%%%%
           %%%%%%%%%%%%%%%%%%%%%%%%%%%%%%%%%%%%%%%%%%%%%%%%%%%%%%%%%%%%%%%%    
%            [ mu(:,i) , mu_back(i) , acceptance_rate_mu ] = Sample_emission( ...
%            ...
%            ...
%            mu_proposal         , mu_back_proposal , mu_alpha  , mu_beta     , ...
%            mu_back_alpha       , mu_back_beta     , mu(:,i)     , mu_back(i)  , ...
%            acceptance_rate_mu  , observation      , Time      , b(:,i)      , ...
%            constant            , Number_molecules , Max_Species);
    
           %%%%%%%%%%%%%%%%%%%%%%%%%%%%%%%%%%%%%%%%%%%%%%%%%%%%%%%%%%%%%%%%
           %%%%%%%% Sampling the weights over loads (hyper priors) %%%%%%%%
           %%%%%%%%%%%%%%%%%%%%%%%%%%%%%%%%%%%%%%%%%%%%%%%%%%%%%%%%%%%%%%%%
           
           % This step will be a direct sampling from a beta distribution q ~ p(b|q) p(q)
           q(:,i) = betarnd(...
                 (alpha_q/(Number_molecules))+b(:,i),...
                 (beta_q*(Number_molecules-1)/(Number_molecules))+1-b(:,i)...
                 );
           
           %%%%%%%%%%%%%%%%%%%%%%%%%%%%%%%%%%%%%%%%%%%%%%%%%%%%%%%%%%%%%%%%
           %%%%%%%%%%%%%%%%%%% Sampling the loads  %%%%%%%%%%%%%%%%%%%%%%%%
           %%%%%%%%%%%%%%%%%%%%%%%%%%%%%%%%%%%%%%%%%%%%%%%%%%%%%%%%%%%%%%%%
              for m=1:Max_Species
                 [ b(:,i) , acceptance_rate_b ] = Sample_loads( ...
                  ...
                  ...
                  b(:,i)       , q(:,i)             , mu(:,i) , mu_back(i)        , ...
                  observation  , acceptance_rate_b  , Time    , Number_molecules  , ...
                  Max_Species  , constant           , m      ); 
              end
           
           end

%%%%%%%%%%%%%%%%%%%%%%%%%%%%%%%%%%%%%%%%%%%%%%%%%%%%%%%%%%%%%%%%%%%%%%%%%%% 
%%%%%%%%%%%%%%%%%%%%%%%%%%%%%% Save  data %%%%%%%%%%%%%%%%%%%%%%%%%%%%%%%%%
%%%%%%%%%%%%%%%%%%%%%%%%%%%%%%%%%%%%%%%%%%%%%%%%%%%%%%%%%%%%%%%%%%%%%%%%%%% 
           if find((size_iter_old+floor(save_bar.*Max_iter./save_size))==i)>=1
               
               % Plot the remaining time
                 time_loop=time_loop+toc;
                 remaining_time=floor((time_loop/(save_bar.*Max_iter./save_size))*((save_size-save_bar).*Max_iter./save_size)*10)/10;
                 
                 waitbar(save_bar/save_size,hbar,['Remaining time = ', num2str(remaining_time),' s']);
                 save_bar=save_bar+1;
                 tic
                
               % Calculate the acceptance rates of the loads and emission rates metropolis
                 acceptance_rate_muu  = acceptance_rate_mu / (10*(i-1))  ;
                 acceptance_rate_bb   = acceptance_rate_b  / (10*Max_Species*(i-1))  ;

               % Save the results based on the sive size
                 save('results' , 'D'                   , 'mu'                 , 'mu_back'   ...
                                , 'b'                   , 'concentration'      , 'x'         ...
                                , 'acceptance_rate_muu' , 'acceptance_rate_bb' , 'q'         ...
                                , 'Time'                , 'trace'              )
           end
           
           
           
           
    end
    % Terminate the time bar
     delete(hbar);
     
     
    % Normalize the acceptance rates and the concentration of teh particels
    acceptance_rate_mu  = acceptance_rate_mu / (10*(Max_iter-1))  ;
    acceptance_rate_b   = acceptance_rate_b  / (10*(Max_iter-1))  ;
end
```
