## Supplementary material for "An Alternative Framework for Fluorescence Correlation Spectroscopy": Source Code: Inactive_molecules.m

```
function[ x  ] = Inactive_molecules( ...
...
x           , sign_siz     , Number_molecules , Time          , ...
b           , mu_prior_xy  , mu_prior_z       , var_prior_xy  , ...
var_prior_z , wxy          , wz               , D             , ...
Max_species )   

%%%%%%%%%%%%%%%%%%%%%%%%%%%%%%%%%%%%%%%%%%%%%%%%%%%%%%%%%%%%%%%%%%%%%%%%%%%
%%%%%%%%%%%%%%%%%%%%%%%%%%%%%%% INPUT %%%%%%%%%%%%%%%%%%%%%%%%%%%%%%%%%%%%%
%%%%%%%%%%%%%%%%%%%%%%%%%%%%%%%%%%%%%%%%%%%%%%%%%%%%%%%%%%%%%%%%%%%%%%%%%%%
     % x                             :  Positions of the particles (Normalized)
     % sign_siz                      :  Length of the observation trace
     % Number_molecules              :  Number of defined molecules for each species
     % Time                          :  Data acquisition time
     % b                             :  Loads for active particles
     % mu_prior_xy                   :  Mean value for X0 prior in normalized scale for x and y coordintes
     % mu_prior_z                    :  Mean value for X0 prior in normalized scale for z coordinte
     % var_prior_xy                  :  Variance for X0 prior in normalized scale
     % var_prior_z                   :  Variance for X0 prior in normalized scale
     % wxy                           :  Minor Semi-axis of the confocal volume
     % wz                            :  Major Semi-axis of the confocal volume
     % D                             :  Values for Diffusion coefficient

%%%%%%%%%%%%%%%%%%%%%%%%%%%%%%%%%%%%%%%%%%%%%%%%%%%%%%%%%%%%%%%%%%%%%%%%%%%
%%%%%%%%%%%%%%%%%%%%%%%%%%%%%% OUTPUT %%%%%%%%%%%%%%%%%%%%%%%%%%%%%%%%%%%%%
%%%%%%%%%%%%%%%%%%%%%%%%%%%%%%%%%%%%%%%%%%%%%%%%%%%%%%%%%%%%%%%%%%%%%%%%%%%
     % x                             :  Positions of the particles (Normalized)
  
kxy = sqrt(2*bsxfun(@times,D,Time(2:end)))/wxy                            ;
kz  = kxy*(wxy/wz)                                                        ;

for m=1:Max_species
    for n=1:Number_molecules
         if b((m-1)*Number_molecules +n)==0
                x(3*Number_molecules*(m-1)+3*n-2 , :)   = ...
                    cumsum([((-1)^(floor(2*rand())))*mu_prior_xy + sqrt(var_prior_xy)*randn(), ...
                                    kxy(m,:).*randn(1,sign_siz-1)])       ;
                                
                x(3*Number_molecules*(m-1)+3*n-1 , :)   = ...
                    cumsum([((-1)^(floor(2*rand())))*mu_prior_xy + sqrt(var_prior_xy)*randn(), ...
                                    kxy(m,:).*randn(1,sign_siz-1)])       ;
                                
                x(3*Number_molecules*(m-1)+3*n   , :)   = ...
                    cumsum([((-1)^(floor(2*rand())))*mu_prior_z  + sqrt(var_prior_z) *randn(), ...
                                    kz(m,:).*randn(1,sign_siz-1)])       ; 
        end
    end
end

end
```
