## Supplementary material for "An Alternative Framework for Fluorescence Correlation Spectroscopy": Source Code: nwspgr.m

```
%*****************************************************************************
%nwSpGr: Nodes and weights for numerical integration on sparse grids (Smolyak)
%(c) 2007 Florian Heiss, Viktor Winschel
%Nov 11, 2007
%*****************************************************************************
%************************************************************************************
%nwspgr(): main function for generating nodes & weights for sparse grids intergration 
%Syntax: 
%    [n w] = nwSpGr(type, dim, k, sym)
%Input: 
%    type = String for type of 1D integration rule:
%           "KPU": Nested rule for unweighted integral over [0,1]
%           "KPN": Nested rule for integral with Gaussian weight
%           "GQU": Gaussian quadrature for unweighted integral over [0,1] (Gauss-Legendre)
%           "GQN": Gaussian quadrature for integral with Gaussian weight (Gauss-Hermite)
%           func:  any function name. Function must accept level l and
%                  return nodes n and weights w for univariate quadrature rule with
%                  polynomial exactness 2l-1 as [n w] = feval(func,level)
%                  Example: If you have installed the CompEcon Toolbox of Paul Fackler and Mario Miranda
%                           (http://www4.ncsu.edu/~pfackler/compecon/toolbox.html)
%                           and you want to integrate with a lognormal weight function, you can use
%                           nwSpGr('qnwlogn', dim, k)
%    dim  : dimension of the integration problem
%    k    : Accuracy level. The rule will be exact for polynomial up to total order 2k-1
%    sym  : (optional) only used for own 1D quadrature rule (type not "KPU",...). If
%           sym is supplied and not=0, the code will run faster but will
%           produce incorrect results if 1D quadrature rule is asymmetric
%Output:
%    n    = matrix of nodes with dim columns 
%    w    = row vector of corresponding weights
%*************************************************************************************
function [nodes, weights] = nwspgr(type, dim, k, sym)

  % interpret inputs
  if (nargin < 3); error('not enough input arguments')
  end
  builtinfct = (strcmp(type,'GQU') || strcmp(type,'GQN') || strcmp(type,'KPU') || strcmp(type,'KPN'));
  if (nargin == 3)
      if builtinfct; sym = true;
      else sym = false;
      end
  else sym = logical(sym);
  end
  
  % get 1D nodes & weights
  try 
      n1D = cell(k,1); w1D = cell(k,1); R1D = zeros(1,k);
      for level=1:k
        [n w] = feval(type,level);
        % if user supplied symmetric 1D rule: keep only positive orthant
        if ~builtinfct && sym
            numnew = rows(n);
            [n sortvec] = sortrows(n);
            w = w(sortvec);
            n = n((floor(numnew/2)+1):numnew,:);
            w = w((floor(numnew/2)+1):numnew,:);
        end
        R1D(level) = length(w);
        n1D{level} = n; w1D{level} = w;
      end
  catch error('Error evaluating the 1D rule');      
  end
  
  % initialization
  minq = max(0,k-dim);
  maxq = k-1;
  nodes = []; weights = [];
  % outer loop over q
  for q = minq:maxq
    r = length(weights);
    bq = (-1)^(maxq-q) * nchoosek(dim-1,dim+q-k);
    % matrix of all rowvectors in N^D_{q}
    is = SpGrGetSeq(dim,dim+q);
    % preallocate new rows for nodes & weights
    Rq = prod(R1D(is),2);
    sRq = sum(Rq);
    nodes = [nodes ; zeros(sRq,dim)];
    weights = [weights ; zeros(sRq,1)];
    % inner loop collecting product rules
    for j=1:size(is,1)
      midx = is(j,:);
      [newn,neww] = SpGrKronProd(n1D(midx),w1D(midx));
      nodes((r+1):(r+Rq(j)),:) = newn;
      weights((r+1):(r+Rq(j))) = bq .* neww;
      r = r + Rq(j);
    end
    % collect equal nodes: first sort
    [nodes sortvec] = sortrows(nodes);
    weights = weights(sortvec);
    keep = 1; 
    lastkeep = 1;
    % then make a list of rows to keep and sum weights of equal nodes
    for j=2:size(nodes,1)
      if nodes(j,:)==nodes(j-1,:) 
          weights(lastkeep)=weights(lastkeep)+weights(j);
      else
        lastkeep = j;
        keep = [keep ; j ];
      end
    end
    % drop equal rows
    nodes = nodes(keep,:);
    weights = weights(keep);
  end
  
  % If symmetric rule: so far, it's for the positive orthant. Copy to other
  % orthants!
  if sym
	  nr = length(weights);
    m = n1D{1};
		for j=1:dim
			keep = zeros(nr,1);
			numnew = 0;
		  for r=1:nr 
        if nodes(r,j) ~= m
  				numnew=numnew+1;
    			keep(numnew)=r;
        end
      end
      if numnew>0
				nodes = [nodes ; nodes(keep(1:numnew),:)];
				nodes(nr+1:nr+numnew,j) = 2*m - nodes(nr+1:nr+numnew,j);
				weights = [weights ; weights(keep(1:numnew))]; 
				nr=nr+numnew;
      end	
    end
	  % 3. final sorting
    [nodes sortvec] = sortrows(nodes);
    weights = weights(sortvec);
  end
  
  % normalize weights to account for rounding errors
  weights = weights / sum(weights);
end

%**************************************************************************************
%SpGrGetSeq(): function for generating matrix of all rowvectors in N^D_{norm} 
%Syntax: 
%    out = nwSpGr(d,norm)
%Input: 
%    d    = dimension, will be #columns in output
%    norm = row sum of elements each of the rows has to have
%Output:
%    out  = matrix with d columns. Each row represents one vector with all elements >=1
%           and the sum of elements == norm
%**************************************************************************************
function fs = SpGrGetSeq(d, norm)
  seq = zeros(1,d);
  a=norm-d;
  seq(1)=a;
  fs = seq;
  c=1;
  while seq(d)<a
    if (c==d) 
        for i=(c-1):-1:1
            c=i;
            if seq(i)~=0, break, end;
        end
    end
    seq(c) = seq(c)-1;
    c=c+1;
    seq(c) = a - sum(seq(1:(c-1)));
    if (c<d) 
        seq((c+1):d)=zeros(1,d-c);
    end
    fs = [fs;seq];
  end 
  fs = fs+1;
end

%************************************************************************************
%SpGrKronProd(): function for generating tensor product quadrature rule 
%Syntax: 
%    [nodes,weights] = SpGrKronProd(n1d,w1D)
%Input: 
%    n1D = cell array of 1D nodes  
%    w1D = cell array of 1D weights 
%Output:
%    nodes   = matrix of tensor product nodes 
%    weights = vector of tensor product weights 
%*************************************************************************************

function [nodes,weights] = SpGrKronProd(n1D,w1D)
  nodes = n1D{1} ; weights = w1D{1};
  for j=2:length(n1D)
    newnodes = n1D{j};
    nodes = [kron(nodes,ones(size(newnodes,1),1)) kron(ones(size(nodes,1),1),newnodes)];
    weights = kron(weights,w1D{j});
  end
end


function [n w] = GQU(l)
  switch l
  case 1
    n = [5.0000000000000000e-001];
    w = [1.0000000000000000e+000];
  case 2
    n = [7.8867513459481287e-001];
    w = [5.0000000000000000e-001];
  case 3
    n = [5.0000000000000000e-001; 8.8729833462074170e-001];
    w = [4.4444444444444570e-001; 2.7777777777777712e-001];
  case 4
    n = [6.6999052179242813e-001; 9.3056815579702623e-001];
    w = [3.2607257743127516e-001; 1.7392742256872484e-001];
  case 5
    n = [5.0000000000000000e-001; 7.6923465505284150e-001; 9.5308992296933193e-001];
    w = [2.8444444444444655e-001; 2.3931433524968501e-001; 1.1846344252809174e-001];
  case 6
    n = [6.1930959304159849e-001; 8.3060469323313235e-001; 9.6623475710157603e-001];
    w = [2.3395696728634746e-001; 1.8038078652407072e-001; 8.5662246189581834e-002];
  case 7
    n = [5.0000000000000000e-001; 7.0292257568869854e-001; 8.7076559279969723e-001; 9.7455395617137919e-001];
    w = [2.0897959183673620e-001; 1.9091502525256090e-001; 1.3985269574463935e-001; 6.4742483084431701e-002];
  case 8
    n = [5.9171732124782495e-001; 7.6276620495816450e-001; 8.9833323870681348e-001; 9.8014492824876809e-001];
    w = [1.8134189168918213e-001; 1.5685332293894469e-001; 1.1119051722668793e-001; 5.0614268145185180e-002];
  case 9
    n = [5.0000000000000000e-001; 6.6212671170190451e-001; 8.0668571635029518e-001; 9.1801555366331788e-001; 9.8408011975381304e-001];
    w = [1.6511967750063075e-001; 1.5617353852000226e-001; 1.3030534820146844e-001; 9.0324080347429253e-002; 4.0637194180784583e-002];
  case 10
    n = [5.7443716949081558e-001; 7.1669769706462361e-001; 8.3970478414951222e-001; 9.3253168334449232e-001; 9.8695326425858587e-001];
    w = [1.4776211235737713e-001; 1.3463335965499873e-001; 1.0954318125799158e-001; 7.4725674575290599e-002; 3.3335672154342001e-002];
  case 11
    n = [5.0000000000000000e-001; 6.3477157797617245e-001; 7.5954806460340585e-001; 8.6507600278702468e-001; 9.4353129988404771e-001; 9.8911432907302843e-001];
    w = [1.3646254338895086e-001; 1.3140227225512388e-001; 1.1659688229599563e-001; 9.3145105463867520e-002; 6.2790184732452625e-002; 2.7834283558084916e-002];
  case 12
    n = [5.6261670425573451e-001; 6.8391574949909006e-001; 7.9365897714330869e-001; 8.8495133709715235e-001; 9.5205862818523745e-001; 9.9078031712335957e-001];
    w = [1.2457352290670189e-001; 1.1674626826917781e-001; 1.0158371336153328e-001; 8.0039164271673444e-002; 5.3469662997659276e-002; 2.3587668193254314e-002];
  case 13
    n = [5.0000000000000000e-001; 6.1522915797756739e-001; 7.2424637551822335e-001; 8.2117466972017006e-001; 9.0078904536665494e-001; 9.5879919961148907e-001; 9.9209152735929407e-001];
    w = [1.1627577661543741e-001; 1.1314159013144903e-001; 1.0390802376844462e-001; 8.9072990380973202e-002; 6.9436755109893875e-002; 4.6060749918864378e-002; 2.0242002382656228e-002];
  case 14
    n = [5.5402747435367183e-001; 6.5955618446394482e-001; 7.5762431817907705e-001; 8.4364645240584268e-001; 9.1360065753488251e-001; 9.6421744183178681e-001; 9.9314190434840621e-001];
    w = [1.0763192673157916e-001; 1.0259923186064811e-001; 9.2769198738969161e-002; 7.8601583579096995e-002; 6.0759285343951711e-002; 4.0079043579880291e-002; 1.7559730165874574e-002];
  case 15
    n = [5.0000000000000000e-001; 6.0059704699871730e-001; 6.9707567353878175e-001; 7.8548608630426942e-001; 8.6220886568008503e-001; 9.2410329170521366e-001; 9.6863669620035298e-001; 9.9399625901024269e-001];
    w = [1.0128912096278091e-001; 9.9215742663556039e-002; 9.3080500007781286e-002; 8.3134602908497196e-002; 6.9785338963077315e-002; 5.3579610233586157e-002; 3.5183023744054159e-002; 1.5376620998057434e-002];
  case 16
    n = [5.4750625491881877e-001; 6.4080177538962946e-001; 7.2900838882861363e-001; 8.0893812220132189e-001; 8.7770220417750155e-001; 9.3281560119391593e-001; 9.7228751153661630e-001; 9.9470046749582497e-001];
    w = [9.4725305227534431e-002; 9.1301707522462000e-002; 8.4578259697501462e-002; 7.4797994408288562e-002; 6.2314485627767105e-002; 4.7579255841246545e-002; 3.1126761969323954e-002; 1.3576229705875955e-002];
  case 17
    n = [5.0000000000000000e-001; 5.8924209074792389e-001; 6.7561588172693821e-001; 7.5634526854323847e-001; 8.2883557960834531e-001; 8.9075700194840068e-001; 9.4011957686349290e-001; 9.7533776088438384e-001; 9.9528773765720868e-001];
    w = [8.9723235178103419e-002; 8.8281352683496447e-002; 8.4002051078225143e-002; 7.7022880538405308e-002; 6.7568184234262890e-002; 5.5941923596702053e-002; 4.2518074158589644e-002; 2.7729764686993612e-002; 1.2074151434273140e-002];
  case 18
    n = [5.4238750652086765e-001; 6.2594311284575277e-001; 7.0587558073142131e-001; 7.7988541553697377e-001; 8.4584352153017661e-001; 9.0185247948626157e-001; 9.4630123324877791e-001; 9.7791197478569880e-001; 9.9578258421046550e-001];
    w = [8.4571191481571939e-002; 8.2138241872916504e-002; 7.7342337563132801e-002; 7.0321457335325452e-002; 6.1277603355739306e-002; 5.0471022053143716e-002; 3.8212865127444665e-002; 2.4857274447484968e-002; 1.0808006763240719e-002];
  case 19
    n = [5.0000000000000000e-001; 5.8017932282011264e-001; 6.5828204998181494e-001; 7.3228537068798050e-001; 8.0027265233084055e-001; 8.6048308866761469e-001; 9.1135732826857141e-001; 9.5157795180740901e-001; 9.8010407606741501e-001; 9.9620342192179212e-001];
    w = [8.0527224924391946e-002; 7.9484421696977337e-002; 7.6383021032929960e-002; 7.1303351086803413e-002; 6.4376981269668232e-002; 5.5783322773667113e-002; 4.5745010811225124e-002; 3.4522271368820669e-002; 2.2407113382849821e-002; 9.7308941148624341e-003];
  case 20
    n = [5.3826326056674867e-001; 6.1389292557082253e-001; 6.8685304435770977e-001; 7.5543350097541362e-001; 8.1802684036325757e-001; 8.7316595323007540e-001; 9.1955848591110945e-001; 9.5611721412566297e-001; 9.8198596363895696e-001; 9.9656429959254744e-001];
    w = [7.6376693565363113e-002; 7.4586493236301996e-002; 7.1048054659191187e-002; 6.5844319224588346e-002; 5.9097265980759248e-002; 5.0965059908620318e-002; 4.1638370788352433e-002; 3.1336024167054569e-002; 2.0300714900193556e-002; 8.8070035695753026e-003];
  case 21
    n = [5.0000000000000000e-001; 5.7278092708044759e-001; 6.4401065840120053e-001; 7.1217106010371944e-001; 7.7580941794360991e-001; 8.3356940209870611e-001; 8.8421998173783889e-001; 9.2668168229165859e-001; 9.6004966707520034e-001; 9.8361341928315316e-001; 9.9687608531019478e-001];
    w = [7.3040566824845346e-002; 7.2262201994985134e-002; 6.9943697395536658e-002; 6.6134469316668845e-002; 6.0915708026864350e-002; 5.4398649583574356e-002; 4.6722211728016994e-002; 3.8050056814189707e-002; 2.8567212713428641e-002; 1.8476894885426285e-002; 8.0086141288864491e-003];
  case 22
    n = [5.3486963665986109e-001; 6.0393021334411068e-001; 6.7096791044604209e-001; 7.3467791899337853e-001; 7.9382020175345580e-001; 8.4724363159334137e-001; 8.9390840298960406e-001; 9.3290628886015003e-001; 9.6347838609358694e-001; 9.8503024891771429e-001; 9.9714729274119962e-001];
    w = [6.9625936427816129e-002; 6.8270749173007697e-002; 6.5586752393531317e-002; 6.1626188405256251e-002; 5.6466148040269712e-002; 5.0207072221440600e-002; 4.2970803108533975e-002; 3.4898234212260300e-002; 2.6146667576341692e-002; 1.6887450792407110e-002; 7.3139976491353280e-003];
  case 23
    n = [5.0000000000000000e-001; 5.6662841214923310e-001; 6.3206784048517251e-001; 6.9515051901514546e-001; 7.5475073892300371e-001; 8.0980493788182306e-001; 8.5933068156597514e-001; 9.0244420080942001e-001; 9.3837617913522076e-001; 9.6648554341300807e-001; 9.8627123560905761e-001; 9.9738466749877608e-001];
    w = [6.6827286093053176e-002; 6.6231019702348404e-002; 6.4452861094041150e-002; 6.1524542153364815e-002; 5.7498320111205814e-002; 5.2446045732270824e-002; 4.6457883030017563e-002; 3.9640705888359551e-002; 3.2116210704262994e-002; 2.4018835865542369e-002; 1.5494002928489686e-002; 6.7059297435702412e-003];
  case 24
    n = [5.3202844643130276e-001; 5.9555943373680820e-001; 6.5752133984808170e-001; 7.1689675381302254e-001; 7.7271073569441984e-001; 8.2404682596848777e-001; 8.7006209578927718e-001; 9.1000099298695147e-001; 9.4320776350220048e-001; 9.6913727600136634e-001; 9.8736427798565474e-001; 9.9759360999851066e-001];
    w = [6.3969097673376246e-002; 6.2918728173414318e-002; 6.0835236463901793e-002; 5.7752834026862883e-002; 5.3722135057982914e-002; 4.8809326052057039e-002; 4.3095080765976693e-002; 3.6673240705540205e-002; 2.9649292457718385e-002; 2.2138719408709880e-002; 1.4265694314466934e-002; 6.1706148999928351e-003];
  case 25
    n = [5.0000000000000000e-001; 5.6143234630535521e-001; 6.2193344186049426e-001; 6.8058615290469393e-001; 7.3650136572285752e-001; 7.8883146512061142e-001; 8.3678318423673415e-001; 8.7962963151867890e-001; 9.1672131438041693e-001; 9.4749599893913761e-001; 9.7148728561448716e-001; 9.8833196072975871e-001; 9.9777848489524912e-001];
    w = [6.1588026863357799e-002; 6.1121221495155122e-002; 5.9727881767892461e-002; 5.7429129572855862e-002; 5.4259812237131867e-002; 5.0267974533525363e-002; 4.5514130991481903e-002; 4.0070350167500532e-002; 3.4019166906178545e-002; 2.7452347987917691e-002; 2.0469578350653148e-002; 1.3177493307516108e-002; 5.6968992505125535e-003];
  end
end
function [n w] = GQN(l)
  switch l
  case 1
    n = [0.0000000000000000e+000];
    w = [1.0000000000000000e+000];
  case 2
    n = [1.0000000000000002e+000];
    w = [5.0000000000000000e-001];
  case 3
    n = [0.0000000000000000e+000; 1.7320508075688772e+000];
    w = [6.6666666666666663e-001; 1.6666666666666674e-001];
  case 4
    n = [7.4196378430272591e-001; 2.3344142183389773e+000];
    w = [4.5412414523193145e-001; 4.5875854768068498e-002];
  case 5
    n = [0.0000000000000000e+000; 1.3556261799742659e+000; 2.8569700138728056e+000];
    w = [5.3333333333333344e-001; 2.2207592200561263e-001; 1.1257411327720691e-002];
  case 6
    n = [6.1670659019259422e-001; 1.8891758777537109e+000; 3.3242574335521193e+000];
    w = [4.0882846955602919e-001; 8.8615746041914523e-002; 2.5557844020562431e-003];
  case 7
    n = [0.0000000000000000e+000; 1.1544053947399682e+000; 2.3667594107345415e+000; 3.7504397177257425e+000];
    w = [4.5714285714285757e-001; 2.4012317860501250e-001; 3.0757123967586491e-002; 5.4826885597221875e-004];
  case 8
    n = [5.3907981135137517e-001; 1.6365190424351082e+000; 2.8024858612875416e+000; 4.1445471861258945e+000];
    w = [3.7301225767907736e-001; 1.1723990766175897e-001; 9.6352201207882630e-003; 1.1261453837536784e-004];
  case 9
    n = [0.0000000000000000e+000; 1.0232556637891326e+000; 2.0768479786778302e+000; 3.2054290028564703e+000; 4.5127458633997826e+000];
    w = [4.0634920634920685e-001; 2.4409750289493909e-001; 4.9916406765217969e-002; 2.7891413212317675e-003; 2.2345844007746563e-005];
  case 10
    n = [4.8493570751549764e-001; 1.4659890943911582e+000; 2.4843258416389546e+000; 3.5818234835519269e+000; 4.8594628283323127e+000];
    w = [3.4464233493201940e-001; 1.3548370298026730e-001; 1.9111580500770317e-002; 7.5807093431221972e-004; 4.3106526307183106e-006];
  case 11
    n = [0.0000000000000000e+000; 9.2886899738106388e-001; 1.8760350201548459e+000; 2.8651231606436447e+000; 3.9361666071299775e+000; 5.1880012243748714e+000];
    w = [3.6940836940836957e-001; 2.4224029987397003e-001; 6.6138746071057644e-002; 6.7202852355372697e-003; 1.9567193027122324e-004; 8.1218497902149036e-007];
  case 12
    n = [4.4440300194413901e-001; 1.3403751971516167e+000; 2.2594644510007993e+000; 3.2237098287700974e+000; 4.2718258479322815e+000; 5.5009017044677480e+000];
    w = [3.2166436151283007e-001; 1.4696704804532995e-001; 2.9116687912364138e-002; 2.2033806875331849e-003; 4.8371849225906076e-005; 1.4999271676371597e-007];
  case 13
    n = [0.0000000000000000e+000; 8.5667949351945005e-001; 1.7254183795882394e+000; 2.6206899734322149e+000; 3.5634443802816347e+000; 4.5913984489365207e+000; 5.8001672523865011e+000];
    w = [3.4099234099234149e-001; 2.3787152296413588e-001; 7.9168955860450141e-002; 1.1770560505996543e-002; 6.8123635044292619e-004; 1.1526596527333885e-005; 2.7226276428059039e-008];
  case 14
    n = [4.1259045795460181e-001; 1.2426889554854643e+000; 2.0883447457019444e+000; 2.9630365798386675e+000; 3.8869245750597696e+000; 4.8969363973455646e+000; 6.0874095469012914e+000];
    w = [3.0263462681301945e-001; 1.5408333984251366e-001; 3.8650108824253432e-002; 4.4289191069474066e-003; 2.0033955376074381e-004; 2.6609913440676334e-006; 4.8681612577483872e-009];
  case 15
    n = [0.0000000000000000e+000; 7.9912906832454811e-001; 1.6067100690287301e+000; 2.4324368270097581e+000; 3.2890824243987664e+000; 4.1962077112690155e+000; 5.1900935913047821e+000; 6.3639478888298378e+000];
    w = [3.1825951825951820e-001; 2.3246229360973222e-001; 8.9417795399844444e-002; 1.7365774492137616e-002; 1.5673575035499571e-003; 5.6421464051890157e-005; 5.9754195979205961e-007; 8.5896498996331805e-010];
  case 16
    n = [3.8676060450055738e-001; 1.1638291005549648e+000; 1.9519803457163336e+000; 2.7602450476307019e+000; 3.6008736241715487e+000; 4.4929553025200120e+000; 5.4722257059493433e+000; 6.6308781983931295e+000];
    w = [2.8656852123801241e-001; 1.5833837275094925e-001; 4.7284752354014067e-002; 7.2669376011847411e-003; 5.2598492657390979e-004; 1.5300032162487286e-005; 1.3094732162868203e-007; 1.4978147231618314e-010];
  case 17
    n = [0.0000000000000000e+000; 7.5184260070389630e-001; 1.5098833077967408e+000; 2.2810194402529889e+000; 3.0737971753281941e+000; 3.9000657171980104e+000; 4.7785315896299840e+000; 5.7444600786594071e+000; 6.8891224398953330e+000];
    w = [2.9953837012660756e-001; 2.2670630846897877e-001; 9.7406371162718081e-002; 2.3086657025711152e-002; 2.8589460622846499e-003; 1.6849143155133945e-004; 4.0126794479798725e-006; 2.8080161179305783e-008; 2.5843149193749151e-011];
  case 18
    n = [3.6524575550769767e-001; 1.0983955180915013e+000; 1.8397799215086457e+000; 2.5958336889112403e+000; 3.3747365357780907e+000; 4.1880202316294044e+000; 5.0540726854427405e+000; 6.0077459113595975e+000; 7.1394648491464796e+000];
    w = [2.7278323465428789e-001; 1.6068530389351263e-001; 5.4896632480222654e-002; 1.0516517751941352e-002; 1.0654847962916496e-003; 5.1798961441161962e-005; 1.0215523976369816e-006; 5.9054884788365484e-009; 4.4165887693587078e-012];
  case 19
    n = [0.0000000000000000e+000; 7.1208504404237993e-001; 1.4288766760783731e+000; 2.1555027613169351e+000; 2.8980512765157536e+000; 3.6644165474506383e+000; 4.4658726268310316e+000; 5.3205363773360386e+000; 6.2628911565132519e+000; 7.3825790240304316e+000];
    w = [2.8377319275152108e-001; 2.2094171219914366e-001; 1.0360365727614400e-001; 2.8666691030118496e-002; 4.5072354203420355e-003; 3.7850210941426759e-004; 1.5351145954666744e-005; 2.5322200320928681e-007; 1.2203708484474786e-009; 7.4828300540572308e-013];
  case 20
    n = [3.4696415708135592e-001; 1.0429453488027509e+000; 1.7452473208141270e+000; 2.4586636111723679e+000; 3.1890148165533900e+000; 3.9439673506573163e+000; 4.7345813340460552e+000; 5.5787388058932015e+000; 6.5105901570136551e+000; 7.6190485416797591e+000];
    w = [2.6079306344955544e-001; 1.6173933398400026e-001; 6.1506372063976029e-002; 1.3997837447101043e-002; 1.8301031310804918e-003; 1.2882627996192898e-004; 4.4021210902308646e-006; 6.1274902599829597e-008; 2.4820623623151838e-010; 1.2578006724379305e-013];
  case 21
    n = [0.0000000000000000e+000; 6.7804569244064405e-001; 1.3597658232112304e+000; 2.0491024682571628e+000; 2.7505929810523733e+000; 3.4698466904753764e+000; 4.2143439816884216e+000; 4.9949639447820253e+000; 5.8293820073044706e+000; 6.7514447187174609e+000; 7.8493828951138225e+000];
    w = [2.7026018357287707e-001; 2.1533371569505982e-001; 1.0839228562641938e-001; 3.3952729786542839e-002; 6.4396970514087768e-003; 7.0804779548153736e-004; 4.2192347425515866e-005; 1.2253548361482522e-006; 1.4506612844930740e-008; 4.9753686041217464e-011; 2.0989912195656652e-014];
  case 22
    n = [3.3117931571527381e-001; 9.9516242227121554e-001; 1.6641248391179071e+000; 2.3417599962877080e+000; 3.0324042278316763e+000; 3.7414963502665177e+000; 4.4763619773108685e+000; 5.2477244337144251e+000; 6.0730749511228979e+000; 6.9859804240188152e+000; 8.0740299840217116e+000];
    w = [2.5024359658693501e-001; 1.6190629341367538e-001; 6.7196311428889891e-002; 1.7569072880805774e-002; 2.8087610475772107e-003; 2.6228330325596416e-004; 1.3345977126808712e-005; 3.3198537498140043e-007; 3.3665141594582109e-009; 9.8413789823460105e-012; 3.4794606478771428e-015];
  case 23
    n = [0.0000000000000000e+000; 6.4847115353449580e-001; 1.2998764683039790e+000; 1.9573275529334242e+000; 2.6243236340591820e+000; 3.3050400217529652e+000; 4.0047753217333044e+000; 4.7307241974514733e+000; 5.4934739864717947e+000; 6.3103498544483996e+000; 7.2146594350518622e+000; 8.2933860274173536e+000];
    w = [2.5850974080883904e-001; 2.0995966957754261e-001; 1.1207338260262091e-001; 3.8867183703480947e-002; 8.5796783914656640e-003; 1.1676286374978613e-003; 9.3408186090312983e-005; 4.0899772449921549e-006; 8.7750624838617161e-008; 7.6708888623999076e-010; 1.9229353115677913e-012; 5.7323831678020873e-016];
  case 24
    n = [3.1737009662945231e-001; 9.5342192293210926e-001; 1.5934804298164202e+000; 2.2404678516917524e+000; 2.8977286432233140e+000; 3.5693067640735610e+000; 4.2603836050199053e+000; 4.9780413746391208e+000; 5.7327471752512009e+000; 6.5416750050986341e+000; 7.4378906660216630e+000; 8.5078035191952583e+000];
    w = [2.4087011554664056e-001; 1.6145951286700025e-001; 7.2069364017178436e-002; 2.1126344408967029e-002; 3.9766089291813113e-003; 4.6471871877939763e-004; 3.2095005652745989e-005; 1.2176597454425830e-006; 2.2674616734804651e-008; 1.7186649279648690e-010; 3.7149741527624159e-013; 9.3901936890419202e-017];
  case 25
    n = [0.0000000000000000e+000; 6.2246227918607611e-001; 1.2473119756167892e+000; 1.8770583699478387e+000; 2.5144733039522058e+000; 3.1627756793881927e+000; 3.8259005699724917e+000; 4.5089299229672850e+000; 5.2188480936442794e+000; 5.9660146906067020e+000; 6.7674649638097168e+000; 7.6560379553930762e+000; 8.7175976783995885e+000];
    w = [2.4816935117648548e-001; 2.0485102565034041e-001; 1.1488092430395164e-001; 4.3379970167644971e-002; 1.0856755991462316e-002; 1.7578504052637961e-003; 1.7776690692652660e-004; 1.0672194905202536e-005; 3.5301525602454978e-007; 5.7380238688993763e-009; 3.7911500004771871e-011; 7.1021030370039253e-014; 1.5300389979986825e-017];
  end
end
function [n w] = KPU(l)
  switch l
  case 1
    n = [5.0000000000000000e-001];
    w = [1.0000000000000000e+000];
  case 2
    n = [5.0000000000000000e-001; 8.8729829999999998e-001];
    w = [4.4444440000000002e-001; 2.7777780000000002e-001];
  case 3
    n = [5.0000000000000000e-001; 8.8729829999999998e-001];
    w = [4.4444440000000002e-001; 2.7777780000000002e-001];
  case 4
    n = [5.0000000000000000e-001; 7.1712189999999998e-001; 8.8729829999999998e-001; 9.8024560000000005e-001];
    w = [2.2545832254583223e-001; 2.0069872006987199e-001; 1.3424401342440134e-001; 5.2328105232810521e-002];
  case 5
    n = [5.0000000000000000e-001; 7.1712189999999998e-001; 8.8729829999999998e-001; 9.8024560000000005e-001];
    w = [2.2545832254583223e-001; 2.0069872006987199e-001; 1.3424401342440134e-001; 5.2328105232810521e-002];
  case 6
    n = [5.0000000000000000e-001; 7.1712189999999998e-001; 8.8729829999999998e-001; 9.8024560000000005e-001];
    w = [2.2545832254583223e-001; 2.0069872006987199e-001; 1.3424401342440134e-001; 5.2328105232810521e-002];
  case 7
    n = [5.0000000000000000e-001; 6.1169330000000000e-001; 7.1712189999999998e-001; 8.1055149999999998e-001; 8.8729829999999998e-001; 9.4422960000000000e-001; 9.8024560000000005e-001; 9.9691600000000002e-001];
    w = [1.1275520000000000e-001; 1.0957840000000001e-001; 1.0031430000000000e-001; 8.5755999999999999e-002; 6.7207600000000006e-002; 4.6463600000000001e-002; 2.5801600000000001e-002; 8.5009000000000005e-003];
  case 8
    n = [5.0000000000000000e-001; 6.1169330000000000e-001; 7.1712189999999998e-001; 8.1055149999999998e-001; 8.8729829999999998e-001; 9.4422960000000000e-001; 9.8024560000000005e-001; 9.9691600000000002e-001];
    w = [1.1275520000000000e-001; 1.0957840000000001e-001; 1.0031430000000000e-001; 8.5755999999999999e-002; 6.7207600000000006e-002; 4.6463600000000001e-002; 2.5801600000000001e-002; 8.5009000000000005e-003];
  case 9
    n = [5.0000000000000000e-001; 6.1169330000000000e-001; 7.1712189999999998e-001; 8.1055149999999998e-001; 8.8729829999999998e-001; 9.4422960000000000e-001; 9.8024560000000005e-001; 9.9691600000000002e-001];
    w = [1.1275520000000000e-001; 1.0957840000000001e-001; 1.0031430000000000e-001; 8.5755999999999999e-002; 6.7207600000000006e-002; 4.6463600000000001e-002; 2.5801600000000001e-002; 8.5009000000000005e-003];
  case 10
    n = [5.0000000000000000e-001; 6.1169330000000000e-001; 7.1712189999999998e-001; 8.1055149999999998e-001; 8.8729829999999998e-001; 9.4422960000000000e-001; 9.8024560000000005e-001; 9.9691600000000002e-001];
    w = [1.1275520000000000e-001; 1.0957840000000001e-001; 1.0031430000000000e-001; 8.5755999999999999e-002; 6.7207600000000006e-002; 4.6463600000000001e-002; 2.5801600000000001e-002; 8.5009000000000005e-003];
  case 11
    n = [5.0000000000000000e-001; 6.1169330000000000e-001; 7.1712189999999998e-001; 8.1055149999999998e-001; 8.8729829999999998e-001; 9.4422960000000000e-001; 9.8024560000000005e-001; 9.9691600000000002e-001];
    w = [1.1275520000000000e-001; 1.0957840000000001e-001; 1.0031430000000000e-001; 8.5755999999999999e-002; 6.7207600000000006e-002; 4.6463600000000001e-002; 2.5801600000000001e-002; 8.5009000000000005e-003];
  case 12
    n = [5.0000000000000000e-001; 6.1169330000000000e-001; 7.1712189999999998e-001; 8.1055149999999998e-001; 8.8729829999999998e-001; 9.4422960000000000e-001; 9.8024560000000005e-001; 9.9691600000000002e-001];
    w = [1.1275520000000000e-001; 1.0957840000000001e-001; 1.0031430000000000e-001; 8.5755999999999999e-002; 6.7207600000000006e-002; 4.6463600000000001e-002; 2.5801600000000001e-002; 8.5009000000000005e-003];
  case 13
    n = [5.0000000000000000e-001; 5.5624450000000003e-001; 6.1169330000000000e-001; 6.6556769999999998e-001; 7.1712189999999998e-001; 7.6565989999999995e-001; 8.1055149999999998e-001; 8.5124809999999995e-001; 8.8729829999999998e-001; 9.1836300000000004e-001; 9.4422960000000000e-001; 9.6482740000000000e-001; 9.8024560000000005e-001; 9.9076560000000002e-001; 9.9691600000000002e-001; 9.9954909999999997e-001];
    w = [5.6377600000000014e-002; 5.5978400000000011e-002; 5.4789200000000017e-002; 5.2834900000000011e-002; 5.0157100000000017e-002; 4.6813600000000004e-002; 4.2878000000000006e-002; 3.8439800000000010e-002; 3.3603900000000006e-002; 2.8489800000000006e-002; 2.3231400000000003e-002; 1.7978600000000004e-002; 1.2903800000000003e-002; 8.2230000000000011e-003; 4.2173000000000011e-003; 1.2724000000000001e-003];
  case 14
    n = [5.0000000000000000e-001; 5.5624450000000003e-001; 6.1169330000000000e-001; 6.6556769999999998e-001; 7.1712189999999998e-001; 7.6565989999999995e-001; 8.1055149999999998e-001; 8.5124809999999995e-001; 8.8729829999999998e-001; 9.1836300000000004e-001; 9.4422960000000000e-001; 9.6482740000000000e-001; 9.8024560000000005e-001; 9.9076560000000002e-001; 9.9691600000000002e-001; 9.9954909999999997e-001];
    w = [5.6377600000000014e-002; 5.5978400000000011e-002; 5.4789200000000017e-002; 5.2834900000000011e-002; 5.0157100000000017e-002; 4.6813600000000004e-002; 4.2878000000000006e-002; 3.8439800000000010e-002; 3.3603900000000006e-002; 2.8489800000000006e-002; 2.3231400000000003e-002; 1.7978600000000004e-002; 1.2903800000000003e-002; 8.2230000000000011e-003; 4.2173000000000011e-003; 1.2724000000000001e-003];
  case 15
    n = [5.0000000000000000e-001; 5.5624450000000003e-001; 6.1169330000000000e-001; 6.6556769999999998e-001; 7.1712189999999998e-001; 7.6565989999999995e-001; 8.1055149999999998e-001; 8.5124809999999995e-001; 8.8729829999999998e-001; 9.1836300000000004e-001; 9.4422960000000000e-001; 9.6482740000000000e-001; 9.8024560000000005e-001; 9.9076560000000002e-001; 9.9691600000000002e-001; 9.9954909999999997e-001];
    w = [5.6377600000000014e-002; 5.5978400000000011e-002; 5.4789200000000017e-002; 5.2834900000000011e-002; 5.0157100000000017e-002; 4.6813600000000004e-002; 4.2878000000000006e-002; 3.8439800000000010e-002; 3.3603900000000006e-002; 2.8489800000000006e-002; 2.3231400000000003e-002; 1.7978600000000004e-002; 1.2903800000000003e-002; 8.2230000000000011e-003; 4.2173000000000011e-003; 1.2724000000000001e-003];
  case 16
    n = [5.0000000000000000e-001; 5.5624450000000003e-001; 6.1169330000000000e-001; 6.6556769999999998e-001; 7.1712189999999998e-001; 7.6565989999999995e-001; 8.1055149999999998e-001; 8.5124809999999995e-001; 8.8729829999999998e-001; 9.1836300000000004e-001; 9.4422960000000000e-001; 9.6482740000000000e-001; 9.8024560000000005e-001; 9.9076560000000002e-001; 9.9691600000000002e-001; 9.9954909999999997e-001];
    w = [5.6377600000000014e-002; 5.5978400000000011e-002; 5.4789200000000017e-002; 5.2834900000000011e-002; 5.0157100000000017e-002; 4.6813600000000004e-002; 4.2878000000000006e-002; 3.8439800000000010e-002; 3.3603900000000006e-002; 2.8489800000000006e-002; 2.3231400000000003e-002; 1.7978600000000004e-002; 1.2903800000000003e-002; 8.2230000000000011e-003; 4.2173000000000011e-003; 1.2724000000000001e-003];
  case 17
    n = [5.0000000000000000e-001; 5.5624450000000003e-001; 6.1169330000000000e-001; 6.6556769999999998e-001; 7.1712189999999998e-001; 7.6565989999999995e-001; 8.1055149999999998e-001; 8.5124809999999995e-001; 8.8729829999999998e-001; 9.1836300000000004e-001; 9.4422960000000000e-001; 9.6482740000000000e-001; 9.8024560000000005e-001; 9.9076560000000002e-001; 9.9691600000000002e-001; 9.9954909999999997e-001];
    w = [5.6377600000000014e-002; 5.5978400000000011e-002; 5.4789200000000017e-002; 5.2834900000000011e-002; 5.0157100000000017e-002; 4.6813600000000004e-002; 4.2878000000000006e-002; 3.8439800000000010e-002; 3.3603900000000006e-002; 2.8489800000000006e-002; 2.3231400000000003e-002; 1.7978600000000004e-002; 1.2903800000000003e-002; 8.2230000000000011e-003; 4.2173000000000011e-003; 1.2724000000000001e-003];
  case 18
    n = [5.0000000000000000e-001; 5.5624450000000003e-001; 6.1169330000000000e-001; 6.6556769999999998e-001; 7.1712189999999998e-001; 7.6565989999999995e-001; 8.1055149999999998e-001; 8.5124809999999995e-001; 8.8729829999999998e-001; 9.1836300000000004e-001; 9.4422960000000000e-001; 9.6482740000000000e-001; 9.8024560000000005e-001; 9.9076560000000002e-001; 9.9691600000000002e-001; 9.9954909999999997e-001];
    w = [5.6377600000000014e-002; 5.5978400000000011e-002; 5.4789200000000017e-002; 5.2834900000000011e-002; 5.0157100000000017e-002; 4.6813600000000004e-002; 4.2878000000000006e-002; 3.8439800000000010e-002; 3.3603900000000006e-002; 2.8489800000000006e-002; 2.3231400000000003e-002; 1.7978600000000004e-002; 1.2903800000000003e-002; 8.2230000000000011e-003; 4.2173000000000011e-003; 1.2724000000000001e-003];
  case 19
    n = [5.0000000000000000e-001; 5.5624450000000003e-001; 6.1169330000000000e-001; 6.6556769999999998e-001; 7.1712189999999998e-001; 7.6565989999999995e-001; 8.1055149999999998e-001; 8.5124809999999995e-001; 8.8729829999999998e-001; 9.1836300000000004e-001; 9.4422960000000000e-001; 9.6482740000000000e-001; 9.8024560000000005e-001; 9.9076560000000002e-001; 9.9691600000000002e-001; 9.9954909999999997e-001];
    w = [5.6377600000000014e-002; 5.5978400000000011e-002; 5.4789200000000017e-002; 5.2834900000000011e-002; 5.0157100000000017e-002; 4.6813600000000004e-002; 4.2878000000000006e-002; 3.8439800000000010e-002; 3.3603900000000006e-002; 2.8489800000000006e-002; 2.3231400000000003e-002; 1.7978600000000004e-002; 1.2903800000000003e-002; 8.2230000000000011e-003; 4.2173000000000011e-003; 1.2724000000000001e-003];
  case 20
    n = [5.0000000000000000e-001; 5.5624450000000003e-001; 6.1169330000000000e-001; 6.6556769999999998e-001; 7.1712189999999998e-001; 7.6565989999999995e-001; 8.1055149999999998e-001; 8.5124809999999995e-001; 8.8729829999999998e-001; 9.1836300000000004e-001; 9.4422960000000000e-001; 9.6482740000000000e-001; 9.8024560000000005e-001; 9.9076560000000002e-001; 9.9691600000000002e-001; 9.9954909999999997e-001];
    w = [5.6377600000000014e-002; 5.5978400000000011e-002; 5.4789200000000017e-002; 5.2834900000000011e-002; 5.0157100000000017e-002; 4.6813600000000004e-002; 4.2878000000000006e-002; 3.8439800000000010e-002; 3.3603900000000006e-002; 2.8489800000000006e-002; 2.3231400000000003e-002; 1.7978600000000004e-002; 1.2903800000000003e-002; 8.2230000000000011e-003; 4.2173000000000011e-003; 1.2724000000000001e-003];
  case 21
    n = [5.0000000000000000e-001; 5.5624450000000003e-001; 6.1169330000000000e-001; 6.6556769999999998e-001; 7.1712189999999998e-001; 7.6565989999999995e-001; 8.1055149999999998e-001; 8.5124809999999995e-001; 8.8729829999999998e-001; 9.1836300000000004e-001; 9.4422960000000000e-001; 9.6482740000000000e-001; 9.8024560000000005e-001; 9.9076560000000002e-001; 9.9691600000000002e-001; 9.9954909999999997e-001];
    w = [5.6377600000000014e-002; 5.5978400000000011e-002; 5.4789200000000017e-002; 5.2834900000000011e-002; 5.0157100000000017e-002; 4.6813600000000004e-002; 4.2878000000000006e-002; 3.8439800000000010e-002; 3.3603900000000006e-002; 2.8489800000000006e-002; 2.3231400000000003e-002; 1.7978600000000004e-002; 1.2903800000000003e-002; 8.2230000000000011e-003; 4.2173000000000011e-003; 1.2724000000000001e-003];
  case 22
    n = [5.0000000000000000e-001; 5.5624450000000003e-001; 6.1169330000000000e-001; 6.6556769999999998e-001; 7.1712189999999998e-001; 7.6565989999999995e-001; 8.1055149999999998e-001; 8.5124809999999995e-001; 8.8729829999999998e-001; 9.1836300000000004e-001; 9.4422960000000000e-001; 9.6482740000000000e-001; 9.8024560000000005e-001; 9.9076560000000002e-001; 9.9691600000000002e-001; 9.9954909999999997e-001];
    w = [5.6377600000000014e-002; 5.5978400000000011e-002; 5.4789200000000017e-002; 5.2834900000000011e-002; 5.0157100000000017e-002; 4.6813600000000004e-002; 4.2878000000000006e-002; 3.8439800000000010e-002; 3.3603900000000006e-002; 2.8489800000000006e-002; 2.3231400000000003e-002; 1.7978600000000004e-002; 1.2903800000000003e-002; 8.2230000000000011e-003; 4.2173000000000011e-003; 1.2724000000000001e-003];
  case 23
    n = [5.0000000000000000e-001; 5.5624450000000003e-001; 6.1169330000000000e-001; 6.6556769999999998e-001; 7.1712189999999998e-001; 7.6565989999999995e-001; 8.1055149999999998e-001; 8.5124809999999995e-001; 8.8729829999999998e-001; 9.1836300000000004e-001; 9.4422960000000000e-001; 9.6482740000000000e-001; 9.8024560000000005e-001; 9.9076560000000002e-001; 9.9691600000000002e-001; 9.9954909999999997e-001];
    w = [5.6377600000000014e-002; 5.5978400000000011e-002; 5.4789200000000017e-002; 5.2834900000000011e-002; 5.0157100000000017e-002; 4.6813600000000004e-002; 4.2878000000000006e-002; 3.8439800000000010e-002; 3.3603900000000006e-002; 2.8489800000000006e-002; 2.3231400000000003e-002; 1.7978600000000004e-002; 1.2903800000000003e-002; 8.2230000000000011e-003; 4.2173000000000011e-003; 1.2724000000000001e-003];
  case 24
    n = [5.0000000000000000e-001; 5.5624450000000003e-001; 6.1169330000000000e-001; 6.6556769999999998e-001; 7.1712189999999998e-001; 7.6565989999999995e-001; 8.1055149999999998e-001; 8.5124809999999995e-001; 8.8729829999999998e-001; 9.1836300000000004e-001; 9.4422960000000000e-001; 9.6482740000000000e-001; 9.8024560000000005e-001; 9.9076560000000002e-001; 9.9691600000000002e-001; 9.9954909999999997e-001];
    w = [5.6377600000000014e-002; 5.5978400000000011e-002; 5.4789200000000017e-002; 5.2834900000000011e-002; 5.0157100000000017e-002; 4.6813600000000004e-002; 4.2878000000000006e-002; 3.8439800000000010e-002; 3.3603900000000006e-002; 2.8489800000000006e-002; 2.3231400000000003e-002; 1.7978600000000004e-002; 1.2903800000000003e-002; 8.2230000000000011e-003; 4.2173000000000011e-003; 1.2724000000000001e-003];
  case 25
    n = [5.0000000000000000e-001; 5.2817210000000003e-001; 5.5624450000000003e-001; 5.8411769999999996e-001; 6.1169330000000000e-001; 6.3887490000000002e-001; 6.6556769999999998e-001; 6.9167970000000001e-001; 7.1712189999999998e-001; 7.4180900000000005e-001; 7.6565989999999995e-001; 7.8859789999999996e-001; 8.1055149999999998e-001; 8.3145480000000005e-001; 8.5124809999999995e-001; 8.6987800000000004e-001; 8.8729829999999998e-001; 9.0347029999999995e-001; 9.1836300000000004e-001; 9.3195399999999995e-001; 9.4422960000000000e-001; 9.5518559999999997e-001; 9.6482740000000000e-001; 9.7317140000000002e-001; 9.8024560000000005e-001; 9.8609139999999995e-001; 9.9076560000000002e-001; 9.9434239999999996e-001; 9.9691600000000002e-001; 9.9860309999999997e-001; 9.9954909999999997e-001; 9.9993650000000001e-001];
    w = [2.8188799999999993e-002; 2.8138799999999992e-002; 2.7989199999999992e-002; 2.7740699999999993e-002; 2.7394599999999995e-002; 2.6952699999999993e-002; 2.6417499999999993e-002; 2.5791599999999994e-002; 2.5078599999999993e-002; 2.4282199999999993e-002; 2.3406799999999995e-002; 2.2457299999999996e-002; 2.1438999999999996e-002; 2.0357799999999995e-002; 1.9219899999999998e-002; 1.8032199999999998e-002; 1.6801899999999998e-002; 1.5536799999999996e-002; 1.4244899999999996e-002; 1.2934799999999996e-002; 1.1615699999999998e-002; 1.0297099999999998e-002; 8.9892999999999987e-003; 7.7033999999999983e-003; 6.4518999999999983e-003; 5.2490999999999987e-003; 4.1114999999999988e-003; 3.0577999999999990e-003; 2.1087999999999997e-003; 1.2894999999999998e-003; 6.3259999999999987e-004; 1.8159999999999997e-004];
  end
end
function [n w] = KPN(l)
  switch l
  case 1
    n = [0.0000000000000000e+000];
    w = [1.0000000000000000e+000];
  case 2
    n = [0.0000000000000000e+000; 1.7320508075688772e+000];
    w = [6.6666666666666663e-001; 1.6666666666666666e-001];
  case 3
    n = [0.0000000000000000e+000; 1.7320508075688772e+000];
    w = [6.6666666666666674e-001; 1.6666666666666666e-001];
  case 4
    n = [0.0000000000000000e+000; 7.4109534999454085e-001; 1.7320508075688772e+000; 4.1849560176727323e+000];
    w = [4.5874486825749189e-001; 1.3137860698313561e-001; 1.3855327472974924e-001; 6.9568415836913987e-004];
  case 5
    n = [0.0000000000000000e+000; 7.4109534999454085e-001; 1.7320508075688772e+000; 2.8612795760570582e+000; 4.1849560176727323e+000];
    w = [2.5396825396825407e-001; 2.7007432957793776e-001; 9.4850948509485125e-002; 7.9963254708935293e-003; 9.4269457556517470e-005];
  case 6
    n = [0.0000000000000000e+000; 7.4109534999454085e-001; 1.7320508075688772e+000; 2.8612795760570582e+000; 4.1849560176727323e+000];
    w = [2.5396825396825429e-001; 2.7007432957793776e-001; 9.4850948509485070e-002; 7.9963254708935293e-003; 9.4269457556517551e-005];
  case 7
    n = [0.0000000000000000e+000; 7.4109534999454085e-001; 1.7320508075688772e+000; 2.8612795760570582e+000; 4.1849560176727323e+000];
    w = [2.5396825396825418e-001; 2.7007432957793781e-001; 9.4850948509485014e-002; 7.9963254708935311e-003; 9.4269457556517592e-005];
  case 8
    n = [0.0000000000000000e+000; 7.4109534999454085e-001; 1.7320508075688772e+000; 2.8612795760570582e+000; 4.1849560176727323e+000];
    w = [2.5396825396825418e-001; 2.7007432957793781e-001; 9.4850948509485042e-002; 7.9963254708935276e-003; 9.4269457556517375e-005];
  case 9
    n = [0.0000000000000000e+000; 7.4109534999454085e-001; 1.2304236340273060e+000; 1.7320508075688772e+000; 2.5960831150492023e+000; 2.8612795760570582e+000; 4.1849560176727323e+000; 5.1870160399136562e+000; 6.3633944943363696e+000];
    w = [2.6692223033505302e-001; 2.5456123204171222e-001; 1.4192654826449365e-002; 8.8681002152028010e-002; 1.9656770938777492e-003; 7.0334802378279075e-003; 1.0563783615416941e-004; -8.2049207541509217e-007; 2.1136499505424257e-008];
  case 10
    n = [0.0000000000000000e+000; 7.4109534999454085e-001; 1.2304236340273060e+000; 1.7320508075688772e+000; 2.5960831150492023e+000; 2.8612795760570582e+000; 3.2053337944991944e+000; 4.1849560176727323e+000; 5.1870160399136562e+000; 6.3633944943363696e+000];
    w = [3.0346719985420623e-001; 2.0832499164960877e-001; 6.1151730125247716e-002; 6.4096054686807610e-002; 1.8085234254798462e-002; -6.3372247933737571e-003; 2.8848804365067559e-003; 6.0123369459847997e-005; 6.0948087314689840e-007; 8.6296846022298632e-010];
  case 11
    n = [0.0000000000000000e+000; 7.4109534999454085e-001; 1.2304236340273060e+000; 1.7320508075688772e+000; 2.5960831150492023e+000; 2.8612795760570582e+000; 3.2053337944991944e+000; 4.1849560176727323e+000; 5.1870160399136562e+000; 6.3633944943363696e+000];
    w = [3.0346719985420623e-001; 2.0832499164960872e-001; 6.1151730125247709e-002; 6.4096054686807541e-002; 1.8085234254798459e-002; -6.3372247933737545e-003; 2.8848804365067555e-003; 6.0123369459847922e-005; 6.0948087314689830e-007; 8.6296846022298839e-010];
  case 12
    n = [0.0000000000000000e+000; 7.4109534999454085e-001; 1.2304236340273060e+000; 1.7320508075688772e+000; 2.5960831150492023e+000; 2.8612795760570582e+000; 3.2053337944991944e+000; 4.1849560176727323e+000; 5.1870160399136562e+000; 6.3633944943363696e+000];
    w = [3.0346719985420623e-001; 2.0832499164960872e-001; 6.1151730125247716e-002; 6.4096054686807624e-002; 1.8085234254798466e-002; -6.3372247933737545e-003; 2.8848804365067559e-003; 6.0123369459847841e-005; 6.0948087314689830e-007; 8.6296846022298963e-010];
  case 13
    n = [0.0000000000000000e+000; 7.4109534999454085e-001; 1.2304236340273060e+000; 1.7320508075688772e+000; 2.5960831150492023e+000; 2.8612795760570582e+000; 3.2053337944991944e+000; 4.1849560176727323e+000; 5.1870160399136562e+000; 6.3633944943363696e+000];
    w = [3.0346719985420600e-001; 2.0832499164960883e-001; 6.1151730125247730e-002; 6.4096054686807638e-002; 1.8085234254798459e-002; -6.3372247933737580e-003; 2.8848804365067555e-003; 6.0123369459847868e-005; 6.0948087314689830e-007; 8.6296846022298756e-010];
  case 14
    n = [0.0000000000000000e+000; 7.4109534999454085e-001; 1.2304236340273060e+000; 1.7320508075688772e+000; 2.5960831150492023e+000; 2.8612795760570582e+000; 3.2053337944991944e+000; 4.1849560176727323e+000; 5.1870160399136562e+000; 6.3633944943363696e+000];
    w = [3.0346719985420617e-001; 2.0832499164960874e-001; 6.1151730125247702e-002; 6.4096054686807596e-002; 1.8085234254798459e-002; -6.3372247933737563e-003; 2.8848804365067555e-003; 6.0123369459847936e-005; 6.0948087314689851e-007; 8.6296846022298322e-010];
  case 15
    n = [0.0000000000000000e+000; 7.4109534999454085e-001; 1.2304236340273060e+000; 1.7320508075688772e+000; 2.5960831150492023e+000; 2.8612795760570582e+000; 3.2053337944991944e+000; 4.1849560176727323e+000; 5.1870160399136562e+000; 6.3633944943363696e+000];
    w = [3.0346719985420612e-001; 2.0832499164960874e-001; 6.1151730125247723e-002; 6.4096054686807652e-002; 1.8085234254798459e-002; -6.3372247933737597e-003; 2.8848804365067563e-003; 6.0123369459848091e-005; 6.0948087314689851e-007; 8.6296846022298983e-010];
  case 16
    n = [0.0000000000000000e+000; 2.4899229757996061e-001; 7.4109534999454085e-001; 1.2304236340273060e+000; 1.7320508075688772e+000; 2.2336260616769419e+000; 2.5960831150492023e+000; 2.8612795760570582e+000; 3.2053337944991944e+000; 3.6353185190372783e+000; 4.1849560176727323e+000; 5.1870160399136562e+000; 6.3633944943363696e+000; 7.1221067008046166e+000; 7.9807717985905606e+000; 9.0169397898903032e+000];
    w = [2.5890005324151566e-001; 2.8128101540033167e-002; 1.9968863511734550e-001; 6.5417392836092561e-002; 6.1718532565867179e-002; 1.7608475581318002e-003; 1.6592492698936010e-002; -5.5610063068358157e-003; 2.7298430467334002e-003; 1.5044205390914219e-005; 5.9474961163931621e-005; 6.1435843232617913e-007; 7.9298267864869338e-010; 5.1158053105504208e-012; -1.4840835740298868e-013; 1.2618464280815118e-015];
  case 17
    n = [0.0000000000000000e+000; 2.4899229757996061e-001; 7.4109534999454085e-001; 1.2304236340273060e+000; 1.7320508075688772e+000; 2.2336260616769419e+000; 2.5960831150492023e+000; 2.8612795760570582e+000; 3.2053337944991944e+000; 3.6353185190372783e+000; 4.1849560176727323e+000; 5.1870160399136562e+000; 5.6981777684881099e+000; 6.3633944943363696e+000; 7.1221067008046166e+000; 7.9807717985905606e+000; 9.0169397898903032e+000];
    w = [1.3911022236338039e-001; 1.0387687125574284e-001; 1.7607598741571459e-001; 7.7443602746299481e-002; 5.4677556143463042e-002; 7.3530110204955076e-003; 1.1529247065398790e-002; -2.7712189007789243e-003; 2.1202259559596325e-003; 8.3236045295766745e-005; 5.5691158981081479e-005; 6.9086261179113738e-007; -1.3486017348542930e-008; 1.5542195992782658e-009; -1.9341305000880955e-011; 2.6640625166231651e-013; -9.9313913286822465e-016];
  case 18
    n = [0.0000000000000000e+000; 2.4899229757996061e-001; 7.4109534999454085e-001; 1.2304236340273060e+000; 1.7320508075688772e+000; 2.2336260616769419e+000; 2.5960831150492023e+000; 2.8612795760570582e+000; 3.2053337944991944e+000; 3.6353185190372783e+000; 4.1849560176727323e+000; 4.7364330859522967e+000; 5.1870160399136562e+000; 5.6981777684881099e+000; 6.3633944943363696e+000; 7.1221067008046166e+000; 7.9807717985905606e+000; 9.0169397898903032e+000];
    w = [5.1489450806921377e-004; 1.9176011588804434e-001; 1.4807083115521585e-001; 9.2364726716986353e-002; 4.5273685465150391e-002; 1.5673473751851151e-002; 3.1554462691875513e-003; 2.3113452403522071e-003; 8.1895392750226735e-004; 2.7524214116785131e-004; 3.5729348198975332e-005; 2.7342206801187888e-006; 2.4676421345798140e-007; 2.1394194479561062e-008; 4.6011760348655917e-010; 3.0972223576062995e-012; 5.4500412650638128e-015; 1.0541326582334014e-018];
  case 19
    n = [0.0000000000000000e+000; 2.4899229757996061e-001; 7.4109534999454085e-001; 1.2304236340273060e+000; 1.7320508075688772e+000; 2.2336260616769419e+000; 2.5960831150492023e+000; 2.8612795760570582e+000; 3.2053337944991944e+000; 3.6353185190372783e+000; 4.1849560176727323e+000; 4.7364330859522967e+000; 5.1870160399136562e+000; 5.6981777684881099e+000; 6.3633944943363696e+000; 7.1221067008046166e+000; 7.9807717985905606e+000; 9.0169397898903032e+000];
    w = [5.1489450806921377e-004; 1.9176011588804437e-001; 1.4807083115521585e-001; 9.2364726716986353e-002; 4.5273685465150523e-002; 1.5673473751851151e-002; 3.1554462691875604e-003; 2.3113452403522050e-003; 8.1895392750226670e-004; 2.7524214116785131e-004; 3.5729348198975447e-005; 2.7342206801187884e-006; 2.4676421345798140e-007; 2.1394194479561056e-008; 4.6011760348656077e-010; 3.0972223576063011e-012; 5.4500412650637663e-015; 1.0541326582337958e-018];
  case 20
    n = [0.0000000000000000e+000; 2.4899229757996061e-001; 7.4109534999454085e-001; 1.2304236340273060e+000; 1.7320508075688772e+000; 2.2336260616769419e+000; 2.5960831150492023e+000; 2.8612795760570582e+000; 3.2053337944991944e+000; 3.6353185190372783e+000; 4.1849560176727323e+000; 4.7364330859522967e+000; 5.1870160399136562e+000; 5.6981777684881099e+000; 6.3633944943363696e+000; 7.1221067008046166e+000; 7.9807717985905606e+000; 9.0169397898903032e+000];
    w = [5.1489450806925551e-004; 1.9176011588804440e-001; 1.4807083115521585e-001; 9.2364726716986298e-002; 4.5273685465150537e-002; 1.5673473751851155e-002; 3.1554462691875573e-003; 2.3113452403522080e-003; 8.1895392750226724e-004; 2.7524214116785137e-004; 3.5729348198975352e-005; 2.7342206801187888e-006; 2.4676421345798124e-007; 2.1394194479561056e-008; 4.6011760348656144e-010; 3.0972223576062963e-012; 5.4500412650638365e-015; 1.0541326582335402e-018];
  case 21
    n = [0.0000000000000000e+000; 2.4899229757996061e-001; 7.4109534999454085e-001; 1.2304236340273060e+000; 1.7320508075688772e+000; 2.2336260616769419e+000; 2.5960831150492023e+000; 2.8612795760570582e+000; 3.2053337944991944e+000; 3.6353185190372783e+000; 4.1849560176727323e+000; 4.7364330859522967e+000; 5.1870160399136562e+000; 5.6981777684881099e+000; 6.3633944943363696e+000; 7.1221067008046166e+000; 7.9807717985905606e+000; 9.0169397898903032e+000];
    w = [5.1489450806913744e-004; 1.9176011588804429e-001; 1.4807083115521594e-001; 9.2364726716986312e-002; 4.5273685465150391e-002; 1.5673473751851151e-002; 3.1554462691875565e-003; 2.3113452403522089e-003; 8.1895392750226670e-004; 2.7524214116785142e-004; 3.5729348198975285e-005; 2.7342206801187888e-006; 2.4676421345798119e-007; 2.1394194479561059e-008; 4.6011760348656594e-010; 3.0972223576062950e-012; 5.4500412650638696e-015; 1.0541326582332041e-018];
  case 22
    n = [0.0000000000000000e+000; 2.4899229757996061e-001; 7.4109534999454085e-001; 1.2304236340273060e+000; 1.7320508075688772e+000; 2.2336260616769419e+000; 2.5960831150492023e+000; 2.8612795760570582e+000; 3.2053337944991944e+000; 3.6353185190372783e+000; 4.1849560176727323e+000; 4.7364330859522967e+000; 5.1870160399136562e+000; 5.6981777684881099e+000; 6.3633944943363696e+000; 7.1221067008046166e+000; 7.9807717985905606e+000; 9.0169397898903032e+000];
    w = [5.1489450806903368e-004; 1.9176011588804448e-001; 1.4807083115521574e-001; 9.2364726716986423e-002; 4.5273685465150516e-002; 1.5673473751851161e-002; 3.1554462691875543e-003; 2.3113452403522063e-003; 8.1895392750226713e-004; 2.7524214116785164e-004; 3.5729348198975319e-005; 2.7342206801187905e-006; 2.4676421345798151e-007; 2.1394194479561082e-008; 4.6011760348656005e-010; 3.0972223576063043e-012; 5.4500412650637592e-015; 1.0541326582339926e-018];
  case 23
    n = [0.0000000000000000e+000; 2.4899229757996061e-001; 7.4109534999454085e-001; 1.2304236340273060e+000; 1.7320508075688772e+000; 2.2336260616769419e+000; 2.5960831150492023e+000; 2.8612795760570582e+000; 3.2053337944991944e+000; 3.6353185190372783e+000; 4.1849560176727323e+000; 4.7364330859522967e+000; 5.1870160399136562e+000; 5.6981777684881099e+000; 6.3633944943363696e+000; 7.1221067008046166e+000; 7.9807717985905606e+000; 9.0169397898903032e+000];
    w = [5.1489450806913755e-004; 1.9176011588804442e-001; 1.4807083115521577e-001; 9.2364726716986381e-002; 4.5273685465150468e-002; 1.5673473751851155e-002; 3.1554462691875560e-003; 2.3113452403522045e-003; 8.1895392750226572e-004; 2.7524214116785158e-004; 3.5729348198975298e-005; 2.7342206801187892e-006; 2.4676421345798129e-007; 2.1394194479561072e-008; 4.6011760348656103e-010; 3.0972223576062963e-012; 5.4500412650638207e-015; 1.0541326582338368e-018];
  case 24
    n = [0.0000000000000000e+000; 2.4899229757996061e-001; 7.4109534999454085e-001; 1.2304236340273060e+000; 1.7320508075688772e+000; 2.2336260616769419e+000; 2.5960831150492023e+000; 2.8612795760570582e+000; 3.2053337944991944e+000; 3.6353185190372783e+000; 4.1849560176727323e+000; 4.7364330859522967e+000; 5.1870160399136562e+000; 5.6981777684881099e+000; 6.3633944943363696e+000; 7.1221067008046166e+000; 7.9807717985905606e+000; 9.0169397898903032e+000];
    w = [5.1489450806914438e-004; 1.9176011588804442e-001; 1.4807083115521577e-001; 9.2364726716986340e-002; 4.5273685465150509e-002; 1.5673473751851155e-002; 3.1554462691875586e-003; 2.3113452403522058e-003; 8.1895392750226551e-004; 2.7524214116785142e-004; 3.5729348198975386e-005; 2.7342206801187884e-006; 2.4676421345798082e-007; 2.1394194479561059e-008; 4.6011760348656382e-010; 3.0972223576062942e-012; 5.4500412650638381e-015; 1.0541326582336941e-018];
  case 25
    n = [0.0000000000000000e+000; 2.4899229757996061e-001; 7.4109534999454085e-001; 1.2304236340273060e+000; 1.7320508075688772e+000; 2.2336260616769419e+000; 2.5960831150492023e+000; 2.8612795760570582e+000; 3.2053337944991944e+000; 3.6353185190372783e+000; 4.1849560176727323e+000; 4.7364330859522967e+000; 5.1870160399136562e+000; 5.6981777684881099e+000; 6.3633944943363696e+000; 7.1221067008046166e+000; 7.9807717985905606e+000; 9.0169397898903032e+000];
    w = [5.1489450806919989e-004; 1.9176011588804437e-001; 1.4807083115521580e-001; 9.2364726716986395e-002; 4.5273685465150426e-002; 1.5673473751851158e-002; 3.1554462691875539e-003; 2.3113452403522054e-003; 8.1895392750226681e-004; 2.7524214116785142e-004; 3.5729348198975292e-005; 2.7342206801187884e-006; 2.4676421345798108e-007; 2.1394194479561056e-008; 4.6011760348655901e-010; 3.0972223576062975e-012; 5.4500412650638412e-015; 1.0541326582337527e-018];
  end
end
```
