## Supplementary material for "An Alternative Framework for Fluorescence Correlation Spectroscopy": Source Code: Sample_emission.m

```
% Metropolis-Hastings algorithm to sample molecule and background photon emission rates. 
% Written by Sina Jazani

function[ MU, MU_BACK , mu_acceptance_rate ] = Sample_emission( ...
...
...
mu_proposal        , mu_back_proposal , mu_alpha  , mu_beta      , ...
mu_back_alpha      , mu_back_beta     , MU_OLD    , MU_BACK_OLD  , ...
mu_acceptance_rate , trace            , Time      , b            , ...
constant           , Number_molecules , Max_Species)  


%%%%%%%%%%%%%%%%%%%%%%%%%%%%%%%%%%%%%%%%%%%%%%%%%%%%%%%%%%%%%%%%%%%%%%%%%%%
%%%%%%%%%%%%%%%%%%%%%%%%%%%     INPUT      %%%%%%%%%%%%%%%%%%%%%%%%%%%%%%%%
%%%%%%%%%%%%%%%%%%%%%%%%%%%%%%%%%%%%%%%%%%%%%%%%%%%%%%%%%%%%%%%%%%%%%%%%%%%
     % mu_proposal                   :  Proposal parameter for MU           ( Gamma distribution)
     % mu_back_proposal              :  Proposal parameter for MU_BACK      ( Gamma distribution)
     % mu_alpha                      :  Alpha parameter of MU prior         ( Gamma distribution)
     % mu_beta                       :  Beta parameter of MU prior          ( Gamma distribution)
     % mu_back_alpha                 :  Alpha parameter of MU_BACK prior    ( Gamma distribution)
     % mu_back_beta                  :  Alpha parameter of MU_BACK prior    ( Gamma distribution)
     % x                             :  Positions of the particles (Normalized)
     % MU_OLD                        :  Previously sampled molecule photon emission rate 
     % MU_BACK_OLD                   :  Previously sampled background photon emission rate
     % sign_siz                      :  Length of the observation trace
     % Number_molecules              :  Number of defined molecules for each species
     % mu_acceptance_rate            :  Previous acceptance rate 
     % trace                         :  Observed number of photons per bin (trace)
     % Time                          :  Data acquisition time
     % b                             :  Loads for active particles
     % Max_species                   :  Total number of species
     % PSF_func                      :  Point Spread Function type ( 'Gaussian' or 'Lorentzian Gaussian')
     
%%%%%%%%%%%%%%%%%%%%%%%%%%%%%%%%%%%%%%%%%%%%%%%%%%%%%%%%%%%%%%%%%%%%%%%%%%%
%%%%%%%%%%%%%%%%%%%%%%%%%%%     OUTPUT     %%%%%%%%%%%%%%%%%%%%%%%%%%%%%%%%
%%%%%%%%%%%%%%%%%%%%%%%%%%%%%%%%%%%%%%%%%%%%%%%%%%%%%%%%%%%%%%%%%%%%%%%%%%%
     % MU                            :  Sampled molcule photon emission rate 
     % MU_BACK                       :  Sampled background photon emission rate
     % mu_acceptance_rate            :  Updated acceptance rate


     
% Sample proposals for mu and mu_back from gamma distributions
     MU_NEW       =  (MU_OLD/mu_proposal)           .* randg(mu_proposal,Max_Species,1)      ;
     MU_BACK_NEW  =  (MU_BACK_OLD/mu_back_proposal)  * randg(mu_back_proposal) ;

     Tnew=sqrt(4*(MU_BACK_NEW + (repelem(MU_NEW,Number_molecules,1).*b)'*constant ).*Time+1.5) ...
         -0.25./sqrt((MU_BACK_NEW+ (repelem(MU_NEW,Number_molecules,1).*b)'*constant).*Time);   
     
     Told=sqrt(4*(MU_BACK_OLD + (repelem(MU_OLD,Number_molecules,1).*b)'*constant).*Time+1.5) ...
         -0.25./sqrt((MU_BACK_OLD+ (repelem(MU_OLD,Number_molecules,1).*b)'*constant).*Time);
                      
     constant1 = sum( (Told.^2 - Tnew.^2)./2 + trace.*(Tnew-Told) );
              

% Calculate the logarithm of the new posterior over old posterior and proposal ratio                             
     logr = constant1 + sum(((2*mu_proposal-mu_alpha)*log(MU_OLD./MU_NEW))+((MU_OLD-MU_NEW)/mu_beta)+...
                (mu_proposal*((MU_NEW./MU_OLD)-(MU_OLD./MU_NEW))))+...
                ...
                ((2*mu_back_proposal-mu_back_alpha)*log(MU_BACK_OLD/MU_BACK_NEW))+...   
                ((MU_BACK_OLD-MU_BACK_NEW)/mu_back_beta)+...
                (mu_back_proposal*((MU_BACK_NEW/MU_BACK_OLD)-(MU_BACK_OLD/MU_BACK_NEW)));

% Accept or reject the proposals
         if  logr>(-exprnd(1))
             MU = MU_NEW;
             MU_BACK=MU_BACK_NEW;
             mu_acceptance_rate=mu_acceptance_rate+1;
         else
             MU = MU_OLD;
             MU_BACK=MU_BACK_OLD;
         end
end
```
