## Supplementary material for "An Alternative Framework for Fluorescence Correlation Spectroscopy": Source Code: Sample_Generator.m

```
function [ X , Y , Z , trace ] = ...
    ...
    ...
Sample_Generator( Lxy                , Lz                   , Length_signal , Number_molecules , ...
                  wxy                , wz                   , D             , Delta_t          , ...
                  Molecule_emission  , Background_emission  , PSF_func )
 
%%%%%%%%%%%%%%%%%%%%%%%%%%%%%%%%%%%%%%%%%%%%%%%%%%%%%%%%%%%%%%%%%%%%%%%%%%%
%%%%%%%%%%%%%%%%%%%%%%%%%%%%%% INPUT %%%%%%%%%%%%%%%%%%%%%%%%%%%%%%%%%%%%%%
%%%%%%%%%%%%%%%%%%%%%%%%%%%%%%%%%%%%%%%%%%%%%%%%%%%%%%%%%%%%%%%%%%%%%%%%%%%
     % Lxy                   :  Periodic boundary in  x and y coordinates
     % Lz                    :  Periodic boundary in  z coordinate
     % Length_signal         :  Length of the trace
     % Number_molecules      :  Number of molecules for each species
     % wxy                   :  Minor Semi-axis of the confocal volume
     % wz                    :  Major Semi-axis of the confocal volume
     % D                     :  Diffusion coeffcient(s)
     % Delta_t               :  Data acquisition time
     % Molecule_emission     :  Molecule photon emission rates
     % Background_emission   :  Backgrouund photon emission rates
     % PSF_func              :  Point spread function type (3DG = 1 and 2DG-L =2 )

           
%%%%%%%%%%%%%%%%%%%%%%%%%%%%%%%%%%%%%%%%%%%%%%%%%%%%%%%%%%%%%%%%%%%%%%%%%%%
%%%%%%%%%%%%%%%%%%%%%%%%%%%%%% OUTPUT %%%%%%%%%%%%%%%%%%%%%%%%%%%%%%%%%%%%%
%%%%%%%%%%%%%%%%%%%%%%%%%%%%%%%%%%%%%%%%%%%%%%%%%%%%%%%%%%%%%%%%%%%%%%%%%%%
     % X                     :  Location molecuules in x coordinate
     % Y                     :  Location molecuules in y coordinate
     % Z                     :  Location molecuules in z coordinate
     % trace                 :  Single photon arrival time trace

     
     
     
% Total number of species based on number of defined difussion coefficients
num_species=length(D);

% Creat empty vactors for the locatoins of molecules
X = zeros ( sum(Number_molecules) , Length_signal+1 ) ;
Y = zeros ( sum(Number_molecules) , Length_signal+1 ) ;
Z = zeros ( sum(Number_molecules) , Length_signal+1 ) ;


Emission=[];


for num = 1:num_species
    kxy  = sqrt(2*D(num)*Delta_t)/wxy ;
    kz   = sqrt(2*D(num)*Delta_t)/wz ;
    
    Emission = [Emission , repmat(Molecule_emission(num),1,Number_molecules(num))];
     for n  = 1:Number_molecules(num)
         
% Initialize location of molecules
        nump=(num-1)*sum(Number_molecules(1:num-1))+n;
        
        X(nump,1) = Lxy*(1-2*rand());
        Y(nump,1) = Lxy*(1-2*rand());
        Z(nump,1) = Lz *(1-2*rand());
    
        for i=2:Length_signal+1 
    
% Brownian motion
            X(nump,i) = X(nump,i-1) + kxy*randn();
            Y(nump,i) = Y(nump,i-1) + kxy*randn();
            Z(nump,i) = Z(nump,i-1) + kz *randn();

% Periodic boundaries
            if  X(nump,i) >= Lxy
                X(nump,i)  = X(nump,i)-2*Lxy;
            end  
            if  X(nump,i) <= -Lxy
                X(nump,i)  = X(nump,i)+2*Lxy;
            end 
            if  Y(nump,i) >= Lxy
                Y(nump,i)  = Y(nump,i)-2*Lxy;
            end  
            if  Y(nump,i) <= -Lxy
                Y(nump,i)  = Y(nump,i)+2*Lxy;
            end 
            if  Z(nump,i) >= Lz
                Z(nump,i)  = Z(nump,i)-2*Lz;
            end  
            if  Z(nump,i) <= -Lz
                Z(nump,i)  = Z(nump,i)+2*Lz;
            end
            
        end
    end

end % end of time loop


    
switch PSF_func
    
   case 1         % 3D Gaussian PSF
              constant = Emission*(exp(-2*( X.^2 + Y.^2 + Z.^2 )));
    case 2        % 2D Gaussian-Lorentzian PSF
              constant = Emission*(exp(-2*(X.^2+Y.^2)./(1+Z.^2))./(1+Z.^2)) ;
    otherwise     % 2D Gaussian-Cylindrical PSF
              constant = Emission*(exp(-2*( X.^2 + Y.^2  )));
end

   trace_0_1 = binornd(ones(1,Length_signal+1) ,1-exp(-Delta_t*(Background_emission+constant)) );
   
 % Arrival photons locations in time
   photon_loc = find(trace_0_1>0)  ;
   trace      = photon_loc*Delta_t ;
   
   X          = X(:,photon_loc)    ;
   Y          = Y(:,photon_loc)    ;
   Z          = Z(:,photon_loc)    ;
   
end
```
