## Supplementary material for "An Alternative Framework for Fluorescence Correlation Spectroscopy": Source Code: Sample_loads.m

```
% Metropolis-Hastings algorithm to sample loads. 
% Written by Sina Jazani

function[ b , acceptance_rate_b ]    = Sample_loads( ...
...
...
b_old        , q                  , I     , I_back            , ...
trace        , acceptance_rate_b  , Time  , Number_molecules  , ...
Max_Species  , constant           , m     ) 

% %%%%%%%%% INPUT %%%%%%%%%
     % b_old                         :  Previously sampled load b
     % q                             :  Weights of loads
     % I                             :  Molecule photon emission rate
     % I_back                        :  Background photon emission rate
     % alpha_b                       :  Alpha parameter of load b prior         ( Gamma distribution)
     % beta_b                        :  Beta parameter of load b prior          ( Gamma distribution)
     % tace                          :  Observed number of photons per bin (trace)
     % x                             :  Positions of the molecules (Normalized)
     % acceptance_rate_b             :  Previous acceptance rate 
     % Time                          :  Data acquisition time
     % sign_siz                      :  Length of the observation trace
     % Number_molecules              :  Number of defined molecules for each species
     % Max_species                   :  Total number of species
     % PSF_func                      :  Point Spread Function type ( 'Gaussian' or 'Lorentzian Gaussian')
     
 
% %%%%%%%% OUTPUT %%%%%%%%%
     % b                             :  Samopled load b
     % acceptance_rate_b             :  Updated acceptance rate
     

   
   % Propose the new loads
   b_new = b_old;
   b     = b_old;
   
   b_new((m-1)*Number_molecules+1:m*Number_molecules,1) = ...
       rand(Number_molecules,1)<q((m-1)*Number_molecules+1:m*Number_molecules,1);
   

   Tnew = sqrt(4*(I_back + (repelem(I,Number_molecules,1).*b_new)'*constant).*Time + 1.5)...
       -0.25./sqrt((I_back + (repelem(I,Number_molecules,1).*b_new)'*constant).*Time )  ; 
   
   Told = sqrt(4*(I_back + (repelem(I,Number_molecules,1).*b_old)'*constant).*Time + 1.5)...
       -0.25./sqrt((I_back + (repelem(I,Number_molecules,1).*b_old)'*constant).*Time )   ;
         
   % Logarithm of the Posterior ratio
   logr = sum( (Told.^2 - Tnew.^2)./2 + trace.*(Tnew-Told) );


   % Check the acceptance         
   if  logr>(-exprnd(1))
       b = b_new ;   %% Accept %%
       acceptance_rate_b = acceptance_rate_b + 1 ;
   end
   
end
```
