## Supplementary material for "An Alternative Framework for Fluorescence Correlation Spectroscopy": Source Code: run_Main.m

```
%%%%%%%%%%%%%%% Multi-diffusive species analysis v1.1 %%%%%%%%%%%%%%%%%%%%%
%%%% In the case of problem, please contact 
%%%% Sina Jazani or Dr. Steve Presse.
%%%% The code written by Sina Jazani.


%%%%%%%%%%%%%%%%%%%%% Import experimental traces %%%%%%%%%%%%%%%%%%%%%%%%%% 
%%%%  In the case that the user want to import an experimental trace:
%%%%
%%%%    -  The imported trace should ba a vector of single photon arrival
%%%%       times.
%%%%    -  Start from step 3 which the user choose a segment of the
%%%%       imported trace and bin it based on initial bin size.
%%%%    -  In step 4 we apply the addaptive bin and the outputs are bbon
%%%%       sizes and the related trace.
%%%%    -  In step 5 user will provide parameter which are needed in the
%%%%       algorithm.
%%%%    -  It is necessary for the user to define wxy, wz, and PSF_func
%%%%       type before run step 6.
%%%%
%%%%          PSF_func = 1 -> 3D gaussian
%%%%          PSF_func = 2 -> 2D Gaussian-Lorentzian
%%%%          PSF_func = 2 -> 2D Gaussian-cylindrical


%%%%%%%%%%%%%%%%%%%%%%%%%%%% Clear the history %%%%%%%%%%%%%%%%%%%%%%%%%%%% Step 0
clc
clear
close all

% Add the function folder to the path
addpath('Functions')
%%%%%%%%%%%%%%%%%%%%%%%%%%%%%%%%%%%%%%%%%%%%%%%%%%%%%%%%%%%%%%%%%%%%%%%%%%% Step 1
%%%%%%%%%%%%%%%%%%%%%%%%%%%% Define parameters %%%%%%%%%%%%%%%%%%%%%%%%%%%%
%%%%%%%%%%%%%%%%%%%%%%%%%%%%%%%%%%%%%%%%%%%%%%%%%%%%%%%%%%%%%%%%%%%%%%%%%%%
Delta_t              =  10^-7      ;    % camera dead time
Diffusion            =  [5,10]     ;    % Deffusion coefficient(s) of species
Molecule_emission    = [5,10].*(10^4) ;    % Molecule photon emission rate (photons/s)
Background_emission  =    1000     ;    % Background photon emission rate (photons/s)
Lxy                  =       2     ;    % Periodic boundary in x and y axis
Lz                   =       3     ;    % Periodic boundary in z axis

PSF_func             =  1          ;    % Point spread function type ( 3D Gaussian = 1 , 2D Gaussian Lorentzian = 2 )

wxy                  =  0.3     ;   % Semi-axis of confocal in xy plane (micro meter)
wz                   =  1.5     ;   % Semi-axis of confocal in xy plane (micro meter)


Number_molecules_sim =       [2,1]   ;    % Number of molecule for each species

Length_signal        =  2000000   ;    % Number of steps
%%
%%%%%%%%%%%%%%%%%%%%%%%%%%%%%%%%%%%%%%%%%%%%%%%%%%%%%%%%%%%%%%%%%%%%%%%%%%% Step 2
%%%%%%%%%%%%%%%%%%%%%%%% Generate synthetic trace %%%%%%%%%%%%%%%%%%%%%%%%%
%%%%%%%%%%%%%%%%%%%%%%%%%%%%%%%%%%%%%%%%%%%%%%%%%%%%%%%%%%%%%%%%%%%%%%%%%%%

[ X , Y , Z , Trace ] = ...
    ...
Sample_Generator( Lxy               , Lz                  , Length_signal , Number_molecules_sim  , ...
                  wxy               , wz                  , Diffusion     , Delta_t               , ...
                  Molecule_emission , Background_emission , PSF_func);


for m=1:sum(Number_molecules_sim)
    subplot(sum(Number_molecules_sim),1,m)
    stairs(cumsum(Trace),sqrt((X(m,:).^2)+(Y(m,:).^2)+(Z(m,:).^2) ))
    ylim([0 2])
end
%%
%%%%%%%%%%%%%%%%%%%%%%%%%%%%%%%%%%%%%%%%%%%%%%%%%%%%%%%%%%%%%%%%%%%%%%%%%%% Step 3
%%%% Choose the segment of interest from the signal and bin size %%%%%%%%%%
%%%%%%%%%%%%%%%%%%%%%%%%%%%%%%%%%%%%%%%%%%%%%%%%%%%%%%%%%%%%%%%%%%%%%%%%%%%

bin_size = 0.0001    ;             % Initial bin size

minn     = 1         ;             % minimum time step
maxx     = 1000000   ;             % Maximuum time step

% Find the partial trace
mminn=min(find(Trace>=(minn-1)*bin_size));
mmaxx=max(find(Trace<=maxx*bin_size));
edge=cumsum([0,repmat(bin_size,1,floor(Trace(mmaxx)/bin_size)-floor(Trace(mminn)/bin_size))]);
if mminn==1
    sss=0;
else
    sss=Trace(mminn);
end

[trace_binned_partial,~] = histcounts(Trace(mminn:mmaxx)-sss,edge);
plot(trace_binned_partial)

trace_single_partial=Trace(mminn:mmaxx)-sss;
%%
%%%%%%%%%%%%%%%%%%%%%%%%%%%%%%%%%%%%%%%%%%%%%%%%%%%%%%%%%%%%%%%%%%%%%%%%%%% Step 4
%%%%%%%%%%%%%%%%%% Applying addaptive binsize %%%%%%%%%%%%%%%%%%%%%%%%%%%%%
%%%%%%%%%%%%%%%%%%%%%%%%%%%%%%%%%%%%%%%%%%%%%%%%%%%%%%%%%%%%%%%%%%%%%%%%%%%

threshold = 0;                % Adaptive bin size threshold

[Time , trace] = Adaptive_bin(threshold , trace_binned_partial , trace_single_partial , bin_size);


subplot(3,1,1)
plot(trace)

subplot(3,1,2)
plot(Time)

subplot(3,1,3)
plot(trace_binned_partial)

%%
%%%%%%%%%%%%%%%%%%%%%%%%%%%%%%%%%%%%%%%%%%%%%%%%%%%%%%%%%%%%%%%%%%%%%%%%%%% Step 5
%%%%%%%%%%%%%%% Define parameters of the model %%%%%%%%%%%%%%%%%%%%%%%%%%%%
%%%%%%%%%%%%%%%%%%%%%%%%%%%%%%%%%%%%%%%%%%%%%%%%%%%%%%%%%%%%%%%%%%%%%%%%%%%

% Run the Gibbs sampler function based on the number of the iteration.
% There are no automatic MCMC termination for this case (multi-diffusive case) and the user 
% needs to choose a large number of iteration.

save_size         =         10        ;

Number_molecules  =          20        ;     % Maximum number of molecules which will be considered
Max_Species       =          2        ;     % Number of diffusive species

D_alpha_prior     =          1        ;     % Alpha parameter of inverse gamma distr. as the prior of Diffusion coefficient  
D_beta_prior      =          1        ;     % Beta parameter of inverse gamma ditsr. as the prior of Diffusion coefficient

mu_back_proposal  =       1000        ;     % Proposal parameter of molecule photon emission rate
mu_back_alpha     =          2        ;     % Alpha parameter of gamma distr. as the prior of molecule photon emission rate
mu_back_beta      =        400        ;     % Beta parameter of gamma distr. as the prior of molecule photon emission rate

mu_proposal       =       1000        ;     % Proposal parameter of background photon emission rate
mu_alpha          =          2        ;     % Alpha parameter of gamma distr. as the prior of background photon emission rate
mu_beta           =      50000        ;     % Beta parameter of gamma distr. as the prior of background photon emission rate

alpha_q           =          2        ;     % Alpha parameter of hyper prior on the loads
beta_q            =          1        ;     % Beta parameter of hyper prior on the loads

mu_prior_xy       =          0.1      ;     % Mean value for X0 prior in x and y coordintes
mu_prior_z        =          0.1      ;     % Mean value for X0 prior in z coordinte
var_prior_xy      =          2        ;     % Variance for X0 prior in x and y coordintes
var_prior_z       =          2        ;     % Variance for X0 prior in z coordinte
%
%%%%%%%%%%%%%%%%%%%%%%%%%%%%%%%%% Initial vslues %%%%%%%%%%%%%%%%%%%%%%%%%%%%%%%%%%%
D                 =   [5;10]    ;     % Initial Value(s) of Diffusion coefficient
mu                =   [5 ;10]*(10^4)             ;     % Initial Value of molecule photon emission rate
mu_back           =       1000        ;     % Initial value of background photon emission rate

b                 = zeros(Number_molecules*Max_Species,1)                    ;    % Initial value of loads
q                 = zeros(Number_molecules*Max_Species,1)                    ;    % Initial value of weights on loads
x                 = 0.1*randn(3*Max_Species*Number_molecules,length(trace))  ;    % Initial locations of molecules

%
% Define the normalize distances to measure the concentrations (1 = effective volume of FCS)
concen_radious    = [ 0.5 , 1 , 1.5 ] ;

concentration     = zeros(length(concen_radious),length(trace),1) ;    % Set the concentration to zero

PSF_func          = 1      ; % Point spread function type ( 3D Gaussian = 1 , 2D Gaussian Lorentzian = 2 )

%%
%%%%%%%%%%%%%%%%%%%%%%%%%%%%%%%%%%%%%%%%%%%%%%%%%%%%%%%%%%%%%%%%%%%%%%%%%%% Step 6
%%%%%%%%%%%%%%%%%%%%%%%%% Run the algorithm %%%%%%%%%%%%%%%%%%%%%%%%%%%%%%%
%%%%%%%%%%%%%%%%%%%%%%%%%%%%%%%%%%%%%%%%%%%%%%%%%%%%%%%%%%%%%%%%%%%%%%%%%%%

% Since this process is computationaly heavy, we strongly suggest to
% measure the running time with low nuumber of iterations first.


Max_iter          =      1000 ;    % Number of iteration 


[ D                    , mu             , mu_back    , b       , ...
  acceptance_rate_mu   , acceptance_b   , x          , q       , ...
  concentration       ] =  Gibbs_sampler(...
...
...
trace              , Time          , D_alpha_prior     , D_beta_prior     , ...
D                  , mu            , mu_back           , Number_molecules , ...
Max_iter           , wxy           , wz                , mu_proposal      , ...
mu_back_proposal   , mu_alpha      , mu_beta           , mu_back_alpha    , ...
mu_back_beta       , alpha_q       , beta_q            , mu_prior_xy      , ...
mu_prior_z         , var_prior_xy  , var_prior_z       , b                , ...
x                  , q             , concen_radious    , concentration    , ...
Max_Species        , PSF_func      , save_size);
%%
%%%%%%%%%%%%%%%%%%%%%%%%%%%%%%%%%%%%%%%%%%%%%%%%%%%%%%%%%%%%%%%%%%%%%%%%%%%
%%%%%%%%%%%%%%%%%%%%%%%%%%% Show resuults %%%%%%%%%%%%%%%%%%%%%%%%%%%%%%%%%
%%%%%%%%%%%%%%%%%%%%%%%%%%%%%%%%%%%%%%%%%%%%%%%%%%%%%%%%%%%%%%%%%%%%%%%%%%%


%%%%%%%%%%%%%%%%%%%%%%%% Diffusion coefficients %%%%%%%%%%%%%%%%%%%%%%%%%%%
subplot(2,1,1)
for m=1:Max_Species
plot(D(m,:))
hold on
end
hold off
xlabel('Iteration')
ylabel('Diffusion Coef.')

subplot(2,1,2)
dbnd = logspace(-1 , 2 , 100);
% for m=1:Max_Species
%     histogram(D(m,:),dbnd,'Normalization','pdf')
%     hold on
% end
histogram(D,dbnd,'Normalization','pdf')

hold off
set(gca,'XScale','log')
xlabel('Diffusion coefficient (\mum^2/s)')
ylabel('Post. prob. distr.')
hold on
plot(dbnd,((dbnd.^(-D_alpha_prior-1)).*exp(-(D_beta_prior./dbnd)))/...
    ((D_beta_prior.^D_alpha_prior)./gamma(D_alpha_prior)),'linewidth',2,'color','c')
%%
%%%%%%%%%%% molecule and background photon emission rates %%%%%%%%%%%%%%%%%
subplot(2,2,[1,2])
for m=1:Max_Species
plot(mu(m,:))
hold on
end
plot(mu_back,'r') 
hold off
xlabel('Iteration')
ylabel('Photon emision rates (photons/s)')
set(gca,'YScale','log')

subplot(2,2,3)
dbnd = logspace(1 , 5 , 100);
histogram(mu_back,dbnd,'Normalization','pdf','facecolor','r')
set(gca,'XScale','log')
xlabel('Molecule photon emission rates (photons/s)')
ylabel('Post. prob. distr.')
hold on
plot(dbnd,((dbnd.^(mu_back_alpha-1)).*exp(-(dbnd./mu_back_beta)))/...
    ((mu_back_beta.^mu_back_alpha)*gamma(mu_back_alpha)),'linewidth',2,'color','c')

subplot(2,2,4)
dbnd = logspace(2 , 6 , 100);
% for m=1:Max_Species
%     histogram(mu(m,:),dbnd,'Normalization','pdf')
%     hold on
% end
histogram(mu,dbnd,'Normalization','pdf')
hold on
set(gca,'XScale','log')
xlabel('Background photon emission rate (photons/s)')
ylabel('Post. prob. distr.')
plot(dbnd,((dbnd.^(mu_alpha-1)).*exp(-(dbnd./mu_beta)))/...
    ((mu_beta.^mu_alpha)*gamma(mu_alpha)),'linewidth',2,'color','c')
%%
%%%%%%%%%%%%%%%%% Number of active molecules (loads) %%%%%%%%%%%%%%%%%%%%%%
subplot(2,1,1)
for m=1:Max_Species
    plot(sum(b((m-1)*Number_molecules+1:m*Number_molecules,:),1))
    hold on
end
xlabel('Iteration')
ylabel('Number of active molecules')

subplot(2,1,2)
for m=1:Max_Species
histogram(sum(b((m-1)*Number_molecules+1:m*Number_molecules,:),1),'Normalization','pdf')
hold on
end
% histogram(sum(b(:,:),1),'Normalization','pdf')
ylabel('Post. prob. distr.')
xlabel('Number of active molecules')
%%
%%%%%%%%%%%%%%%%%%%%%%%%%%%% Concentration(s) %%%%%%%%%%%%%%%%%%%%%%%%%%%%%

% Pre-calculations
sign_siz  = length(Trace);
radi_size = length(concen_radious);
concentration_real = zeros(length(concen_radious),sign_siz);

% calculate the real concentrations
for num_spe = 1:length(Number_molecules_sim)

    for num_mol = 1:Number_molecules_sim(num_spe)
        
        nump=(num_spe-1)*sum(Number_molecules_sim(1:num_spe-1))+num_mol;
        
        for num_radi = 1:radi_size
         
            concentration_real(num_radi,:) = concentration_real(num_radi,:)+...
                                            ( sqrt(((X(nump,:)./wxy).^2)+...
                                                   ((Y(nump,:)./wxy).^2)+...
                                                   ((Z(nump,:)./wz ).^2) )<=concen_radious(num_radi) );
                             
        end
    
    end
end

% Cacluate the learned concentrations
mean25   =  [];
mean50   =  [];
mean95   =  [];

for k=1:size(concentration,2)
    for l = 1 : radi_size
            mean25(l,k)=quantile(concentration(l,k,:),0.025)      ;
            mean50(l,k)=quantile(concentration(l,k,:),0.5)     ;
            mean95(l,k)=quantile(concentration(l,k,:),0.975)     ;
   end
end


mean25_stairs=[];
mean50_stairs=[];
mean95_stairs=[];
timee=[];
for i=1:length(Time)
    mean25_stairs(:,2*i-1)=mean25(:,i);
    mean50_stairs(:,2*i-1)=mean50(:,i);
    mean95_stairs(:,2*i-1)=mean95(:,i);
    
    if  i<length(Time)
        mean25_stairs(:,2*i)=mean25(:,i+1);
        mean50_stairs(:,2*i)=mean50(:,i+1);
        mean95_stairs(:,2*i)=mean95(:,i+1);
    else
        mean25_stairs(:,2*i)=mean25(:,i);
        mean50_stairs(:,2*i)=mean50(:,i);
        mean95_stairs(:,2*i)=mean95(:,i);
    end
    
    timee(2*i-1)=Time(i);
    timee(2*i)=0;
end


for l = 1 : length(concen_radious)
    
    subplot(length(concen_radious)+1,1,l)
    hold on
    stairs(Trace,concentration_real(l,:),'g');
    stairs(cumsum(timee),mean50_stairs(l,:),'color','b')
    
    x2 = [cumsum(timee), fliplr(cumsum(timee))];
    inBetween = [mean25_stairs(l,:), fliplr(mean95_stairs(l,:))];
    h1=fill(x2, inBetween, 'm','facealpha', 0.2,'edgecolor', 'none');
    hold on
    
    xlabel('Time step (ms)')
    ylabel('concentration')
    box off
    legend('Estimated value of concentration')
end

subplot(length(concen_radious)+1,2,[2*length(concen_radious)+1,2*length(concen_radious)+2])
plot(trace_binned_partial)

xlabel(['Time step (',num2str(bin_size),'\mus)'])
ylabel('Observed photons')


%%
%%%%%%%%%%%%%%%%%%% Joint probablity distributions %%%%%%%%%%%%%%%%%%%%%%%%
Results=[];

for m=1:Max_Species
    Titles{m}=['D',num2str(m)];
    Results(m,:)=D(m,:);
end

Titles{Max_Species+1}='\mu_{mol}';
Titles{Max_Species+2}='\mu_{back}';

Results(3:4,:)=[mu;mu_back];
Results=Results';

numParams=size(Results,2);

for i=1:numParams
    for j=i:numParams
        subplot(numParams,numParams,numParams*(i-1)+i+(j-i));
        if i~=j
            scatter(Results(:,i),Results(:,j),'SizeData',1,'Marker','.');
            title([Titles{i},' , ',Titles{j}])
            ylabel(Titles{j})
            xlabel(Titles{i})
            zlabel('Joint Posterior')
            hold on
        end
        if i==j
           histogram(Results(:,j),'Normalization','pdf')
           ylabel('Post. prob. distr.')
           xlabel(Titles{i})
           box off
           hold on 
        end
    end
end
%%
%%%%%%%%%%%%%%%%%%%%%%%%% Save results %%%%%%%%%%%%%%%%%%%%%%%%%%%%%%%%%%%%
save('Results',...
    ...
    'D','mu','mu_back','b','q','trace','Trace',...
    'wxy','wz','mu_alpha','mu_beta','mu_proposal',...
    'mu_back_alpha','mu_back_beta','mu_back_proposal','mu_prior_xy',...
    'mu_prior_z','D_alpha_prior','D_beta_prior','acceptance_rate_mu',...
    'acceptance_b','x','Time','Signal','trace_single_partial','minn','maxx','trace_binned_partial')
```
